# deSALT: fast and accurate long transcriptomic read alignment with de Bruijn graph-based index

**DOI:** 10.1101/612176

**Authors:** Bo Liu, Yadong Liu, Junyi Li, Hongzhe Guo, Tianyi Zang, Yadong Wang

## Abstract

Long-read RNA sequencing (RNA-seq) is promising to transcriptomics studies, however, the alignment of long RNA-seq reads is still non-trivial due to high sequencing errors and complicated gene structures. Herein, we propose deSALT, a tailored two-pass alignment approach, which constructs graph-based alignment skeletons to infer exons and uses them to generate spliced reference sequences to produce refined alignments. deSALT addresses several difficult technical issues, such as small exons and sequencing errors, which breakthroughs the bottlenecks of long RNA-seq read alignment. Benchmarks demonstrate that deSALT has a greater ability to produce accurate and homogeneous full-length alignments. deSALT is available at: https://github.com/hitbc/deSALT.

## Background

RNA sequencing (RNA-seq) has become a fundamental approach to characterize transcriptomes. It reveals precise gene structures and quantifies gene/transcript expressions [1-5] in various applications, such as variant calling [6], RNA editing analysis [7, 8], and gene fusion detection [9, 10]. However, current widely used short read sequencing technologies have limited read length and systematic bias from library preparation. These drawbacks limit more accurate alignment [11] and precise gene isoform analysis [12], thus creating a bottleneck for transcriptomic studies.

Two kinds of long read sequencing technologies, i.e., single molecule real time (SMRT) sequencing produced by Pacific Biosciences (PacBio) [13] and nanopore sequencing produced by Oxford Nanopore Technologies (ONT) [14], are emerging and promising to breakthrough the bottleneck of short reads in transcriptomic analysis. Both of them enable the production of much longer reads, the mean and maximum lengths of the reads being over ten to hundreds of thousands of base pairs (bp) [15, 16], respectively. Taking this advantage, full-length transcripts can be sequenced by single reads, which is promising for substantially improving the accuracy of gene isoform reconstruction. Furthermore, there is less systematic bias in the sequencing procedure [17], which is also beneficial to gene/transcript expression quantification.

Besides their advantages, PacBio and ONT reads have much higher sequencing error rates than that of short reads. For PacBio SMRT sequencing, the sequencing error rate of raw reads (“subreads”) is about 10% to 20% [16]; for ONT nanopore sequencing, the sequencing error rates of 1D and 2D (also known as 1D2) reads are about 25% and 12% [18, 19], respectively. PacBio SMRT platforms can produce reads of inserts (ROIs) by sequencing circular fragments multiple times to largely reduce sequencing errors. However, this technology has lower sequencing yields and reduced read lengths. Therefore, these high sequencing errors raise new technical challenges for RNA-seq data analysis. Read alignment could be the most affected one, and the effect may not be limited to the read alignment itself since it is fundamental to many downstream analyses.

Previous studies [20-22] have demonstrated that noisy DNA-seq long read alignment is a non-trivial task. Many technical issues, such as the high sequencing errors, potential genome variants, and large read lengths, need to be handled well. For RNA-seq long read alignment, the task is even more difficult since the aligner has to deal with numerous splicing events besides the issues mentioned above. This requires the aligner to have a strong ability to implement a highly complicated split alignment (also called “spliced alignment”) to correctly recognize many splicing junctions and map the bases to the corresponding exons. Although most of the proposed DNA-seq long read alignment approaches have the ability to implement split alignment to handle genome structure variations (SVs) [21-23], tailored algorithms are still in high demand because splicing junctions occur more frequently and the lengths of exons are much shorter and divergent.

There have been several approaches supporting RNA-seq long read alignment, such as BBMap [24], GMAP [25], STAR [26], BLAT [27], GraphMap2 [28] and Minimap2 [29]. All of these approaches are based on the commonly used seed-and-extension strategy, by which various seeding and extension methods are implemented to address the technical issues. They all have the ability to handle splicing junctions. However, most of them have relatively slow speed [28, 29], mainly because of the numerous short matches in the seeding step and the time-consuming local alignment in the extension step. Moreover, some of the algorithms have lower sensitivity [30], i.e., many reads are unaligned or only partially aligned, potentially due to their relatively poor ability to handle sequencing errors. An outstanding algorithm is Minimap2, which simultaneously achieves tens of times faster speed and similar or higher sensitivity than other state-of-the-art aligners. This algorithm mainly benefits from its well-designed minimizer-based indexing [31] and SSE-based local alignment methods [32], which greatly improve the efficiency of the seeding and extension steps. Furthermore, its specifically designed local extension method is suited to handling splicing junctions.

In absolute terms, the ultimate goal of the task is to map all the bases for all the reads correctly. However, this could still be non-trivial to state-of-the-art aligners in several aspects. One problem is the alignment of the bases from relatively short exons (e.g., exons having only a few tens of bp). It is extremely hard to find seeds in the read parts from such short exons under the circumstances of high sequencing errors and potential variants, so that the read parts are usually unaligned or mistakenly aligned. Another issue is that it is difficult to align the bases near the splicing junctions correctly. This problem also exists in short RNA-seq read alignment; however, it is more serious in the alignment of noisy long RNA-seq reads. Moreover, with the effect of sequencing errors, the alignments of the reads from the same gene isoform are usually divergent from each other, which is also misleading downstream analysis.

Herein, we propose the de Bruijn graph-based spliced aligner for long transcriptome reads (deSALT). deSALT is a fast and accurate RNA-seq long read alignment approach which takes the advantages of a novel two-pass read alignment strategy based on the de Bruijn graph-based index. It has a strong ability to handle complicated gene structures and high sequencing errors to produce sensitive, accurate and homogeneous alignments. For most of the reads, deSALT can produce full-length alignments to recover the exons and splicing junctions thoroughly along the entire reads. Moreover, the speed of deSALT is also faster than or comparable to state-of-the-art approaches. We believe that it has the potential to play an important role in many forthcoming transcriptomic studies.

## Results

### Overview of the deSALT approach

The seed-and-extension approach is suited to spliced alignment since it is able to match the short tokens of the read to its spanning exons first (i.e., seeding) and then implement base-level alignment between the read and the matched exons (i.e., extension). However, under the circumstances of frequent splicing events and high sequencing errors, this task is non-trivial in practice. For a single read, it is usually difficult to find matches between the read and all of its spanning exons accurately, especially for the read parts with high sequencing errors and relatively short exons. Since it is hard to compose a local reference sequence containing all the spanning exons of the read, the extension alignment would be less accurate, or some of the read parts could be unaligned or clipped. Moreover, due to the randomness of the sequencing errors, various mistakes could be made in the seeding and extension phases in regard to multiple reads from the same gene isoform, and the produced alignments of the reads could be divergent from each other.

Motivated by these technical problems and existing short RNA-seq read alignment algorithms [26, 33], deSALT uses a two-pass approach to align the noisy long reads (a schematic illustration is in Figure 1). In the first pass, it employs a graph-based genome index [34] to find match blocks (MBs) between the read and the reference and uses a sparse dynamic programming (SDP) approach to compose the MBs into alignment skeletons (referred to as the “alignment skeleton generation” step). All the alignment skeletons of all the reads are then integrated to comprehensively detect the exon regions (referred to as the “exon inference” step). In the second pass, deSALT relocates the short matches between the read and the detected exons to compose a local spliced reference sequence (LSRS), which is expected to be a concatenation of all the spanning exons of the read. The read is aligned against the LSRS to produce a refined base-level alignment (referred to as the “refined alignment” step).

**Figure 1.**
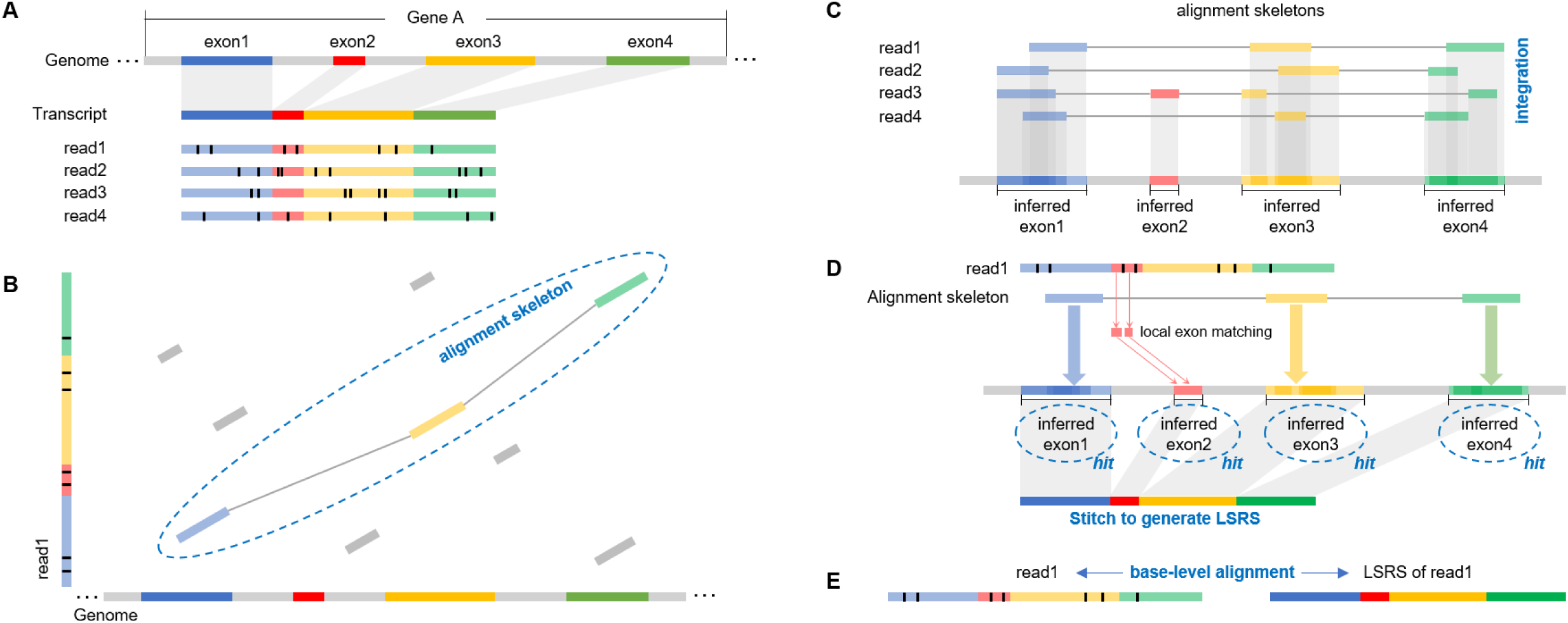
A schematic illustration of the deSALT approach. There is a gene (termed “Gene A”) with four exons (respectively marked by the colors blue, red, yellow and green; introns are marked by grey color) and four reads that all sequence throughout the whole transcript (assuming that Gene A has only one isoform). Moreover, each of the reads has some sequencing errors (marked by the short black bars in the reads). **(B)** Alignment skeleton generation (first-pass alignment): for each of the reads (read1 is employed as an example), deSALT finds the MBs between it and the reference genome (marked as colored bars) and connects them to build an optimized alignment skeleton using an SDP approach. **(C)** Exon inference: deSALT integrates all of the generated alignment skeletons by mapping their involved MBs to the reference genome. The projections of the MBs are analyzed to infer exon regions in the reference genome. **(D-E)** Refined alignment (second-pass alignment): for each of the reads, deSALT finds additional local matches on the exons between or near the exons involved in the alignment skeleton. Further, it recognizes all the inferred exons related to the alignment skeleton or the newly found local matches as “hit exons” and stitches all of them to generate an LSRS. (The figure shows that there are two newly found matches on exon 2, and they help to recuse this exon to build a correct LSRS.) The read is then aligned with the LSRS to produce a refined alignment.

The key point of deSALT is its comprehensive analysis of the local matches of all the reads through alignment skeletons. Since the sequencing errors are random [17], each of the alignment skeletons contains some distinct as well as some complementary information about exon regions, which look like puzzle pieces, and the integration of alignment skeletons can effectively filter the sequencing errors to implement a sensitive and noise-robust detection of exons. In the later step, the detected exons help to narrow down the searching space to find additional short matches which cannot be detected by the relatively longer seeds used in the initial step. With these local matches, deSALT enables the effective inference of all the spanning exons of a given read and the composition of a high-quality spliced reference sequence to produce accurate full-length alignment. This approach is robust to very short exons, frequent splicing events, potential small variants, as well as sequencing errors. Furthermore, deSALT generates homogeneous LSRSs for the reads from the same gene isoform with the integrated information, which enables the production of more homogeneous alignments.

deSALT has fast speed with its tailored design. Unlike conventional two-pass alignment approaches [26, 33] where both of the two passes produce base-level alignment, deSALT only uses pseudo-alignment (alignment skeleton) in the first pass, the operation of which is similar to the seeding process. Thus, the whole process is like a one-pass alignment plus a fast integration of the alignment skeletons. Moreover, some optimized implementations, for example, graph-index-based skeleton generation and SIMD-based local alignment [29, 32], also help to accelerate the speed.

### Results on simulated datasets

We simulated 60 long RNA-seq read datasets with 6 simulation models, respectively termed as “PacBio ROI reads”, “PacBio subreads”, “ONT 2D reads”, “ONT 1D reads”, “PS-ONT reads” and “NS-ONT reads”, which have specific sequencing error profiles and read lengths to mimic various sequencing platforms. The error models were configured according to previous studies [17, 35, 36]. More precisely, “PacBio ROI reads” (error rate: 2%) and “PacBio subreads” (error rate: 15%) models were respectively generated by PBSim [37] with fixed parameters to mimic PacBio ROIs and subreads; “ONT 2D reads” (error rate: 12%) and “ONT 1D reads” (error rate: 25%) models were respectively generated by PBSim with fixed parameter to mimic ONT 2D and ONT 1D reads; and “PS-ONT reads” and “NS-ONT reads” were respectively generated by PBSim and NanoSim [36] based on a real ONT dataset (SRA Accession Number: SRR2848544) to simulate more realistic ONT datasets. Refer to Supplementary Table 1 and Supplementary Notes for the used command lines and parameters of the simulators.

For each model, there were 10 simulated datasets from three species based on Ensembl gene annotations [38] (human: GRCh38, version 94; mouse: GRCm38, version 94; fruit fly: BDGP6, version 94). One was generated by all coding genes of human to mimic the variable isoform level expressions. In details, for each of the genes with multiple isoforms, one of the isoforms was randomly selected as “highly expressed” and the other ones were selected as “lowly expressed”. And for the genes with single isoforms, all their isoforms were selected as “highly expressed”. Then high (30X) and low (4X) coverage simulations were implemented for the highly and the lowly expressed isoforms, respectively, and the simulated reads were mixed to build the dataset. To benchmark the aligners on the datasets from various species and in various coverages more comprehensively, we referred to a previous study [30] to randomly select three sets of genes respectively from the three species to simulate 9 datasets. Each of the datasets was built with one of the gene-sets and a fixed coverage (4X, 10X or 30X). Also refer to Methods section for more details about the implementation of the benchmarking.

deSALT and three state-of-the-art approaches, Minimap2, GraphMap2 and GMAP, were applied to all 60 datasets for comparison. In the benchmark, the indexes of all the aligners were pre-built. Five metrics (i.e., Base%, Exon%, Read80%, Read100%, and #Bases/m) were used to assess the sensitivity, accuracy and performance of the aligners (refer to the Methods section for the definitions). The results of the simulated datasets are provided in Figure 2 and Supplementary Tables 2-8. Mainly, five observations were made.

**Figure 2.**
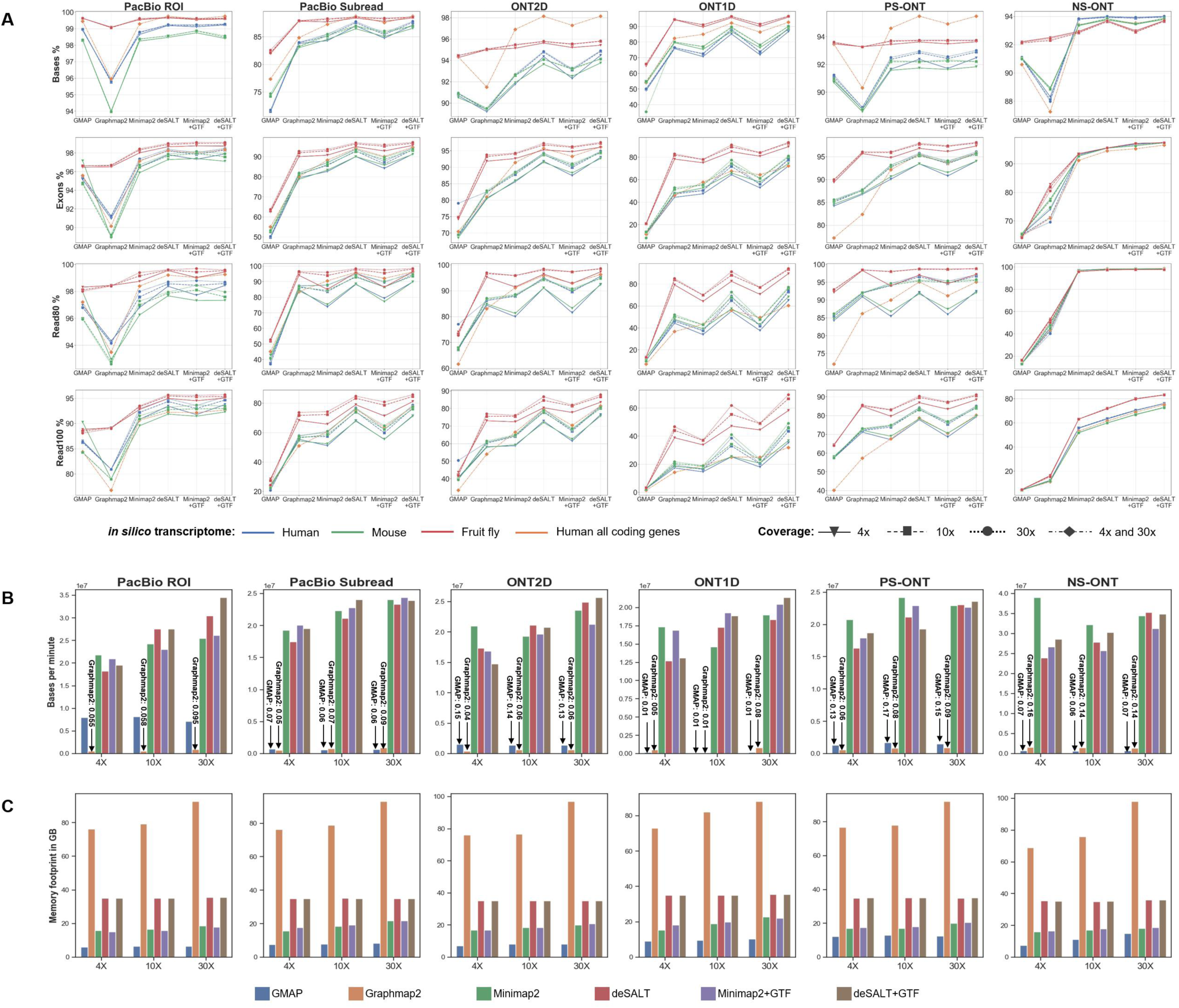
Results on simulated datasets. The figure depicts the yields (A), speed (B) and memory footprints (C) of the aligners on the simulated datasets. It is worth noting that for deSALT and Minimap2, both the results with and without gene annotations are shown (indicated as “deSALT+GTF”, “deSALT”, “Minimap2”, “Minimap2+GTF”, respectively). **(A)** Each of the subplots indicates one of the four metrics (Base%, Exon%, Read80%, and Read100%, respectively) of the aligners on the datasets in a specific sequencing model (PacBio ROI, PacBio subreads, ONT2D, ONT1D, PS-ONT and NS-ONT, respectively). In each subplot, the blue, green and red lines respectively correspond to the results of the datasets from randomly selected genes of human, mouse and fruit fly, and the orange lines correspond to the results of the dataset from all the protein coding genes of human. Moreover, the shapes (reversed triangles, rectangles, circles and rhombuses) indicate the datasets in various sequencing coverages. In **(B)** and **(C)**, each of the subplots indicates the speed (#Base/m) and the memory footprint (in GB) of the aligners (in 8 CPU threads) on the datasets simulated from human (randomly selected genes) with a specific sequencing model. The datasets in various sequencing coverages are shown separately, and the bars in different colors refer to various aligners. Also refer to Supplementary Table 8 for a more comprehensive assessment of the alignment speeds and memory footprints in various numbers of CPU threads (1, 4, 8, and 16 CPU threads).

1. deSALT has outstanding alignment yields. The results of the aligners without gene annotations (the first four columns of all the subplots of Figure 2) indicate that deSALT had higher or comparable base% statistics on all the datasets (the results with gene annotations are described in the fifth observation below). In particular, the outperformance of deSALT was more obvious on the datasets having medium and high error rates, and this is preferable for handling real noisy long reads. Moreover, deSALT showed its advantages in the exon% statistics, suggesting that it has a stronger ability to recover the exons and splicing junctions within the reads. deSALT also outperformed other state-of-the-art aligners on the Read80% and Read100% statistics, indicating that it is good at producing full-length alignments. In particular, the highest Read100% statistics of deSALT suggests that, for more reads, it can correctly align all of their exons without introducing false positives. It is also worth noting that both of GraphMap2 and Minimap2 outperformed GMAP. Moreover, GraphMap2 and Minimap2 outperformed each other on various datasets and various metrics, and they overall had comparable yields. In absolute terms, deSALT correctly aligned most of the bases as well as the exons for the reads with low and medium error rates. But for the error-prone ONT 1D reads (error rate: 25%), the exon% statistics of deSALT is about 64%–90%, lower than that of deSALT on other simulated datasets. This indicates that deSALT was to some extent affected by the high sequencing errors. It is observed that the most affected datasets are the low coverage (4X) mammalian (human and mouse) datasets. This was mainly due to the fact that the bases near exon boundaries are very difficult to confidently align under the circumstance of serious noise. However, the yield on these error-prone reads improved with the increase in read depth. It is a feature of the two-pass approach that the effect of sequencing errors can be better mitigated and the detection of exons can be improved with more available reads, and all the reads share this profit to compose more sensitive alignments. Furthermore, deSALT has the ability to produce not only accurate, but also homogeneous alignments. This is also an advantage of the two-pass alignment, that deSALT tends to compose homogeneous LSRSs for the reads from same gene isoforms, which helps to align them to the correct positions simultaneously. However, one-pass approaches are more easily affected by sequencing errors and other factors, such as very small exons and frequent splicing events, which usually produce more heterogeneous alignments potentially with more mistakes. Figure 3 shows a typical example describing the characteristics of deSALT. Besides the yields, deSALT also has good performance. Its speed is similar to that of Minimap2 and tens of times faster than that of GraphMap2 and GMAP (Figure 2B and Supplementary Table 8). This speed is suited to large-scale datasets. Moreover, the aligners have different memory footprints (Figure 2C and Supplementary Table 8). The memory footprints of deSALT is higher than that of Minimap2 and GMAP, since the RdBG-index of deSALT uses a hash table-based data structure to index all the k-mers of RdBG, which needs larger RAM space. However, the memory use of deSALT is still acceptable to most of modern servers (e.g., about 35GB for human). It is observed that the memory footprints of GraphMap2 increased with the number of input reads, and higher than that of all the other benchmarked aligners.
2. deSALT has good ability to align the reads spanning short exons. We specifically assessed the alignment of the reads spanning short exons (exons < 31 bp), and the results (Supplementary Table 3) suggest that deSALT outperformed other aligners. This is derived from the two-pass approach that short exons can be better detected with the generation and integration of alignment skeletons, and the discovered exons are fully considered by the shorter local matches used in the second pass. Thus, high-quality LSRSs can be composed and help the alignment of the read parts spanning short exons. However, other state-of-the-art aligners that use a one-pass strategy are more likely to be affected by splicing events and sequencing errors. This results in reduced ability to find local matches on short exons and some of the read parts from short exons are mistakenly aligned. Two examples are provided in Figure 3B and Supplementary Figure 1.
3. deSALT has good ability to handle multiple splicing events. We assessed the alignment of the reads from the transcripts with various number of exons (2–5 exons, 6–9 exons, and >9 exons). deSALT can produce equally good alignments for all the three read groups (Supplementary Table 4), indicating that it enables the handling of numerous splicing events within the reads (an example is provided in Supplementary Figure 2). Minimap2 showed a similar trend, but its Read80% and Read100% statistics were lower. GMAP showed significant decreases in the Read80% and Read100% statistics as the number of exons increased, indicating that it might not be good at handling reads with many splicing events. GraphMap2 also showed such decreases, but not as significant as that of GMAP.
4. deSALT has good ability to handle genes with multiple isoforms. We separately assessed the alignments of the reads from the genes with single and multiple isoforms (Supplementary Table 5), and found that overall deSALT has equally good yields for both of the two categories of reads, indicating that it has the ability to handle genes with various numbers of isoforms. An example of deSALT to align the reads from a gene with many isoforms is in Supplementary Figure 3. To further investigate the ability of the aligners on the reads from alternative splicing genes, we assessed the alignments of the reads from “highly expressed isoforms” and “lowly expressed isoforms” of all human coding genes datasets separately (Supplementary Table 6). The yields of deSALT were still better than those of other aligners, and the difference between the reads from highly- and lowly expressed isoforms was overall quite small. However, for error-prone ONT 1D reads, the difference between lowly- and highly expressed isoforms was larger, e.g., there was a 10% decrease for the exon% statistics (Supplementary Figure 4). We investigated some intermediate results of deSALT, and found that the decreased statistics are not due to a poor ability of handling alternative splicing genes, but the low coverage of the datasets, which is similar to that of the low coverage ONT 1D datasets. That is, there were only a small number of reads being from lowly expressed isoforms, and they were not enough to mitigate the sequencing noise with their own alignment skeletons. Thus, some difficult and isoform-specific cases could not be well handled, e.g., some read parts from isoform-specific short exons. To further justify this issue, we used the ONT 1D reads from lowly expressed isoforms as an independent dataset and asked deSALT to run it. Similar results were obtained (the red bars in Supplementary Figure 4), and this suggested that deSALT was not affected by the variable expression levels of the isoforms. Moreover, there is also a category of special cases that pairs of genes overlap in the genome but are on opposite strands. The alignment of the reads from such genes are to some extent similar to that of the reads from alternative splicing genes. We implemented an assessment on the alignments of these reads (Supplementary Table 7), and found that there was no significant difference to that of deSALT’s alignment results on the reads from non-overlapped genes (Supplementary Figure 5), indicating that deSALT can also produce accurate alignments for such reads. An example is in Supplementary Figure 6.
5. deSALT can further improve the alignment of error-prone reads with gene annotations. We used Ensembl gene annotations as input to benchmark the alignments of deSALT and Minimap2. The results (Figure 2A and Supplementary Table 2) demonstrate that both of deSALT and Minimap2 had better yields with the help of gene annotations, especially for the alignment of the error-prone ONT 1D reads. Moreover, the difference between the yields of the two aligners is also smaller. For deSALT, the improvement comes from that gene annotations help to rescue missed exons. In details, deSALT usually finds only a few matches on very noisy reads to build incomplete alignment skeletons which lower the sensitivity of exon detection. In this situation, gene annotations supply additional information to find matches for those read parts from missed exon regions. This solves many error-prone read parts (an example is provided in Supplementary Figure 7) and helps to produce full-length alignments (see the gains in the Read80% and Read100% statistics).

**Figure 3.**
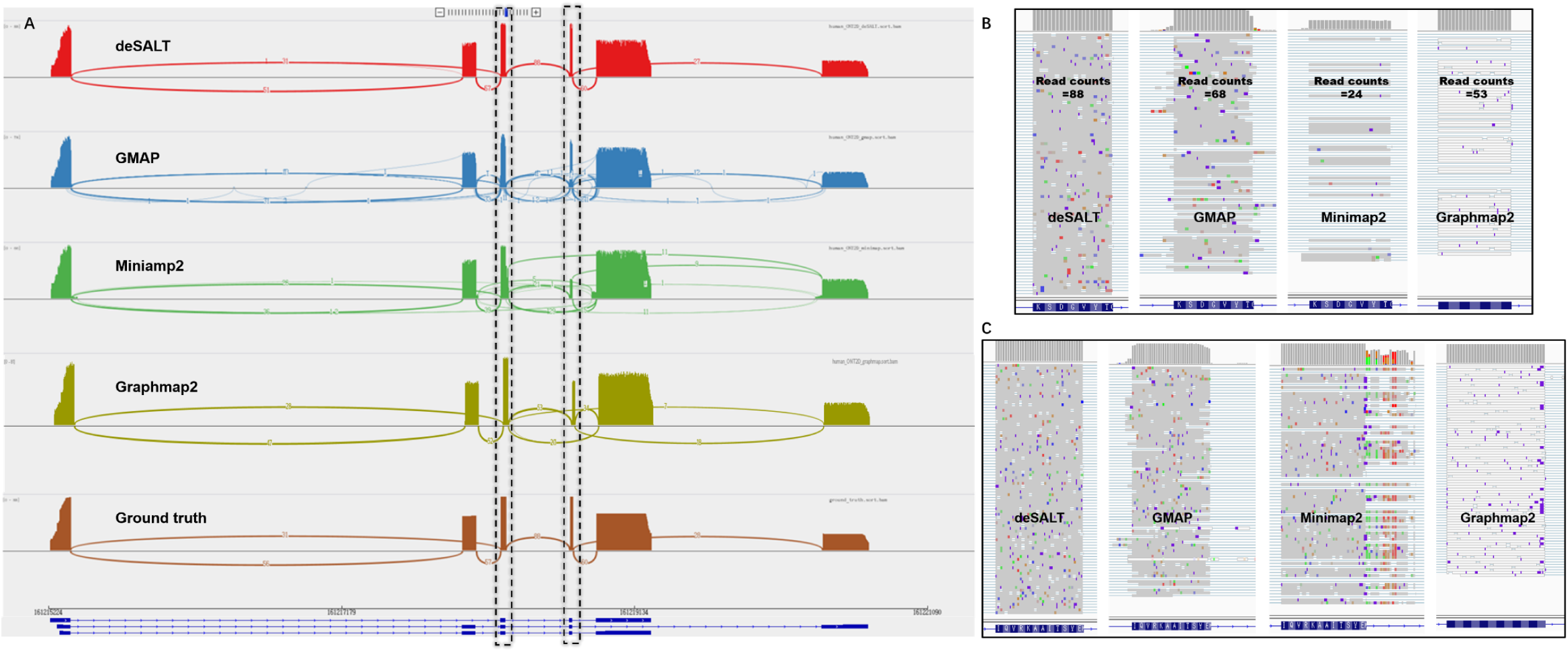
An example of the alignments of simulated reads by various aligners. This figure represents the snapshots of the alignments of the reads from the simulated 30X ONT 2D human dataset, around FCER1G gene (Chr1: 161215297–161219248) of GRCh38. FCER1G gene has 6 exons and 3 isoforms. According to the ground truth, there are 88 reads in this region. The numbers of Read100 and Read80 reads of deSALT are 84 and 88, respectively, higher than those of GMAP (#Read100: 45 and #Read80: 54), Minimap2 (#Read100: 19 and #Read80: 23) and Graphmap2 (#Read100: 50 and #Read80: 78). **(A)** The Sashimi plots represent the overall views of the alignments. Compared to the ground truth (the bottom track), it is observed that the deSALT alignments are more homogenous, i.e., at each splicing site, most of the reads have similar breakpoints, which also coincide with the ground truth. The more heterogeneous alignments of GMAP, Minimap2 and Graphmap2 are usually due to some less accurate alignments at small exons and exon boundaries. **(B)** A detailed view at the fourth exon of the FCER1G gene (length: 21 bp). deSALT correctly aligns all of the 88 reads spanning this exon; however, the corresponding numbers for GMAP (68), Minimap2 (24) and Graphmap2 (53) are lower. **(C)** A detailed view at the third exon of the FCER1G gene (length: 36 bp). It is observed that the reads have nearly the same breakpoints with the homogeneous alignments of deSALT. However, for the other three aligners, the breakpoints of the reads are more divergent, and some of them are less accurate, which could be due to the effect of sequencing errors as well as to the nearby small exons.

Overall, the simulation results demonstrate that deSALT is able to achieve high sensitivity, accuracy and performance simultaneously. Especially, it has the ability to address many difficult issues, such as sequencing errors, short exons, frequent splicing events, multiple isoforms and so on, which is useful to breakthrough the bottlenecks of long RNA-seq read alignment.

### Results on real sequencing datasets

We assessed the aligners with three real sequencing datasets. The first two datasets are from a well-studied CEPH sample (NA12878), and respectively produced by ONT cDNA sequencing (containing 15,152,101 reads and 14,134,831,170 bases in total) and ONT direct RNA sequencing (containing 10,302,647 reads and 10,614,186,428 bases in total). The two datasets are available at https://github.com/nanopore-wgs-consortium/NA12878. The third dataset is from a mouse sample produced by the PacBio platform [39] (SRA Accession Number: SRR6238555; containing 2,269,795 reads and 3,213,849,871 bases in total).

We used a series of metrics based on gene annotations to evaluate the alignments (i.e., #BaseA, #BaseGA, #ExonP, #ExonGO, #ExonGA, #ExonGA(x), #ReadGA) due to a lack of ground truth (refer to Methods section for definitions). Ensembl gene annotations (human: GRCh38, version 94 and mouse: GRCm38, version 94) were employed for the assessment. It is also worth noting that we only showed the results of GraphMap2 on the ONT cDNA dataset, since it unaligned most of the reads of the ONT direct RNA dataset and raised a “segmentation fault” for the PacBio dataset during benchmarking. The results are provided in Figure 4 and Supplementary Tables 9 and 10. Four observations were made as follows.

**Figure 4.**
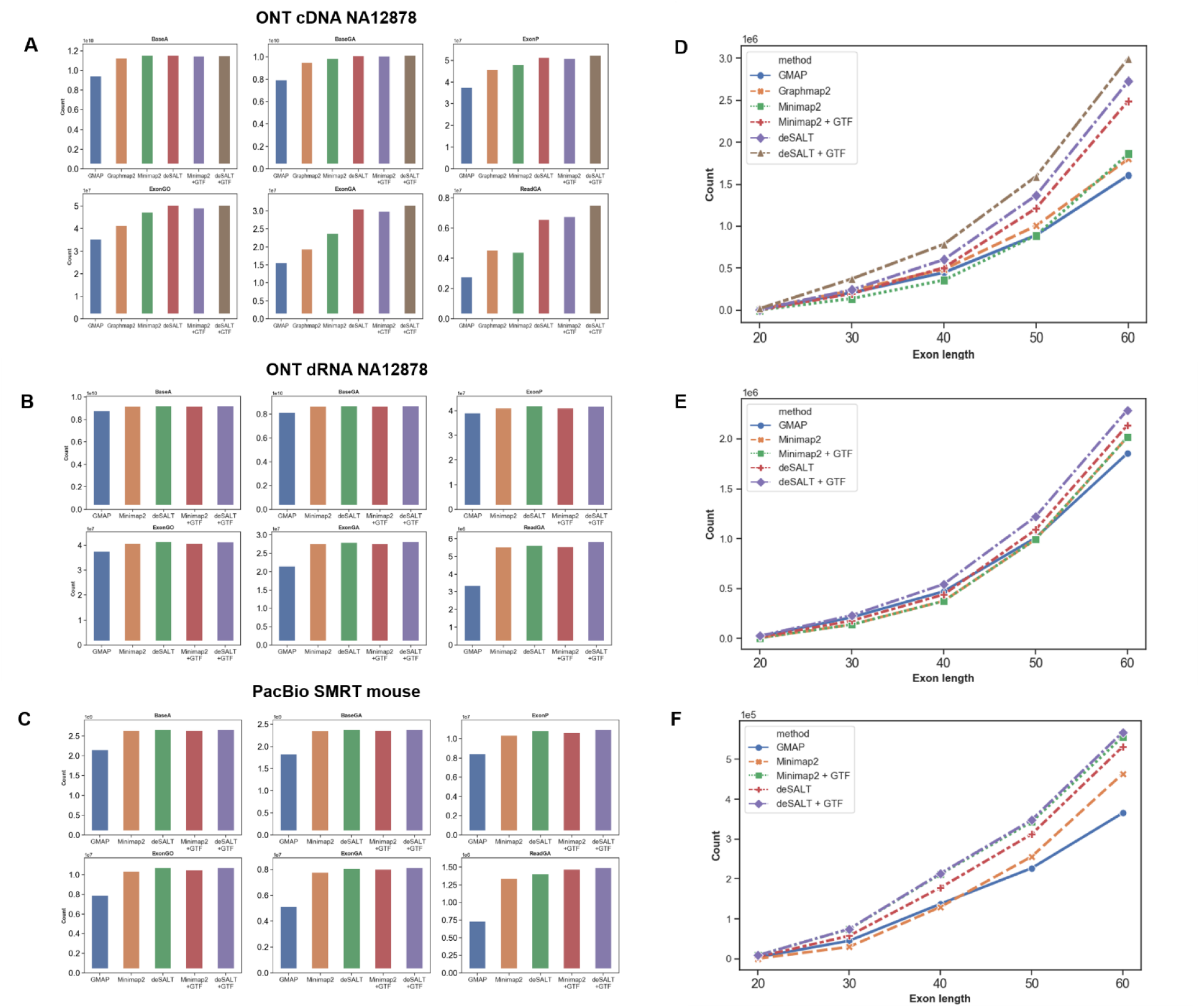
Results on real datasets. The figure depicts the yields of the aligners and the sensitivity of the aligners on short exons. **(A-C)** The six metrics (BaseA, BaseGA, ExonP, ExonGO, ExonGA, and ReadGA) of the aligners on the human ONT cDNA (A) and direct RNA datasets, and the mouse PacBio dataset (C). Each bar in a subplot indicates the result of a specific aligner. **(D-F)** The ExonGA(x) metrics, i.e., ExonGA(20), ExonGA(30), ExonGA(40), ExonGA(50), and ExonGA(60), of the aligners on the human ONT cDNA (A) and direct RNA (B) datasets, and the mouse PacBio dataset (C), which depicts the sensitivities of the aligners for relatively short (up to 60 bp) exons.

1. deSALT still has the best alignment yields. For the three real datasets, deSALT achieved the highest #BaseGA statistics (i.e., it aligned most bases to the annotated exon regions). Moreover, deSALT also had the highest numbers of predicted exons being overlapped by (#ExonGO) and exactly matched to (#ExonGA) annotated exons. These statistics indicate that deSALT achieves good sensitivity. Furthermore, deSALT had the highest #ReadGA statistics, indicating that it has better ability to produce correct full-length read alignments. The time costs with 24 threads were also assessed (both wall clock time and CPU time, Supplementary Table 9), and it suggests that deSALT and Minimap2 has similar speed and both of them are faster than GMAP and GraphMap2. It was also observed that the #BaseGA of Minimap2 was close to that of deSALT, indicating that the two approaches have similar alignment yields overall. However, deSALT outperformed Minimap2 on #ExonGO, #ExonGA, and #ReadGA statistics. We investigated the detailed alignment results and found that, similar to that of the simulated reads, this derives from deSALT’s ability to handle short exons and produce homogeneous alignments (see below for details). A typical example of the alignment of real sequencing reads is in Figure 5.
2. deSALT enables to handle relatively short exons. We assessed the alignment of the bases putatively from short exons by a series of #ExonGA(x) statistics (Figures 4D-F), i.e., ExonGA(20), ExonGA(30), ExonGA(40), ExonGA(50) and ExonGA(60). The results demonstrate that deSALT enables the recovery of a higher number of short exons. It is worth noting that although only a small proportion of exons are short, they are important to the study of gene splicing, and so it is of great value to correctly align such read parts. However, this is still a difficult task for other state-of-the-art aligners. Furthermore, this advantage helps deSALT to produce better full-length alignments for reads from the genes with small exons (an example is shown in Supplementary Figure 8) and to achieve overall higher #ReadGA statistics.
3. deSALT produces homogeneous alignments. It can be observed from the alignments of deSALT that in local regions, various reads usually have highly similar alignments and exon boundary predictions which also coincide with gene annotations. However, for other aligners, the predicted exon boundaries of the same reads could be more divergent from each other. As shown in the example in Supplementary Figure 9, the homogeneous alignments of deSALT could be more accurate overall, especially for those bases near exon boundaries. The homogeneous alignments are more useful to the study of splicing events since there is less noise in these alignments than in ambiguous alignments.
4. A proportion of bases are aligned to unannotated regions. According to the Ensembl gene annotations, there were about 10% of the bases aligned by deSALT to regions other than the annotated exons: 1) a proportion of the bases (5.60% and 5.21% for the human ONT cDNA and direct RNA sequencing datasets, respectively, and 5.13% for the mouse PacBio dataset) were aligned to intron regions; 2) a proportion of the bases (4.61% and 0.9% for the human ONT cDNA and direct RNA sequencing datasets, respectively, and 4.02% for the mouse PacBio dataset) were aligned to intergenic regions. Minimap2 also had similar proportions of bases aligned to such regions. We found that the alignments of these read parts were highly clustered: i.e., in most cases, there were multiple reads aligned in a local region, indicating that there could be unannotated exons or novel transcripts. Furthermore, we found that deSALT and Minimap2 had similar outputs for these read parts, which also indicates that the alignments are plausible. Two examples in intragenic and intergenic regions are shown in Supplementary Figures 10 and 11, respectively.

**Figure 5.**
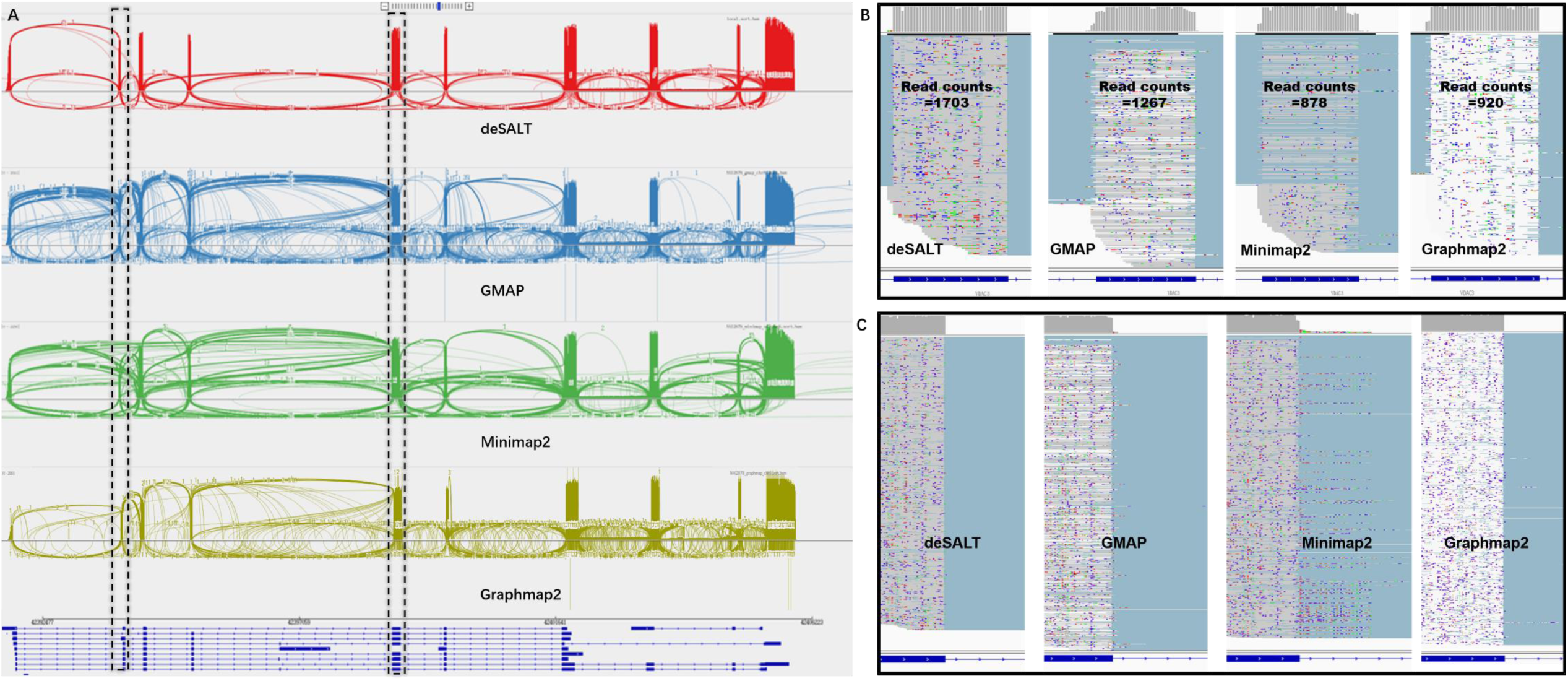
An example of the alignments of real sequencing reads by various aligners. This figure represents the snapshots of the alignments of the reads from the human ONT cDNA dataset around the VDAC3 gene (Chr8: 42391761–42405937) of reference GRCh38. The VDAC3 gene has 10 exons and 12 isoforms (according to Ensembl gene annotation). deSALT, GMAP, Minimap2 and Graphmap2 respectively mapped 2652, 2639, 2462 and 2480 reads to this region. The ratios #BaseGA/#BaseT of the aligners are respectively 78.93% (deSALT), 74.2% (GMAP), 72.69% (Minimap2), 74.02% (Graphmap2), where #BaseT is the total number of bases aligned to the VDAC3 region. This indicates that deSALT produces overall more accurate split alignments. Moreover, the #ReadGA statistics of the aligners are respectively 1630 (deSALT), 889 (GMAP), 751 (Minimap2) and 1030 (Graphmap2), also indicating that deSALT produces better full-length alignments. **(A)** The overall views (sashimi plots) of the alignments indicate that deSALT produces more homogenous alignments. Considering the higher #BaseGA/# BaseT and #ReadGA statistics, such alignments could be more plausible. **(B)** A detailed view at the second exon of the VDAC3 gene (exon length: 40 bp). deSALT aligns much more reads (i.e., 1703 reads) to this short exon than that of GMAP (1267 reads), Minimap2 (878 reads) and Graphmap2 (920 reads), indicating that deSALT potentially handles it better. **(C)** A detailed view at the 3’ splicing site of the fifth exon of the VDAC3 gene (exon length: 153 bp). It shows that the alignments of deSALT near the splicing site are more homogeneous, and the breakpoints of the reads coincide with the annotation, while the alignments of other approaches are more heterogeneous and seem less accurate.

## Discussion

Long read sequencing technologies provide the opportunity to break the limitations of short reads and improve transcriptomics studies. However, complex gene structures and high sequencing errors make it still a non-trivial task to produce accurate full-length alignments to exert the advantages of long RNA-seq reads. So, there is wide demand for the development of more advanced read alignment algorithms to breakthrough this bottleneck. Herein, we proposed deSALT, a novel read alignment algorithm using the de Bruijn graph-based index and a tailored two-pass strategy, as a solution to this important open problem. Mainly, we show how to build and integrate spliced alignment skeletons to handle sequencing errors and complex gene structures in order to generate high-quality spliced reference sequences and use them to produce accurate and homogeneous full-length alignments for long RNA-seq reads.

On both the simulated and real datasets, the deSALT results demonstrate its good sensitivity and accuracy. For most of the datasets, it maps the highest number of bases to their ground truth positions or the positions supported by gene annotations. Its advantage with regard to the recovery of exons and splicing junctions is more obvious, suggesting that deSALT has the ability to produce spliced alignments. This is further demonstrated by several kinds of difficult scenarios, such as very short exons, numerous splicing events, and genes with multiple isoforms.

A more important feature of deSALT is its ability to produce accurate and homogeneous full-length alignments. With the ever-increasing length of reads, this feature is in great demand since it provides the opportunity to investigate gene structures directly. However, it requires the employed aligner to handle many technical issues well and simultaneously. deSALT improves full-length alignment by using several key techniques such as sensitive exon detection, local exon matching, and LSRS generation. For larger numbers of reads, deSALT can recover their splicing junctions by single alignments comprehensively and accurately. And the produced alignments are homogeneous and confident. This contribution has the potential to facilitate many downstream analyses.

According to gene annotations, there are still a proportion of reads and bases being mapped to intron and intergenic regions. Considering the similar results independently produced by deSALT and Minimap2, there could be some unknown transcripts being sequenced, and the alignments are plausible. Moreover, we found that deSALT and Minimap2 similarly clipped a proportion of bases. We tried to align some of the clipped read parts with BLAT [27]. However, no successful alignment was produced (data not shown). In this situation, we realized that these read parts could be extremely low quality.

deSALT does not only use reference genome, it supports the use of gene annotations to enhance the alignment. However, the benchmarking results were to some extent unexpected in that there was no significant difference between the alignments with and without gene annotations, only except for the low depth, high error rate (e.g., ONT 1D) datasets. This is also reasonable since the two-pass strategy enables to mitigate the effect of moderate sequencing errors even if the read depth is low. Moreover, this ability can be enhanced with the increase of sequencing depth, so that high coverage high error rate datasets can also be sensitively aligned without gene annotations. However, this function of deSALT is still useful since gene expression is uneven, i.e., there are always less expressed genes with fewer reads being sequenced, and gene annotations could make their own contributions to the alignment of those reads.

## Conclusion

Overall, the benchmark results on simulated and real datasets demonstrate that the two-pass approach of deSALT is suited to address several difficult technical issues in long RNA-seq read alignment such as sequencing errors, small exons and numerous splicing events. It is able to produce high quality full-length alignments. Moreover, its tailored implementations also enable to achieve good performance. We believe deSALT will be a useful alignment tool to play an important role in many cutting-edge transcriptomics studies.

## Methods

### Genome indexing

The reference genome is indexed by reference de Bruijn graph indexing (RdBG-index) approach, which was initially used by a short read aligner deBGA [34]. Given a reference, a de Bruijn graph of the reference (also called as “RdBG”) is constructed with a user-defined k-mer size, and the unitigs of RdBG are extracted. A RdBG-index is then constructed to index all the vertices (k-mers) of the graph as well as their unitig IDs and offsets. A RdBG-index is composed by several hash table and linear table-based data structures, and it enables to fast retrieve and merge short token matches between reads and reference. Refer to Supplementary Notes and Supplementary Figure 12 for more detailed information about the data structures of RdBG-index and their functions.

It is worth noting that, the construction of RdBG-index for large genomes could cost a couple of hours (129, 112 and 4 minutes for human, mouse and fruit fly, respectively) and several tens of GB RAM space (73GB, 63GB and 5.5GB for human, mouse and fruit fly, respectively), depending on the number of distinct k-mers in the genome. This is mainly because that it needs to extract and sort all the k-mers to construct RdBG and analyze its unitigs at first. However, it is also affordable since the index needs only to be built once before use, and we also provide pre-built RdBG-indexes for human, mouse and fruit fly reference genomes (Supplementary Notes).

### Steps of the deSALT approach

deSALT aligns input reads in three major steps as follows:

1. Alignment skeleton generation (first-pass alignment): for each of the reads, deSALT uses the RdBG-index [34] to find the maximal exact matches between the unitigs of a reference de Buijn graph (RdBG) and the read (termed as U-MEMs) and to build one or more alignment skeletons using an SDP approach.
2. Exon inference: deSALT maps all the alignment skeletons to the reference and infers potential exons from the projections of the skeletons. A local sequence-based scoring system [40] is employed to refine the inferred exons. Moreover, it is optional to introduce gene annotations as additional information to enhance exon detection.
3. Refined alignment (second-pass alignment): for each of the reads, deSALT finds additional local matches to the inferred exons with shorter tokens (seeds) than the ones used in the first step. Further, it combines the newly found matches and the alignment skeleton to retrieve and stich all the spanning exons to build an LSRS and implement a base-level read alignment.

### Alignment skeleton generation (first-pass alignment)

For a read, deSALT extracts *l*-mers (*l*<*k*, default value: *l*=15) at every *m* bp (default value: *m*=5) as seeds and matches them to the unitigs of RdBG with the RdBG-index. The matches are extended in both directions to generate U-MEMs. deSALT then merges co-linear U-MEMs on the same unitigs as super U-MEMs (SU-MEMs) and maps the SU-MEMs, as well as the U-MEMs that cannot be merged, to reference genome as MBs to build alignment skeletons.

deSALT uses the MBs as vertices to build a direct acyclic graph (DAG). The edges of the DAG are defined by the pairs of MBs whose distances are no longer than a pre-defined maximum intron length, T_intron_(default value: T_intron_=200,000 bp). A weight is assigned to each of the edges on the basis of the sizes of the two corresponding MBs and their distances (Supplementary Notes). An SDP approach is then used to find the path with the largest sum weight as the alignment skeleton. It is also worth noting that deSALT could produce multiple alignment skeletons with very similar scores (sum weights) for some of the reads, considering that such reads possibly have multiple “equally best” alignments.

### Exon inference

deSALT maps all the alignment skeletons to the reference genome and uses a set of pre-defined rules (Section 3.1 of the Supplementary Notes) to iteratively combine the genomic regions covered by alignment skeletons from upstream to downstream. It is optional to introduce a gene annotation file (in GTF format) into this process. deSALT treats known gene isoforms as a special kind of alignment skeletons, and it also maps them to the reference genome so that the genomic regions covered by known gene isoforms and the alignment skeletons are combined together. The combined regions are then recognized as draft exons, and their lengths and alignment skeleton coverages are calculated. The ones with too short a length and too low coverage are then filtered out.

A local sequence-based scoring system [40] is then employed to refine the draft exons (Section 3.2 of Supplementary Notes). For each of the draft exons, deSALT selects two small flanking regions. The scoring system uses pre-defined acceptor and donor scoring matrixes to score each of the positions in the upstream and downstream regions respectively. The positions with the highest scores in the two regions are recognized as acceptor and donor splicing sites, and the region in between is determined to be a refined exon.

### Refined alignment (second-pass alignment)

Refined alignment is mainly implemented in two sub-steps as follows:

1. LSRS generation: deSALT splits the read into a series of parts and composes partial LSRSs for each of them separately (Section 4.1 of the Supplementary Notes). Each read part is defined as a specific substring of the read within two neighboring MBs of its alignment skeleton. For a read part, deSALT detects a set of exons (termed “spanning exons”) which are placed in between or nearby the two corresponding MBs and have short matches to the read part. The spanning exons are then stitched together as the whole LSRS.
2. Base-level alignment: deSALT aligns each of the read parts to its corresponding LSRS using a SIMD-based implementation [29, 32] of semi-global alignment (Section 4.2 of the Supplementary Notes). Furthermore, deSALT checks if there are large deletions in the CIGAR information; if there are, deSALT removes the corresponding deletion part(s) in the LSRS and realigns the read with the updated LSRS. This process is helpful for handling exons with alternative splicing sites. A schematic illustration is in Supplementary Figure 13.

It is also worth noting that for the reads with multiple alignment skeletons, deSALT processes each of the skeletons separately and possibly produces multiple alignments for one read. In this situation, deSALT chooses the alignment with the highest score as the primary alignment and outputs other alignments as secondary alignments.

### Implementation of the simulation benchmark

All the benchmarks were implemented on a server with Intel Xeon E4280 CPU at 2.0GHZ and 1 Terabytes RAM, running Linux Ubuntu 16.04. The simulated datasets were generated from the reference of three organisms: Homo sapiens GRCh38 (human), Mus musculus GRCm38 (mouse), and Drosophila melanogaster r6 (fruit fly), with corresponding Ensembl gene annotations [38]. There are 60 datasets used for the benchmark, and each of them was generated by a specific combination of sequencing model, simulated transcriptome and coverage.

6 sequencing models were built according to previous studies [17, 35, 36], to comprehensively benchmark the aligners on the datasets produced by various long read sequencing platforms. For PacBio platforms, there were two models built with fixed parameters: “PacBio ROI reads” (error rate = 2%, mean read length = 2000 bp) and “PacBio subreads” (error rate = 15%, mean read length = 8000 bp). For ONT platforms, four models were built. Two of them were also with fix parameters: “ONT 2D reads” (error rate = 13%, mean read length = 7800 bp) and “ONT 1D reads” (error rate = 25%, mean read length = 7800 bp). And the other two models, “PS-ONT reads” and “NS-ONT reads” were automatically built by PBSim [37] and NanoSim [36] based on a real ONT sequencing dataset (SRA Accession Number: SRR2848544), respectively. Although previous studies [36] indicate that data-based models more coincide with the characteristics of real sequencing, we considered that real ONT datasets could also have various characteristics [35] and it is hard to build a number of models for benchmark. Thus, we used the two parameter-based models, “ONT 2D reads” and “ONT 1D reads”, as a complement, where the 25% and 12% error rates coincide with typical error rates of ONT 2D and 1D reads [18, 19]. The parameters and command lines of PBSim and NanoSim are in Supplementary Notes.

We built two categories of simulated transcriptomes. The first category only has one simulated transcriptome (called as “H-all” transcriptome) which is from all the coding genes of human, and the second category has three simulated transcriptomes (called as “H-se”, “M-se” and “F-se” transcriptomes respectively) which are from three sets of randomly selected genes of human, mouse and fruit fly, respectively. Each of the transcriptomes was used to generate a series of simulated datasets with various sequencing models. Some details are as following.

H-all transcriptome was composed by all the coding genes recorded in Ensembl gene annotations of human. For each of the genes with alternative splicing, one of the isoforms was randomly selected as “highly expressed” and the other isoforms were “lowly expressed”, and all the genes with single isoform were “highly expressed”. Then transcript sequences were made for all the isoforms of all the genes, i.e., for a specific gene isoform, all its exons were concatenated according to the Ensembl annotations to build a transcript sequence.

Given a sequencing model, the sequences of highly and lowly expressed isoforms were input to the specifically configured simulator (PBSim or NanoSim) to implement a 30X and a 4X coverage *in silico* sequencing, respectively, and the two generated datasets were mixed as one for the use of benchmark. Thus, there were in total 6 datasets simulated with H-all transcriptome by various sequencing models. Herein, we used the term “coverage” only as a measure of the sizes of the datasets. That means, a specific sequencing coverage dX (e.g., *d* = 30) indicates that a set of transcript sequences were input into the simulator to produce a dataset whose total size is *L*_*TS*_ × *d*, where *L*_*TS*_ is the total length of the transcript sequences.

H-se, M-se and F-se transcriptomes were composed in a similar way to a previous study [30]. For a specific species, the gene annotations were scanned to extract three sets of genes. Each of them corresponds to a specific type, i.e., genes with single splicing isoforms, genes with multiple splicing isoforms, and genes with short exons (<31 bp), respectively. A number of genes were randomly selected for each of the gene sets and other genes were no longer used (refer to Supplementary Table 11 for the numbers of selected genes). Transcript sequences were then made for the selected genes with the same approach, i.e., given a gene isoform, a transcript sequence was made by concatenating all its exons, and the sequences were made for all the isoforms of all the selected genes. H-se, M-se and F-se transcriptomes were then composed by the generated transcript sequences of human, mouse and fruit fly, respectively.

For each of the transcriptomes (H-se, M-se or F-se), 3 datasets with various coverages, i.e., 4X, 10X and 30X were simulated by a given sequencing model. Thus, with the 3 transcriptomes, 3 kinds of coverages and 6 models, there were in total 54 datasets produced. These datasets are used to assess the ability of the aligners on various species, coverage and platforms.

The following five metrics were used to evaluate the alignment results of the simulated reads.

Base%: the proportion of bases being correctly aligned to their ground truth positions (i.e., the mapped positions of the bases were within 5 bp of their ground truth positions).

Exon%: the proportion of exons being correctly mapped. An exon in a certain read was considered to be correctly mapped only if its two boundaries were mapped within 5 bp of their ground truth positions.

Read80%: the proportion of Read80% reads. A read was considered to be a Read80% read only if it met two conditions, namely *N*_*T*_/*N*_*G*_ > 80% and *N*_*T*_/*N*_*P*_ > 80%, where *N*_*G*_ is the number of ground truth exons within the read, *N*_*P*_ is the number of exons predicted by the alignment, and *N*_*T*_ is the number of true positive exons. Herein, a predicted exon is considered to be a true positive exon only if there was a ground truth exon in the read, and the distance between the corresponding boundaries of the predicted exon and the ground truth exon were within 5 bp.

Read100%: the proportion of Read100% reads. A read was considered to be a Read100% read only if it met two conditions, namely *N*_*T*_/*N*_*G*_ = 100% and *N*_*T*_/*N*_*P*_ = 100%. It is worth noting that a Read100% read indicates that the read has a highly correct full-length alignment.

#Bases/m: the number of bases aligned per minute, which depicts the alignment speed and is computed by *N*_*base*_/*T*_*aln*_, where *N*_*base*_ is the total number of bases in the dataset and *T*_*aln*_ is the wall clock time.

### Implementation of the real data benchmark

The benchmarks were implemented with the same hardware environment as that used for the simulated datasets. Three real datasets respectively produced by ONT and PacBio platforms were used. Two of them are from the NA12878 sample, and produced by cDNA sequencing and direct RNA sequencing, respectively. They were sequenced by the ONT MinION sequencer by using direct RNA sequencing kits (30 flowcells) and the 1D ligation kit (SQK-LSK108) on R9.4 flowcells with R9.4 chemistry (FLO-MIN106). More detailed information about this dataset is available at https://github.com/nanopore-wgs-consortium/NA12878. The third dataset (SRA Accession Number: SRR6238555) is a full-length isoform sequencing of total mouse RNA using standard PacBio-seq protocols [39]. The availability of the two real datasets is provided in the Supplementary Notes.

The following metrics were used to evaluate the alignment results of the real sequencing reads.

#BaseA: the number of bases being aligned.

#BaseGA: the number of bases aligned to the positions within annotated exons.

#ExonP: the number of exons predicted by the alignments (also termed “predicted exons”). Here, the predicted exons in various reads were independently considered.

#ExonGO: the number of predicted exons being overlapped by annotated exons (also termed “overlapped exons”). Herein, a predicted exon was considered to be overlapped by annotated exons only if there was at least one annotated exon and at least 10 bp overlapping between the predicted exon and the annotated exon.

#ExonGA: the number of predicted exons being exactly matched by annotated exons (also termed “exactly matched exons”). Herein, a predicted exon was considered to be exactly matched by annotated exons only if there was an annotated exon and the distance between the corresponding boundaries of the predicted exon and the annotated exon were within 5 bp.

#ExonGA(x): the number of exactly matched exons whose lengths were shorter than x bp.

#ReadGA: the number of ReadGA reads. A read was considered to be a ReadGA read only if each of the intron boundaries implied by its alignment was within 5 bp of an annotated exon. Herein, a ReadGA read indicates that the read could has a correct full-length alignment.

## Declarations

### Ethics approval and consent to participate

Not applicable.

### Consent for publication

Not applicable.

### Availability of data and material

The source code of deSALT and the scripts of data simulations and benchmarking are available at https://github.com/hitbc/deSALT (DOI: 10.5281/zenodo.3479485).

Please refer to the Supplementary Notes for the availability of the simulated and real sequencing datasets.

### Author contributions

BL designed the method, YL implemented the method, and BL, YL, JL and HG performed the analysis. All of the authors wrote the manuscript. BL and YL contributed equally to this work.

## Acknowledgements

We are very grateful to Prof. Yi Xing of the University of Pennsylvania and the Children’s Hospital of Philadelphia and Dr. Mingxiang Teng of Moffitt Cancer Center for their helpful discussion.

