## supplementary for "deSALT: fast and accurate long transcriptomic read alignment with de Bruijn graph-based index"

**Supplementary Material**

Bo Liu[1, 2], Yadong Liu[1, 2], Junyi Li[1], Hongzhe Guo[1], Tianyi Zang[1, *], and Yadong Wang[1, *]

[1]Center for Bioinformatics, School of Computer Science and Technology, Harbin Institute of Tech-nology, Harbin, Heilongjiang 150001, China
[2] These authors contributed equally to this work.

Contact:, and

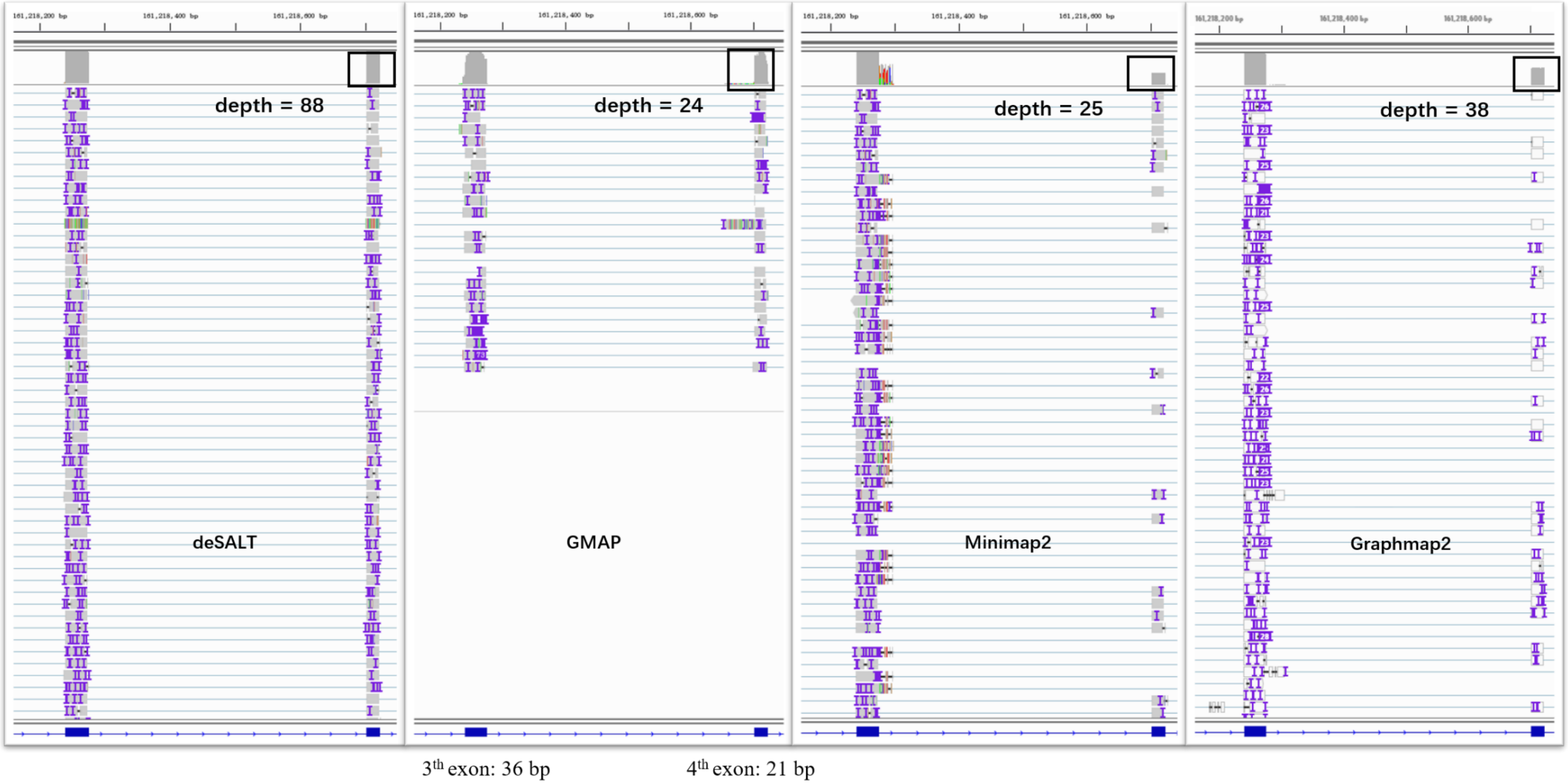

**Supplementary Figure 1. An example of the alignment of simulated reads around short exons**

This figure represents the Integrative Genome Viewer (IGV) [1] snapshots of the alignments of the reads from the simulated 30X PacBio subread human dataset (from randomly selected genes), around the FECR1G gene (Chr1: 161215297–161219248) of reference GRCh38. FECR1G has 5 exons, and the snapshots focus on the third and fourth exons whose lengths are 36 bp and 21 bp respectively. According to the ground truth, there are 88 reads spanning these two short exons, and deSALT aligns all of them correctly. However, GMAP only aligns a proportion (i.e., 24) of these reads to these two exons correctly. Minimap2 and Graphmap2 align most of the reads to the third exon, but the breakpoints of the reads are more divergent from each other. Moreover, both of them only align a few (i.e., 25 and 38) reads to the fourth exon. This case suggests the outstanding ability of deSALT to handle the short exon parts of reads, which is beneficial to the production of accurate alignments.

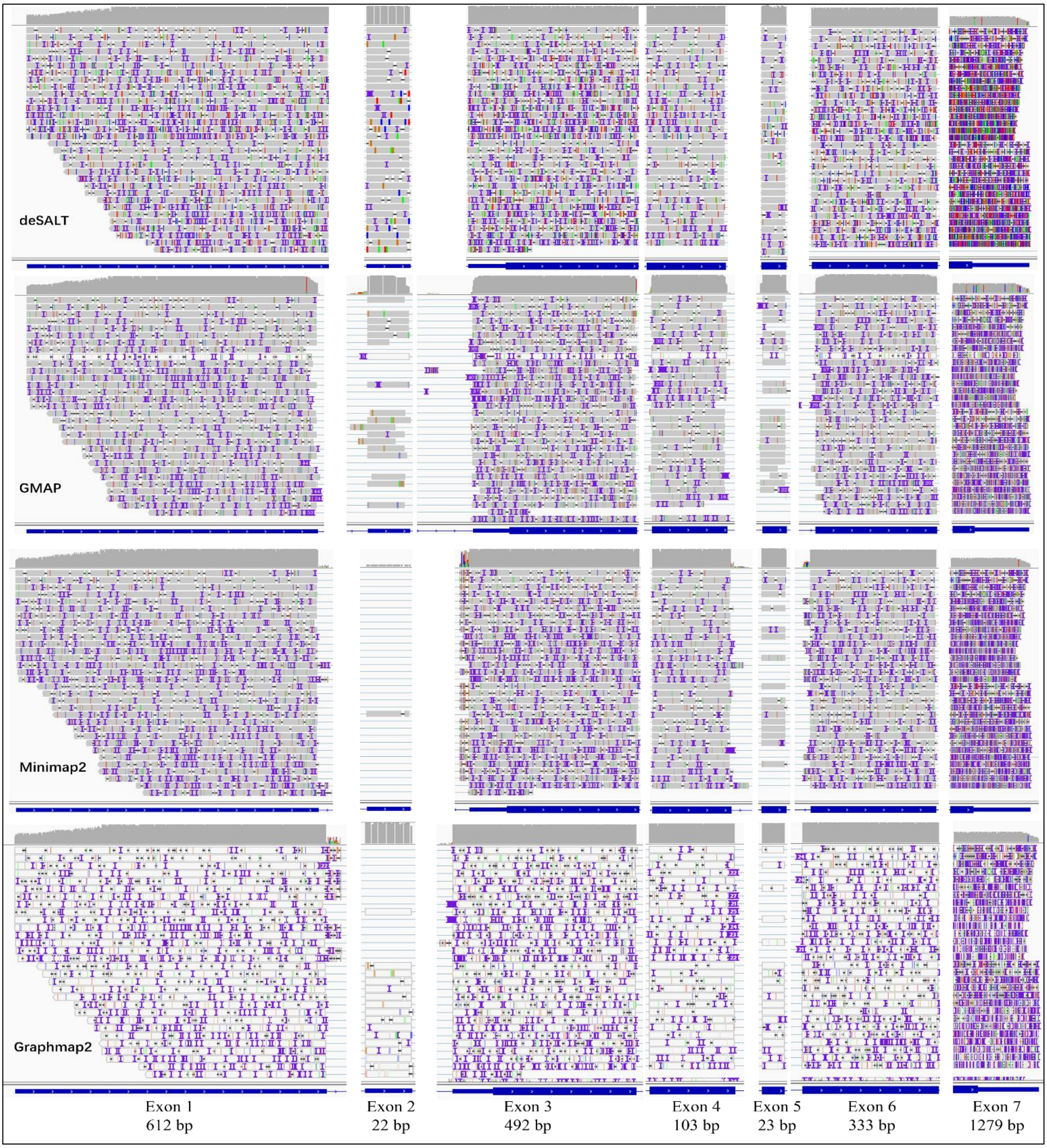

**Supplementary Figure 2. An example of the alignment of simulated reads from transcripts with multiple exons**
This figure represents the Integrative Genome Viewer (IGV) [1] snapshots of the alignments of the reads from the simulated 30X ONT 2D fruit fly dataset, around the CG42747 gene (Chr3L: 6400590–6485964) of reference BDGP6 (worth noting that the intron regions are removed in the figure). The region contains 7 exons of gene CG42747 (their lengths are given at the bottom of the figure), and there are 32 reads in total in this gene region according to the ground truth. It is observed from the figure that deSALT can correctly produce full-length alignments for all the reads spanning the multiple exons of the CG42747 gene. However, for GMAP, Minimap2 and Graphmap2, some of the reads are not aligned correctly. GMAP misses the short exon parts for a proportion of the reads, and for two of the reads it only aligns a few bases. Minimap2 correctly aligns only one of the reads for its exon2 part and nearly 50% of the reads for their exon5 parts. Graphmap2 also only aligns a proportion of the reads to exon2 and exon5. This case suggests the ability of deSALT to handle the reads from transcripts with multiple exons.

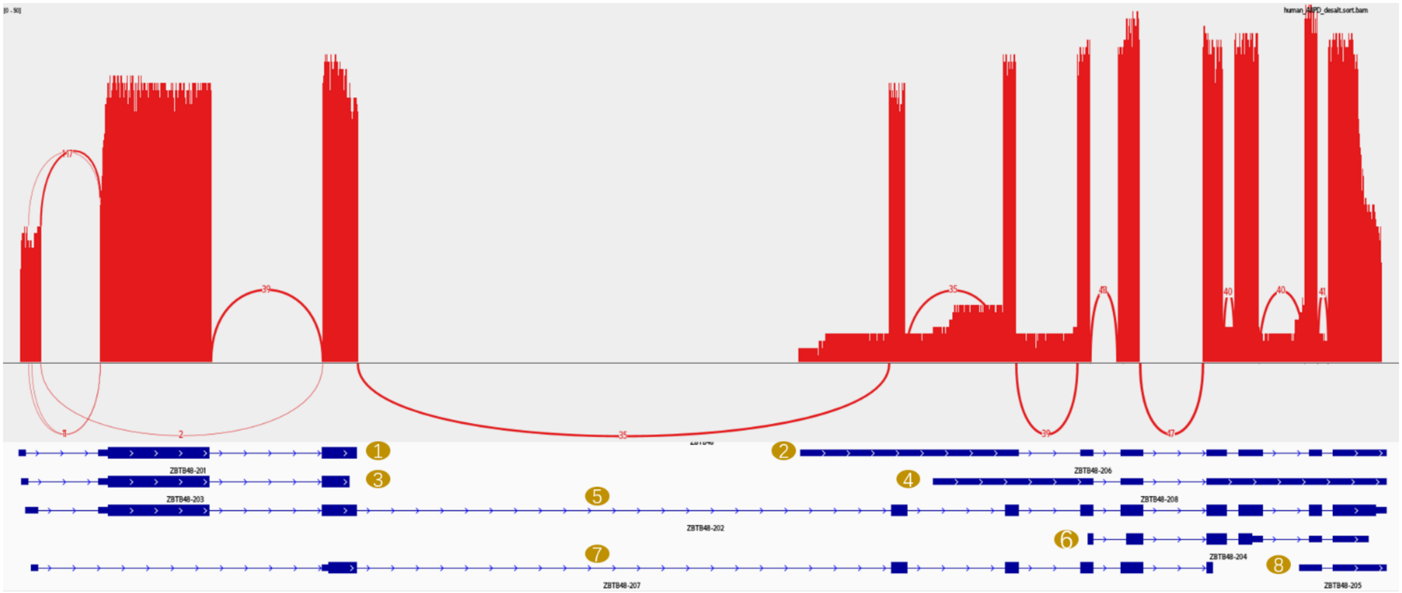

**Supplementary Figure 3. An example of the alignment by deSALT for the reads from genes with alternative splicing**

This figure represents the sashimi plot of the alignments of the reads from simulated human all protein coding genes PS-ONT dataset, around the ZBTB48 gene (Chr1: 6579994–6589280) of reference GRCh38. The ZBTB48 gene has 8 isoforms (marked by yellow ovals). The fifth isoform was selected as “highly expressed” (33 reads were simulated in total) and the other ones were “lowly expressed” (34 reads were simulated in total for the 7 isofroms). deSALT aligned all the 67 reads to ZBTB48 region correctly. The numbers of Read100 reads for “highly expressed” and “lowly expressed” isoforms are high (32 and 32, respectively). This indicates that deSALT enables to handle the reads from lowly expressed isoforms as well as highly expressed isoforms.

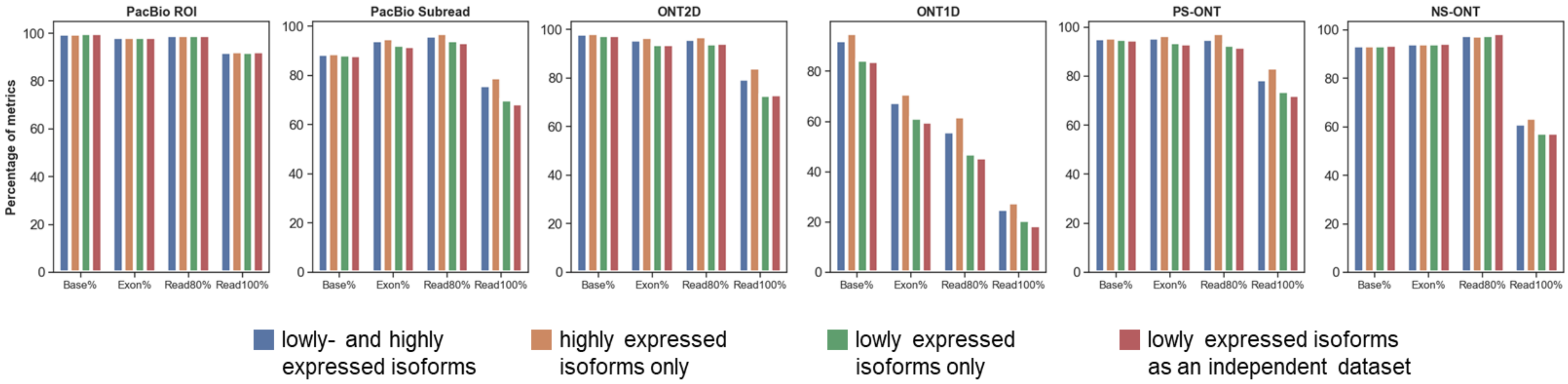

**Supplementary Figure 4. Assessmenet on the reads from lowly- and highly expressed isoforms of the human all protein coding gene datasets**

The figure depicts the yields of deSALT on the reads from lowly- and highly expressed isoforms of the human all protein coding gene datasets. The blue bars indicate the overall yields on the human all protein coding gene datasets. The orange bars indicate the yields on the reads from highly expressed isoforms. The green bars indicate the yields on the reads from lowly expressed isoforms. Moreover, deSALT was asked to align the reads from lowly expressed isoforms as independent datasets, and the results are shown by the red bars.

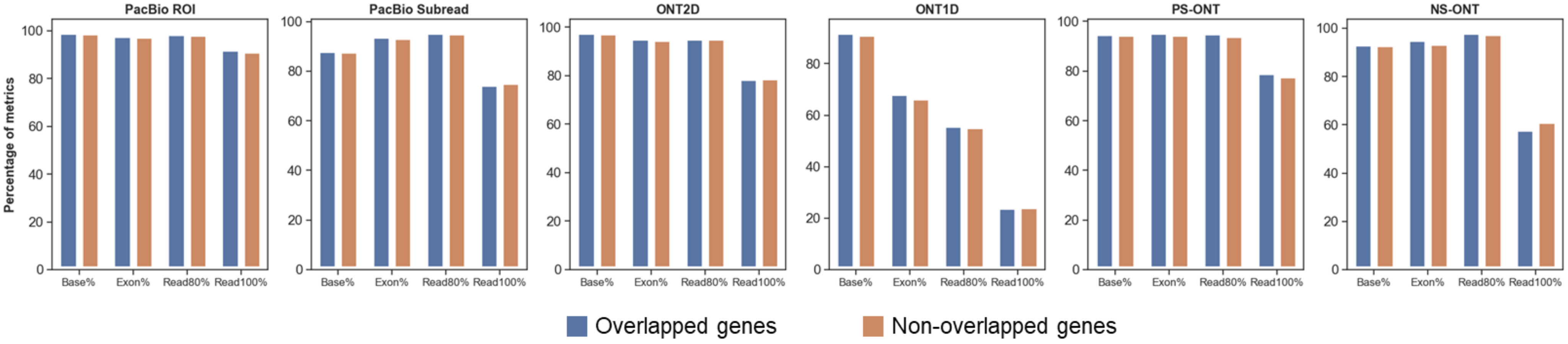

**Supplementary Figure 5. Assessmenet on the reads from genomically overlapped genes of the human all protein coding gene datasets**

The figure depicts the yields of deSALT on the reads from genomically overlapped genes and non-overlapped genes. Blue bars indicate the yields on the reads from genomically overlapped genes, and orange bars indicate the yields on the reads from non-overlapped genes.

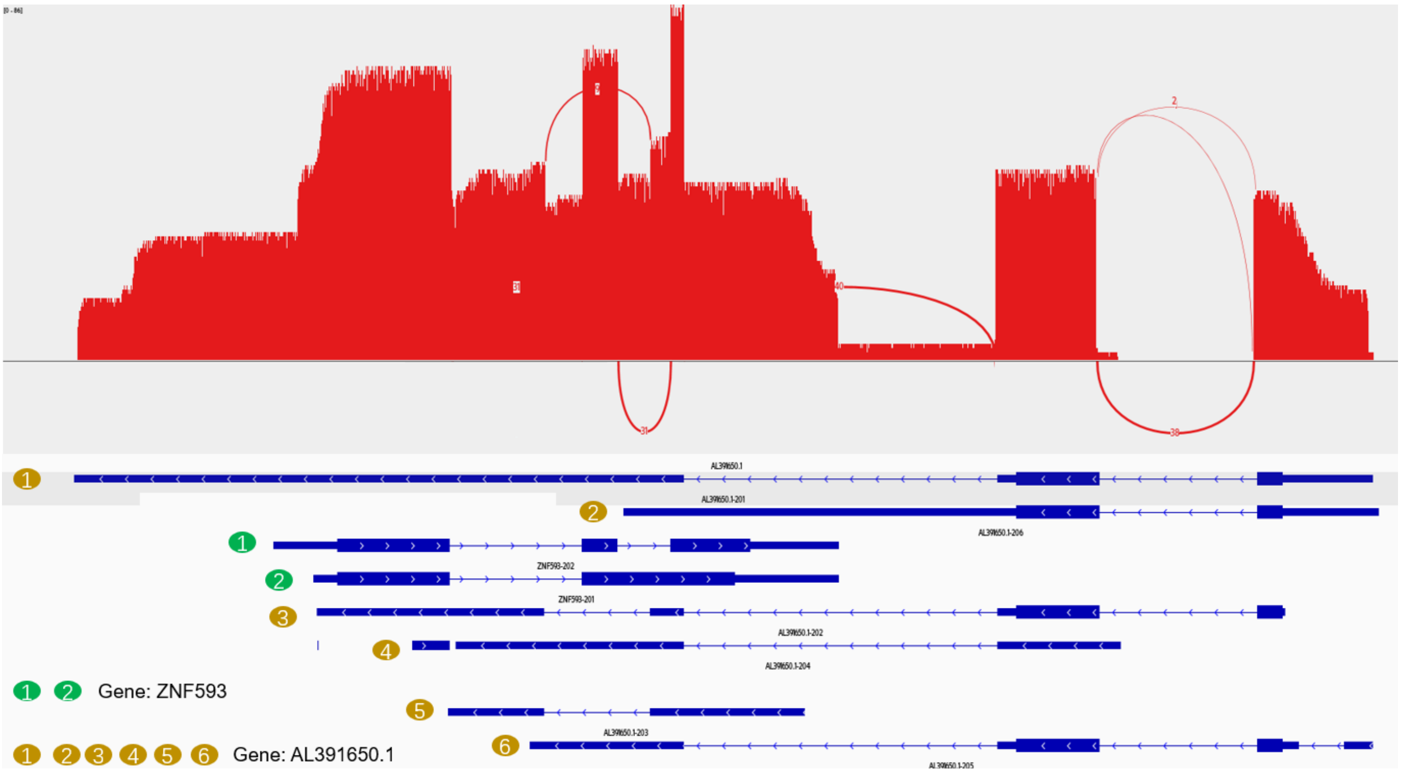

**Supplementary Figure 6. An example of the alignment by deSALT for the reads from genomically overlapped genes**

This figure represents the sashimi plot of the alignments of the reads from simulated all human protein coding gene PS-ONT dataset, around ZNF593 gene (Chr1: 26169908–26170873) and AL391650.1 gene (Chr1: 26169516–26171831) of reference GRCh38. AL391650.1 gene has 6 isoforms in the reverse strand (marked by yellow ovals) and the ZNF593 gene has 2 isoforms in the forward strand (marked by green ovals). Moreover, ZNF593 and AL391650.1 overlap. There are totally 93 reads (36 from ZNF593 gene and 57 from AL391650.1 gene) according to the ground truth. deSALT aligned all the 93 reads to AL391650.1 and ZNF593 regions correctly. The numbers of Read100 and Read80 reads are 90 and 93 respectively. This suggests that deSALT enables to handle the reads from pairs of genes that overlap genomically but are on opposite strands.

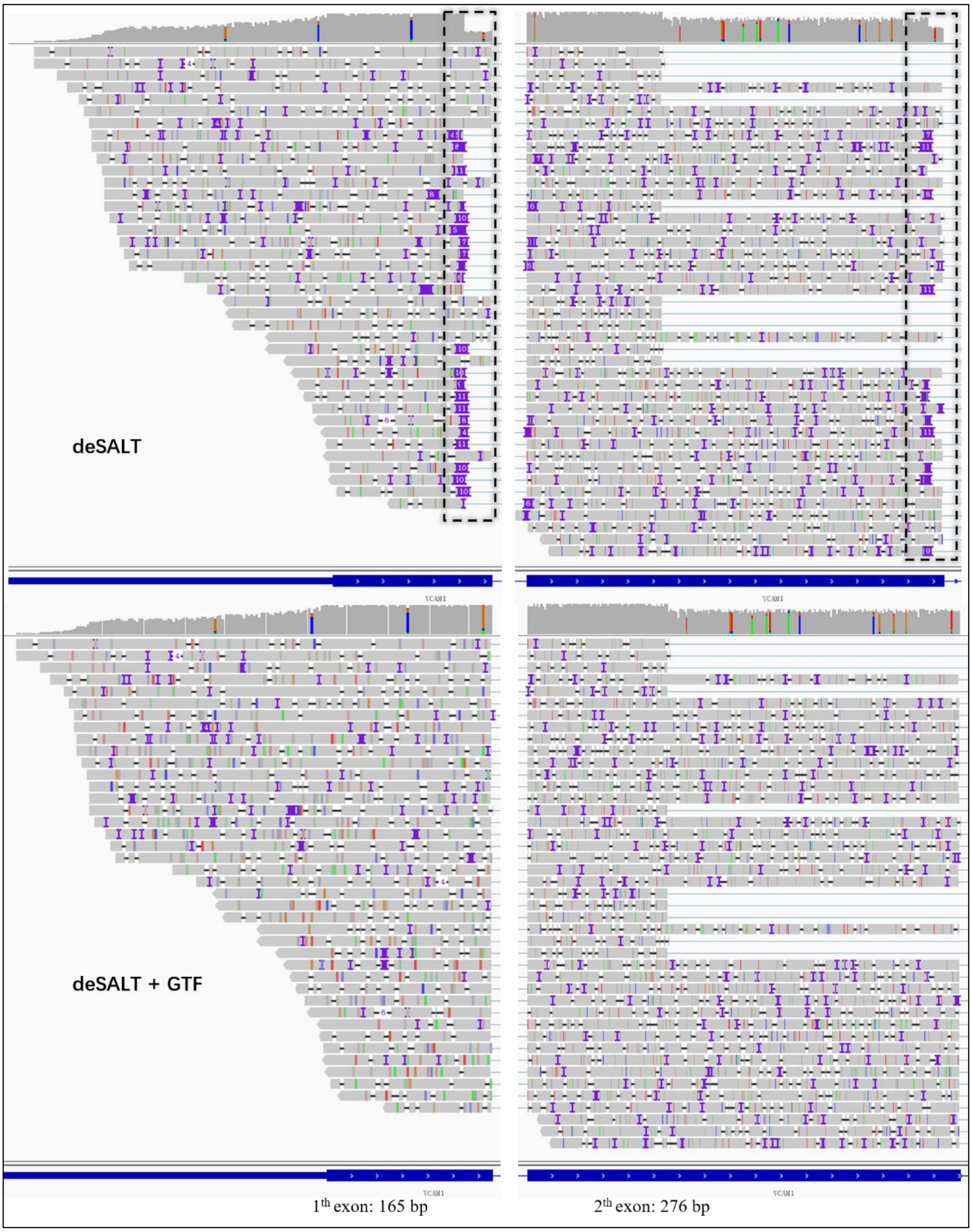

**Supplementary Figure 7. An example of the alignment of error-prone reads by deSALT with and without gene annotations**

This figure represents the Integrative Genome Viewer (IGV) [1] snapshots of the alignments of the reads from the simulated 10X ONT 1D human dataset (from randomly selected genes), around the VCAM1 gene (Chr1: 100719640–100739046) of reference GRCh38. VCAM1 gene has 9 exons, and the snapshots focus on the first two exons; moreover, it is also worth noting that the second exon has two alternative splicing sites (which are respectively at Chr1: 100720565 and Chr1: 100720751). According to the ground truth, there are 43 reads in total in this gene region, and the alignments of deSALT with (termed "deSALT+GTF" in the figure) and without (termed "deSALT" in the figure) annotations are represented in the lower and upper parts of the figure, respectively. It is observed that the high sequencing errors (error rate: 25%) affect the alignment of the reads, especially the bases near the exon boundaries (marked by dashed rectangles in the upper part of the figure). The bases can be aligned with the help of the gene annotation more confidently, and the alignment is obviously improved.

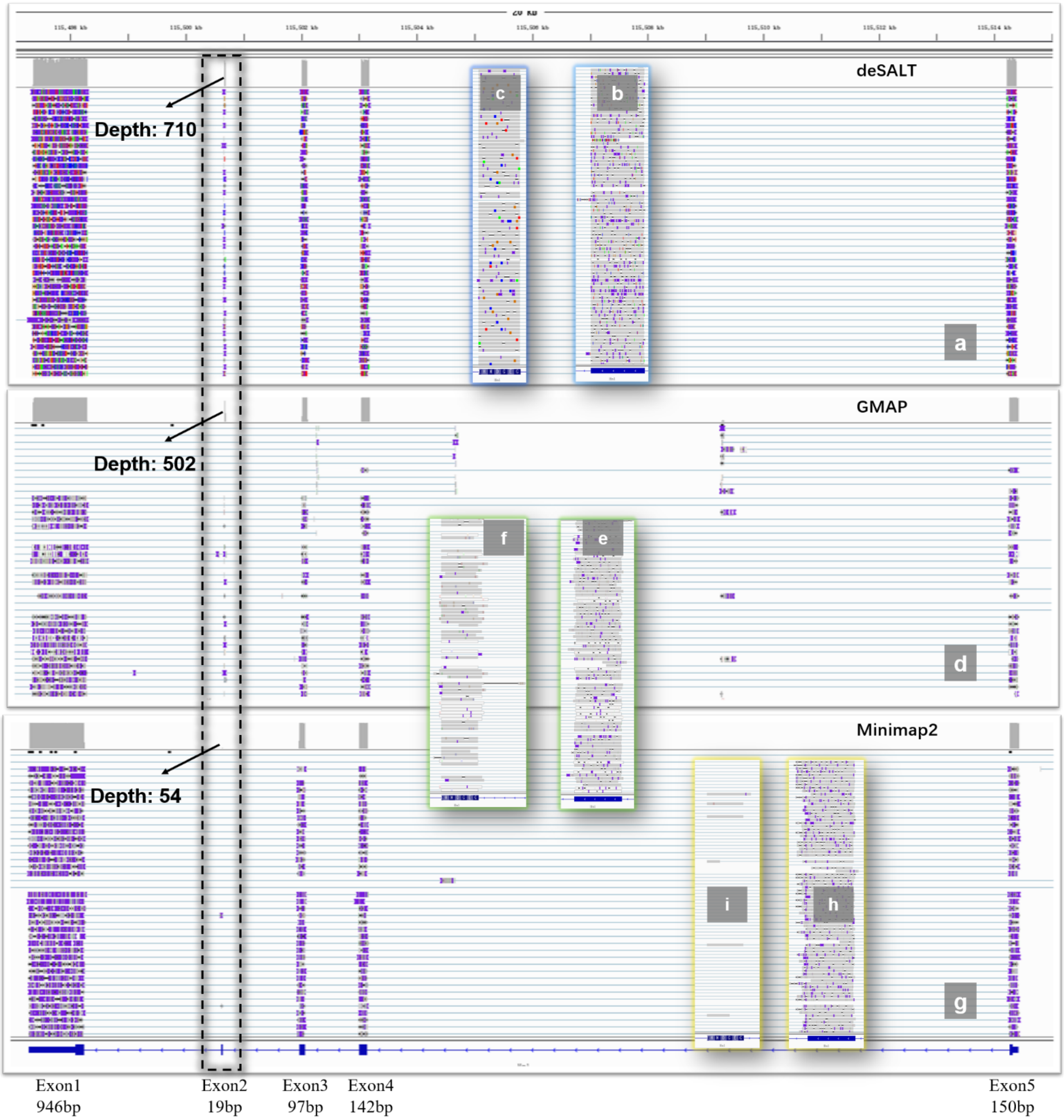

**Supplementary Figure 8. An example of the alignment of real sequencing reads from transcripts with small exons**

This figure represents the Integrative Genome Viewer (IGV) [1] snapshots of the alignments of the reads from the real mouse PacBio dataset, around the Hn1 gene (Chr11: 115497353–115514370) of reference GRCm38. The Hn1 gene has one isoform containing 5 exons (according to Ensembl gene annotation; the lengths of the exons are marked at the bottom of the figure), and exon2 is very short (i.e., 19 bp). In the alignment results, deSALT, GMAP, and Minimap2 map 816, 863, and 857 reads to the Hn1 gene region respectively. However, the number of reads being aligned to exon2 by GMAP (i.e., 502 reads) and Minimap2 (i.e., 54 reads) is less than that of deSALT (710 reads). Moreover, the #ReadGA statistics of the aligners in the Hn1 region are 704 (deSALT), 467 (GMAP), and 48 (Minimap2). This case indicates that deSALT has better ability to handle small exons in real sequencing reads and this ability is also helpful in producing full-length alignments for more reads. It is also worth noting that GraphMap2 raised a “segement fault” during running this dataset, so that its alignment is not shown.

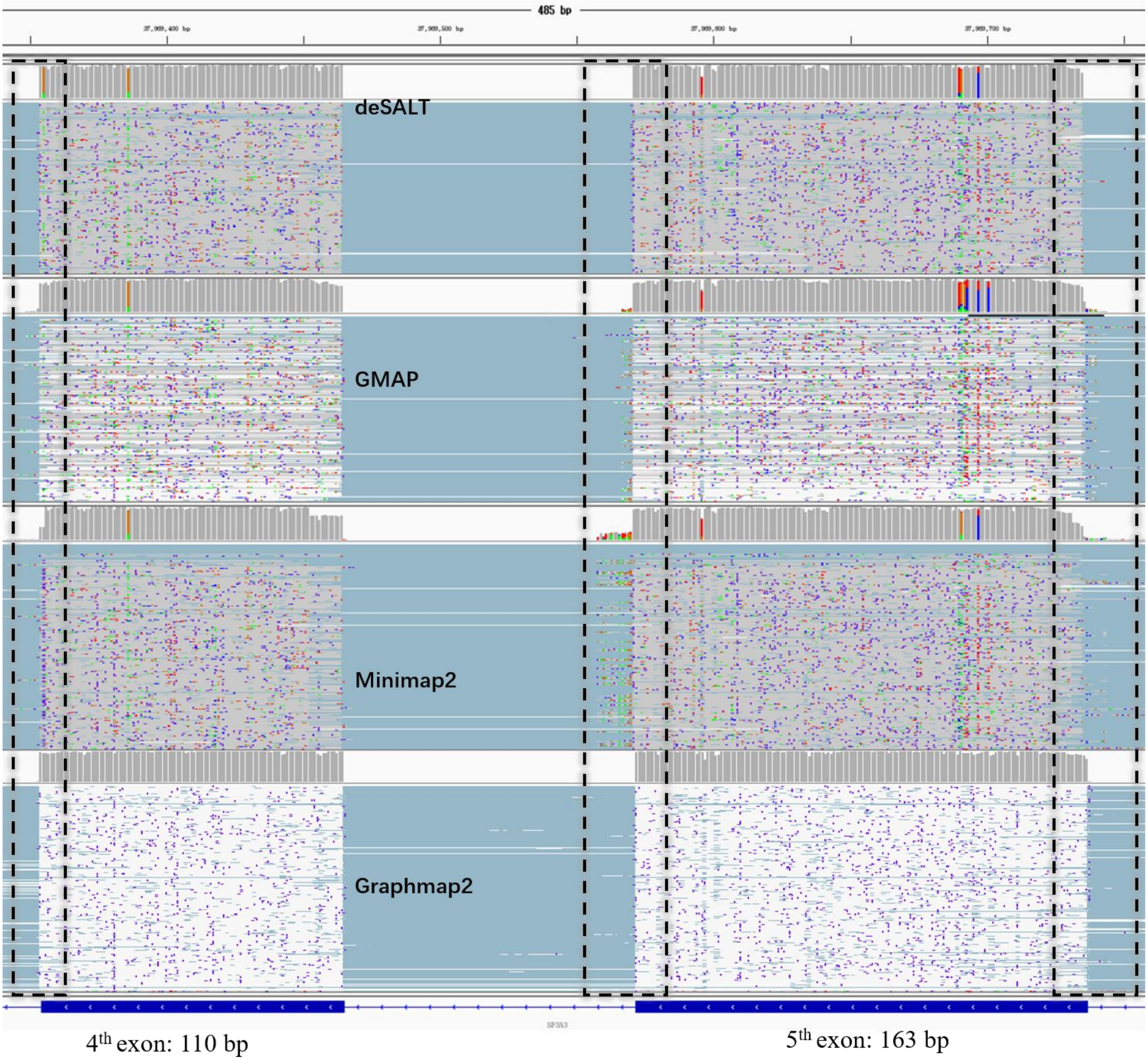

**Supplementary Figure 9. An example of the homogeneous alignment of real sequencing reads by deSALT**
This figure represents the Integrative Genome Viewer (IGV) [1] snapshots of the alignments of the reads from the real human ONT cDNA dataset, around the SF3A3 gene (Chr1: 37956980–37990110) of reference GRCh38. SF3A3 gene has 17 exons (according to Ensembl gene annotation), and the snapshots focus on the fourth and fifth exons whose lengths are 110 bp and 163 bp, respectively. In the alignment results, there are 658, 426, 633 and 797 reads mapped to this region by deSALT, GMAP, Minimap2 and Graphmap2, respectively. It is observed that the alignments produced by deSALT are highly homogeneous. However, the alignments produced by GMAP and GraphMap2 are more heterogenous for both the inner regions and boundaries of the two exons. The alignments produced by Minimap2 are not as heterogenous as those produced by GMAP and GraphMap2 in the inner regions, but they are very heterogenous at the exon boundaries. Moreover, the #ReadGA statistics of the aligners in the SF3A3 region are respectively 411 (deSALT), 109 (GMAP), 173 (Minimap2) and 298 (Graphmap2), and the ratios #BaseGA/#BaseT of the aligners are respectively 89.82% (deSALT), 86.92% (GMAP), 83.7% (Minimap2) and 81.8% (Graphmap2), where #BaseT is the total number of bases aligned to the SF3A3 region by the corresponding aligner. This case indicates that deSALT enables the production of homogeneous read alignments and it has the potential to improve the overall alignment accuracy.

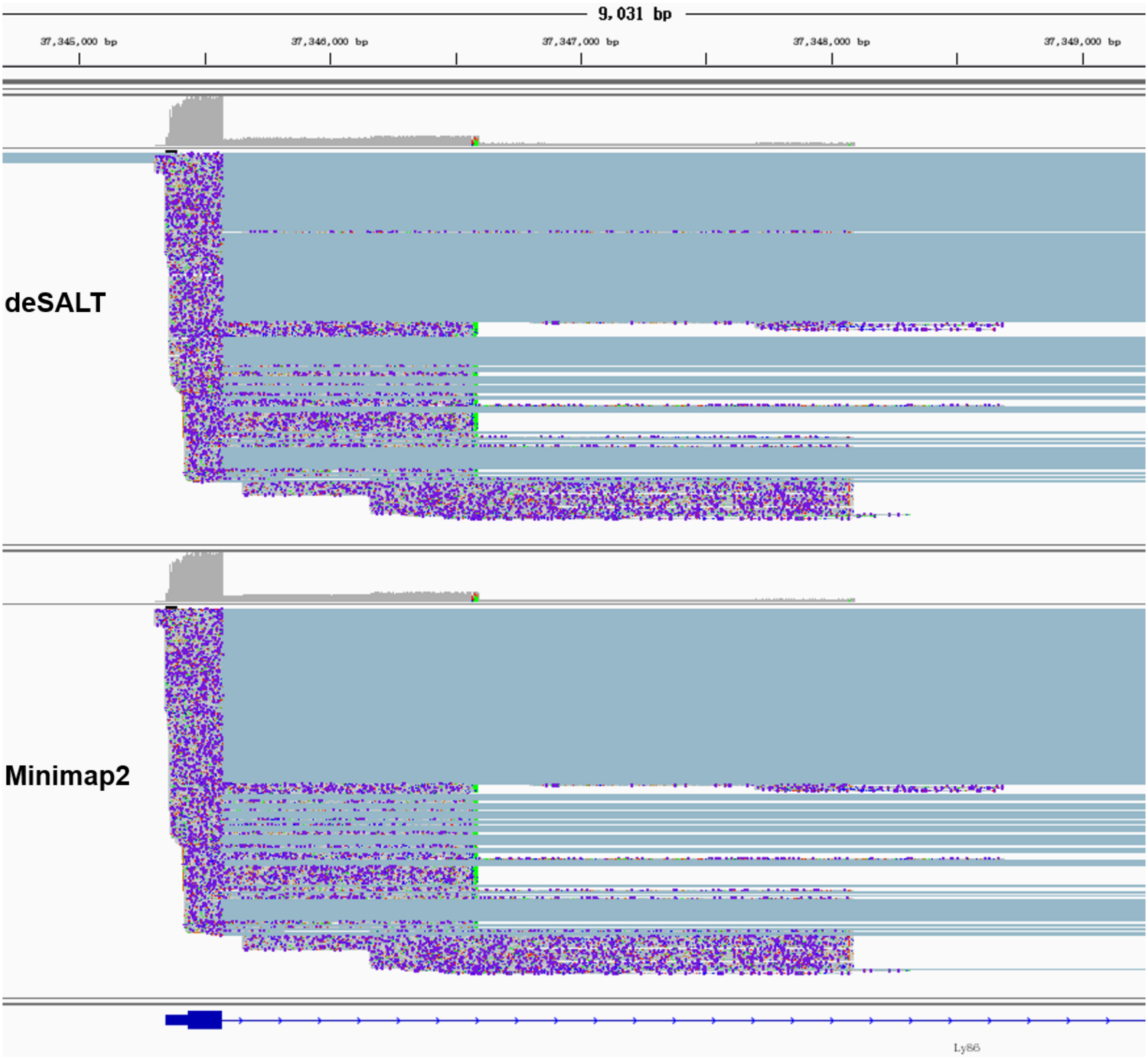

**Supplementary Figure 10. An example of the reads aligned to intron regions**

This figure represents the Integrative Genome Viewer (IGV) [1] snapshots of the alignments of the reads from the real mouse PacBio dataset, around the Ly86 gene (Chr13: 37345345–37419036) of reference GRCm38. The Ly86 gene has 5 exons (according to Ensembl gene annotation), and the snapshots focus on the intron region between the first and second exons. It is observed from the alignment results that both deSALT and Minimap2 align 430 reads to this region. Moreover, similar numbers of reads are aligned to the intron region between the first and second exons, and the detailed alignments (i.e., CIGARs) produced by the two aligners are also similar. Considering the numerous aligned reads and the similar alignments independently produced by various aligners, there could be unannotated splicing events in this region.

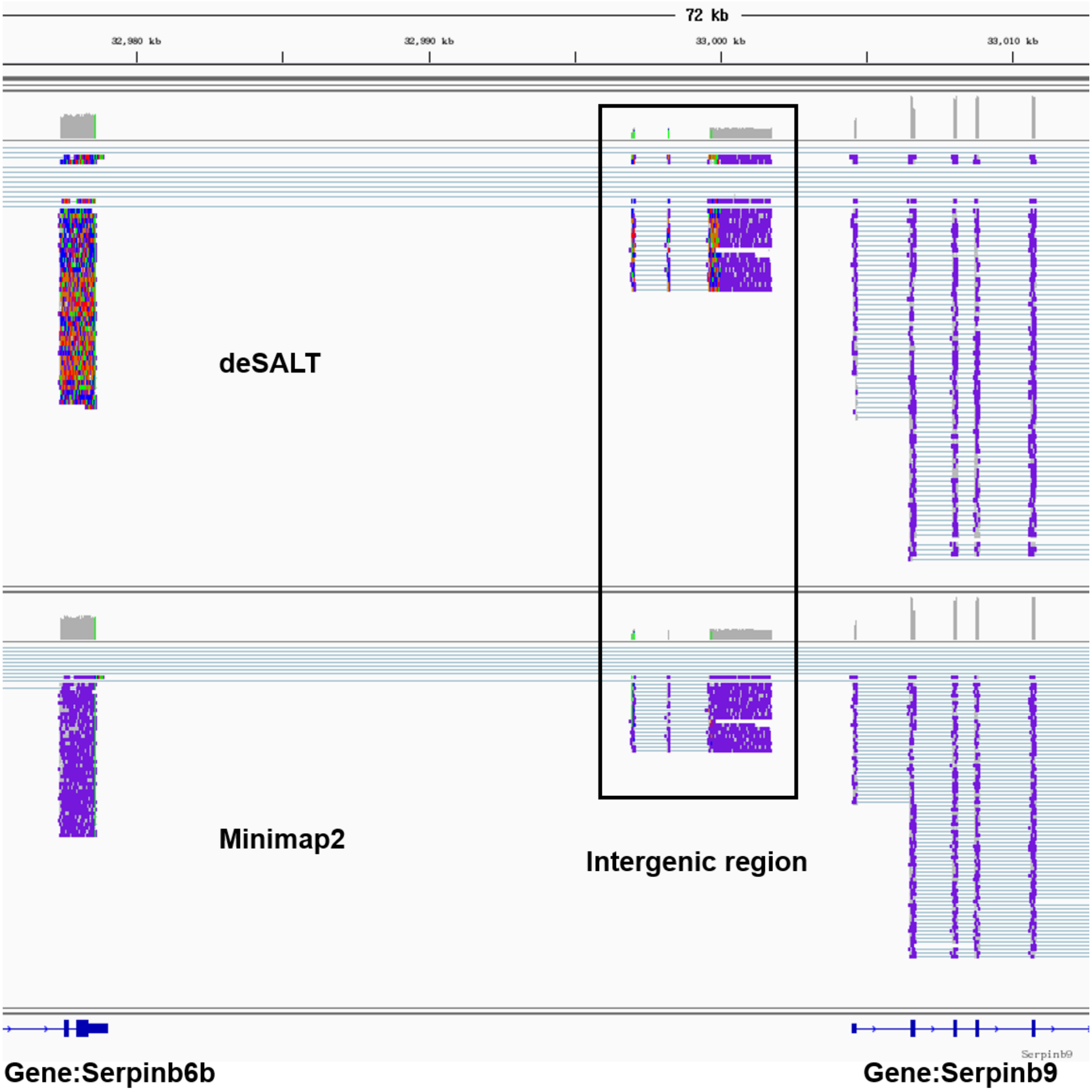

**Supplementary Figure 11. An example of the reads aligned to intergenic regions**

This figure represents the Integrative Genome Viewer (IGV) [1] snapshots of the alignments of the reads from the real mouse PacBio dataset, in the intergenic region between the Serpinb6b gene (Chr13: 32965513–32979037) and Serpinb9 gene (Chr13: 33004541–33017955) of reference GRCm38. It is observed from the alignment results that deSALT and Minimaps2 align 25 and 27 reads to this region, respectively, and the alignments of the reads are also highly similar. This indicates that there could be unannotated transcripts in this region.

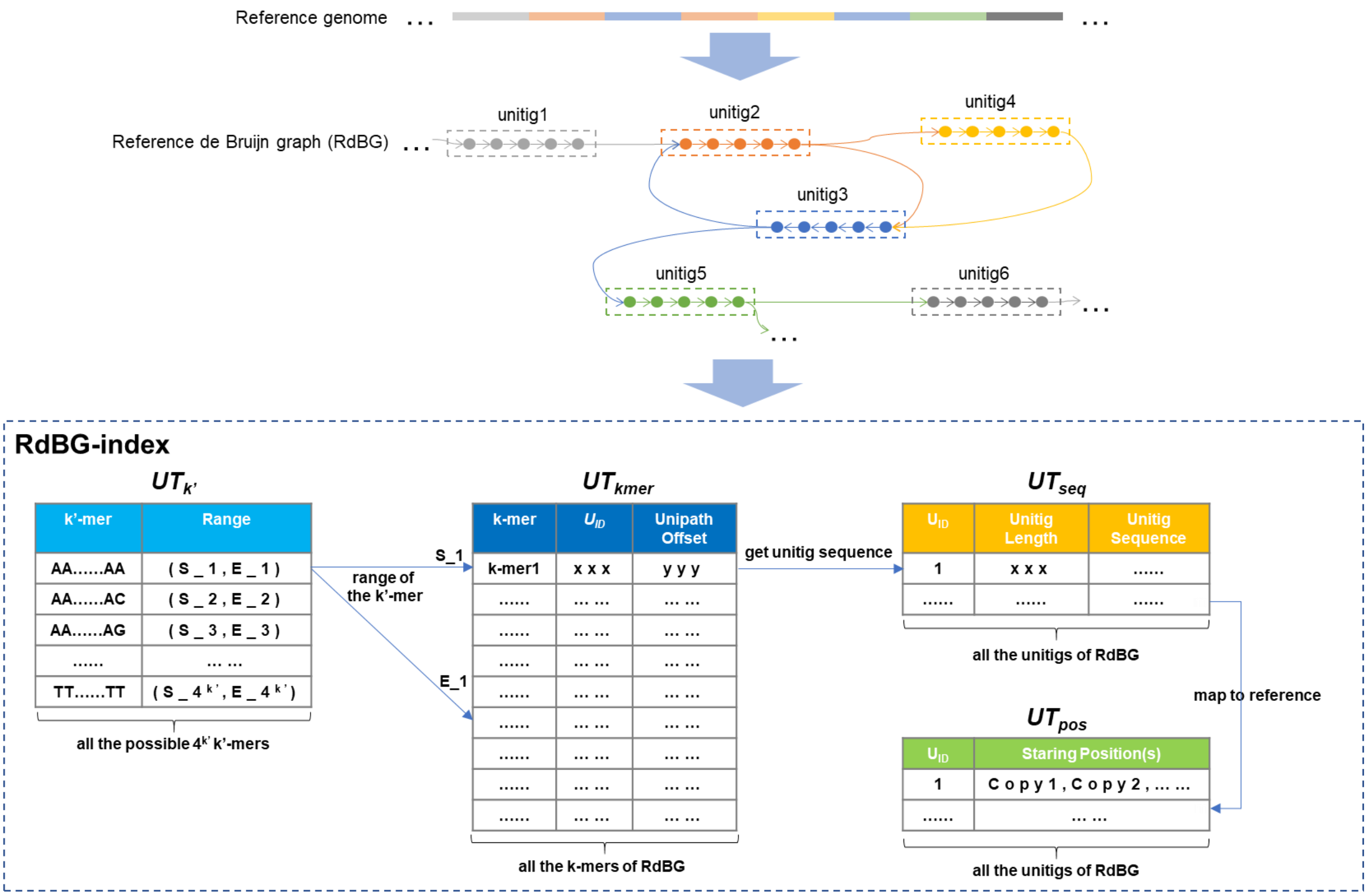

**Supplementary Figure 12. A schematic illustration of the data structures of RdBG-index**

There is a reference genome in the figure (the blocks with various colors indicate the various repeats in the reference sequence). A RdBG is constructed according to the reference sequence (the vertices are represented by the colored round dots). In the RdBG, the repeats are collapsed into 6 unitigs. The RdBG is then indexed by RdBG-index with four major data structures, $UT_{seq}$, $UT_{kmer}$, $UT_{k'}$ and $UT_{pos}$, where $UT_{seq}$, $UT_{kmer}$ and $UT_{pos}$ are linear tables and $UT_{k'}$ is a hash table. The main functions of these data structures are that, $UT_{seq}$ records the sequences and the IDs (called as $U_{ID}$) of the unitigs; $UT_{kmer}$ records the lexicographically sorted list of all the k-mers of RdBG; $UT_{k'}$ records the ranges of the k-mers having identical initial k' charaters in $UT_{kmer}$; and $UT_{pos}$ records the genomic positions of all the copies of the untigis.

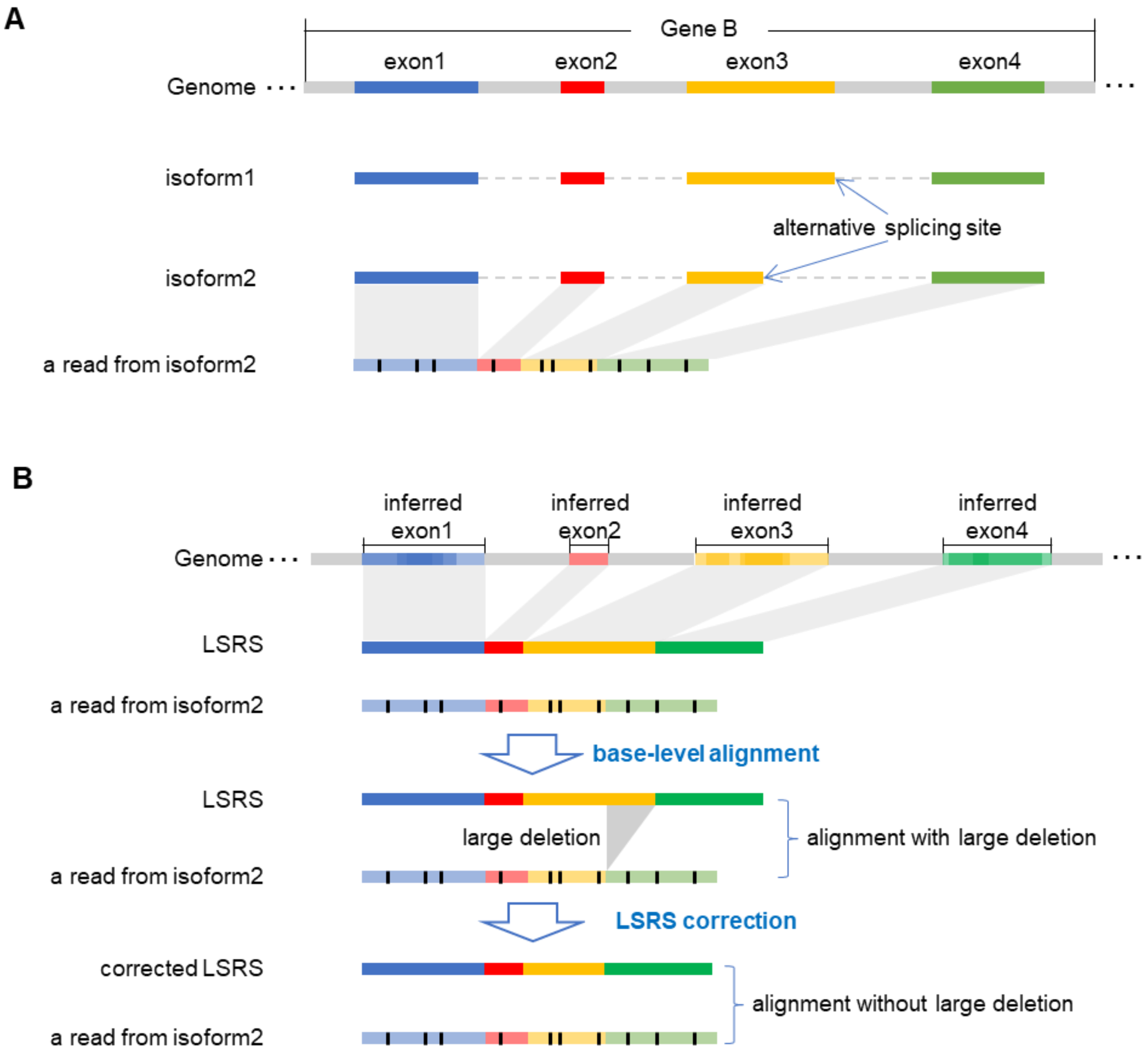

**Supplementary Figure 13. A schematic illustration of the alignment of read against LSRS**

**(A)** There is a gene (called Gene B) having two isoforms which the third exon has an alternative splicing site. And there is a read from isoform2, which has a substring from the alternatively spliced exon3. **(B)** deSALT successfully infers all the four exons of Gene A. With the matched blocks of the read, deSALT composes the LSRS that the whole sequence of exon3 is involved in. Thus, a large deletion occurs in the alignment between the read and the LSRS. deSALT considers it as that there is alternative splicing site around the large deletion. In this situation, it corrects the LSRS by removing the bases corresponding to the large deletion and realigns the read with the corrected LSRS. With these operations, a more accurate alignment is produced.

#### Supplementary Table 1. List of simulated datasets

| No. | Sequencing model [a] | Sequencing error rate [a] | Error ratio [a] | *In silico* transcriptome [b] | Coverage | Average length(bp) | #of reads | #of bases | Reference/ Annotations |
|---|---|---|---|---|---|---|---|---|---|
| 1 | PacBio ROI | 2% | 75:5:20 | Random selected genes | 4X | 6000 | 46783 | 57203247 | GRCh38, version 94 |
| 2 | PacBio ROI | 2% | 75:5:20 | Random selected genes | 10X | 6000 | 103339 | 141769367 | GRCh38, version 94 |
| 3 | PacBio ROI | 2% | 75:5:20 | Random selected genes | 30X | 6000 | 292117 | 424010883 | GRCh38, version 94 |
| 4 | PacBio ROI | 2% | 75:5:20 | All protein coding genes | 4x & 30x | 6000 | 1376093 | 2419998570 | GRCh38, version 94 |
| 5 | PacBio subread | 15% | 1:12:2 | Random selected genes | 4X | 7800 | 46537 | 56809800 | GRCh38, version 94 |
| 6 | PacBio subread | 15% | 1:12:2 | Random selected genes | 10X | 7800 | 102533 | 141326345 | GRCh38, version 94 |
| 7 | PacBio subread | 15% | 1:12:2 | Random selected genes | 30X | 7800 | 289375 | 423841890 | GRCh38, version 94 |
| 8 | PacBio subread | 15% | 1:12:2 | All protein coding genes | 4x & 30x | 7800 | 1390350 | 2415995346 | GRCh38, version 94 |
| 9 | ONT 2D | 13% | 41:23:36 | Random selected genes | 4X | 7800 | 46594 | 56604793 | GRCh38, version 94 |
| 10 | ONT 2D | 13% | 41:23:36 | Random selected genes | 10X | 7800 | 102920 | 141283445 | GRCh38, version 94 |
| 11 | ONT 2D | 13% | 41:23:36 | Random selected genes | 30X | 7800 | 299397 | 423898156 | GRCh38, version 94 |
| 12 | ONT 2D | 13% | 41:23:36 | All protein coding genes | 4x & 30x | 7800 | 1101686 | 2414703724 | GRCh38, version 94 |
| 13 | ONT1D | 25% | 48:15:37 | Random selected genes | 4X | 7800 | 46644 | 56517199 | GRCh38, version 94 |
| 14 | ONT1D | 25% | 48:15:37 | Random selected genes | 10X | 7800 | 103532 | 141296228 | GRCh38, version 94 |
| 15 | ONT1D | 25% | 48:15:37 | Random selected genes | 30X | 7800 | 309135 | 423843578 | GRCh38, version 94 |
| 16 | ONT1D | 25% | 48:15:37 | All protein coding genes | 4x & 30x | 7800 | 1126328 | 2414327781 | GRCh38, version 94 |
| 17 | PS-ONT | -- | -- | Random selected genes | 4X | -- | 47186 | 56649799 | GRCh38, version 94 |
| 18 | PS-ONT | -- | -- | Random selected genes | 10X | -- | 105043 | 141294844 | GRCh38, version 94 |
| 19 | PS-ONT | -- | -- | Random selected genes | 30X | -- | 304133 | 424093591 | GRCh38, version 94 |
| 20 | PS-ONT | -- | -- | All protein coding genes | 4x & 30x | -- | 1367081 | 2624659295 | GRCh38, version 94 |
| 21 | NS-ONT | -- | -- | Random selected genes | 4X | -- | 37000 | 216047531 | GRCh38, version 94 |
| 22 | NS-ONT | -- | -- | Random selected genes | 10X | -- | 90000 | 525306565 | GRCh38, version 94 |
| 23 | NS-ONT | -- | -- | Random selected genes | 30X | -- | 270000 | 1575546383 | GRCh38, version 94 |
| 24 | NS-ONT | -- | -- | All protein coding genes | 4x & 30x | NA | 987268 | 5782765576 | GRCh38, version 94 |
| 25 | PacBio ROI | 2% | 75:5:20 | Random selected genes | 4X | 6000 | 46812 | 66151862 | GRCm38, version 94 |
| 26 | PacBio ROI | 2% | 75:5:20 | Random selected genes | 10X | 6000 | 103498 | 164193767 | GRCm38, version 94 |
| 27 | PacBio ROI | 2% | 75:5:20 | Random selected genes | 30X | 6000 | 292784 | 491418994 | GRCm38, version 94 |
| 28 | PacBio subread | 15% | 1:12:2 | Random selected genes | 4X | 7800 | 46481 | 65756249 | GRCm38, version 94 |
| 29 | PacBio subread | 15% | 1:12:2 | Random selected genes | 10X | 7800 | 102433 | 163803693 | GRCm38, version 94 |
| 30 | PacBio subread | 15% | 1:12:2 | Random selected genes | 30X | 7800 | 289221 | 491278535 | GRCm38, version 94 |
| 31 | ONT 2D | 13% | 41:23:36 | Random selected genes | 4X | 7800 | 46591 | 65584331 | GRCm38, version 94 |
| 32 | ONT 2D | 13% | 41:23:36 | Random selected genes | 10X | 7800 | 102879 | 163762584 | GRCm38, version 94 |

| | | | | | | | | | |
|---|---|---|---|---|---|---|---|---|---|
| 33 | | 13% | 41:23:36 | | 30X | 7800 | 299413 | 491329071 | |
| 34 | ONT1D | 25% | 48:15:37 | Random selected genes | 4X | 7800 | 46600 | 65508866 | |
| 35 | | 25% | 48:15:37 | | 10X | 7800 | 103647 | 163775669 | |
| 36 | | 25% | 48:15:37 | | 30X | 7800 | 309276 | 491282118 | |
| 37 | PS-ONT | -- | -- | Random selected genes | 4X | -- | 47348 | 65619390 | |
| 38 | | -- | -- | | 10X | -- | 105717 | 163772057 | |
| 39 | | -- | -- | | 30X | -- | 305944 | 491502093 | |
| 40 | NS-ONT | -- | -- | Random selected genes | 4X | -- | 37000 | 216240439 | |
| 41 | | -- | -- | | 10X | -- | 90000 | 526100252 | |
| 42 | | -- | -- | | 30X | -- | 270000 | 1575147063 | |
| 43 | PacBio ROI | 2% | 75:5:20 | Random selected genes | 4X | 6000 | 45296 | 102311172 | BDGP6, version 94 |
| 44 | | 2% | 75:5:20 | | 10X | 6000 | 101290 | 254849528 | |
| 45 | | 2% | 75:5:20 | | 30X | 6000 | 288355 | 763849030 | |
| 46 | PacBio subread | 15% | 1:12:2 | Random selected genes | 4X | 7800 | 44169 | 101945130 | |
| 47 | | 15% | 1:12:2 | | 10X | 7800 | 97910 | 254613734 | |
| 48 | | 15% | 1:12:2 | | 30X | 7800 | 277541 | 763798783 | |
| 49 | ONT 2D | 13% | 41:23:36 | Random selected genes | 4X | 7800 | 44445 | 101867685 | |
| 50 | | 13% | 41:23:36 | | 10X | 7800 | 98980 | 254601058 | |
| 51 | | 13% | 41:23:36 | | 30X | 7800 | 288337 | 763814989 | |
| 52 | ONT1D | 25% | 48:15:37 | Random selected genes | 4X | 7800 | 44546 | 101842292 | |
| 53 | | 25% | 48:15:37 | | 10X | 7800 | 100127 | 254608663 | |
| 54 | | 25% | 48:15:37 | | 30X | 7800 | 297821 | 763802581 | |
| 55 | PS-ONT | -- | -- | Random selected genes | 4X | -- | 46305 | 101883320 | |
| 56 | | -- | -- | | 10X | -- | 104775 | 254610343 | |
| 57 | | -- | -- | | 30X | -- | 304510 | 763907660 | |
| 58 | NS-ONT | -- | -- | Random selected genes | 4X | -- | 34000 | 198110605 | |
| 59 | | -- | -- | | 10X | -- | 86000 | 503566482 | |
| 60 | | -- | -- | | 30X | -- | 250000 | 1460343906 | |

a) “PacBio ROI”, “PacBio Subread”, “ONT 2D” and “ONT 1D” models are generated by PBSim with fixed parameters, and the total sequencing error rates and the ratios of the sequencing errors (represented as mismatches: insertions: deletions) are configured by referring to previous studies. More precisely, the error models of the PacBio ROI datasets, ONT 2D datasets, and ONT1D datasets are configured by referring to the previous study conducted by Weirather *et al.* [2], and the error model of the PacBio subread datasets are configured by referring to the previous study conducted by Carneiro *et al.* [3]. “PS-ONT” and “NS-ONT” models are respectively generated by PBSim and NanoSim based on a real ONT dataset. Also refer to the Supplementary Notes for the used command lines of the simulators.

b) “Radomly selected genes” indicates the dataset was simulated from a set of randomly selected genes of some species (human, mouse or fruit fly), i.e., the H-se, M-se or F-se transcriptome, and “All protein coding genes” indicates that the dataset was simulated from all the protein coding genes of human, i.e., the H-all transcriptome.

#### Supplementary Table 2. Overall benchmark results on the simulated datasets [a]

| Aligner | Parameters [b] | Base% [c] | Exons% [d] | Read80% [e] | Read100% [f] |
|---|---|---|---|---|---|
| | | **Simulated PacBio ROI datasets** | | | |
| | | **Fruit fly PacBio ROI, coverage = 4X, 45296 reads/102311172 bases/230387 exons** | | | |
| GMAP | -f samse --cross-species -z sense_force | 99.63 | 96.47 | 98.32 | 88.83 |
| Graphmap2 | -x rnaseq | 99.09 | 96.51 | 98.44 | 89.16 |
| Minimap2 | -ax splice | 99.55 | 98.01 | 98.75 | 92.78 |
| Minimap2 + GTF | -ax splice –junc-bed anno.bed | 99.55 | 98.76 | 99.03 | 94.51 |
| deSALT | -d 10 -x ccs | 99.66 | 98.67 | 99.55 | 94.95 |
| deSALT + GTF | -d 10 -x ccs -G anno.info | 99.59 | 98.74 | 99.44 | 95.03 |
| | | **Fruit fly PacBio ROI, coverage = 10X, 101290 reads/254849528 bases/561473 exons** | | | |
| GMAP | -f samse --cross-species -z sense_force | 99.63 | 96.62 | 98.11 | 88.5 |
| Graphmap2 | -x rnaseq | 99.06 | 96.64 | 98.43 | 89.14 |
| Minimap2 | -ax splice | 99.57 | 98.33 | 99.15 | 93.32 |
| Minimap2 + GTF | -ax splice –junc-bed anno.bed | 99.58 | 99.03 | 99.44 | 95.28 |
| deSALT | -d 10 -x ccs | 99.67 | 98.91 | 99.59 | 95.44 |
| deSALT + GTF | -d 10 -x ccs -G anno.info | 99.62 | 99.04 | 99.54 | 95.33 |
| | | **Fruit fly PacBio ROI, coverage = 30X, 288355 reads/763849030 bases/1666790 exons** | | | |
| GMAP | -f samse --cross-species -z sense_force | 99.64 | 96.62 | 98.01 | 88.1 |
| Graphmap2 | -x rnaseq | 99.06 | 96.69 | 98.39 | 88.98 |
| Minimap2 | -ax splice | 99.62 | 98.44 | 99.4 | 93.49 |
| Minimap2 + GTF | -ax splice –junc-bed anno.bed | 99.63 | 99.14 | 99.7 | 95.58 |
| deSALT | -d 10 -x ccs | 99.69 | 99.02 | 99.61 | 95.66 |
| deSALT + GTF | -d 10 -x ccs -G anno.info | 99.64 | 99.13 | 99.59 | 95.7 |
| | | **Mouse PacBio ROI, coverage = 4X, 46812 reads/66151862 bases/232811 exons** | | | |
| GMAP | -f samse --cross-species -z sense_force | 98.31 | 97.16 | 97.96 | 90.25 |
| Graphmap2 | -x rnaseq | 94.02 | 88.88 | 92.95 | 79.04 |
| Minimap2 | -ax splice | 98.25 | 95.88 | 96.25 | 89.55 |
| Minimap2 + GTF | -ax splice –junc-bed anno.bed | 98.75 | 97.4 | 97.32 | 91.44 |
| deSALT | -d 10 -x ccs | 98.47 | 97.28 | 97.67 | 91.84 |

| | | | | | |
|---|---|---|---|---|---|
| deSALT + GTF | -d 10 -x ccs -G anno.info | 98.41 | 97.11 | 97.34 | 90.27 |
| **Mouse PacBio ROI, coverage = 10X, 103498 reads/164193767 bases/564165 exons** | | | | | |
| GMAP | -f samse --cross-species -z sense_force | 98.3 | 94.68 | 96 | 84.27 |
| Graphmap2 | -x rnaseq | 93.97 | 89.02 | 92.76 | 78.94 |
| Minimap2 | -ax splice | 98.37 | 96.47 | 96.99 | 90.88 |
| Minimap2 + GTF | -ax splice –junc-bed anno.bed | 98.87 | 97.95 | 98.1 | 92.9 |
| deSALT | -d 10 -x ccs | 98.57 | 97.68 | 97.87 | 92.74 |
| deSALT + GTF | -d 10 -x ccs -G anno.info | 98.49 | 97.58 | 97.6 | 92.96 |
| **Mouse PacBio ROI, coverage = 30X, 292784 reads/491418994 bases/1669598 exons** | | | | | |
| GMAP | -f samse --cross-species -z sense_force | 98.32 | 94.8 | 95.93 | 84.29 |
| Graphmap2 | -x rnaseq | 93.98 | 89.22 | 92.6 | 78.96 |
| Minimap2 | -ax splice | 98.38 | 96.72 | 97.29 | 91.52 |
| Minimap2 + GTF | -ax splice –junc-bed anno.bed | 98.88 | 98.17 | 98.45 | 93.64 |
| deSALT | -d 10 -x ccs | 98.56 | 97.88 | 97.95 | 93.22 |
| deSALT + GTF | -d 10 -x ccs -G anno.info | 98.57 | 97.92 | 97.96 | 93.47 |
| **Human PacBio ROI, coverage = 4X, 46783 reads/57203247 bases/222656 exons** | | | | | |
| GMAP | -f samse --cross-species -z sense_force | 98.9 | 95.14 | 96.97 | 86.51 |
| Graphmap2 | -x rnaseq | 95.86 | 90.96 | 94.32 | 80.92 |
| Minimap2 | -ax splice | 98.65 | 96.48 | 96.8 | 90.81 |
| Minimap2 + GTF | -ax splice –junc-bed anno.bed | 99.11 | 97.32 | 97.72 | 91.76 |
| deSALT | -d 10 -x ccs | 99.19 | 97.81 | 98.38 | 93.44 |
| deSALT + GTF | -d 10 -x ccs -G anno.info | 99.23 | 97.9 | 98.44 | 93.75 |
| **Human PacBio ROI, coverage = 10X, 103339 reads/141769367 bases/537916 exons** | | | | | |
| GMAP | -f samse --cross-species -z sense_force | 98.96 | 95.39 | 96.81 | 86.23 |
| Graphmap2 | -x rnaseq | 95.76 | 91.15 | 94.21 | 80.93 |
| Minimap2 | -ax splice | 98.77 | 97.16 | 97.59 | 92.2 |
| Minimap2 + GTF | -ax splice –junc-bed anno.bed | 99.19 | 97.84 | 98.45 | 92.97 |
| deSALT | -d 10 -x ccs | 99.22 | 98.23 | 98.57 | 94.34 |
| deSALT + GTF | -d 10 -x ccs -G anno.info | 99.26 | 98.32 | 98.59 | 94.63 |
| **Human PacBio ROI, coverage = 30X, 292117 reads/424010883 bases/1591417 exons** | | | | | |
| GMAP | -f samse --cross-species -z sense_force | 98.96 | 95.46 | 96.79 | 86.37 |
| Graphmap2 | -x rnaseq | 95.82 | 91.26 | 94.15 | 80.91 |

| | | | | | |
|---|---|---|---|---|---|
| Minimap2 | -ax splice | 98.8 | 97.36 | 98 | 92.95 |
| Minimap2 + GTF | -ax splice –junc-bed anno.bed | 99.24 | 98.05 | 98.8 | 93.56 |
| deSALT | -d 10 -x ccs | 99.25 | 98.39 | 98.71 | 94.91 |
| deSALT + GTF | -d 10 -x ccs -G anno.info | 99.28 | 98.48 | 98.7 | 95.29 |
| **Human all protein coding genes PacBio ROI, 1376093 reads/ 2419998570 bases/ 8750772 exons** | | | | | |
| GMAP | -f samse --cross-species -z sense_force | 99.45 | 95.58 | 97.21 | 84.39 |
| Graphmap2 | -x rnaseq | 95.97 | 90.15 | 93.49 | 76.76 |
| Minimap2 | -ax splice | 99.29 | 97.13 | 98.4 | 90.67 |
| Minimap2 + GTF | -ax splice –junc-bed anno.bed | 99.59 | 97.98 | 99.03 | 92.16 |
| deSALT | -d 10 -x ccs | 99.76 | 98.27 | 99.21 | 92.19 |
| deSALT + GTF | -d 10 -x ccs -G anno.info | 99.76 | 98.37 | 99.26 | 92.66 |
| **Simulated ONT 2D ($1D^2$) datasets** | | | | | |
| **Fruit fly ONT 2D ($1D^2$), coverage = 4X, 44445 reads/101867685 bases/229107 exons** | | | | | |
| GMAP | -f samse --cross-species -z sense_force | 94.26 | 74.17 | 74.08 | 43.78 |
| Graphmap2 | -x rnaseq | 95.01 | 91.83 | 95.28 | 73.01 |
| Minimap2 | -ax splice | 95.14 | 92.64 | 91.47 | 72.43 |
| Minimap2 + GTF | -ax splice –junc-bed anno.bed | 95.21 | 94.71 | 92.81 | 77.75 |
| deSALT | -d 10 -x ont2d -s 2 | 95.63 | 95.28 | 96.45 | 80.43 |
| deSALT + GTF | -d 10 -x ont2d -s 2 -G anno.info | 95.42 | 96.09 | 97.31 | 83.55 |
| **Fruit fly ONT 2D ($1D^2$), coverage = 10X, 98980 reads/254601058 bases/560709 exons** | | | | | |
| GMAP | -f samse --cross-species -z sense_force | 94.46 | 74.86 | 73.28 | 42.3 |
| Graphmap2 | -x rnaseq | 95.02 | 93.26 | 96.67 | 75.52 |
| Minimap2 | -ax splice | 95.45 | 94.08 | 95.87 | 75.59 |
| Minimap2 + GTF | -ax splice –junc-bed anno.bed | 95.52 | 96.07 | 97.31 | 81.46 |
| deSALT | -d 10 -x ont2d -s 2 | 95.76 | 96.79 | 98.1 | 84.62 |
| deSALT + GTF | -d 10 -x ont2d -s 2 -G anno.info | 95.79 | 97.26 | 98.5 | 86.71 |
| **Fruit fly ONT 2D ($1D^2$), coverage = 30X, 288337 reads/763814989 bases/1670965 exons** | | | | | |
| GMAP | -f samse --cross-species -z sense_force | 94.45 | 74.79 | 72.76 | 41.83 |
| Graphmap2 | -x rnaseq | 95.08 | 93.76 | 96.99 | 77.01 |
| Minimap2 | -ax splice | 95.48 | 94.29 | 95.9 | 76.1 |

| | | | | | |
|---|---|---|---|---|---|
| Minimap2 + GTF | -ax splice –junc-bed anno.bed | 95.55 | 96.28 | 97.35 | 81.99 |
| deSALT | -d 10 -x ont2d -s 2 | 95.8 | 97.28 | 98.47 | 86.7 |
| deSALT + GTF | -d 10 -x ont2d -s 2 -G anno.info | 95.82 | 97.57 | 98.62 | 87.85 |
| **Mouse ONT 2D (1D$^2$), coverage = 4X, 46591 reads/65584331 bases/232141 exons** | | | | | |
| GMAP | -f samse --cross-species -z sense_force | 90.51 | 68.52 | 67.98 | 40.13 |
| Graphmap2 | -x rnaseq | 89.4 | 80.36 | 84.88 | 57.94 |
| Minimap2 | -ax splice | 91.92 | 85.88 | 81.23 | 59.36 |
| Minimap2 + GTF | -ax splice –junc-bed anno.bed | 92.51 | 88.27 | 83.21 | 62.52 |
| deSALT | -d 10 -x ont2d -s 2 | 93.64 | 91.5 | 91.1 | 72.01 |
| deSALT + GTF | -d 10 -x ont2d -s 2 -G anno.info | 93.76 | 92.65 | 92.3 | 75.73 |
| **Mouse ONT 2D (1D$^2$), coverage = 10X, 102879 reads/163762584 bases/565884 exons** | | | | | |
| GMAP | -f samse --cross-species -z sense_force | 90.84 | 69.48 | 67.89 | 39.59 |
| Graphmap2 | -x rnaseq | 89.48 | 82.47 | 86.79 | 60.76 |
| Minimap2 | -ax splice | 92.59 | 88.31 | 88.24 | 64.64 |
| Minimap2 + GTF | -ax splice –junc-bed anno.bed | 93.18 | 90.69 | 90.36 | 68.13 |
| deSALT | -d 10 -x ont2d -s 2 | 94.09 | 93.77 | 94.26 | 77.73 |
| deSALT + GTF | -d 10 -x ont2d -s 2 -G anno.info | 94.15 | 94.37 | 94.81 | 80.08 |
| **Mouse ONT 2D (1D$^2$), coverage = 30X, 299413 reads/491329071 bases/1688076 exons** | | | | | |
| GMAP | -f samse --cross-species -z sense_force | 90.92 | 69.46 | 67.28 | 39.42 |
| Graphmap2 | -x rnaseq | 89.51 | 82.89 | 87.09 | 61.48 |
| Minimap2 | -ax splice | 92.68 | 88.57 | 88.57 | 65 |
| Minimap2 + GTF | -ax splice –junc-bed anno.bed | 93.28 | 91.08 | 90.81 | 68.64 |
| deSALT | -d 10 -x ont2d -s 2 | 94.08 | 94.12 | 94.58 | 79.06 |
| deSALT + GTF | -d 10 -x ont2d -s 2 -G anno.info | 94.12 | 94.54 | 94.91 | 80.69 |
| **Human ONT 2D (1D$^2$), coverage = 4X, 46594 reads/56604793 bases/223025 exons** | | | | | |
| GMAP | -f samse --cross-species -z sense_force | 90.7 | 68.99 | 67.62 | 40.72 |
| Graphmap2 | -x rnaseq | 89.19 | 80.62 | 84.39 | 58.24 |
| Minimap2 | -ax splice | 91.77 | 85.57 | 79.95 | 58.76 |
| Minimap2 + GTF | -ax splice –junc-bed anno.bed | 92.31 | 87.46 | 81.45 | 61.46 |
| deSALT | -d 10 -x ont2d -s 2 | 94.36 | 91.81 | 91.11 | 72.83 |
| deSALT + GTF | -d 10 -x ont2d -s 2 -G anno.info | 94.48 | 93.01 | 92.55 | 76.51 |
| **Human ONT 2D (1D$^2$), coverage = 10X, 102920 reads/141283445 bases/543017 exons** | | | | | |

| | | | | | |
|---|---|---|---|---|---|
| GMAP | -f samse --cross-species -z sense_force | 90.88 | 69.22 | 67.14 | 40.21 |
| Graphmap2 | -x rnaseq | 89.34 | 82.38 | 86.29 | 60.31 |
| Minimap2 | -ax splice | 92.57 | 87.83 | 87.85 | 64.07 |
| Minimap2 + GTF | -ax splice –junc-bed anno.bed | 93.11 | 89.91 | 89.61 | 67.13 |
| deSALT | -d 10 -x ont2d -s 2 | 94.78 | 93.86 | 94.84 | 78.28 |
| deSALT + GTF | -d 10 -x ont2d -s 2 -G anno.info | 94.86 | 94.64 | 95.59 | 81.05 |
| **Human ONT 2D (1D$^2$), coverage = 30X, 299397 reads/423898156 bases/1615900 exons** | | | | | |
| GMAP | -f samse --cross-species -z sense_force | 90.96 | 79.05 | 77.04 | 50.44 |
| Graphmap2 | -x rnaseq | 89.35 | 82.5 | 86.18 | 60.77 |
| Minimap2 | -ax splice | 92.69 | 88.29 | 88.03 | 64.53 |
| Minimap2 + GTF | -ax splice –junc-bed anno.bed | 93.22 | 90.33 | 89.77 | 67.74 |
| deSALT | -d 10 -x ont2d -s 2 | 94.85 | 94.33 | 95.06 | 79.63 |
| deSALT + GTF | -d 10 -x ont2d -s 2 -G anno.info | 94.91 | 94.92 | 95.63 | 81.6 |
| **Human all protein coding genes ONT 2D (1D$^2$), 1101686 reads/ 2414703724 bases/ 8422080 exons** | | | | | |
| GMAP | -f samse --cross-species -z sense_force | 94.33 | 70.48 | 61.57 | 33.45 |
| Graphmap2 | -x rnaseq | 91.5 | 80.96 | 83.03 | 54.03 |
| Minimap2 | -ax splice | 96.91 | 91.44 | 91.18 | 66.52 |
| Minimap2 + GTF | -ax splice –junc-bed anno.bed | 97.29 | 93.31 | 92.91 | 70.41 |
| deSALT | -d 10 -x ont2d -s 2 | 98.14 | 95.71 | 95.93 | 79.59 |
| deSALT + GTF | -d 10 -x ont2d -s 2 -G anno.info | 98.15 | 96.18 | 96.31 | 81.47 |
| **Simulated PacBio subread datasets** | | | | | |
| **Fruit fly PacBio subread, coverage = 4X, 44169 reads/101945130 bases/212417 exons** | | | | | |
| GMAP | -f samse --cross-species -z sense_force | 81.93 | 62.44 | 52.23 | 28.47 |
| Graphmap2 | -x rnaseq | 87.84 | 90.03 | 94.07 | 68.29 |
| Minimap2 | -ax splice | 87.7 | 90.76 | 85.07 | 65.73 |
| Minimap2 + GTF | -ax splice –junc-bed anno.bed | 87.79 | 92.84 | 86.53 | 71.33 |
| deSALT | -d 10 -x clr -s 2 | 88.39 | 95.01 | 95.71 | 78.98 |
| deSALT + GTF | -d 10 -x clr -s 2 -G anno.info | 88.42 | 95.62 | 96.29 | 81.12 |
| **Fruit fly PacBio subread, coverage = 10X, 97910 reads/254613734 bases/518798 exons** | | | | | |
| GMAP | -f samse --cross-species -z sense_force | 82.13 | 62.94 | 51.9 | 27.45 |

| | | | | | |
|---|---|---|---|---|---|
| Graphmap2 | -x rnaseq | 87.87 | 91.93 | 96.13 | 71.7 |
| Minimap2 | -ax splice | 88.15 | 92.95 | 93.98 | 72.28 |
| Minimap2 + GTF | -ax splice –junc-bed anno.bed | 88.24 | 95.21 | 95.52 | 78.65 |
| deSALT | -d 10 -x clr -s 2 | 88.55 | 96.4 | 97.83 | 83.32 |
| deSALT + GTF | -d 10 -x clr -s 2 -G anno.info | 88.56 | 96.71 | 98.09 | 84.55 |
| **Fruit fly PacBio subread, coverage = 30X, 277541 reads/763798783 bases/1540635 exons** | | | | | |
| GMAP | -f samse --cross-species -z sense_force | 82.58 | 85.25 | 52.79 | 27.41 |
| Graphmap2 | -x rnaseq | 87.94 | 92.72 | 96.58 | 73.7 |
| Minimap2 | -ax splice | 88.3 | 93.68 | 96 | 74.27 |
| Minimap2 + GTF | -ax splice –junc-bed anno.bed | 88.38 | 95.94 | 97.57 | 80.84 |
| deSALT | -d 10 -x clr -s 2 | 88.58 | 96.92 | 98.34 | 84.78 |
| deSALT + GTF | -d 10 -x clr -s 2 -G anno.info | 88.59 | 97.16 | 98.44 | 85.76 |
| **Mouse PacBio subread, coverage = 4X, 46481 reads/65756249 bases/216580 exons** | | | | | |
| GMAP | -f samse --cross-species -z sense_force | 74.2 | 52.37 | 42.76 | 24.27 |
| Graphmap2 | -x rnaseq | 83.03 | 78.73 | 84.16 | 54.31 |
| Minimap2 | -ax splice | 84.5 | 83.3 | 75.27 | 52.45 |
| Minimap2 + GTF | -ax splice –junc-bed anno.bed | 85.04 | 85.77 | 77.15 | 55.51 |
| deSALT | -d 10 -x clr -s 2 | 86.45 | 90.06 | 88.57 | 67.79 |
| deSALT + GTF | -d 10 -x clr -s 2 -G anno.info | 86.53 | 91.19 | 89.86 | 71.17 |
| **Mouse PacBio subread, coverage = 10X, 102433 reads/163803693 bases/526135 exons** | | | | | |
| GMAP | -f samse --cross-species -z sense_force | 74.2 | 52.55 | 41.02 | 22.68 |
| Graphmap2 | -x rnaseq | 83.22 | 81.03 | 86.29 | 56.39 |
| Minimap2 | -ax splice | 85.18 | 85.98 | 84.42 | 58.65 |
| Minimap2 + GTF | -ax splice –junc-bed anno.bed | 85.75 | 88.62 | 86.63 | 62.2 |
| deSALT | -d 10 -x clr -s 2 | 86.89 | 92.56 | 92.89 | 73.95 |
| deSALT + GTF | -d 10 -x clr -s 2 -G anno.info | 86.93 | 93.16 | 93.56 | 76.08 |
| **Mouse PacBio subread, coverage = 30X, 289221 reads/491278535 bases/1559863 exons** | | | | | |
| GMAP | -f samse --cross-species -z sense_force | 74.7 | 53.32 | 40.85 | 22.39 |
| Graphmap2 | -x rnaseq | 83.26 | 81.83 | 87.08 | 57.63 |
| Minimap2 | -ax splice | 85.49 | 87.22 | 87.99 | 60.97 |
| Minimap2 + GTF | -ax splice –junc-bed anno.bed | 86.04 | 89.74 | 90.26 | 64.69 |
| deSALT | -d 10 -x clr -s 2 | 86.98 | 93.5 | 94.49 | 76.25 |

| | | | | | |
|---|---|---|---|---|---|
| deSALT + GTF | -d 10 -x clr -s 2 -G anno.info | 87 | 93.9 | 94.79 | 77.68 |
| **Human PacBio subread, coverage = 4X, 46537 reads/56809800 bases/208419 exons** | | | | | |
| GMAP | -f samse --cross-species -z sense_force | 71.43 | 49.52 | 39.77 | 22.84 |
| Graphmap2 | -x rnaseq | 83.79 | 79.54 | 85 | 55.48 |
| Minimap2 | -ax splice | 84.29 | 82.65 | 73.78 | 51.12 |
| Minimap2 + GTF | -ax splice –junc-bed anno.bed | 84.75 | 84.16 | 79.3 | 55.36 |
| deSALT | -d 10 -x clr -s 2 | 86.98 | 90.05 | 88.24 | 68.13 |
| deSALT + GTF | -d 10 -x clr -s 2 -G anno.info | 87.12 | 91.33 | 89.84 | 71.52 |
| **Human PacBio subread, coverage = 10X, 102533 reads/141326345 bases/504951 exons** | | | | | |
| GMAP | -f samse --cross-species -z sense_force | 71.47 | 49.79 | 37.66 | 21.19 |
| Graphmap2 | -x rnaseq | 83.92 | 81.27 | 86.83 | 56.95 |
| Minimap2 | -ax splice | 85.13 | 85.64 | 83.57 | 57.45 |
| Minimap2 + GTF | -ax splice –junc-bed anno.bed | 85.4 | 86.42 | 86.53 | 59.84 |
| deSALT | -d 10 -x clr -s 2 | 87.57 | 92.84 | 93.44 | 74.29 |
| deSALT + GTF | -d 10 -x clr -s 2 -G anno.info | 87.64 | 93.59 | 94.32 | 76.62 |
| **Human PacBio subread, coverage = 30X, 289375 reads/423841890 bases/1496976 exons** | | | | | |
| GMAP | -f samse --cross-species -z sense_force | 71.75 | 50.28 | 37.04 | 20.8 |
| Graphmap2 | -x rnaseq | 83.98 | 81.97 | 87.41 | 57.99 |
| Minimap2 | -ax splice | 85.48 | 86.92 | 87.58 | 60.13 |
| Minimap2 + GTF | -ax splice –junc-bed anno.bed | 85.68 | 87.51 | 89.62 | 62.06 |
| deSALT | -d 10 -x clr -s 2 | 87.76 | 93.85 | 95.39 | 76.87 |
| deSALT + GTF | -d 10 -x clr -s 2 -G anno.info | 87.8 | 94.4 | 95.94 | 78.75 |
| **Human all protein coding genes PacBio subread, 1390350 reads/ 2415995346 bases/ 8156901 exons** | | | | | |
| GMAP | -f samse --cross-species -z sense_force | 77.36 | 55.05 | 45.11 | 24.01 |
| Graphmap2 | -x rnaseq | 84.84 | 80.03 | 83.54 | 50.98 |
| Minimap2 | -ax splice | 87.25 | 88.3 | 90.53 | 60.74 |
| Minimap2 + GTF | -ax splice –junc-bed anno.bed | 87.57 | 90.11 | 92.1 | 63.64 |
| deSALT | -d 10 -x clr -s 2 | 88.74 | 94.12 | 95.99 | 75.84 |
| deSALT + GTF | -d 10 -x clr -s 2 -G anno.info | 88.74 | 94.64 | 96.33 | 77.54 |
| **Simulated ONT 1D datasets** | | | | | |

| | | | | | |
|---|---|---|---|---|---|
| **Fruit fly ONT 1D, coverage = 4X, 44546 reads/101842292 bases/240905 exons** | | | | | |
| GMAP | -f samse --cross-species -z sense_force | 64.73 | 20.37 | 13.58 | 3.4 |
| Graphmap2 | -x rnaseq | 94.32 | 78.28 | 79.21 | 38.8 |
| Minimap2 | -ax splice | 89.58 | 75.05 | 64.21 | 33.72 |
| Minimap2 + GTF | -ax splice –junc-bed anno.bed | 89.98 | 80.88 | 70.58 | 44.36 |
| deSALT | -d 10 -x ont1d -s 2 -l 14 | 95.54 | 84.89 | 82.46 | 47.1 |
| deSALT + GTF | -d 10 -x ont1d -s 2 -l 14 -G anno.info | 95.94 | 89.32 | 89.39 | 58.13 |
| **Fruit fly ONT 1D, coverage = 10X, 100127 reads/254608663 bases/589616 exons** | | | | | |
| GMAP | -f samse --cross-species -z sense_force | 65.95 | 20.86 | 13.25 | 3.25 |
| Graphmap2 | -x rnaseq | 94.39 | 81.92 | 83.86 | 44.03 |
| Minimap2 | -ax splice | 90.98 | 77.97 | 69.81 | 36.9 |
| Minimap2 + GTF | -ax splice –junc-bed anno.bed | 91.39 | 83.97 | 76.75 | 48.9 |
| deSALT | -d 10 -x ont1d -s 2 -l 14 | 96.24 | 89.12 | 88 | 55.5 |
| deSALT + GTF | -d 10 -x ont1d -s 2 -l 14 -G anno.info | 96.47 | 92.51 | 93.29 | 66.57 |
| **Fruit fly ONT 1D, coverage = 30X, 297821 reads/763802581 bases/1764192 exons** | | | | | |
| GMAP | -f samse --cross-species -z sense_force | 66.01 | 20.91 | 13.24 | 3.2 |
| Graphmap2 | -x rnaseq | 94.41 | 82.95 | 85.03 | 46.67 |
| Minimap2 | -ax splice | 91.05 | 78.03 | 69.96 | 37.11 |
| Minimap2 + GTF | -ax splice –junc-bed anno.bed | 91.46 | 84.1 | 76.93 | 49.1 |
| deSALT | -d 10 -x ont1d -s 2 -l 14 | 96.43 | 90.94 | 91.28 | 61.83 |
| deSALT + GTF | -d 10 -x ont1d -s 2 -l 14 -G anno.info | 96.56 | 93.23 | 94.1 | 69.45 |
| **Mouse ONT 1D, coverage = 4X, 46600 reads/65508866 bases/243889 exons** | | | | | |
| GMAP | -f samse --cross-species -z sense_force | 53.66 | 12.83 | 13.09 | 2.86 |
| Graphmap2 | -x rnaseq | 79.57 | 47.85 | 47.8 | 18.51 |
| Minimap2 | -ax splice | 75.32 | 52.55 | 39.15 | 16.76 |
| Minimap2 + GTF | -ax splice –junc-bed anno.bed | 76.37 | 57.95 | 43.14 | 20.65 |
| deSALT | -d 10 -x ont1d -s 2 -l 14 | 86.68 | 65.86 | 57.44 | 25.43 |
| deSALT + GTF | -d 10 -x ont1d -s 2 -l 14 -G anno.info | 88.04 | 74.35 | 68.35 | 36.75 |
| **Mouse ONT 1D, coverage = 10X, 103647 reads/163775669 bases/595736 exons** | | | | | |
| GMAP | -f samse --cross-species -z sense_force | 54.88 | 13.51 | 12.34 | 2.66 |
| Graphmap2 | -x rnaseq | 79.97 | 51.47 | 50.53 | 20.41 |
| Minimap2 | -ax splice | 77.03 | 55.27 | 42.83 | 18.66 |

| | | | | | |
|---|---|---|---|---|---|
| Minimap2 + GTF | -ax splice –junc-bed anno.bed | 78.12 | 61.03 | 47.54 | 23.2 |
| deSALT | -d 10 -x ont1d -s 2 -l 14 | 89.22 | 74.21 | 67.75 | 34.25 |
| deSALT + GTF | -d 10 -x ont1d -s 2 -l 14 -G anno.info | 89.98 | 80.11 | 76.05 | 45.76 |
| **Mouse ONT 1D, coverage = 30X, 309276 reads/491282118 bases/1783381 exons** | | | | | |
| GMAP | -f samse --cross-species -z sense_force | 35.45 | 8.14 | 9.68 | 2.02 |
| Graphmap2 | -x rnaseq | 80.03 | 52.88 | 51.99 | 21.69 |
| Minimap2 | -ax splice | 77.13 | 55.53 | 42.97 | 18.87 |
| Minimap2 + GTF | -ax splice –junc-bed anno.bed | 78.19 | 61.29 | 47.65 | 23.4 |
| deSALT | -d 10 -x ont1d -s 2 -l 14 | 89.75 | 77.5 | 72.65 | 40.99 |
| deSALT + GTF | -d 10 -x ont1d -s 2 -l 14 -G anno.info | 90.16 | 81.1 | 77.19 | 49.07 |
| **Human ONT 1D, coverage = 4X, 46644 reads/56517199 bases/233621 exons** | | | | | |
| GMAP | -f samse --cross-species -z sense_force | 49.06 | 11.52 | 10.59 | 2.5 |
| Graphmap2 | -x rnaseq | 75.78 | 44.3 | 44.24 | 17.29 |
| Minimap2 | -ax splice | 70.68 | 47.38 | 33.84 | 14.5 |
| Minimap2 + GTF | -ax splice –junc-bed anno.bed | 71.88 | 52.75 | 37.54 | 17.92 |
| deSALT | -d 10 -x ont1d -s 2 -l 14 | 85.16 | 64.34 | 55.25 | 25.24 |
| deSALT + GTF | -d 10 -x ont1d -s 2 -l 14 -G anno.info | 86.64 | 72.08 | 65.13 | 35.27 |
| **Human ONT 1D, coverage = 10X, 103532 reads/141296228 bases/569988 exons** | | | | | |
| GMAP | -f samse --cross-species -z sense_force | 49.92 | 12.13 | 10.17 | 2.38 |
| Graphmap2 | -x rnaseq | 76.28 | 47.17 | 46.09 | 18.69 |
| Minimap2 | -ax splice | 72.45 | 50.34 | 37.46 | 16.32 |
| Minimap2 + GTF | -ax splice –junc-bed anno.bed | 73.68 | 55.88 | 41.63 | 20.27 |
| deSALT | -d 10 -x ont1d -s 2 -l 14 | 87.73 | 71.89 | 65.03 | 32.99 |
| deSALT + GTF | -d 10 -x ont1d -s 2 -l 14 -G anno.info | 88.59 | 77.52 | 72.78 | 43.43 |
| **Human ONT 1D, coverage = 30X, 309135 reads/423843578 bases/1706339 exons** | | | | | |
| GMAP | -f samse --cross-species -z sense_force | 50.12 | 12.05 | 10.19 | 2.36 |
| Graphmap2 | -x rnaseq | 76.49 | 48.26 | 47.13 | 19.63 |
| Minimap2 | -ax splice | 72.6 | 50.47 | 37.54 | 16.4 |
| Minimap2 + GTF | -ax splice –junc-bed anno.bed | 73.82 | 56.02 | 41.78 | 20.37 |
| deSALT | -d 10 -x ont1d -s 2 -l 14 | 88.36 | 74.52 | 69.05 | 38.61 |
| deSALT + GTF | -d 10 -x ont1d -s 2 -l 14 -G anno.info | 88.88 | 78.36 | 73.71 | 46.23 |
| **Human all protein coding genes ONT 1D, 1126328 reads/ 2414327781 bases/ 8884153 exons** | | | | | |

| | | | | | |
|---|---|---|---|---|---|
| GMAP | -f samse --cross-species -z sense_force | 54.03 | 11.59 | 7.01 | 1.37 |
| Graphmap2 | -x rnaseq | 82.36 | 46.01 | 36.75 | 14.22 |
| Minimap2 | -ax splice | 84.96 | 57.83 | 43.34 | 18.91 |
| Minimap2 + GTF | -ax splice –junc-bed anno.bed | 86.37 | 64.45 | 49.48 | 24.87 |
| deSALT | -d 10 -x ont1d -s 2 -l 14 | 92.16 | 67.58 | 56.1 | 25.04 |
| deSALT + GTF | -d 10 -x ont1d -s 2 -l 14 -G anno.info | 92.63 | 72.25 | 60.34 | 31.81 |
| **Simulated datasets by PBSim based on a real ONT dataset (PS-ONT)** | | | | | |
| **Fruit fly PS-ONT, coverage = 4X, 46305 reads/101883320 bases/293757 exons** | | | | | |
| GMAP | -f samse --cross-species -z sense_force | 93.48 | 89.39 | 92.1 | 64.47 |
| Graphmap2 | -x rnaseq | 93.28 | 95.68 | 98.27 | 84.94 |
| Minimap2 | -ax splice | 93.44 | 94.78 | 93.99 | 79.64 |
| Minimap2 + GTF | -ax splice –junc-bed anno.bed | 93.48 | 96.14 | 94.63 | 83.4 |
| deSALT | -d 10 | 93.62 | 96.82 | 96.98 | 86.89 |
| deSALT + GTF | -d 10 -G anno.info | 93.63 | 97.34 | 97.18 | 88.48 |
| **Fruit fly PS-ONT, coverage = 10X, 104775 reads/ 254610343 bases/ 557257 exons** | | | | | |
| GMAP | -f samse --cross-species -z sense_force | 93.56 | 89.86 | 92.86 | 64.14 |
| Graphmap2 | -x rnaseq | 93.27 | 95.91 | 98.42 | 85.19 |
| Minimap2 | -ax splice | 93.67 | 95.92 | 97.94 | 82.98 |
| Minimap2 + GTF | -ax splice –junc-bed anno.bed | 93.71 | 97.28 | 98.58 | 86.99 |
| deSALT | -d 10 | 93.71 | 97.73 | 98.67 | 89.63 |
| deSALT + GTF | -d 10 -G anno.info | 93.71 | 98.01 | 98.77 | 90.6 |
| **Fruit fly PS-ONT, coverage = 30X, 304510 reads/ 763907660 bases/ 1660900 exons** | | | | | |
| GMAP | -f samse --cross-species -z sense_force | 93.61 | 89.99 | 92.97 | 64.04 |
| Graphmap2 | -x rnaseq | 93.26 | 96.07 | 98.45 | 85.64 |
| Minimap2 | -ax splice | 93.71 | 96.06 | 98.03 | 83.18 |
| Minimap2 + GTF | -ax splice –junc-bed anno.bed | 93.75 | 97.41 | 98.69 | 87.29 |
| deSALT | -d 10 | 93.75 | 97.95 | 98.75 | 90.4 |
| deSALT + GTF | -d 10 -G anno.info | 93.75 | 98.16 | 98.84 | 91.13 |
| **Mouse PS-ONT, coverage = 4X, 47348 reads/ 65619390 bases/ 229025 exons** | | | | | |
| GMAP | -f samse --cross-species -z sense_force | 90.68 | 84.6 | 84.69 | 58.17 |

| | | | | | |
|---|---|---|---|---|---|
| Graphmap2 | -x rnaseq | 88.59 | 86.94 | 91.44 | 72.16 |
| Minimap2 | -ax splice | 91.58 | 90.62 | 86.75 | 68.98 |
| Minimap2 + GTF | -ax splice –junc-bed anno.bed | 91.64 | 91.53 | 87.41 | 70.61 |
| deSALT | -d 10 | 91.74 | 93.45 | 91.73 | 78.24 |
| deSALT + GTF | -d 10 -G anno.info | 91.83 | 94.03 | 92.04 | 79.74 |
| **Mouse PS-ONT, coverage = 10X, 105717 reads/ 163772057 bases/ 559198 exons** | | | | | |
| GMAP | -f samse --cross-species -z sense_force | 90.86 | 85.28 | 86.04 | 57.69 |
| Graphmap2 | -x rnaseq | 88.77 | 87.56 | 92.06 | 72.82 |
| Minimap2 | -ax splice | 92.19 | 92.84 | 94.18 | 74.62 |
| Minimap2 + GTF | -ax splice –junc-bed anno.bed | 92.24 | 93.76 | 94.88 | 76.44 |
| deSALT | -d 10 | 92.18 | 95.19 | 95.36 | 83.03 |
| deSALT + GTF | -d 10 -G anno.info | 92.2 | 95.51 | 95.55 | 83.96 |
| **Mouse PS-ONT, coverage = 30X, 305944 reads/ 491502093 bases/ 1665740 exons** | | | | | |
| GMAP | -f samse --cross-species -z sense_force | 90.97 | 85.59 | 86.15 | 57.44 |
| Graphmap2 | -x rnaseq | 88.75 | 87.78 | 92.08 | 73.15 |
| Minimap2 | -ax splice | 92.27 | 93.19 | 94.68 | 74.97 |
| Minimap2 + GTF | -ax splice –junc-bed anno.bed | 92.32 | 94.12 | 95.4 | 76.87 |
| deSALT | -d 10 | 92.21 | 95.54 | 95.56 | 83.87 |
| deSALT + GTF | -d 10 -G anno.info | 92.22 | 95.8 | 95.75 | 84.6 |
| **Human PS-ONT, coverage = 4X, 47186 reads/ 56649799 bases/ 220303 exons** | | | | | |
| GMAP | -f samse --cross-species -z sense_force | 90.83 | 84.2 | 84.22 | 57.39 |
| Graphmap2 | -x rnaseq | 88.76 | 86.74 | 90.91 | 71.15 |
| Minimap2 | -ax splice | 91.66 | 90.07 | 85.42 | 67.15 |
| Minimap2 + GTF | -ax splice –junc-bed anno.bed | 91.71 | 90.79 | 85.96 | 68.6 |
| deSALT | -d 10 | 92.37 | 93.46 | 92.11 | 77.78 |
| deSALT + GTF | -d 10 -G anno.info | 92.48 | 94.01 | 92.43 | 79.28 |
| **Human PS-ONT, coverage = 10X, 105043 reads/ 141294844 bases/ 535559 exons;** | | | | | |
| GMAP | -f samse --cross-species -z sense_force | 91.11 | 85.04 | 85.33 | 57.51 |
| Graphmap2 | -x rnaseq | 88.91 | 87.61 | 91.92 | 72.02 |
| Minimap2 | -ax splice | 92.36 | 92.72 | 94.06 | 73.83 |
| Minimap2 + GTF | -ax splice –junc-bed anno.bed | 92.42 | 93.47 | 94.63 | 75.39 |
| deSALT | -d 10 | 92.84 | 95.45 | 96.53 | 83.42 |

| | | | | | |
|---|---|---|---|---|---|
| deSALT + GTF | -d 10 -G anno.info | 92.89 | 95.75 | 96.75 | 84.38 |
| **Human PS-ONT, coverage = 30X, 304133 reads/ 424093591 bases/ 1594954 exons** | | | | | |
| GMAP | -f samse --cross-species -z sense_force | 91.24 | 85.4 | 85.64 | 57.61 |
| Graphmap2 | -x rnaseq | 88.9 | 87.78 | 91.87 | 72.39 |
| Minimap2 | -ax splice | 92.5 | 93.12 | 94.64 | 74.49 |
| Minimap2 + GTF | -ax splice –junc-bed anno.bed | 92.56 | 93.89 | 95.27 | 76.12 |
| deSALT | -d 10 | 92.95 | 95.79 | 96.87 | 84.23 |
| deSALT + GTF | -d 10 -G anno.info | 93.01 | 96.11 | 97.1 | 85.15 |
| **Human all protein coding genes PS-ONT, 1367081 reads/ 2624659295 bases/ 9310212 exons** | | | | | |
| GMAP | -f samse --cross-species -z sense_force | 93.46 | 77.2 | 72.01 | 40.15 |
| Graphmap2 | -x rnaseq | 90.31 | 82.38 | 86.2 | 57.31 |
| Minimap2 | -ax splice | 94.61 | 92.16 | 89.98 | 67.59 |
| Minimap2 + GTF | -ax splice –junc-bed anno.bed | 94.92 | 93.59 | 91.2 | 70.53 |
| deSALT | -d 10 -x ont1d -s 2 -l 14 | 95.48 | 95.50 | 95.08 | 78.68 |
| deSALT + GTF | -d 10 -x ont1d -s 2 -l 14 -G anno.info | 95.48 | 95.83 | 95.07 | 80.21 |
| **Simulated datasets by NanoSim based on a real ONT dataset (NS-ONT)** | | | | | |
| **Fruit fly NS-ONT, coverage = 4X, 34000 reads/198110604 bases/293757 exons** | | | | | |
| GMAP | -f samse --cross-species -z sense_force | 92.21 | 64.05 | 16.41 | 4.81 |
| Graphmap2 | -x rnaseq | 92.51 | 82.79 | 53.18 | 16.38 |
| Minimap2 | -ax splice | 92.94 | 93.55 | 95.78 | 63.06 |
| Minimap2 + GTF | -ax splice –junc-bed anno.bed | 92.99 | 97.15 | 97.68 | 79.91 |
| deSALT | -d 10 | 92.69 | 95.56 | 97.42 | 71.59 |
| deSALT + GTF | -d 10 -G anno.info | 92.7 | 97.39 | 97.81 | 83.01 |
| **Fruit fly NS-ONT, coverage = 10X, 86000 reads/ 503566481 bases/ 744738 exons** | | | | | |
| GMAP | -f samse --cross-species -z sense_force | 92.12 | 64.18 | 16.19 | 4.62 |
| Graphmap2 | -x rnaseq | 92.3 | 81.88 | 52.2 | 15.94 |
| Minimap2 | -ax splice | 92.84 | 93.54 | 95.81 | 63.17 |
| Minimap2 + GTF | -ax splice –junc-bed anno.bed | 92.89 | 97.14 | 97.68 | 80.02 |
| deSALT | -d 10 | 92.63 | 95.58 | 97.37 | 72.06 |
| deSALT + GTF | -d 10 -G anno.info | 92.65 | 97.39 | 97.68 | 83.3 |

| | | | | | |
|---|---|---|---|---|---|
| **Fruit fly NS-ONT, coverage = 30X, 250000 reads/ 1460343905 bases/ 2158993 exons** | | | | | |
| GMAP | -f samse --cross-species -z sense_force | 92.17 | 64.12 | 16.28 | 4.73 |
| Graphmap2 | -x rnaseq | 92.43 | 80.52 | 50.09 | 15.31 |
| Minimap2 | -ax splice | 92.88 | 93.49 | 95.75 | 63.06 |
| Minimap2 + GTF | -ax splice –junc-bed anno.bed | 92.92 | 97.13 | 97.68 | 79.58 |
| deSALT | -d 10 | 92.69 | 95.56 | 97.38 | 71.82 |
| deSALT + GTF | -d 10 -G anno.info | 92.7 | 97.39 | 97.72 | 83.12 |
| **Mouse NS-ONT, coverage = 4X, 37000 reads/ 216240438 bases/ 565964 exons** | | | | | |
| GMAP | -f samse --cross-species -z sense_force | 91.06 | 65.54 | 13.12 | 4.28 |
| Graphmap2 | -x rnaseq | 88.95 | 77.66 | 49.1 | 12.14 |
| Minimap2 | -ax splice | 93.35 | 92.71 | 96.92 | 51.85 |
| Minimap2 + GTF | -ax splice –junc-bed anno.bed | 93.44 | 96.33 | 98.26 | 67.01 |
| deSALT | -d 10 | 93.76 | 95.48 | 97.78 | 59.85 |
| deSALT + GTF | -d 10 -G anno.info | 93.79 | 97.33 | 98.15 | 72.63 |
| **Mouse NS-ONT, coverage = 10X, 90000 reads/ 526100251 bases/ 1374454 exons** | | | | | |
| GMAP | -f samse --cross-species -z sense_force | 91.01 | 65.47 | 12.75 | 4.13 |
| Graphmap2 | -x rnaseq | 88.85 | 77.12 | 47.84 | 12.06 |
| Minimap2 | -ax splice | 93.39 | 92.76 | 96.91 | 51.99 |
| Minimap2 + GTF | -ax splice –junc-bed anno.bed | 93.48 | 96.39 | 98.21 | 66.89 |
| deSALT | -d 10 | 93.81 | 95.52 | 97.89 | 59.98 |
| deSALT + GTF | -d 10 -G anno.info | 93.84 | 97.4 | 98.24 | 72.88 |
| **Mouse NS-ONT, coverage = 30X, 270000 reads/ 1575147062 bases/ 4108056 exons** | | | | | |
| GMAP | -f samse --cross-species -z sense_force | 90.99 | 65.35 | 12.78 | 4.25 |
| Graphmap2 | -x rnaseq | 88.87 | 74.73 | 44.79 | 11.64 |
| Minimap2 | -ax splice | 93.38 | 92.68 | 96.76 | 51.75 |
| Minimap2 + GTF | -ax splice –junc-bed anno.bed | 93.47 | 96.33 | 98.17 | 66.65 |
| deSALT | -d 10 | 93.86 | 95.54 | 98.02 | 59.86 |
| deSALT + GTF | -d 10 -G anno.info | 93.89 | 97.41 | 98.27 | 72.47 |
| **Human NS-ONT, coverage = 4X, 37000 reads/ 216047530 bases/ 496385 exons** | | | | | |
| GMAP | -f samse --cross-species -z sense_force | 91.06 | 65.19 | 13.32 | 4.46 |
| Graphmap2 | -x rnaseq | 88.32 | 74.01 | 45.55 | 12.46 |
| Minimap2 | -ax splice | 93.82 | 93.12 | 97.01 | 55.52 |

| | | | | | |
|---|---|---|---|---|---|
| Minimap2 + GTF | -ax splice –junc-bed anno.bed | 93.9 | 96.83 | 98.35 | 70.23 |
| deSALT | -d 10 | 93.96 | 95.6 | 98.25 | 63.23 |
| deSALT + GTF | -d 10 -G anno.info | 93.98 | 97.5 | 98.53 | 76.01 |
| **Human NS-ONT, coverage = 10X, 90000 reads/ 525306564 bases/ 1201021 exons** | | | | | |
| GMAP | -f samse --cross-species -z sense_force | 91.01 | 65.05 | 13.35 | 4.48 |
| Graphmap2 | -x rnaseq | 88.11 | 71.26 | 42.68 | 12.4 |
| Minimap2 | -ax splice | 93.8 | 93.17 | 97.03 | 55.84 |
| Minimap2 + GTF | -ax splice –junc-bed anno.bed | 93.88 | 96.87 | 98.32 | 70.49 |
| deSALT | -d 10 | 93.93 | 95.62 | 98.29 | 63.69 |
| deSALT + GTF | -d 10 -G anno.info | 93.95 | 97.52 | 98.56 | 76.11 |
| **Human NS-ONT, coverage = 30X, 270000 reads/ 1575546382 bases/ 227570 exons** | | | | | |
| GMAP | -f samse --cross-species -z sense_force | 91.12 | 65.19 | 13.31 | 4.47 |
| Graphmap2 | -x rnaseq | 87.99 | 69.6 | 40.12 | 11.73 |
| Minimap2 | -ax splice | 93.85 | 93.18 | 96.96 | 55.77 |
| Minimap2 + GTF | -ax splice –junc-bed anno.bed | 93.94 | 96.87 | 98.31 | 70.49 |
| deSALT | -d 10 | 94.01 | 95.58 | 98.33 | 63.36 |
| deSALT + GTF | -d 10 -G anno.info | 94.02 | 97.49 | 98.59 | 75.88 |
| **Human all protein coding genes NS-ONT, 987268 reads/ 5782765576 bases/ 14480231 exons** | | | | | |
| GMAP | -f samse --cross-species -z sense_force | 90.6 | 64.6 | 13.41 | 4.36 |
| Graphmap2 | -x rnaseq | 87.26 | 70.98 | 43.55 | 11.2 |
| Minimap2 | -ax splice | 93.42 | 91.19 | 96.55 | 53.18 |
| Minimap2 + GTF | -ax splice –junc-bed anno.bed | 93.53 | 95.25 | 98.18 | 68.87 |
| deSALT | -d 10 -s 2 -l 14 | 93.73 | 94.51 | 98.33 | 61.52 |
| deSALT + GTF | -d 10 -s 2 -l 14 -G anno.info | 93.77 | 96.54 | 98.62 | 75.01 |

a) The table depicts the results of the aligners on the 60 simulated datasets.

b) Refer to the Supplementary Notes for the command lines of the benchmarked aligners.

c) Base%: the proportion of bases being correctly aligned to their ground truth positions, i.e., the mapped positions of the bases are within 5 bp of their ground truth positions.

d) Exon%: the proportion of exons being correctly mapped. An exon in a certain read is considered to be correctly mapped only if its two boundaries are mapped within 5 bp of their ground truth positions in the reference genome.

e) Read80%: the proportion of Read80% reads. A read is considered to be a Read80% read only if it meets two conditions: $N_T/N_G > 80\%$ and $N_T/N_P > 80\%$, where $N_G$ is the number of ground truth exons within the read, $N_P$ is the number of exons predicted by the alignment, and $N_T$ is the number of true positive exons. Herein, a predicted exon is considered to be a true positive exon only if there is a ground truth exon in the read and the corresponding boundaries of the predicted exon and the ground truth exon are within 5 bp.

f) Read100%: the proportion of Read100% reads. A read is considered to be a Read100% read only if it meets two conditions: $N_T/N_G = 100\%$ and $N_T/N_P = 100\%$. It is worth noting that a Read100% read indicates that the read has a highly correct full-length alignment.

.

#### Supplementary Table 3. Benchmark results on the simulated reads with short exons [a]

| Aligner | Parameters [b] | Base% [c] | Exons% [d] | Read80% [e] | Read100% [f] |
|---|---|---|---|---|---|
| | | **Simulated PacBio ROI datasets** | | | |
| | | **Fruit fly PacBio ROI, coverage = 4X, 2657 reads/9799450 bases/29172 exons** | | | |
| GMAP | -f samse --cross-species -z sense_force | 99.17 | 95.43 | 96.39 | 70.91 |
| Graphmap2 | -x rnaseq | 98.31 | 92.02 | 92.54 | 53.69 |
| Minimap2 | -ax splice | 99.23 | 93.47 | 95.22 | 54.12 |
| Minimap2 + GTF | -ax splice –junc-bed anno.bed | 99.87 | 98.15 | 99.74 | 82.44 |
| deSALT | -d 10 -x ccs | 99.31 | 96.42 | 98.27 | 69.74 |
| deSALT + GTF | -d 10 -x ccs -G anno.info | 99.85 | 99.53 | 99.89 | 95.44 |
| | | **Fruit fly PacBio ROI, coverage = 10X, 6624 reads/25034865 bases/73558 exons** | | | |
| GMAP | -f samse --cross-species -z sense_force | 99.88 | 95.27 | 96.78 | 70.73 |
| Graphmap2 | -x rnaseq | 98.26 | 92.07 | 93.2 | 54.11 |
| Minimap2 | -ax splice | 99.91 | 93.47 | 95.79 | 54.63 |
| Minimap2 + GTF | -ax splice –junc-bed anno.bed | 99.87 | 98.24 | 99.74 | 82.92 |
| deSALT | -d 10 -x ccs | 99.97 | 96.73 | 98.96 | 72.95 |
| deSALT + GTF | -d 10 -x ccs -G anno.info | 99.84 | 99.51 | 99.94 | 95.31 |
| | | **Fruit fly PacBio ROI, coverage = 30X, 19879 reads/76182159 bases/222242 exons** | | | |
| GMAP | -f samse --cross-species -z sense_force | 99.61 | 94.62 | 96.34 | 69.9 |
| Graphmap2 | -x rnaseq | 98.34 | 92.4 | 93.48 | 54.94 |
| Minimap2 | -ax splice | 99.68 | 92.96 | 95.8 | 54.1 |
| Minimap2 + GTF | -ax splice –junc-bed anno.bed | 99.87 | 98.25 | 99.79 | 82.92 |
| deSALT | -d 10 -x ccs | 99.76 | 96.29 | 99.01 | 72.42 |
| deSALT + GTF | -d 10 -x ccs -G anno.info | 99.83 | 99.52 | 99.9 | 95.54 |
| | | **Mouse PacBio ROI, coverage = 4X, 1369 reads/3348645 bases/18578 exons** | | | |
| GMAP | -f samse --cross-species -z sense_force | 96.43 | 95.13 | 95.03 | 73.05 |
| Graphmap2 | -x rnaseq | 86.46 | 78.93 | 73.89 | 40.65 |
| Minimap2 | -ax splice | 94.09 | 88.44 | 85.24 | 48.58 |
| Minimap2 + GTF | -ax splice –junc-bed anno.bed | 98.15 | 97.4 | 97.4 | 80.34 |
| deSALT | -d 10 -x ccs | 96.04 | 94.1 | 92.7 | 70.27 |

| | | | | | |
|---|---|---|---|---|---|
| deSALT + GTF | -d 10 -x ccs -G anno.info | 98.24 | 97.94 | 97.4 | 85.01 |
| **Mouse PacBio ROI, coverage = 10X, 3332 reads/8371168 bases/45615 exons** | | | | | |
| GMAP | -f samse --cross-species -z sense_force | 99.06 | 94.34 | 91.42 | 64.2 |
| Graphmap2 | -x rnaseq | 86.18 | 79.54 | 75.16 | 42.18 |
| Minimap2 | -ax splice | 97.12 | 91.36 | 87.76 | 51.89 |
| Minimap2 + GTF | -ax splice –junc-bed anno.bed | 98.24 | 97.5 | 97.58 | 81.49 |
| deSALT | -d 10 -x ccs | 99.86 | 97.66 | 96.82 | 74.67 |
| deSALT + GTF | -d 10 -x ccs -G anno.info | 98.81 | 98.37 | 98.06 | 86.36 |
| **Mouse PacBio ROI, coverage = 30X, 10012 reads/25528211 bases/138591 exons** | | | | | |
| GMAP | -f samse --cross-species -z sense_force | 98.66 | 93.86 | 90.86 | 63.18 |
| Graphmap2 | -x rnaseq | 86.24 | 79.56 | 75.7 | 42.06 |
| Minimap2 | -ax splice | 96.53 | 90.93 | 87.4 | 51.32 |
| Minimap2 + GTF | -ax splice –junc-bed anno.bed | 98.15 | 97.53 | 97.9 | 81.16 |
| deSALT | -d 10 -x ccs | 99.32 | 97.09 | 96.47 | 72.87 |
| deSALT + GTF | -d 10 -x ccs -G anno.info | 98.69 | 98.38 | 98.22 | 86.28 |
| **Human PacBio ROI, coverage = 4X, 1244 reads/2645652 bases/16104 exons** | | | | | |
| GMAP | -f samse --cross-species -z sense_force | 97.35 | 93.01 | 93.25 | 70.34 |
| Graphmap2 | -x rnaseq | 92.52 | 87.09 | 88.09 | 55.48 |
| Minimap2 | -ax splice | 95.35 | 91.77 | 91.88 | 63.42 |
| Minimap2 + GTF | -ax splice –junc-bed anno.bed | 98.88 | 96.86 | 98.08 | 81.45 |
| deSALT | -d 10 -x ccs | 99.08 | 96.57 | 97.99 | 75.56 |
| deSALT + GTF | -d 10 -x ccs -G anno.info | 99.14 | 98.02 | 98.56 | 86.81 |
| **Human PacBio ROI, coverage = 10X, 3085 reads/6673839 bases/40650 exons** | | | | | |
| GMAP | -f samse --cross-species -z sense_force | 98.33 | 93.53 | 93.45 | 69.14 |
| Graphmap2 | -x rnaseq | 91.97 | 86.6 | 87.43 | 57.46 |
| Minimap2 | -ax splice | 96.22 | 92.49 | 91.64 | 63.4 |
| Minimap2 + GTF | -ax splice –junc-bed anno.bed | 99.04 | 97.05 | 98.22 | 81.9 |
| deSALT | -d 10 -x ccs | 99.7 | 97.24 | 97.67 | 77.18 |
| deSALT + GTF | -d 10 -x ccs -G anno.info | 99.21 | 98.06 | 98.63 | 92.56 |
| **Human PacBio ROI, coverage = 30X, 9243 reads/20206849 bases/122929 exons** | | | | | |
| GMAP | -f samse --cross-species -z sense_force | 98.68 | 93.87 | 93.48 | 69.14 |
| Graphmap2 | -x rnaseq | 92.58 | 87.38 | 87.7 | 57.09 |

| | | | | | |
|---|---|---|---|---|---|
| Minimap2 | -ax splice | 96.23 | 92.45 | 91.5 | 63.09 |
| Minimap2 + GTF | -ax splice –junc-bed anno.bed | 99.07 | 97.08 | 98.32 | 82.29 |
| deSALT | -d 10 -x ccs | 99.8 | 97.48 | 98.05 | 77.72 |
| deSALT + GTF | -d 10 -x ccs -G anno.info | 99.2 | 98.05 | 98.77 | 86.99 |
| **Human all protein coding genes PacBio ROI, 49618 reads/ 94213686 bases/ 620094 exons** | | | | | |
| GMAP | -f samse –cross-species -z sense_force | 98.87 | 94.11 | 94.71 | 66.74 |
| Graphmap2 | -x rnaseq | 89.72 | 82.68 | 81.52 | 47.3 |
| Minimap2 | -ax splice | 97.88 | 91.42 | 91.27 | 54.25 |
| Minimap2 + GTF | -ax splice –junc-bed anno.bed | 98.92 | 96.87 | 97.62 | 79.96 |
| deSALT | -d 10 -x ccs | 99.56 | 96.92 | 97.75 | 76.64 |
| deSALT + GTF | -d 10 -x ccs -G anno.info | 99.59 | 98.12 | 98.44 | 86.62 |
| **Simulated ONT 2D (1D$^2$) datasets** | | | | | |
| **Fruit fly ONT 2D (1D$^2$), coverage = 4X, 2669 reads/101867685 bases/229107 exons** | | | | | |
| GMAP | -f samse –cross-species -z sense_force | 94.69 | 70.17 | 49.42 | 15.51 |
| Graphmap2 | -x rnaseq | 94.07 | 86.29 | 82.52 | 31.61 |
| Minimap2 | -ax splice | 96.01 | 86.63 | 85.09 | 22.22 |
| Minimap2 + GTF | -ax splice –junc-bed anno.bed | 95.77 | 96.03 | 96.83 | 69.33 |
| deSALT | -d 10 -x ont2d -s 2 | 96.32 | 92.64 | 92.13 | 47.28 |
| deSALT + GTF | -d 10 -x ont2d -s 2 -G anno.info | 95.78 | 96.4 | 97.73 | 70.03 |
| **Fruit fly ONT 2D (1D$^2$), coverage = 10X, 6632 reads/25802683 bases/74427 exons** | | | | | |
| GMAP | -f samse –cross-species -z sense_force | 94.05 | 71.2 | 50.9 | 16.3 |
| Graphmap2 | -x rnaseq | 94.37 | 88.16 | 87.05 | 34.96 |
| Minimap2 | -ax splice | 95.55 | 86.56 | 85.57 | 23.72 |
| Minimap2 + GTF | -ax splice –junc-bed anno.bed | 95.89 | 96.3 | 97.33 | 69.83 |
| deSALT | -d 10 -x ont2d -s 2 | 95.88 | 94.18 | 95.75 | 56.77 |
| deSALT + GTF | -d 10 -x ont2d -s 2 -G anno.info | 95.94 | 96.95 | 98.6 | 72.65 |
| **Fruit fly ONT 2D (1D$^2$), coverage = 30X, 19914 reads/78707188 bases/225342 exons** | | | | | |
| GMAP | -f samse --cross-species -z sense_force | 94.6 | 71.09 | 50.21 | 16.26 |
| Graphmap2 | -x rnaseq | 94.36 | 88.62 | 88.4 | 35.78 |
| Minimap2 | -ax splice | 95.99 | 86.85 | 85.87 | 23.67 |

| | | | | | |
|---|---|---|---|---|---|
| Minimap2 + GTF | -ax splice –junc-bed anno.bed | 95.91 | 96.37 | 97.52 | 70.03 |
| deSALT | -d 10 -x ont2d -s 2 | 96.34 | 94.92 | 96.66 | 60.5 |
| deSALT + GTF | -d 10 -x ont2d -s 2 -G anno.info | 95.93 | 97.11 | 98.7 | 73.67 |
| **Mouse ONT 2D (1D²), coverage = 4X, 1326 reads/3335501 bases/18089 exons** | | | | | |
| GMAP | -f samse --cross-species -z sense_force | 88.38 | 64.7 | 40.57 | 12.9 |
| Graphmap2 | -x rnaseq | 81.69 | 71.68 | 59.8 | 21.35 |
| Minimap2 | -ax splice | 90.03 | 79.91 | 65.08 | 15.08 |
| Minimap2 + GTF | -ax splice –junc-bed anno.bed | 92.69 | 92.36 | 87.16 | 55.32 |
| deSALT | -d 10 -x ont2d -s 2 | 92.8 | 88.6 | 82.5 | 39.14 |
| deSALT + GTF | -d 10 -x ont2d -s 2 -G anno.info | 93.41 | 93.61 | 89.92 | 61.63 |
| **Mouse ONT 2D (1D²), coverage = 10X, 3298 reads/8408061 bases/45667 exons** | | | | | |
| GMAP | -f samse --cross-species -z sense_force | 90.22 | 66.14 | 40.39 | 12.49 |
| Graphmap2 | -x rnaseq | 81.96 | 73.03 | 62.49 | 22.69 |
| Minimap2 | -ax splice | 91.39 | 81.76 | 67.4 | 16.16 |
| Minimap2 + GTF | -ax splice –junc-bed anno.bed | 92.66 | 92.48 | 87.84 | 55.62 |
| deSALT | -d 10 -x ont2d -s 2 | 94.34 | 92.01 | 85.63 | 47.63 |
| deSALT + GTF | -d 10 -x ont2d -s 2 -G anno.info | 93.53 | 94.25 | 90.9 | 63.75 |
| **Mouse ONT 2D (1D²), coverage = 30X, 9934 reads/25658107 bases/139226 exons** | | | | | |
| GMAP | -f samse --cross-species -z sense_force | 89.4 | 66.1 | 41.68 | 12.86 |
| Graphmap2 | -x rnaseq | 81.94 | 73.55 | 63.53 | 23.59 |
| Minimap2 | -ax splice | 90.45 | 80.59 | 67.3 | 15.93 |
| Minimap2 + GTF | -ax splice –junc-bed anno.bed | 92.78 | 92.65 | 88.63 | 56.55 |
| deSALT | -d 10 -x ont2d -s 2 | 93.14 | 91.02 | 86.65 | 48.88 |
| deSALT + GTF | -d 10 -x ont2d -s 2 -G anno.info | 93.65 | 94.57 | 91.59 | 65.41 |
| **Human ONT 2D (1D²), coverage = 4X, 1246 reads/2631404 bases/16424 exons** | | | | | |
| GMAP | -f samse --cross-species -z sense_force | 89.1 | 67.68 | 45.83 | 17.26 |
| Graphmap2 | -x rnaseq | 85.42 | 77.02 | 68.1 | 25.89 |
| Minimap2 | -ax splice | 90.25 | 82.56 | 71.27 | 27.61 |
| Minimap2 + GTF | -ax splice –junc-bed anno.bed | 94.15 | 93.62 | 89.61 | 58.85 |
| deSALT | -d 10 -x ont2d -s 2 | 95.06 | 90.99 | 87.24 | 47.99 |
| deSALT + GTF | -d 10 -x ont2d -s 2 -G anno.info | 94.75 | 94.83 | 92.61 | 64.2 |
| **Human ONT 2D (1D²), coverage = 10X, 3099 reads/6631932 bases/41196 exons** | | | | | |

| | | | | | |
|---|---|---|---|---|---|
| GMAP | -f samse --cross-species -z sense_force | 87.85 | 66.83 | 46.21 | 16.65 |
| Graphmap2 | -x rnaseq | 85.94 | 79.22 | 71.42 | 30.19 |
| Minimap2 | -ax splice | 89.13 | 81.67 | 71.64 | 25.69 |
| Minimap2 + GTF | -ax splice –junc-bed anno.bed | 93.9 | 93.05 | 89.87 | 57.54 |
| deSALT | -d 10 -x ont2d -s 2 | 93.98 | 91.64 | 89.64 | 53.05 |
| deSALT + GTF | -d 10 -x ont2d -s 2 -G anno.info | 94.66 | 94.92 | 93.69 | 65.05 |
| **Human ONT 2D (1D$^2$), coverage = 30X, 9258 reads/20030911 bases/123352 exons** | | | | | |
| GMAP | -f samse --cross-species -z sense_force | 88.76 | 77.41 | 62.58 | 23.74 |
| Graphmap2 | -x rnaseq | 85.53 | 78.02 | 71.37 | 28.31 |
| Minimap2 | -ax splice | 93.5 | 85.8 | 77.23 | 26.07 |
| Minimap2 + GTF | -ax splice –junc-bed anno.bed | 94.09 | 93.57 | 91.23 | 58.92 |
| deSALT | -d 10 -x ont2d -s 2 | 94.98 | 92.35 | 90.13 | 52.2 |
| deSALT + GTF | -d 10 -x ont2d -s 2 -G anno.info | 94.83 | 94.94 | 93.99 | 64.64 |
| **Human all protein coding genes ONT 2D (1D$^2$), 48969 reads/ 145210695 bases/ 758229 exons** | | | | | |
| GMAP | -f samse --cross-species -z sense_force | 92.37 | 67.01 | 41.41 | 12.23 |
| Graphmap2 | -x rnaseq | 83.5 | 71.64 | 64.94 | 26.87 |
| Minimap2 | -ax splice | 95.53 | 84.23 | 75.17 | 20.17 |
| Minimap2 + GTF | -ax splice –junc-bed anno.bed | 96.82 | 93.93 | 91.4 | 59.47 |
| deSALT | -d 10 -x ont2d -s 2 | 97.76 | 93.44 | 91.14 | 53.08 |
| deSALT + GTF | -d 10 -x ont2d -s 2 -G anno.info | 97.86 | 96.01 | 95.18 | 69.06 |
| **Simulated PacBio subread datasets** | | | | | |
| **Fruit fly PacBio subread, coverage = 4X, 2425 reads/9649661 bases/26290 exons** | | | | | |
| GMAP | -f samse --cross-species -z sense_force | 82.91 | 61.57 | 31.01 | 6.85 |
| Graphmap2 | -x rnaseq | 87.17 | 85.57 | 81.64 | 29.88 |
| Minimap2 | -ax splice | 87.51 | 85.24 | 81.81 | 21.73 |
| Minimap2 + GTF | -ax splice –junc-bed anno.bed | 88.63 | 95.89 | 96.77 | 68.52 |
| deSALT | -d 10 -x clr -s 2 | 87.86 | 92.52 | 91.42 | 48.95 |
| deSALT + GTF | -d 10 -x clr -s 2 -G anno.info | 88.48 | 95.99 | 96.77 | 67.69 |
| **Fruit fly PacBio subread, coverage = 10X, 5981 reads/24299878 bases/66138 exons** | | | | | |
| GMAP | -f samse --cross-species -z sense_force | 84.08 | 61.94 | 32.28 | 6.83 |

| | | | | | |
|---|---|---|---|---|---|
| Graphmap2 | -x rnaseq | 87.12 | 86.71 | 84.83 | 32.15 |
| Minimap2 | -ax splice | 88.61 | 85.82 | 83.95 | 22.34 |
| Minimap2 + GTF | -ax splice –junc-bed anno.bed | 88.69 | 95.99 | 97.08 | 68.9 |
| deSALT | -d 10 -x clr -s 2 | 89.06 | 94.6 | 95.27 | 58.67 |
| deSALT + GTF | -d 10 -x clr -s 2 -G anno.info | 88.52 | 96.52 | 97.36 | 70.67 |
| **Fruit fly PacBio subread, coverage = 30X, 277541 reads/763798783 bases/1540635 exons** | | | | | |
| GMAP | -f samse --cross-species -z sense_force | 84.08 | 61.94 | 32.28 | 6.83 |
| Graphmap2 | -x rnaseq | 87.29 | 87.5 | 86.02 | 33.24 |
| Minimap2 | -ax splice | 88.61 | 85.82 | 83.95 | 22.34 |
| Minimap2 + GTF | -ax splice –junc-bed anno.bed | 88.67 | 96.13 | 97.4 | 69.13 |
| deSALT | -d 10 -x clr -s 2 | 88.91 | 95.08 | 96.43 | 60.82 |
| deSALT + GTF | -d 10 -x clr -s 2 -G anno.info | 88.53 | 96.64 | 97.95 | 70.75 |
| **Mouse PacBio subread, coverage = 4X, 1223 reads/3211130 bases/16386 exons** | | | | | |
| GMAP | -f samse --cross-species -z sense_force | 76.75 | 54.13 | 20.2 | 4.91 |
| Graphmap2 | -x rnaseq | 76.22 | 71.11 | 56.53 | 20.41 |
| Minimap2 | -ax splice | 83.45 | 79.3 | 61.82 | 13.65 |
| Minimap2 + GTF | -ax splice –junc-bed anno.bed | 85.6 | 91.88 | 86.28 | 51.07 |
| deSALT | -d 10 -x clr -s 2 | 86.4 | 88.48 | 80.95 | 36.47 |
| deSALT + GTF | -d 10 -x clr -s 2 -G anno.info | 86.37 | 93.09 | 89.67 | 56.12 |
| **Mouse PacBio subread, coverage = 10X, 2998 reads/8020991 bases/40870 exons** | | | | | |
| GMAP | -f samse --cross-species -z sense_force | 77.02 | 54.63 | 20.28 | 4.87 |
| Graphmap2 | -x rnaseq | 76.01 | 72.03 | 60.74 | 22.08 |
| Minimap2 | -ax splice | 83.13 | 79.02 | 63.04 | 12.94 |
| Minimap2 + GTF | -ax splice –junc-bed anno.bed | 85.49 | 91.57 | 86.48 | 50.32 |
| deSALT | -d 10 -x clr -s 2 | 86.51 | 90.35 | 84.96 | 45.76 |
| deSALT + GTF | -d 10 -x clr -s 2 -G anno.info | 86.64 | 94.06 | 91.14 | 61.07 |
| **Mouse PacBio subread, coverage = 30X, 9007 reads/24148406 bases/123267 exons** | | | | | |
| GMAP | -f samse --cross-species -z sense_force | 77.42 | 54.43 | 19.77 | 4.74 |
| Graphmap2 | -x rnaseq | 76.39 | 73.02 | 62.4 | 20.42 |
| Minimap2 | -ax splice | 83.46 | 79.34 | 64.32 | 13.08 |
| Minimap2 + GTF | -ax splice –junc-bed anno.bed | 85.48 | 91.54 | 87.09 | 50.54 |
| deSALT | -d 10 -x clr -s 2 | 86.18 | 90.36 | 85.11 | 45.81 |

| | | | | | |
|---|---|---|---|---|---|
| deSALT + GTF | -d 10 -x clr -s 2 -G anno.info | 86.53 | 93.92 | 91.09 | 62.65 |
| **Human PacBio subread, coverage = 4X, 1109 reads/2359725 bases/14010 exons** | | | | | |
| GMAP | -f samse --cross-species -z sense_force | 74.53 | 55.65 | 23.08 | 5.68 |
| Graphmap2 | -x rnaseq | 79.74 | 75.49 | 65.75 | 23.6 |
| Minimap2 | -ax splice | 82.54 | 81.52 | 68.98 | 22.45 |
| Minimap2 + GTF | -ax splice –junc-bed anno.bed | 86.07 | 90.57 | 87.84 | 48.09 |
| deSALT | -d 10 -x clr -s 2 | 86.43 | 90.11 | 85.75 | 44.72 |
| deSALT + GTF | -d 10 -x clr -s 2 -G anno.info | 86.68 | 92.05 | 89.71 | 53.33 |
| **Human PacBio subread, coverage = 10X, 2761 reads/5996015 bases/35333 exons** | | | | | |
| GMAP | -f samse --cross-species -z sense_force | 75.83 | 56.22 | 22.6 | 6.16 |
| Graphmap2 | -x rnaseq | 79.59 | 76.86 | 68.97 | 25.67 |
| Minimap2 | -ax splice | 84.06 | 82.85 | 70.01 | 23.65 |
| Minimap2 + GTF | -ax splice –junc-bed anno.bed | 86.05 | 90.77 | 88.2 | 48.91 |
| deSALT | -d 10 -x clr -s 2 | 88.39 | 92.43 | 88.7 | 48.39 |
| deSALT + GTF | -d 10 -x clr -s 2 -G anno.info | 87.02 | 92.95 | 91.74 | 56.92 |
| **Human PacBio subread, coverage = 30X, 8275 reads/17979517 bases/105647 exons** | | | | | |
| GMAP | -f samse --cross-species -z sense_force | 75.24 | 56.24 | 23.08 | 5.73 |
| Graphmap2 | -x rnaseq | 79.79 | 77.01 | 68.69 | 24.74 |
| Minimap2 | -ax splice | 83.64 | 82.73 | 70.98 | 22.65 |
| Minimap2 + GTF | -ax splice –junc-bed anno.bed | 85.99 | 90.76 | 88.23 | 48.5 |
| deSALT | -d 10 -x clr -s 2 | 87.99 | 93.23 | 90.21 | 51.9 |
| deSALT + GTF | -d 10 -x clr -s 2 -G anno.info | 87.06 | 93.22 | 91.99 | 58.91 |
| **Human all protein coding genes PacBio subread, 45377 reads/ 99585099 bases/ 572446 exons** | | | | | |
| GMAP | -f samse --cross-species -z sense_force | 77.14 | 55.16 | 22.96 | 5.26 |
| Graphmap2 | -x rnaseq | 77.92 | 72.09 | 62.9 | 21.44 |
| Minimap2 | -ax splice | 85.39 | 79.81 | 68.55 | 15.96 |
| Minimap2 + GTF | -ax splice –junc-bed anno.bed | 86.84 | 91.17 | 88.76 | 49.52 |
| deSALT | -d 10 -x clr -s 2 | 88.46 | 91.86 | 89.9 | 49.04 |
| deSALT + GTF | -d 10 -x clr -s 2 -G anno.info | 88.55 | 94.86 | 94.52 | 64.2 |
| **Simulated ONT 1D datasets** | | | | | |

| | | | | | |
|---|---|---|---|---|---|
| **Fruit fly ONT 1D, coverage = 4X, 2829 reads/10366837 bases/31649 exons** | | | | | |
| GMAP | -f samse --cross-species -z sense_force | 64.43 | 18.54 | 1.03 | 0.18 |
| Graphmap2 | -x rnaseq | 92.76 | 74.53 | 56.61 | 10.91 |
| Minimap2 | -ax splice | 92.51 | 72.39 | 55.11 | 2.58 |
| Minimap2 + GTF | -ax splice –junc-bed anno.bed | 93.46 | 86.38 | 79.67 | 38.47 |
| deSALT | -d 10 -x ont1d -s 2 -l 14 | 96.02 | 83.14 | 76.85 | 15.73 |
| deSALT + GTF | -d 10 -x ont1d -s 2 -l 14 -G anno.info | 96.02 | 90.71 | 88.17 | 44.88 |
| **Fruit fly ONT 1D, coverage = 10X, 7112 reads/26737114 bases/80525 exons** | | | | | |
| GMAP | -f samse --cross-species -z sense_force | 64.21 | 18.64 | 1.03 | 0.17 |
| Graphmap2 | -x rnaseq | 92.97 | 76.73 | 62.69 | 10.55 |
| Minimap2 | -ax splice | 92.93 | 73.51 | 56.72 | 2.64 |
| Minimap2 + GTF | -ax splice –junc-bed anno.bed | 93.86 | 87.56 | 82.47 | 39.74 |
| deSALT | -d 10 -x ont1d -s 2 -l 14 | 96.38 | 87.11 | 83.52 | 26.07 |
| deSALT + GTF | -d 10 -x ont1d -s 2 -l 14 -G anno.info | 96.37 | 92.42 | 91.18 | 49.92 |
| **Fruit fly ONT 1D, coverage = 30X, 21225 reads/80613021 bases/242158 exons** | | | | | |
| GMAP | -f samse --cross-species -z sense_force | 64.51 | 18.77 | 1.19 | 0.23 |
| Graphmap2 | -x rnaseq | 92.92 | 77.62 | 65.39 | 11.88 |
| Minimap2 | -ax splice | 92.94 | 73.59 | 57.14 | 2.95 |
| Minimap2 + GTF | -ax splice –junc-bed anno.bed | 93.84 | 87.47 | 82.5 | 39.34 |
| deSALT | -d 10 -x ont1d -s 2 -l 14 | 96.42 | 88.93 | 87.59 | 34.46 |
| deSALT + GTF | -d 10 -x ont1d -s 2 -l 14 -G anno.info | 96.34 | 92.8 | 91.9 | 52.88 |
| **Mouse ONT 1D, coverage = 4X, 1436 reads/3402252 bases/19736 exons** | | | | | |
| GMAP | -f samse --cross-species -z sense_force | 50.38 | 12.69 | 0.49 | 0.14 |
| Graphmap2 | -x rnaseq | 70.58 | 46.36 | 18.29 | 3.06 |
| Minimap2 | -ax splice | 77.72 | 55.96 | 26.39 | 1.11 |
| Minimap2 + GTF | -ax splice –junc-bed anno.bed | 80.41 | 69.64 | 48.33 | 17.87 |
| deSALT | -d 10 -x ont1d -s 2 -l 14 | 87.14 | 66.07 | 40.04 | 3.34 |
| deSALT + GTF | -d 10 -x ont1d -s 2 -l 14 -G anno.info | 86.39 | 76.34 | 57.65 | 23.16 |
| **Mouse ONT 1D, coverage = 10X, 3536 reads/8738830 bases/49776 exons** | | | | | |
| GMAP | -f samse --cross-species -z sense_force | 49.36 | 12.66 | 0.51 | 0.14 |
| Graphmap2 | -x rnaseq | 70.13 | 47.37 | 20.01 | 3.47 |
| Minimap2 | -ax splice | 77.51 | 56.37 | 25.51 | 0.62 |

| | | | | | |
|---|---|---|---|---|---|
| Minimap2 + GTF | -ax splice –junc-bed anno.bed | 80.06 | 69.75 | 48.34 | 18.26 |
| deSALT | -d 10 -x ont1d -s 2 -l 14 | 87.65 | 71.71 | 50.57 | 8.6 |
| deSALT + GTF | -d 10 -x ont1d -s 2 -l 14 -G anno.info | 87.32 | 79.29 | 61.88 | 28.3 |
| **Mouse ONT 1D, coverage = 30X, 10567 reads/26107819 bases/148826 exons** | | | | | |
| GMAP | -f samse --cross-species -z sense_force | 29.87 | 7.28 | 0.29 | 0.03 |
| Graphmap2 | -x rnaseq | 70.47 | 48.87 | 22.87 | 3.9 |
| Minimap2 | -ax splice | 77.74 | 56.62 | 25.69 | 0.68 |
| Minimap2 + GTF | -ax splice –junc-bed anno.bed | 80.3 | 70.07 | 48.49 | 17.94 |
| deSALT | -d 10 -x ont1d -s 2 -l 14 | 88.36 | 74.87 | 56.04 | 13.24 |
| deSALT + GTF | -d 10 -x ont1d -s 2 -l 14 -G anno.info | 88.46 | 81.19 | 65.29 | 31.33 |
| **Human ONT 1D, coverage = 4X, 1317 reads/2603348 bases/16963 exons** | | | | | |
| GMAP | -f samse --cross-species -z sense_force | 47.55 | 12.77 | 0.68 | 0.3 |
| Graphmap2 | -x rnaseq | 67.57 | 45.52 | 17.54 | 3.11 |
| Minimap2 | -ax splice | 73.25 | 51.66 | 22.1 | 1.52 |
| Minimap2 + GTF | -ax splice –junc-bed anno.bed | 78.48 | 69.35 | 47.15 | 17.54 |
| deSALT | -d 10 -x ont1d -s 2 -l 14 | 87.69 | 64.78 | 39.94 | 4.78 |
| deSALT + GTF | -d 10 -x ont1d -s 2 -l 14 -G anno.info | 86.28 | 75.29 | 55.35 | 21.64 |
| **Human ONT 1D, coverage = 10X, 3297 reads/6757929 bases/43499 exons** | | | | | |
| GMAP | -f samse --cross-species -z sense_force | 47.97 | 13.16 | 0.55 | 0.15 |
| Graphmap2 | -x rnaseq | 66.78 | 46.6 | 20.16 | 4.17 |
| Minimap2 | -ax splice | 72.61 | 51.49 | 24.17 | 1.4 |
| Minimap2 + GTF | -ax splice –junc-bed anno.bed | 78.32 | 69.47 | 48.02 | 18.32 |
| deSALT | -d 10 -x ont1d -s 2 -l 14 | 87.61 | 70.77 | 53.11 | 10.01 |
| deSALT + GTF | -d 10 -x ont1d -s 2 -l 14 -G anno.info | 87.73 | 80.26 | 64.84 | 27.68 |
| **Human ONT 1D, coverage = 30X, 9917 reads/20588822 bases/132385 exons** | | | | | |
| GMAP | -f samse --cross-species -z sense_force | 47.94 | 13.15 | 0.67 | 0.17 |
| Graphmap2 | -x rnaseq | 66.7 | 47.54 | 21.94 | 4.07 |
| Minimap2 | -ax splice | 72.96 | 52.07 | 23.59 | 1.27 |
| Minimap2 + GTF | -ax splice –junc-bed anno.bed | 79.19 | 70.28 | 47.97 | 18.09 |
| deSALT | -d 10 -x ont1d -s 2 -l 14 | 88.29 | 72.85 | 55.21 | 13.21 |
| deSALT + GTF | -d 10 -x ont1d -s 2 -l 14 -G anno.info | 87.9 | 80.86 | 64.96 | 28.81 |
| **Human all protein coding genes PacBio subread, 51863 reads/ 148485622 bases/ 815326 exons** | | | | | |

| | | | | | |
|---|---|---|---|---|---|
| GMAP | -f samse --cross-species -z sense_force | 47.74 | 10.13 | 0.38 | 0.07 |
| Graphmap2 | -x rnaseq | 77.92 | 72.09 | 62.9 | 21.44 |
| Minimap2 | -ax splice | 83.49 | 53.88 | 26.48 | 0.9 |
| Minimap2 + GTF | -ax splice –junc-bed anno.bed | 87.89 | 71.2 | 52.66 | 19.05 |
| deSALT | -d 10 -x ont1d -s 2 -l 14 | 91.19 | 64.68 | 40.71 | 3.81 |
| deSALT + GTF | -d 10 -x ont1d -s 2 -l 14 -G anno.info | 93.09 | 75.85 | 58.55 | 22.1 |
| **Simulated datasets by PBSim based on a real ONT dataset (PS-ONT)** | | | | | |
| **Fruit fly PS-ONT, coverage = 4X, 2597 reads/9279341 bases/27717 exons** | | | | | |
| GMAP | -f samse --cross-species -z sense_force | 93.48 | 86.9 | 84.87 | 31.57 |
| Graphmap2 | -x rnaseq | 92.56 | 90.53 | 91.3 | 40.47 |
| Minimap2 | -ax splice | 93.81 | 88.26 | 89.18 | 28.22 |
| Minimap2 + GTF | -ax splice –junc-bed anno.bed | 94.01 | 96.5 | 98.07 | 70.54 |
| deSALT | -d 10 | 93.8 | 94.17 | 95.23 | 55.49 |
| deSALT + GTF | -d 10 -G anno.info | 93.86 | 97.02 | 98.34 | 73.55 |
| **Fruit fly PS-ONT, coverage = 10X, 6513 reads/ 23899485 bases/ 70609 exons** | | | | | |
| GMAP | -f samse --cross-species -z sense_force | 93.38 | 86.86 | 84.17 | 29.68 |
| Graphmap2 | -x rnaseq | 92.55 | 90.92 | 91.91 | 41.1 |
| Minimap2 | -ax splice | 93.87 | 88.62 | 89.85 | 29.1 |
| Minimap2 + GTF | -ax splice –junc-bed anno.bed | 94.05 | 96.67 | 98.23 | 71.49 |
| deSALT | -d 10 | 93.82 | 95.36 | 97.16 | 62.2 |
| deSALT + GTF | -d 10 -G anno.info | 93.85 | 97.19 | 98.66 | 74.56 |
| **Fruit fly PS-ONT, coverage = 30X, 19548 reads/ 72652003 bases/ 212575 exons** | | | | | |
| GMAP | -f samse --cross-species -z sense_force | 93.44 | 86.98 | 84.78 | 31.03 |
| Graphmap2 | -x rnaseq | 92.52 | 91.07 | 92.54 | 42.09 |
| Minimap2 | -ax splice | 90.96 | 85.97 | 78.1 | 29.14 |
| Minimap2 + GTF | -ax splice –junc-bed anno.bed | 94.04 | 96.7 | 98.39 | 71.69 |
| deSALT | -d 10 | 93.78 | 95.72 | 97.51 | 64.97 |
| deSALT + GTF | -d 10 -G anno.info | 93.8 | 97.3 | 98.77 | 75.43 |
| **Mouse PS-ONT, coverage = 4X, 1305 reads/ 3149525 bases/ 17199 exons** | | | | | |
| GMAP | -f samse --cross-species -z sense_force | 90.27 | 83.21 | 93.41 | 24.83 |

| | | | | | |
|---|---|---|---|---|---|
| Graphmap2 | -x rnaseq | 81.91 | 78.1 | 71.49 | 30.57 |
| Minimap2 | -ax splice | 90.92 | 85.63 | 77.48 | 23.14 |
| Minimap2 + GTF | -ax splice –junc-bed anno.bed | 91.34 | 93.56 | 90.5 | 58.7 |
| deSALT | -d 10 | 91.59 | 91.1 | 88.28 | 45.06 |
| deSALT + GTF | -d 10 -G anno.info | 92.49 | 95.06 | 93.33 | 65.62 |
| **Mouse PS-ONT, coverage = 10X, 3236 reads/ 7959175 bases/ 43553 exons** | | | | | |
| GMAP | -f samse --cross-species -z sense_force | 89.9 | 82.96 | 74.04 | 25.4 |
| Graphmap2 | -x rnaseq | 81.32 | 78.02 | 73.3 | 30.47 |
| Minimap2 | -ax splice | 90.96 | 85.97 | 78.1 | 24.17 |
| Minimap2 + GTF | -ax splice –junc-bed anno.bed | 91.45 | 93.93 | 91.9 | 60.48 |
| deSALT | -d 10 | 92.41 | 92.78 | 89.22 | 52.75 |
| deSALT + GTF | -d 10 -G anno.info | 92.57 | 95.51 | 93.67 | 68.63 |
| **Mouse PS-ONT, coverage = 30X, 9728 reads/ 24184055 bases/ 131649 exons** | | | | | |
| GMAP | -f samse --cross-species -z sense_force | 90.37 | 83.54 | 74.35 | 24.67 |
| Graphmap2 | -x rnaseq | 81.49 | 78.44 | 74.03 | 31.13 |
| Minimap2 | -ax splice | 91.14 | 86.42 | 79.52 | 23.65 |
| Minimap2 + GTF | -ax splice –junc-bed anno.bed | 91.67 | 94.32 | 92.73 | 60.68 |
| deSALT | -d 10 | 92.5 | 93.71 | 90.79 | 56.78 |
| deSALT + GTF | -d 10 -G anno.info | 92.61 | 95.67 | 93.77 | 69.68 |
| **Human PS-ONT, coverage = 4X, 1209 reads/ 2394045 bases/ 15126 exons** | | | | | |
| GMAP | -f samse --cross-species -z sense_force | 89.36 | 84.79 | 78.33 | 28.7 |
| Graphmap2 | -x rnaseq | 86.28 | 83.75 | 79.32 | 37.63 |
| Minimap2 | -ax splice | 92.53 | 88.58 | 81.8 | 32.67 |
| Minimap2 + GTF | -ax splice –junc-bed anno.bed | 92.95 | 95.03 | 94.13 | 62.94 |
| deSALT | -d 10 | 92.98 | 93.41 | 92.89 | 54.26 |
| deSALT + GTF | -d 10 -G anno.info | 93.2 | 95.98 | 95.86 | 70.06 |
| **Human PS-ONT, coverage = 10X, 2997 reads/ 6139160 bases/ 38323 exons** | | | | | |
| GMAP | -f samse --cross-species -z sense_force | 89.61 | 84.57 | 78.18 | 28.13 |
| Graphmap2 | -x rnaseq | 85.11 | 83.33 | 81.35 | 37.64 |
| Minimap2 | -ax splice | 92.56 | 88.62 | 82.52 | 31.97 |
| Minimap2 + GTF | -ax splice –junc-bed anno.bed | 92.97 | 94.92 | 94.93 | 60.86 |
| deSALT | -d 10 | 93.03 | 94.11 | 92.56 | 57.66 |

| | | | | | |
|---|---|---|---|---|---|
| deSALT + GTF | -d 10 -G anno.info | 93.15 | 96.03 | 96.53 | 69.27 |
| **Human PS-ONT, coverage = 30X, 8968 reads/ 18471053 bases/ 115592 exons** | | | | | |
| GMAP | -f samse --cross-species -z sense_force | 89.64 | 84.63 | 77.88 | 29.09 |
| Graphmap2 | -x rnaseq | 85.3 | 83.66 | 80.87 | 38.04 |
| Minimap2 | -ax splice | 92.51 | 88.72 | 82.53 | 32.72 |
| Minimap2 + GTF | -ax splice –junc-bed anno.bed | 92.93 | 95.16 | 95.36 | 62.44 |
| deSALT | -d 10 | 93.11 | 94.18 | 92.76 | 57.89 |
| deSALT + GTF | -d 10 -G anno.info | 93.24 | 96.15 | 96.64 | 69.98 |
| **Human all protein coding genes PS-ONT, 51994 reads/ 139988211 bases/ 752983 exons** | | | | | |
| GMAP | -f samse --cross-species -z sense_force | 92.25 | 74.87 | 54.14 | 15.25 |
| Graphmap2 | -x rnaseq | 83.79 | 73.08 | 65.25 | 25.04 |
| Minimap2 | -ax splice | 93.66 | 86.28 | 79.81 | 25.68 |
| Minimap2 + GTF | -ax splice –junc-bed anno.bed | 94.68 | 94.76 | 93.32 | 62.44 |
| deSALT | -d 10 -x ont1d -s 2 -l 14 | 95.3 | 93.46 | 91.35 | 52.33 |
| deSALT + GTF | -d 10 -x ont1d -s 2 -l 14 -G anno.info | 95.37 | 96.21 | 95.47 | 70.69 |
| **Simulated datasets by NanoSim based on a real ONT dataset (NS-ONT)** | | | | | |
| **Fruit fly NS-ONT, coverage = 4X, 2384 reads/13743910 bases/31337 exons** | | | | | |
| GMAP | -f samse --cross-species -z sense_force | 91.79 | 63.49 | 13.38 | 1.13 |
| Graphmap2 | -x rnaseq | 92.66 | 82.29 | 63.38 | 7.09 |
| Minimap2 | -ax splice | 93.87 | 88.01 | 92.53 | 16.4 |
| Minimap2 + GTF | -ax splice –junc-bed anno.bed | 93.99 | 95.9 | 99.16 | 59.98 |
| deSALT | -d 10 | 94.01 | 93.95 | 98.24 | 47.69 |
| deSALT + GTF | -d 10 -G anno.info | 94.04 | 96.71 | 99.37 | 67.49 |
| **Fruit fly NS-ONT, coverage = 10X, 6096 reads/ 35333118 bases/ 80133 exons** | | | | | |
| GMAP | -f samse --cross-species -z sense_force | 91.75 | 63.28 | 12.66 | 1.18 |
| Graphmap2 | -x rnaseq | 92.57 | 81.98 | 62.98 | 6.02 |
| Minimap2 | -ax splice | 93.7 | 87.55 | 91.99 | 17.19 |
| Minimap2 + GTF | -ax splice –junc-bed anno.bed | 93.82 | 95.56 | 98.64 | 61.06 |
| deSALT | -d 10 | 93.84 | 93.7 | 98.06 | 49.72 |
| deSALT + GTF | -d 10 -G anno.info | 93.87 | 96.3 | 98.9 | 67.44 |

| | | | | | |
|---|---|---|---|---|---|
| **Fruit fly NS-ONT, coverage = 30X, 17509 reads/ 101086367 bases/ 229313 exons** | | | | | |
| GMAP | -f samse --cross-species -z sense_force | 91.84 | 63.6 | 13.1 | 1.03 |
| Graphmap2 | -x rnaseq | 92.17 | 78.93 | 56.59 | 6.21 |
| Minimap2 | -ax splice | 93.82 | 87.61 | 91.59 | 17.12 |
| Minimap2 + GTF | -ax splice –junc-bed anno.bed | 93.95 | 95.72 | 98.79 | 59.86 |
| deSALT | -d 10 | 93.97 | 94.18 | 98.36 | 50.95 |
| deSALT + GTF | -d 10 -G anno.info | 93.99 | 96.56 | 99.15 | 67.19 |
| **Mouse NS-ONT, coverage = 4X, 2945 reads/ 18301224 bases/ 95492 exons** | | | | | |
| GMAP | -f samse --cross-species -z sense_force | 89.21 | 65.59 | 7.2 | 0.24 |
| Graphmap2 | -x rnaseq | 81.81 | 73.95 | 55.31 | 1.6 |
| Minimap2 | -ax splice | 93.07 | 89.65 | 91.58 | 4.72 |
| Minimap2 + GTF | -ax splice –junc-bed anno.bed | 93.32 | 95.76 | 98.1 | 32.39 |
| deSALT | -d 10 | 93.39 | 94.32 | 96.43 | 24.31 |
| deSALT + GTF | -d 10 -G anno.info | 93.45 | 96.33 | 98.06 | 37.52 |
| **Mouse NS-ONT, coverage = 10X, 7175 reads/ 44790203 bases/ 234573 exons** | | | | | |
| GMAP | -f samse --cross-species -z sense_force | 89.72 | 65.88 | 7.71 | 0.22 |
| Graphmap2 | -x rnaseq | 81.65 | 73.76 | 54.2 | 1.57 |
| Minimap2 | -ax splice | 93.33 | 89.89 | 91.57 | 4.68 |
| Minimap2 + GTF | -ax splice –junc-bed anno.bed | 93.59 | 95.94 | 98.08 | 31.02 |
| deSALT | -d 10 | 93.62 | 94.28 | 96.43 | 23.25 |
| deSALT + GTF | -d 10 -G anno.info | 93.68 | 96.43 | 98.09 | 37.02 |
| **Mouse NS-ONT, coverage = 30X, 21815 reads/ 136722345 bases/ 711731 exons** | | | | | |
| GMAP | -f samse --cross-species -z sense_force | 89.26 | 65.51 | 6.94 | 0.14 |
| Graphmap2 | -x rnaseq | 80.96 | 69.83 | 46.35 | 1.64 |
| Minimap2 | -ax splice | 93.14 | 89.84 | 91.12 | 4.92 |
| Minimap2 + GTF | -ax splice –junc-bed anno.bed | 93.4 | 95.92 | 98.08 | 32.03 |
| deSALT | -d 10 | 93.95 | 94.62 | 98.3 | 24.36 |
| deSALT + GTF | -d 10 -G anno.info | 94.01 | 96.75 | 98.85 | 37.89 |
| **Human NS-ONT, coverage = 4X, 1978 reads/ 13216642 bases/ 78345 exons** | | | | | |
| GMAP | -f samse --cross-species -z sense_force | 88.45 | 66.95 | 9.2 | 0.35 |
| Graphmap2 | -x rnaseq | 85.85 | 79.5 | 64.56 | 3.08 |
| Minimap2 | -ax splice | 93.92 | 89.8 | 91.3 | 4.55 |

| | | | | | |
|---|---|---|---|---|---|
| Minimap2 + GTF | -ax splice –junc-bed anno.bed | 94.45 | 97.18 | 98.58 | 40.9 |
| deSALT | -d 10 | 94.52 | 96.02 | 98.23 | 28.87 |
| deSALT + GTF | -d 10 -G anno.info | 94.59 | 97.99 | 98.79 | 54.04 |
| **Human NS-ONT, coverage = 10X, 4705 reads/ 32337185 bases/ 191571 exons** | | | | | |
| GMAP | -f samse --cross-species -z sense_force | 88.46 | 66.49 | 7.35 | 0.13 |
| Graphmap2 | -x rnaseq | 85.43 | 75.15 | 52.79 | 2.1 |
| Minimap2 | -ax splice | 93.99 | 90.07 | 92.35 | 4.97 |
| Minimap2 + GTF | -ax splice –junc-bed anno.bed | 94.49 | 97.36 | 98.81 | 42.32 |
| deSALT | -d 10 | 94.56 | 96.16 | 98.6 | 31.05 |
| deSALT + GTF | -d 10 -G anno.info | 94.63 | 98.01 | 99.15 | 55.01 |
| **Human NS-ONT, coverage = 30X, 14180 reads/ 96649265 bases/ 569928 exons** | | | | | |
| GMAP | -f samse --cross-species -z sense_force | 88.33 | 66.59 | 7.63 | 0.16 |
| Graphmap2 | -x rnaseq | 84.42 | 73.26 | 47.85 | 1.42 |
| Minimap2 | -ax splice | 93.77 | 89.92 | 91.18 | 5.03 |
| Minimap2 + GTF | -ax splice –junc-bed anno.bed | 94.28 | 97.28 | 98.62 | 42.17 |
| deSALT | -d 10 | 94.35 | 95.92 | 98.41 | 28.36 |
| deSALT + GTF | -d 10 -G anno.info | 94.41 | 97.88 | 99.1 | 52.72 |
| **Human all protein coding genes NS-ONT, 72905 reads/ 497958564 bases/ 2501454 exons** | | | | | |
| GMAP | -f samse --cross-species -z sense_force | 88.48 | 65.11 | 7.44 | 0.25 |
| Graphmap2 | -x rnaseq | 78.34 | 67.23 | 44.73 | 1.47 |
| Minimap2 | -ax splice | 93.4 | 89.07 | 90.64 | 4.36 |
| Minimap2 + GTF | -ax splice –junc-bed anno.bed | 93.78 | 96.3 | 97.92 | 40.68 |
| deSALT | -d 10 -s 2 -l 14 | 93.89 | 94.83 | 98.08 | 26.48 |
| deSALT + GTF | -d 10 -s 2 -l 14 -G anno.info | 93.96 | 97.18 | 98.55 | 50.9 |

a) This table depicts the results of the aligners on all the reads with short exons (< 31 bp).

b) Refer to the Supplementary Notes for the command lines of the benchmarked aligners.

c) Base%: the proportion of bases being correctly aligned to their ground truth positions, i.e., the mapped positions of the bases are within 5 bp of their ground truth positions.

d) Exon%: the proportion of the exons being correctly mapped. An exon in a certain read is considered to be correctly mapped only if its two boundaries are mapped within 5 bp of their ground truth positions in the reference genome.

e) Read80%: the proportion of Read80% reads. A read is considered to be a Read80% read only if it meets two conditions: $N_T/N_G > 80\%$ and $N_T/N_P > 80\%$, where $N_G$ is the number of ground truth exons within the read, $N_P$ is the number of exons predicted by the alignment, and $N_T$ is the number of true positive exons. Herein, a predicted exon is considered to be a true positive exon only if there is a ground truth exon in the read and the corresponding boundaries of the predicted exon and the ground truth exon are within 5 bp.

f) Read100%: the proportion of Read100% reads. A read is considered to be a Read100% read only if it meets two conditions: $N_T/N_G = 100\%$ and $N_T/N_P = 100\%$. It is worth noting that a Read100% read indicates that the read has a highly correct full-length alignment.

#### Supplementary Table 4. Benchmark results on the simulated reads from the transcripts with various numbers of exons [a]

| Aligner | Parameters [b] | Base% [c] | Exons% [d] | Read80% [e] | Read100% [f] |
|---|---|---|---|---|---|
| | | **Simulated PacBio ROI datasets** | | | |
| | | **Fruit fly PacBio ROI, coverage = 4X, < 6 exons events, 29600 reads/40847769 bases/69512 exons** | | | |
| GMAP | -f samse --cross-species -z sense_force | 99.68 | 96.59 | 98.6 | 93.17 |
| Graphmap2 | -x rnaseq | 99.31 | 96.66 | 98.74 | 93.61 |
| Minimap2 | -ax splice | 99.37 | 97.2 | 98.3 | 94.32 |
| Minimap2 + GTF | -ax splice –junc-bed anno.bed | 99.12 | 97.35 | 98.55 | 94.62 |
| deSALT | -d 10 -x ccs | 99.62 | 98.23 | 99.5 | 96.2 |
| deSALT + GTF | -d 10 -x ccs -G anno.info | 99.24 | 96.73 | 99.17 | 93.08 |
| | | **Fruit fly PacBio ROI, coverage = 4X, 6-9 exons events, 8731 reads/28454497 bases/63936 exons** | | | |
| GMAP | -f samse --cross-species -z sense_force | 99.1 | 95.8 | 97.81 | 84.4 |
| Graphmap2 | -x rnaseq | 99.35 | 96.94 | 98.67 | 85.97 |
| Minimap2 | -ax splice | 99.13 | 97.76 | 99.23 | 92.11 |
| Minimap2 + GTF | -ax splice –junc-bed anno.bed | 99.84 | 99.44 | 100 | 96.07 |
| deSALT | -d 10 -x ccs | 99.15 | 98.29 | 99.2 | 94.61 |
| deSALT + GTF | -d 10 -x ccs -G anno.info | 99.83 | 99.62 | 99.94 | 97.54 |
| | | **Fruit fly PacBio ROI, coverage = 4X, > 9 exons events, 6965 reads/33008906 bases/96939 exons** | | | |
| GMAP | -f samse --cross-species -z sense_force | 99.02 | 96.83 | 97.75 | 75.95 |
| Graphmap2 | -x rnaseq | 98.58 | 96.12 | 96.84 | 74.22 |
| Minimap2 | -ax splice | 99.13 | 98.76 | 99.45 | 87.09 |
| Minimap2 + GTF | -ax splice –junc-bed anno.bed | 99.85 | 99.32 | 99.89 | 92.15 |
| deSALT | -d 10 -x ccs | 99.14 | 99.22 | 99.55 | 90.09 |
| deSALT + GTF | -d 10 -x ccs -G anno.info | 99.84 | 99.61 | 99.93 | 95.48 |
| | | **Fruit fly PacBio ROI, coverage = 10X, < 6 exons events, 62238 reads/100125992 bases/158496 exons** | | | |
| GMAP | -f samse --cross-species -z sense_force | 99.23 | 96.41 | 98.07 | 93.26 |
| Graphmap2 | -x rnaseq | 99.37 | 97.28 | 98.8 | 94.53 |
| Minimap2 | -ax splice | 98.97 | 97.55 | 98.66 | 95.29 |
| Minimap2 + GTF | -ax splice –junc-bed anno.bed | 99.16 | 98.15 | 99.13 | 95.89 |
| deSALT | -d 10 -x ccs | 99.2 | 98.24 | 99.31 | 96.81 |

| | | | | | |
|---|---|---|---|---|---|
| deSALT + GTF | -d 10 -x ccs -G anno.info | 99.29 | 97.5 | 99.28 | 96.85 |
| **Fruit fly PacBio ROI, coverage = 10X, 6-9 exons events, 21463 reads/70301415 bases/157116 exons** | | | | | |
| GMAP | -f samse --cross-species -z sense_force | 99.01 | 96.92 | 98.64 | 85.36 |
| Graphmap2 | -x rnaseq | 99.35 | 96.98 | 98.75 | 85.96 |
| Minimap2 | -ax splice | 99.13 | 98.83 | 100.14 | 93.01 |
| Minimap2 + GTF | -ax splice –junc-bed anno.bed | 99.85 | 99.42 | 99.99 | 95.97 |
| deSALT | -d 10 -x ccs | 99.15 | 99.42 | 99.17 | 95.7 |
| deSALT + GTF | -d 10 -x ccs -G anno.info | 99.82 | 99.65 | 99.94 | 97.69 |
| **Fruit fly PacBio ROI, coverage = 10X, > 9 exons events, 17589 reads/84422121 bases/245861 exon** | | | | | |
| GMAP | -f samse --cross-species -z sense_force | 99.71 | 96.57 | 97.6 | 75.48 |
| Graphmap2 | -x rnaseq | 98.46 | 96.02 | 96.77 | 73.95 |
| Minimap2 | -ax splice | 99.82 | 98.51 | 99.67 | 86.74 |
| Minimap2 + GTF | -ax splice –junc-bed anno.bed | 99.85 | 99.34 | 99.87 | 92.25 |
| deSALT | -d 10 -x ccs | 99.84 | 99.02 | 99.87 | 90.26 |
| deSALT + GTF | -d 10 -x ccs -G anno.info | 99.84 | 99.64 | 99.93 | 95.81 |
| **Fruit fly PacBio ROI, coverage = 30X, < 6 exons events, 171178 reads/296437653 bases/454719 exons** | | | | | |
| GMAP | -f samse --cross-species -z sense_force | 99.39 | 96.68 | 98.02 | 93.26 |
| Graphmap2 | -x rnaseq | 99.36 | 97.44 | 98.78 | 94.72 |
| Minimap2 | -ax splice | 99.19 | 98.09 | 99.08 | 95.89 |
| Minimap2 + GTF | -ax splice –junc-bed anno.bed | 99.27 | 98.52 | 99.54 | 96.49 |
| deSALT | -d 10 -x ccs | 99.36 | 98.63 | 99.37 | 97.18 |
| deSALT + GTF | -d 10 -x ccs -G anno.info | 99.32 | 97.77 | 99.29 | 97.21 |
| **Fruit fly PacBio ROI, coverage = 30X, 6-9 exons events, 64162 reads/211010822 bases/469724 exons** | | | | | |
| GMAP | -f samse --cross-species -z sense_force | 99.67 | 96.49 | 98.36 | 84.77 |
| Graphmap2 | -x rnaseq | 99.36 | 97.02 | 98.75 | 86.07 |
| Minimap2 | -ax splice | 99.72 | 98.48 | 99.89 | 92.58 |
| Minimap2 + GTF | -ax splice –junc-bed anno.bed | 99.87 | 99.42 | 99.99 | 95.94 |
| deSALT | -d 10 -x ccs | 99.74 | 99.13 | 99.92 | 95.6 |
| deSALT + GTF | -d 10 -x ccs -G anno.info | 99.84 | 99.66 | 99.96 | 97.74 |
| **Fruit fly PacBio ROI, coverage = 30X, > 9 exons events, 53015 reads/256400555 bases/742347 exons** | | | | | |
| GMAP | -f samse --cross-species -z sense_force | 99.92 | 96.67 | 97.55 | 75.45 |
| Graphmap2 | -x rnaseq | 98.48 | 96.02 | 96.71 | 74.01 |

| | | | | | |
|---|---|---|---|---|---|
| Minimap2 | -ax splice | 99.03 | 98.62 | 99.84 | 86.85 |
| Minimap2 + GTF | -ax splice –junc-bed anno.bed | 99.85 | 99.34 | 99.88 | 92.23 |
| deSALT | -d 10 -x ccs | 99.93 | 99.19 | 99.89 | 90.8 |
| deSALT + GTF | -d 10 -x ccs -G anno.info | 99.83 | 99.63 | 99.92 | 95.71 |
| **Mouse PacBio ROI, coverage = 4X, < 6 exons events, 34144 reads/33327445 bases/77876 exons** | | | | | |
| GMAP | -f samse --cross-species -z sense_force | 97.75 | 95.04 | 97.72 | 91.76 |
| Graphmap2 | -x rnaseq | 96.27 | 90.13 | 94.58 | 85.04 |
| Minimap2 | -ax splice | 97.97 | 93.37 | 95.87 | 90.31 |
| Minimap2 + GTF | -ax splice –junc-bed anno.bed | 98.21 | 94.61 | 96.64 | 91.23 |
| deSALT | -d 10 -x ccs | 97.69 | 94.85 | 97.21 | 91.93 |
| deSALT + GTF | -d 10 -x ccs -G anno.info | 97.53 | 93.73 | 96.75 | 89.12 |
| **Mouse PacBio ROI, coverage = 4X, 6-9 exons events, 6182 reads/10959544 bases/44384 exons** | | | | | |
| GMAP | -f samse --cross-species -z sense_force | 98.61 | 97.11 | 98.06 | 88.21 |
| Graphmap2 | -x rnaseq | 93.73 | 90.67 | 91.4 | 71.81 |
| Minimap2 | -ax splice | 98.02 | 94.98 | 96.26 | 87.3 |
| Minimap2 + GTF | -ax splice –junc-bed anno.bed | 99.13 | 98.07 | 98.9 | 91.82 |
| deSALT | -d 10 -x ccs | 99.27 | 97.74 | 98.79 | 91.81 |
| deSALT + GTF | -d 10 -x ccs -G anno.info | 99.01 | 98.01 | 98.67 | 92.76 |
| **Mouse PacBio ROI, coverage = 4X, > 9 exons events, 6486 reads/21864873 bases/110551 exons** | | | | | |
| GMAP | -f samse --cross-species -z sense_force | 99 | 98.67 | 99.14 | 84.21 |
| Graphmap2 | -x rnaseq | 90.73 | 87.29 | 85.84 | 54.35 |
| Minimap2 | -ax splice | 98.78 | 98 | 98.21 | 87.73 |
| Minimap2 + GTF | -ax splice –junc-bed anno.bed | 99.38 | 99.1 | 99.38 | 92.19 |
| deSALT | -d 10 -x ccs | 99.26 | 98.81 | 98.98 | 91.4 |
| deSALT + GTF | -d 10 -x ccs -G anno.info | 99.42 | 99.14 | 99.18 | 93.96 |
| **Mouse PacBio ROI, coverage = 10X, < 6 exons events, 71939 reads/81401924 bases/176379 exons** | | | | | |
| GMAP | -f samse --cross-species -z sense_force | 97.99 | 93.62 | 96.03 | 89.12 |
| Graphmap2 | -x rnaseq | 96.28 | 90.9 | 94.51 | 85.97 |
| Minimap2 | -ax splice | 98.36 | 94.94 | 96.89 | 92.32 |
| Minimap2 + GTF | -ax splice –junc-bed anno.bed | 98.39 | 95.96 | 97.62 | 93.23 |
| deSALT | -d 10 -x ccs | 98.05 | 95.91 | 97.45 | 93.25 |
| deSALT + GTF | -d 10 -x ccs -G anno.info | 97.61 | 94.74 | 97.45 | 93.48 |

| | | | | | |
|---|---|---|---|---|---|
| **Mouse PacBio ROI, coverage = 10X, 6-9 exons events, 15326 reads/27583428 bases/110033 exons** | | | | | |
| GMAP | -f samse --cross-species -z sense_force | 97.99 | 94.11 | 95.45 | 78.21 |
| Graphmap2 | -x rnaseq | 93.57 | 90.69 | 90.71 | 71.4 |
| Minimap2 | -ax splice | 97.51 | 94.97 | 96.22 | 87.25 |
| Minimap2 + GTF | -ax splice –junc-bed anno.bed | 99.15 | 98.12 | 98.94 | 91.85 |
| deSALT | -d 10 -x ccs | 98.7 | 97.72 | 98.62 | 91.8 |
| deSALT + GTF | -d 10 -x ccs -G anno.info | 98.97 | 98.01 | 98.69 | 92.75 |
| **Mouse PacBio ROI, coverage = 10X, > 9 exons events, 16233 reads/55208415 bases/277753 exons** | | | | | |
| GMAP | -f samse --cross-species -z sense_force | 98.91 | 95.57 | 96.43 | 68.52 |
| Graphmap2 | -x rnaseq | 90.76 | 87.36 | 86.02 | 54.83 |
| Minimap2 | -ax splice | 98.82 | 98.03 | 98.16 | 87.91 |
| Minimap2 + GTF | -ax splice –junc-bed anno.bed | 99.45 | 99.16 | 99.42 | 92.45 |
| deSALT | -d 10 -x ccs | 99.29 | 98.79 | 99.07 | 91.33 |
| deSALT + GTF | -d 10 -x ccs -G anno.info | 99.54 | 99.21 | 99.27 | 94.12 |
| **Mouse PacBio ROI, coverage = 30X, < 6 exons events, 198377 reads/242311390 bases/506426 exons** | | | | | |
| GMAP | -f samse --cross-species -z sense_force | 97.74 | 93.77 | 95.83 | 89.41 |
| Graphmap2 | -x rnaseq | 96.29 | 91.35 | 94.5 | 86.52 |
| Minimap2 | -ax splice | 98.14 | 95.5 | 97.3 | 93.34 |
| Minimap2 + GTF | -ax splice –junc-bed anno.bed | 98.43 | 96.65 | 98.12 | 94.36 |
| deSALT | -d 10 -x ccs | 97.76 | 96.17 | 97.45 | 93.87 |
| deSALT + GTF | -d 10 -x ccs -G anno.info | 97.78 | 96.3 | 97.47 | 94.18 |
| **Mouse PacBio ROI, coverage = 30X, 6-9 exons events, 45827 reads/82794202 bases/328948 exons** | | | | | |
| GMAP | -f samse --cross-species -z sense_force | 98.61 | 94.35 | 95.85 | 78.75 |
| Graphmap2 | -x rnaseq | 93.75 | 90.77 | 91.53 | 71.67 |
| Minimap2 | -ax splice | 98.03 | 95.12 | 96.3 | 87.48 |
| Minimap2 + GTF | -ax splice –junc-bed anno.bed | 99.12 | 98.11 | 98.92 | 91.95 |
| deSALT | -d 10 -x ccs | 99.22 | 97.87 | 98.76 | 91.97 |
| deSALT + GTF | -d 10 -x ccs -G anno.info | 98.97 | 98.03 | 98.7 | 92.89 |
| **Mouse PacBio ROI, coverage = 30X, > 9 exons events, 48580 reads/166313402 bases/834224 exons** | | | | | |
| GMAP | -f samse --cross-species -z sense_force | 99.01 | 95.6 | 96.4 | 68.59 |
| Graphmap2 | -x rnaseq | 90.73 | 87.31 | 85.86 | 54.96 |
| Minimap2 | -ax splice | 98.9 | 98.09 | 98.19 | 87.93 |

| | | | | | |
|---|---|---|---|---|---|
| Minimap2 + GTF | -ax splice –junc-bed anno.bed | 99.4 | 99.12 | 99.36 | 92.32 |
| deSALT | -d 10 -x ccs | 99.4 | 98.91 | 99.22 | 91.73 |
| deSALT + GTF | -d 10 -x ccs -G anno.info | 99.51 | 99.21 | 99.27 | 94.24 |
| **Human PacBio ROI, coverage = 4X, < 6 exons events, 34881 reads/28674816 bases/83324 exons** | | | | | |
| GMAP | -f samse --cross-species -z sense_force | 98.83 | 94.54 | 96.88 | 90.11 |
| Graphmap2 | -x rnaseq | 97.61 | 91.6 | 95.31 | 86.56 |
| Minimap2 | -ax splice | 98.54 | 94.65 | 96.27 | 91.29 |
| Minimap2 + GTF | -ax splice –junc-bed anno.bed | 98.97 | 95.68 | 97.23 | 92.66 |
| deSALT | -d 10 -x ccs | 98.9 | 96.43 | 98.09 | 93.74 |
| deSALT + GTF | -d 10 -x ccs -G anno.info | 98.93 | 96.6 | 98.16 | 94.05 |
| **Human PacBio ROI, coverage = 4X, 6-9 exons events, 6016 reads/9871523 bases/42525 exons** | | | | | |
| GMAP | -f samse --cross-species -z sense_force | 98.99 | 95.76 | 97.36 | 82.35 |
| Graphmap2 | -x rnaseq | 93.83 | 91.53 | 92.34 | 73.95 |
| Minimap2 | -ax splice | 98.69 | 97.01 | 97.94 | 90.38 |
| Minimap2 + GTF | -ax splice –junc-bed anno.bed | 99.08 | 98.11 | 98.73 | 92.16 |
| deSALT | -d 10 -x ccs | 99.11 | 98.31 | 98.94 | 93.67 |
| deSALT + GTF | -d 10 -x ccs -G anno.info | 99.21 | 98.48 | 98.99 | 93.51 |
| **Human PacBio ROI, coverage = 4X, > 9 exons events, 5886 reads/18656908 bases/96807 exons** | | | | | |
| GMAP | -f samse --cross-species -z sense_force | 98.95 | 95.38 | 97.11 | 69.42 |
| Graphmap2 | -x rnaseq | 93.32 | 90.30 | 90.63 | 55.56 |
| Minimap2 | -ax splice | 98.8 | 97.82 | 98.76 | 88.38 |
| Minimap2 + GTF | -ax splice –junc-bed anno.bed | 99.44 | 98.45 | 99.52 | 86.13 |
| deSALT | -d 10 -x ccs | 99.69 | 98.78 | 99.56 | 91.44 |
| deSALT + GTF | -d 10 -x ccs -G anno.info | 99.47 | 98.76 | 99.52 | 88.95 |
| **Human PacBio ROI, coverage = 10X, < 6 exons events, 73602 reads/69736938 bases/189283 exons** | | | | | |
| GMAP | -f samse --cross-species -z sense_force | 99 | 95.03 | 96.75 | 90.49 |
| Graphmap2 | -x rnaseq | 97.6 | 92.41 | 95.36 | 87.62 |
| Minimap2 | -ax splice | 98.88 | 96.07 | 97.3 | 93.25 |
| Minimap2 + GTF | -ax splice –junc-bed anno.bed | 99.1 | 96.91 | 98.18 | 94.54 |
| deSALT | -d 10 -x ccs | 99.08 | 97.18 | 98.31 | 94.92 |
| deSALT + GTF | -d 10 -x ccs -G anno.info | 99.09 | 97.32 | 98.31 | 95.26 |
| **Human PacBio ROI, coverage = 10X, 6-9 exons events, 15055 reads/25082950 bases/106471 exons** | | | | | |

| | | | | | |
|---|---|---|---|---|---|
| GMAP | -f samse --cross-species -z sense_force | 98.84 | 95.7 | 97.27 | 82.42 |
| Graphmap2 | -x rnaseq | 93.78 | 91.49 | 92.22 | 73.83 |
| Minimap2 | -ax splice | 98.47 | 96.9 | 97.72 | 90.63 |
| Minimap2 + GTF | -ax splice –junc-bed anno.bed | 99.09 | 98.07 | 98.72 | 91.78 |
| deSALT | -d 10 -x ccs | 98.95 | 98.27 | 98.93 | 93.82 |
| deSALT + GTF | -d 10 -x ccs -G anno.info | 99.14 | 98.39 | 98.99 | 93.15 |
| **Human PacBio ROI, coverage = 10X, > 9 exons events, 14682 reads/46949479 bases/242162 exons** | | | | | |
| GMAP | -f samse --cross-species -z sense_force | 98.95 | 95.53 | 96.68 | 68.73 |
| Graphmap2 | -x rnaseq | 93.11 | 89.91 | 90.55 | 54.95 |
| Minimap2 | -ax splice | 98.76 | 98.14 | 98.89 | 88.55 |
| Minimap2 + GTF | -ax splice –junc-bed anno.bed | 99.45 | 98.51 | 99.55 | 86.37 |
| deSALT | -d 10 -x ccs | 99.58 | 99.03 | 99.55 | 91.94 |
| deSALT + GTF | -d 10 -x ccs -G anno.info | 99.48 | 98.79 | 99.56 | 92.25 |
| **Human PacBio ROI, coverage = 30X, < 6 exons events, 203063 reads/207294297 bases/543941 exons** | | | | | |
| GMAP | -f samse --cross-species -z sense_force | 98.99 | 95.37 | 96.74 | 91.01 |
| Graphmap2 | -x rnaseq | 97.67 | 92.78 | 95.4 | 88.11 |
| Minimap2 | -ax splice | 98.93 | 96.78 | 97.92 | 94.45 |
| Minimap2 + GTF | -ax splice –junc-bed anno.bed | 99.17 | 97.48 | 98.66 | 95.47 |
| deSALT | -d 10 -x ccs | 99.09 | 97.62 | 98.47 | 95.75 |
| deSALT + GTF | -d 10 -x ccs -G anno.info | 99.11 | 97.78 | 98.47 | 96.14 |
| **Human PacBio ROI, coverage = 30X, 6-9 exons events, 44964 reads/74851113 bases/317959 exons** | | | | | |
| GMAP | -f samse --cross-species -z sense_force | 99.02 | 95.78 | 97.26 | 82.58 |
| Graphmap2 | -x rnaseq | 93.7 | 91.41 | 92.07 | 73.58 |
| Minimap2 | -ax splice | 98.62 | 96.84 | 97.67 | 90.44 |
| Minimap2 + GTF | -ax splice –junc-bed anno.bed | 99.16 | 98.15 | 98.8 | 91.84 |
| deSALT | -d 10 -x ccs | 99.15 | 98.36 | 99.01 | 93.96 |
| deSALT + GTF | -d 10 -x ccs -G anno.info | 99.2 | 98.39 | 98.98 | 94.14 |
| **Human PacBio ROI, coverage = 30X, > 9 exons events, 44090 reads/141865473 bases/729517 exons** | | | | | |
| GMAP | -f samse --cross-species -z sense_force | 98.88 | 95.38 | 96.56 | 68.85 |
| Graphmap2 | -x rnaseq | 93.27 | 89.95 | 90.53 | 55.16 |
| Minimap2 | -ax splice | 98.69 | 98.01 | 98.71 | 88.6 |
| Minimap2 + GTF | -ax splice –junc-bed anno.bed | 99.44 | 98.48 | 99.47 | 86.55 |

| | | | | | |
|---|---|---|---|---|---|
| deSALT | -d 10 -x ccs | 99.54 | 98.97 | 99.48 | 91.99 |
| deSALT + GTF | -d 10 -x ccs -G anno.info | 99.58 | 99.03 | 99.49 | 92.55 |
| **Human all protein coding genes PacBio ROI, < 6 exons events, 787883 reads/ 1208784180 bases/ 2068575 exons** | | | | | |
| GMAP | -f samse --cross-species -z sense_force | 99.62 | 95.7 | 97.43 | 91.46 |
| Graphmap2 | -x rnaseq | 98.51 | 93.07 | 95.92 | 88.18 |
| Minimap2 | -ax splice | 99.55 | 96.99 | 98.55 | 94.58 |
| Minimap2 + GTF | -ax splice –junc-bed anno.bed | 99.68 | 97.6 | 99.06 | 95.17 |
| deSALT | -d 10 -x ccs | 99.71 | 97.11 | 99.1 | 93.71 |
| deSALT + GTF | -d 10 -x ccs -G anno.info | 99.73 | 97.22 | 99.18 | 93.85 |
| **Human all protein coding genes PacBio ROI, 6-9 exons events, 268961 reads/ 463333791 bases/ 1945073 exons** | | | | | |
| GMAP | -f samse --cross-species -z sense_force | 99.49 | 95.81 | 97.36 | 81.7 |
| Graphmap2 | -x rnaseq | 95.1 | 91.52 | 92.62 | 72.76 |
| Minimap2 | -ax splice | 99.07 | 96.99 | 98.07 | 89.66 |
| Minimap2 + GTF | -ax splice –junc-bed anno.bed | 99.62 | 98.11 | 99.01 | 91.83 |
| deSALT | -d 10 -x ccs | 99.84 | 98.66 | 99.39 | 93.4 |
| deSALT + GTF | -d 10 -x ccs -G anno.info | 99.84 | 98.75 | 99.43 | 93.9 |
| **Human all protein coding genes PacBio ROI, >9 exons events, 319249 reads/ 747880599 bases/ 4737124 exons** | | | | | |
| GMAP | -f samse --cross-species -z sense_force | 99.12 | 95.44 | 96.55 | 69.2 |
| Graphmap2 | -x rnaseq | 94.21 | 88.3 | 88.21 | 51.94 |
| Minimap2 | -ax splice | 99.01 | 97.24 | 98.33 | 81.85 |
| Minimap2 + GTF | -ax splice –junc-bed anno.bed | 99.43 | 98.1 | 98.98 | 85.02 |
| deSALT | -d 10 -x ccs | 99.77 | 98.62 | 99.32 | 87.41 |
| deSALT + GTF | -d 10 -x ccs -G anno.info | 99.75 | 98.72 | 99.32 | 88.66 |
| **Simulated ONT 2D ($1D^2$) datasets** | | | | | |
| **Fruit fly ONT 2D ($1D^2$), coverage = 4X, < 6 exons events, 29228 reads/39799847 bases/70477 exons** | | | | | |
| GMAP | -f samse --cross-species -z sense_force | 93.9 | 72.86 | 81.28 | 55.17 |
| Graphmap2 | -x rnaseq | 95.16 | 88.68 | 95.35 | 77.52 |
| Minimap2 | -ax splice | 94.19 | 85.52 | 87.93 | 72.23 |
| Minimap2 + GTF | -ax splice –junc-bed anno.bed | 94.13 | 87.46 | 89.32 | 75.11 |
| deSALT | -d 10 -x ont2d -s 2 | 95.23 | 90.05 | 94.99 | 79.8 |

| | | | | | |
|---|---|---|---|---|---|
| deSALT + GTF | -d 10 -x ont2d -s 2 -G anno.info | 95.31 | 92.06 | 96.23 | 83.5 |
| **Fruit fly ONT 2D (1D$^2$), coverage = 4X, 6-9 exons events, 8325 reads/26798031 bases/60669 exons** | | | | | |
| GMAP | -f samse --cross-species -z sense_force | 94.91 | 75.98 | 67.1 | 29.45 |
| Graphmap2 | -x rnaseq | 95.41 | 93.42 | 96.09 | 70.88 |
| Minimap2 | -ax splice | 96.01 | 95.39 | 98.28 | 76.88 |
| Minimap2 + GTF | -ax splice –junc-bed anno.bed | 95.86 | 97.41 | 99.47 | 84.63 |
| deSALT | -d 10 -x ont2d -s 2 | 96.18 | 97.3 | 99.33 | 84.11 |
| deSALT + GTF | -d 10 -x ont2d -s 2 -G anno.info | 95.9 | 97.98 | 99.59 | 87.57 |
| **Fruit fly ONT 2D (1D$^2$), coverage = 4X, > 9 exons events, 6892 reads/35269807 bases/97961 exons** | | | | | |
| GMAP | -f samse --cross-species -z sense_force | 94.16 | 73.98 | 51.94 | 12.77 |
| Graphmap2 | -x rnaseq | 94.53 | 93.1 | 94.01 | 56.43 |
| Minimap2 | -ax splice | 95.54 | 96.06 | 98.24 | 67.9 |
| Minimap2 + GTF | -ax splice –junc-bed anno.bed | 95.94 | 98.24 | 99.59 | 80.69 |
| deSALT | -d 10 -x ont2d -s 2 | 95.66 | 97.79 | 99.19 | 78.63 |
| deSALT + GTF | -d 10 -x ont2d -s 2 -G anno.info | 95.97 | 98.58 | 99.74 | 84.25 |
| **Fruit fly ONT 2D (1D$^2$), coverage = 10X, < 6 exons events, 60612 reads/96611912 bases/158794 exons** | | | | | |
| GMAP | -f samse --cross-species -z sense_force | 93.65 | 74.02 | 81.22 | 54.95 |
| Graphmap2 | -x rnaseq | 95.23 | 91.11 | 96.92 | 80.28 |
| Minimap2 | -ax splice | 94.45 | 89.27 | 94.17 | 77.36 |
| Minimap2 + GTF | -ax splice –junc-bed anno.bed | 94.88 | 91.37 | 95.89 | 80.69 |
| deSALT | -d 10 -x ont2d -s 2 | 94.94 | 93.05 | 97.09 | 84.4 |
| deSALT + GTF | -d 10 -x ont2d -s 2 -G anno.info | 94.99 | 94.27 | 97.71 | 87 |
| **Fruit fly ONT 2D (1D$^2$), coverage = 10X, 6-9 exons events, 20799 reads/67295749 bases/151677 exons** | | | | | |
| GMAP | -f samse --cross-species -z sense_force | 95.1 | 75.84 | 66.7 | 29.9 |
| Graphmap2 | -x rnaseq | 95.43 | 94.57 | 97.19 | 74.74 |
| Minimap2 | -ax splice | 96.05 | 95.33 | 98.34 | 76.65 |
| Minimap2 + GTF | -ax splice –junc-bed anno.bed | 95.85 | 97.31 | 99.44 | 84.26 |
| deSALT | -d 10 -x ont2d -s 2 | 96.24 | 97.95 | 99.76 | 87.16 |
| deSALT + GTF | -d 10 -x ont2d -s 2 -G anno.info | 95.89 | 98.18 | 99.63 | 89.01 |
| **Fruit fly ONT 2D (1D$^2$), coverage = 10X, > 9 exons events, 17569 reads/90693397 bases/250238 exons** | | | | | |
| GMAP | -f samse --cross-species -z sense_force | 94.87 | 74.79 | 53.67 | 13.32 |
| Graphmap2 | -x rnaseq | 94.5 | 93.83 | 95.18 | 60.04 |

| | | | | | |
|---|---|---|---|---|---|
| Minimap2 | -ax splice | 96.08 | 96.36 | 98.83 | 68.21 |
| Minimap2 + GTF | -ax splice –junc-bed anno.bed | 95.97 | 98.3 | 99.7 | 80.84 |
| deSALT | -d 10 -x ont2d -s 2 | 96.26 | 98.46 | 99.63 | 82.36 |
| deSALT + GTF | -d 10 -x ont2d -s 2 -G anno.info | 96.06 | 98.8 | 99.78 | 86.28 |
| **Fruit fly ONT 2D (1D$^2$), coverage = 30X, < 6 exons events, 172988 reads/284891590 bases/457490 exons** | | | | | |
| GMAP | -f samse --cross-species -z sense_force | 93.89 | 74.26 | 80.99 | 54.83 |
| Graphmap2 | -x rnaseq | 95.26 | 91.99 | 97.28 | 81.92 |
| Minimap2 | -ax splice | 94.67 | 89.78 | 94.19 | 77.88 |
| Minimap2 + GTF | -ax splice –junc-bed anno.bed | 94.91 | 91.7 | 95.84 | 81.18 |
| deSALT | -d 10 -x ont2d -s 2 | 95.21 | 94.45 | 97.68 | 87 |
| deSALT + GTF | -d 10 -x ont2d -s 2 -G anno.info | 95.24 | 95.09 | 97.89 | 88.35 |
| **Fruit fly ONT 2D (1D$^2$), coverage = 30X, 6-9 exons events, 62337 reads/203862649 bases/455175 exons** | | | | | |
| GMAP | -f samse --cross-species -z sense_force | 94.66 | 75.8 | 66.71 | 30 |
| Graphmap2 | -x rnaseq | 95.43 | 94.98 | 97.61 | 76.4 |
| Minimap2 | -ax splice | 95.69 | 95.27 | 98.17 | 77.32 |
| Minimap2 + GTF | -ax splice –junc-bed anno.bed | 95.87 | 97.47 | 99.59 | 84.85 |
| deSALT | -d 10 -x ont2d -s 2 | 95.89 | 97.93 | 99.44 | 88.68 |
| deSALT + GTF | -d 10 -x ont2d -s 2 -G anno.info | 95.91 | 98.4 | 99.69 | 90.03 |
| **Fruit fly ONT 2D (1D$^2$), coverage = 30X, > 9 exons events, 53012 reads/275060750 bases/758300 exons** | | | | | |
| GMAP | -f samse --cross-species -z sense_force | 94.87 | 74.5 | 53.02 | 13.36 |
| Graphmap2 | -x rnaseq | 94.63 | 94.1 | 95.32 | 61.72 |
| Minimap2 | -ax splice | 96.17 | 96.43 | 98.82 | 68.84 |
| Minimap2 + GTF | -ax splice –junc-bed anno.bed | 95.98 | 98.32 | 99.66 | 81.29 |
| deSALT | -d 10 -x ont2d -s 2 | 96.33 | 98.6 | 99.89 | 83.42 |
| deSALT + GTF | -d 10 -x ont2d -s 2 -G anno.info | 96.05 | 98.84 | 99.74 | 86.8 |
| **Mouse ONT 2D (1D$^2$), coverage = 4X, < 6 exons events, 34108 reads/33184744 bases/79605 exons** | | | | | |
| GMAP | -f samse --cross-species -z sense_force | 89.85 | 64.92 | 74.43 | 48.65 |
| Graphmap2 | -x rnaseq | 91.35 | 76.94 | 86.31 | 63.84 |
| Minimap2 | -ax splice | 90.58 | 73.75 | 77.37 | 59.24 |
| Minimap2 + GTF | -ax splice –junc-bed anno.bed | 90.65 | 75.29 | 78.63 | 60.67 |
| deSALT | -d 10 -x ont2d -s 2 | 92.59 | 83.27 | 89.16 | 71.22 |
| deSALT + GTF | -d 10 -x ont2d -s 2 -G anno.info | 92.75 | 85.75 | 90.64 | 75.33 |

| | | | | | |
|---|---|---|---|---|---|
| **Mouse ONT 2D (1D$^2$), coverage = 4X, 6-9 exons events, 6186 reads/10762016 bases/44279 exons** | | | | | |
| GMAP | -f samse --cross-species -z sense_force | 89.2 | 67.6 | 56.19 | 23.7 |
| Graphmap2 | -x rnaseq | 88.91 | 82.01 | 82.5 | 48.78 |
| Minimap2 | -ax splice | 91.05 | 85.64 | 87.75 | 58.62 |
| Minimap2 + GTF | -ax splice –junc-bed anno.bed | 93.27 | 90.62 | 93.54 | 65.37 |
| deSALT | -d 10 -x ont2d -s 2 | 93.47 | 92.37 | 94.86 | 73.86 |
| deSALT + GTF | -d 10 -x ont2d -s 2 -G anno.info | 93.97 | 93.18 | 95.53 | 74.6 |
| **Mouse ONT 2D (1D$^2$), coverage = 4X, > 9 exons events, 6297 reads/21637571 bases/108257 exons** | | | | | |
| GMAP | -f samse --cross-species -z sense_force | 92.19 | 71.55 | 44.66 | 10.1 |
| Graphmap2 | -x rnaseq | 86.65 | 82.2 | 79.45 | 34.95 |
| Minimap2 | -ax splice | 94.42 | 94.89 | 95.74 | 60.79 |
| Minimap2 + GTF | -ax splice –junc-bed anno.bed | 94.99 | 96.86 | 97.84 | 68.91 |
| deSALT | -d 10 -x ont2d -s 2 | 95.34 | 97.18 | 97.92 | 74.48 |
| deSALT + GTF | -d 10 -x ont2d -s 2 -G anno.info | 95.4 | 97.75 | 98.38 | 78.84 |
| **Mouse ONT 2D (1D$^2$), coverage = 10X, < 6 exons events, 71315 reads/81729449 bases/179515 exons** | | | | | |
| GMAP | -f samse --cross-species -z sense_force | 89.92 | 66.47 | 75.33 | 49.35 |
| Graphmap2 | -x rnaseq | 91.54 | 80.19 | 88.64 | 67.32 |
| Minimap2 | -ax splice | 91.33 | 79.22 | 86.43 | 66.36 |
| Minimap2 + GTF | -ax splice –junc-bed anno.bed | 91.82 | 81.01 | 87.88 | 68.02 |
| deSALT | -d 10 -x ont2d -s 2 | 92.82 | 87.65 | 93.06 | 77.56 |
| deSALT + GTF | -d 10 -x ont2d -s 2 -G anno.info | 92.9 | 89.12 | 93.77 | 80.39 |
| **Mouse ONT 2D (1D$^2$), coverage = 10X, 6-9 exons events, 15705 reads/27421329 bases/112575 exons** | | | | | |
| GMAP | -f samse --cross-species -z sense_force | 90.17 | 68.71 | 57.19 | 24.43 |
| Graphmap2 | -x rnaseq | 89.04 | 83.45 | 83.68 | 53.03 |
| Minimap2 | -ax splice | 92.08 | 86.5 | 88.67 | 59.91 |
| Minimap2 + GTF | -ax splice –junc-bed anno.bed | 93.43 | 90.76 | 93.77 | 65.94 |
| deSALT | -d 10 -x ont2d -s 2 | 94.44 | 93.56 | 95.46 | 77.48 |
| deSALT + GTF | -d 10 -x ont2d -s 2 -G anno.info | 94.49 | 93.84 | 95.7 | 78.39 |
| **Mouse ONT 2D (1D$^2$), coverage = 10X, > 9 exons events, 15859 reads/54611806 bases/273794 exons** | | | | | |
| GMAP | -f samse --cross-species -z sense_force | 92.56 | 71.78 | 45.03 | 10.71 |
| Graphmap2 | -x rnaseq | 86.63 | 83.55 | 81.35 | 39.34 |
| Minimap2 | -ax splice | 94.73 | 95.01 | 95.94 | 61.59 |

| | | | | | |
|---|---|---|---|---|---|
| Minimap2 + GTF | -ax splice –junc-bed anno.bed | 95.07 | 97.01 | 98.12 | 70.78 |
| deSALT | -d 10 -x ont2d -s 2 | 95.83 | 97.86 | 98.47 | 78.75 |
| deSALT + GTF | -d 10 -x ont2d -s 2 -G anno.info | 95.56 | 98.13 | 98.72 | 82.25 |
| **Mouse ONT 2D (1D$^2$), coverage = 30X, < 6 exons events, 204708 reads/242324237 bases/521418 exons** | | | | | |
| GMAP | -f samse --cross-species -z sense_force | 90.12 | 66.84 | 74.97 | 49.55 |
| Graphmap2 | -x rnaseq | 91.62 | 81.03 | 89.06 | 68.23 |
| Minimap2 | -ax splice | 91.56 | 80 | 86.86 | 66.91 |
| Minimap2 + GTF | -ax splice –junc-bed anno.bed | 92.04 | 81.83 | 88.41 | 68.67 |
| deSALT | -d 10 -x ont2d -s 2 | 92.85 | 88.66 | 93.38 | 79.34 |
| deSALT + GTF | -d 10 -x ont2d -s 2 -G anno.info | 92.91 | 89.62 | 93.8 | 81.15 |
| **Mouse ONT 2D (1D$^2$) , coverage = 30X, 6-9 exons events, 46551 reads/82067115 bases/333544 exons** | | | | | |
| GMAP | -f samse --cross-species -z sense_force | 90.85 | 68.82 | 56.81 | 24.85 |
| Graphmap2 | -x rnaseq | 89.2 | 83.79 | 84.44 | 53.89 |
| Minimap2 | -ax splice | 92.54 | 86.55 | 88.81 | 60.26 |
| Minimap2 + GTF | -ax splice –junc-bed anno.bed | 93.44 | 90.89 | 93.84 | 66.49 |
| deSALT | -d 10 -x ont2d -s 2 | 95.05 | 93.93 | 95.9 | 78.17 |
| deSALT + GTF | -d 10 -x ont2d -s 2 -G anno.info | 95.08 | 94.14 | 96.04 | 79.09 |
| **Mouse ONT 2D (1D$^2$), coverage = 30X, > 9 exons events, 48154 reads/166937719 bases/833114 exons** | | | | | |
| GMAP | -f samse --cross-species -z sense_force | 92.12 | 71.35 | 44.75 | 10.42 |
| Graphmap2 | -x rnaseq | 86.59 | 83.71 | 81.28 | 40.17 |
| Minimap2 | -ax splice | 94.38 | 94.75 | 95.61 | 61.45 |
| Minimap2 + GTF | -ax splice –junc-bed anno.bed | 94.99 | 96.96 | 98.06 | 70.58 |
| deSALT | -d 10 -x ont2d -s 2 | 95.39 | 97.62 | 98.4 | 78.73 |
| deSALT + GTF | -d 10 -x ont2d -s 2 -G anno.info | 95.5 | 98.18 | 98.69 | 82.72 |
| **Human ONT 2D (1D$^2$), coverage = 4X, < 6 exons events, 34832 reads/28480470 bases/85272 exons** | | | | | |
| GMAP | -f samse --cross-species -z sense_force | 89.73 | 65.78 | 73.14 | 48.58 |
| Graphmap2 | -x rnaseq | 91.64 | 77.8 | 85.59 | 63.52 |
| Minimap2 | -ax splice | 89.62 | 73.95 | 75.41 | 57.82 |
| Minimap2 + GTF | -ax splice –junc-bed anno.bed | 90.79 | 75.39 | 76.63 | 59.38 |
| deSALT | -d 10 -x ont2d -s 2 | 93.08 | 84.52 | 88.9 | 72.05 |
| deSALT + GTF | -d 10 -x ont2d -s 2 -G anno.info | 93.26 | 86.98 | 90.64 | 76.07 |
| **Human ONT 2D (1D$^2$), coverage = 4X, 6-9 exons events, 6042 reads/9765619 bases/42767 exons** | | | | | |

| | | | | | |
|---|---|---|---|---|---|
| GMAP | -f samse --cross-species -z sense_force | 91.44 | 69.62 | 57.12 | 24.96 |
| Graphmap2 | -x rnaseq | 88.26 | 82.37 | 91.94 | 49.67 |
| Minimap2 | -ax splice | 93.35 | 88.32 | 90.75 | 59.98 |
| Minimap2 + GTF | -ax splice –junc-bed anno.bed | 93.35 | 90.76 | 93.26 | 64.83 |
| deSALT | -d 10 -x ont2d -s 2 | 95.64 | 94.29 | 96.69 | 74.5 |
| deSALT + GTF | -d 10 -x ont2d -s 2 -G anno.info | 95.83 | 95.14 | 97.67 | 77.31 |
| **Human ONT 2D (1D$^2$), coverage = 4X, > 9 exons events, 5720 reads/18358704 bases/94986 exons** | | | | | |
| GMAP | -f samse --cross-species -z sense_force | 91.81 | 71.59 | 45.1 | 9.48 |
| Graphmap2 | -x rnaseq | 85.91 | 82.37 | 79.72 | 35.21 |
| Minimap2 | -ax splice | 94.26 | 94.75 | 96.24 | 63.18 |
| Minimap2 + GTF | -ax splice –junc-bed anno.bed | 95.02 | 96.81 | 98.24 | 70.55 |
| deSALT | -d 10 -x ont2d -s 2 | 95.67 | 97.24 | 98.64 | 75.84 |
| deSALT + GTF | -d 10 -x ont2d -s 2 -G anno.info | 95.35 | 97.69 | 98.73 | 79.04 |
| **Human ONT 2D (1D$^2$), coverage = 10X, < 6 exons events, 73054 reads/69878562 bases/193650 exons** | | | | | |
| GMAP | -f samse --cross-species -z sense_force | 90.78 | 67.28 | 73.9 | 49.47 |
| Graphmap2 | -x rnaseq | 92.04 | 80.88 | 87.95 | 66.77 |
| Minimap2 | -ax splice | 91.69 | 79.78 | 85.76 | 65.67 |
| Minimap2 + GTF | -ax splice –junc-bed anno.bed | 91.81 | 81.26 | 87.15 | 67.35 |
| deSALT | -d 10 -x ont2d -s 2 | 94.32 | 89.3 | 93.71 | 78.92 |
| deSALT + GTF | -d 10 -x ont2d -s 2 -G anno.info | 94.42 | 90.93 | 94.67 | 82.07 |
| **Human ONT 2D (1D$^2$), coverage = 10X, 6-9 exons events, 15404 reads/24970788 bases/109140 exons** | | | | | |
| GMAP | -f samse --cross-species -z sense_force | 89.96 | 68.63 | 56.58 | 24.38 |
| Graphmap2 | -x rnaseq | 88.08 | 83.16 | 83.07 | 51.87 |
| Minimap2 | -ax splice | 92.03 | 87.32 | 89.94 | 59.13 |
| Minimap2 + GTF | -ax splice –junc-bed anno.bed | 93.24 | 90.43 | 93.14 | 64.57 |
| deSALT | -d 10 -x ont2d -s 2 | 94.44 | 94.04 | 96.41 | 76.69 |
| deSALT + GTF | -d 10 -x ont2d -s 2 -G anno.info | 94.53 | 94.5 | 96.75 | 78.26 |
| **Human ONT 2D (1D$^2$), coverage = 10X, > 9 exons events, 14462 reads/46434095 bases/240227 exons** | | | | | |
| GMAP | -f samse --cross-species -z sense_force | 91.51 | 71.05 | 44.27 | 10.28 |
| Graphmap2 | -x rnaseq | 85.97 | 83.24 | 81.36 | 36.68 |
| Minimap2 | -ax splice | 94.19 | 94.56 | 96.18 | 61.19 |
| Minimap2 + GTF | -ax splice –junc-bed anno.bed | 95.01 | 96.65 | 98.23 | 68.74 |

| | | | | | |
|---|---|---|---|---|---|
| deSALT | -d 10 -x ont2d -s 2 | 95.67 | 97.45 | 98.88 | 76.75 |
| deSALT + GTF | -d 10 -x ont2d -s 2 -G anno.info | 95.3 | 97.56 | 98.65 | 78.95 |
| **Human ONT 2D (1D$^2$), coverage = 30X, < 6 exons events, 209725 reads/206910570 bases/562271 exons** | | | | | |
| GMAP | -f samse --cross-species -z sense_force | 90.72 | 76.31 | 80.9 | 60.34 |
| Graphmap2 | -x rnaseq | 92.14 | 81.44 | 88.03 | 67.53 |
| Minimap2 | -ax splice | 91.73 | 80.34 | 85.8 | 66.08 |
| Minimap2 + GTF | -ax splice –junc-bed anno.bed | 91.93 | 81.82 | 87.22 | 67.85 |
| deSALT | -d 10 -x ont2d -s 2 | 94.29 | 90.16 | 93.91 | 80.6 |
| deSALT + GTF | -d 10 -x ont2d -s 2 -G anno.info | 94.38 | 91.27 | 94.59 | 82.66 |
| **Human ONT 2D (1D$^2$), coverage = 30X, 6-9 exons events, 45842 reads/75221616 bases/324667 exons** | | | | | |
| GMAP | -f samse --cross-species -z sense_force | 90.55 | 77.96 | 70.31 | 35.68 |
| Graphmap2 | -x rnaseq | 88.21 | 83.38 | 93.18 | 52.72 |
| Minimap2 | -ax splice | 92.58 | 87.8 | 90.44 | 59.93 |
| Minimap2 + GTF | -ax splice –junc-bed anno.bed | 93.31 | 90.58 | 93.37 | 65.16 |
| deSALT | -d 10 -x ont2d -s 2 | 95.01 | 94.68 | 96.92 | 77.87 |
| deSALT + GTF | -d 10 -x ont2d -s 2 -G anno.info | 95.07 | 95.05 | 97.31 | 79.07 |
| **Human ONT 2D (1D$^2$), coverage = 30X, > 9 exons events, 43830 reads/141765970 bases/728962 exons** | | | | | |
| GMAP | -f samse --cross-species -z sense_force | 91.53 | 81.64 | 65.6 | 18.49 |
| Graphmap2 | -x rnaseq | 85.88 | 82.93 | 80.46 | 36.82 |
| Minimap2 | -ax splice | 94.15 | 94.64 | 96.2 | 61.94 |
| Minimap2 + GTF | -ax splice –junc-bed anno.bed | 95.06 | 96.79 | 98.23 | 69.87 |
| deSALT | -d 10 -x ont2d -s 2 | 95.57 | 97.4 | 98.63 | 76.84 |
| deSALT + GTF | -d 10 -x ont2d -s 2 -G anno.info | 95.6 | 97.68 | 98.81 | 79.21 |
| **Human all protein coding genes ONT 2D (1D$^2$), < 6 exons events, 583561 reads/ 886607912 bases/ 1797858 exons** | | | | | |
| GMAP | -f samse --cross-species -z sense_force | 94.66 | 69.52 | 71.32 | 48.09 |
| Graphmap2 | -x rnaseq | 96.75 | 82.18 | 86.5 | 65.16 |
| Minimap2 | -ax splice | 96.37 | 83.69 | 88.3 | 68.41 |
| Minimap2 + GTF | -ax splice –junc-bed anno.bed | 96.57 | 84.98 | 89.67 | 69.9 |
| deSALT | -d 10 -x ont2d -s 2 | 97.62 | 90.63 | 94.04 | 79.4 |
| deSALT + GTF | -d 10 -x ont2d -s 2 -G anno.info | 97.64 | 91.07 | 94.51 | 80.12 |
| **Human all protein coding genes ONT 2D (1D$^2$), 6-9 exons events, 238407 reads/ 481760161 bases/ 1716649 exons** | | | | | |
| GMAP | -f samse --cross-species -z sense_force | 94.11 | 70.18 | 57.77 | 25.22 |

| | | | | | |
|---|---|---|---|---|---|
| Graphmap2 | -x rnaseq | 92.33 | 83.13 | 82.85 | 51.35 |
| Minimap2 | -ax splice | 96.52 | 89.75 | 92.25 | 65.19 |
| Minimap2 + GTF | -ax splice –junc-bed anno.bed | 97.12 | 92.14 | 94.8 | 69.91 |
| deSALT | -d 10 -x ont2d -s 2 | 98.2 | 95.46 | 97.46 | 80.35 |
| deSALT + GTF | -d 10 -x ont2d -s 2 -G anno.info | 98.24 | 96.01 | 97.84 | 82.33 |
| **Human all protein coding genes ONT 2D (1D$^2$), > 9 exons events, 279718 reads/ 1046335651 bases/ 4907573 exons** | | | | | |
| GMAP | -f samse --cross-species -z sense_force | 94.15 | 70.93 | 44.49 | 9.44 |
| Graphmap2 | -x rnaseq | 87.53 | 79.76 | 76.13 | 33.1 |
| Minimap2 | -ax splice | 97.56 | 94.87 | 96.29 | 63.73 |
| Minimap2 + GTF | -ax splice –junc-bed anno.bed | 97.97 | 96.77 | 98.05 | 71.89 |
| deSALT | -d 10 -x ont2d -s 2 | 98.54 | 97.65 | 98.57 | 79.36 |
| deSALT + GTF | -d 10 -x ont2d -s 2 -G anno.info | 98.54 | 98.11 | 98.77 | 83.55 |
| **Simulated PacBio subread datasets** | | | | | |
| **Fruit fly PacBio subread, coverage = 4X, < 6 exons events, 30278 reads/42410655 bases/69852 exons** | | | | | |
| GMAP | -f samse –cross-species -z sense_force | 77.3 | 54.04 | 56.02 | 36.73 |
| Graphmap2 | -x rnaseq | 87.81 | 85.87 | 93.79 | 73.26 |
| Minimap2 | -ax splice | 85.67 | 80.26 | 78.86 | 63.38 |
| Minimap2 + GTF | -ax splice –junc-bed anno.bed | 86.51 | 82.82 | 80.51 | 66.83 |
| deSALT | -d 10 -x clr -s 2 | 87.06 | 88.58 | 93.84 | 77.42 |
| deSALT + GTF | -d 10 -x clr -s 2 -G anno.info | 87.12 | 89.82 | 94.61 | 79.65 |
| **Fruit fly PacBio subread, coverage = 4X, 6-9 exons events, 7707 reads/26875546 bases/56191 exons** | | | | | |
| GMAP | -f samse –cross-species -z sense_force | 85.99 | 66.81 | 51.81 | 15.67 |
| Graphmap2 | -x rnaseq | 88.24 | 91.98 | 95.4 | 63.64 |
| Minimap2 | -ax splice | 89.52 | 95.21 | 98.31 | 74.35 |
| Minimap2 + GTF | -ax splice –junc-bed anno.bed | 88.6 | 97.12 | 99.43 | 82.58 |
| deSALT | -d 10 -x clr -s 2 | 89.72 | 97.55 | 99.68 | 83.95 |
| deSALT + GTF | -d 10 -x clr -s 2 -G anno.info | 89.74 | 97.89 | 99.92 | 85.62 |
| **Fruit fly PacBio subread, coverage = 4X, > 9 exons events, 6184 reads/32658929 bases/86374 exons** | | | | | |
| GMAP | -f samse –cross-species -z sense_force | 84.61 | 66.39 | 34.22 | 4.03 |
| Graphmap2 | -x rnaseq | 87.54 | 92.1 | 93.79 | 49.93 |

| | | | | | |
|---|---|---|---|---|---|
| Minimap2 | -ax splice | 88.83 | 96.34 | 98.93 | 66.51 |
| Minimap2 + GTF | -ax splice –junc-bed anno.bed | 88.75 | 98.08 | 99.76 | 79.17 |
| deSALT | -d 10 -x clr -s 2 | 89.02 | 98.57 | 99.92 | 80.4 |
| deSALT + GTF | -d 10 -x clr -s 2 -G anno.info | 89.03 | 98.84 | 100 | 82.7 |
| **Fruit fly PacBio subread, coverage = 10X, < 6 exons events, 63048 reads/104050616 bases/159211 exons** | | | | | |
| GMAP | -f samse –cross-species -z sense_force | 78.16 | 56.02 | 56.77 | 36.86 |
| Graphmap2 | -x rnaseq | 87.91 | 89.17 | 96.26 | 76.71 |
| Minimap2 | -ax splice | 87.3 | 87.06 | 91.54 | 73.09 |
| Minimap2 + GTF | -ax splice –junc-bed anno.bed | 87.55 | 89.43 | 93.26 | 77.12 |
| deSALT | -d 10 -x clr -s 2 | 87.98 | 92.35 | 96.82 | 82.58 |
| deSALT + GTF | -d 10 -x clr -s 2 -G anno.info | 88 | 92.99 | 97.2 | 83.85 |
| **Fruit fly PacBio subread, coverage = 10X, 6-9 exons events, 19163 reads/66676969 bases/139521 exons** | | | | | |
| GMAP | -f samse –cross-species -z sense_force | 85.12 | 66.6 | 51.14 | 15.66 |
| Graphmap2 | -x rnaseq | 88.19 | 93.43 | 97.01 | 68.83 |
| Minimap2 | -ax splice | 88.82 | 95.05 | 98.19 | 74.38 |
| Minimap2 + GTF | -ax splice –junc-bed anno.bed | 88.66 | 97.15 | 99.53 | 82.9 |
| deSALT | -d 10 -x clr -s 2 | 89.01 | 98 | 99.62 | 86.73 |
| deSALT + GTF | -d 10 -x clr -s 2 -G anno.info | 89.02 | 98.18 | 99.73 | 87.58 |
| **Fruit fly PacBio subread, coverage = 10X, > 9 exons events, 15699 reads/83886149 bases/220066 exons** | | | | | |
| GMAP | -f samse –cross-species -z sense_force | 84.67 | 65.63 | 33.26 | 4.05 |
| Graphmap2 | -x rnaseq | 87.55 | 92.97 | 94.51 | 55.11 |
| Minimap2 | -ax splice | 88.68 | 95.89 | 98.6 | 66.44 |
| Minimap2 + GTF | -ax splice –junc-bed anno.bed | 88.76 | 98.15 | 99.71 | 79.63 |
| deSALT | -d 10 -x clr -s 2 | 88.88 | 98.31 | 99.68 | 82.14 |
| deSALT + GTF | -d 10 -x clr -s 2 -G anno.info | 88.84 | 98.62 | 99.71 | 84.53 |
| **Fruit fly PacBio subread, coverage = 30X, < 6 exons events, 172125 reads/306614209 bases/452111 exons** | | | | | |
| GMAP | -f samse –cross-species -z sense_force | 79.26 | 58.07 | 58.55 | 37.94 |
| Graphmap2 | -x rnaseq | 87.99 | 90.81 | 96.91 | 79.34 |
| Minimap2 | -ax splice | 87.76 | 89.15 | 94.56 | 76.3 |
| Minimap2 + GTF | -ax splice –junc-bed anno.bed | 87.87 | 91.45 | 96.33 | 80.38 |
| deSALT | -d 10 -x clr -s 2 | 88.18 | 93.68 | 97.53 | 84.8 |
| deSALT + GTF | -d 10 -x clr -s 2 -G anno.info | 88.18 | 94.08 | 97.66 | 85.57 |

| | | | | | |
|---|---|---|---|---|---|
| **Fruit fly PacBio subread, coverage = 30X, 6-9 exons events, 57839 reads/201638578 bases/421444 exons** | | | | | |
| GMAP | -f samse –cross-species -z sense_force | 85.16 | 66.46 | 51.19 | 15.43 |
| Graphmap2 | -x rnaseq | 88.24 | 93.84 | 97.02 | 70.74 |
| Minimap2 | -ax splice | 88.83 | 94.95 | 98.15 | 74.32 |
| Minimap2 + GTF | -ax splice –junc-bed anno.bed | 88.66 | 97.17 | 99.5 | 82.89 |
| deSALT | -d 10 -x clr -s 2 | 89.03 | 98.05 | 99.66 | 86.92 |
| deSALT + GTF | -d 10 -x clr -s 2 -G anno.info | 89.03 | 98.22 | 99.72 | 87.77 |
| **Fruit fly PacBio subread, coverage = 30X, > 9 exons events, 47577 reads/255545996 bases/667078 exons** | | | | | |
| GMAP | -f samse –cross-species -z sense_force | 84.52 | 65.91 | 33.88 | 3.87 |
| Graphmap2 | -x rnaseq | 87.64 | 93.31 | 94.84 | 56.91 |
| Minimap2 | -ax splice | 88.52 | 95.95 | 98.59 | 66.85 |
| Minimap2 + GTF | -ax splice –junc-bed anno.bed | 88.77 | 98.19 | 99.73 | 79.99 |
| deSALT | -d 10 -x clr -s 2 | 88.72 | 98.39 | 99.69 | 82.09 |
| deSALT + GTF | -d 10 -x clr -s 2 -G anno.info | 88.86 | 98.66 | 99.77 | 84.83 |
| **Mouse PacBio subread, coverage = 4X, < 6 exons events, 35080 reads/34447409 bases/78452 exons** | | | | | |
| GMAP | -f samse –cross-species -z sense_force | 68.44 | 37.59 | 46.23 | 29.75 |
| Graphmap2 | -x rnaseq | 84.32 | 74.64 | 85.5 | 60.31 |
| Minimap2 | -ax splice | 82.79 | 68.74 | 69.99 | 51.76 |
| Minimap2 + GTF | -ax splice –junc-bed anno.bed | 83.01 | 70.38 | 71.13 | 53.19 |
| deSALT | -d 10 -x clr -s 2 | 85.21 | 80.67 | 86.16 | 66.92 |
| deSALT + GTF | -d 10 -x clr -s 2 -G anno.info | 85.29 | 82.76 | 87.69 | 70.13 |
| **Mouse PacBio subread, coverage = 4X, 6-9 exons events, 5724 reads/10741011 bases/41117 exons** | | | | | |
| GMAP | -f samse –cross-species -z sense_force | 75.18 | 50.73 | 36.25 | 11.2 |
| Graphmap2 | -x rnaseq | 83.12 | 80.66 | 91.8 | 42.91 |
| Minimap2 | -ax splice | 84.52 | 84.86 | 87.46 | 54 |
| Minimap2 + GTF | -ax splice –junc-bed anno.bed | 86.18 | 89.65 | 93.12 | 60.94 |
| deSALT | -d 10 -x clr -s 2 | 86.97 | 91.68 | 94.13 | 70.18 |
| deSALT + GTF | -d 10 -x clr -s 2 -G anno.info | 87.08 | 92.61 | 94.95 | 73.55 |
| **Mouse PacBio subread, coverage = 4X, > 9 exons events, 5677 reads/20567829 bases/97011 exons** | | | | | |
| GMAP | -f samse –cross-species -z sense_force | 83.33 | 65.01 | 27.85 | 3.61 |
| Graphmap2 | -x rnaseq | 80.82 | 91.22 | 78.25 | 28.61 |
| Minimap2 | -ax splice | 87.34 | 94.4 | 95.61 | 55.15 |

| | | | | | |
|---|---|---|---|---|---|
| Minimap2 + GTF | -ax splice –junc-bed anno.bed | 87.86 | 96.61 | 98.39 | 64.41 |
| deSALT | -d 10 -x clr -s 2 | 88.26 | 96.96 | 97.89 | 70.74 |
| deSALT + GTF | -d 10 -x clr -s 2 -G anno.info | 88.31 | 97.42 | 98.15 | 75.15 |
| **Mouse PacBio subread, coverage = 10X, < 6 exons events, 73842 reads/85200651 bases/179030 exons** | | | | | |
| GMAP | -f samse –cross-species -z sense_force | 68.37 | 37.28 | 44.81 | 28.69 |
| Graphmap2 | -x rnaseq | 84.65 | 77.67 | 87.74 | 62.82 |
| Minimap2 | -ax splice | 83.92 | 74.98 | 81.56 | 59.95 |
| Minimap2 + GTF | -ax splice –junc-bed anno.bed | 84.33 | 77.04 | 83.08 | 61.82 |
| deSALT | -d 10 -x clr -s 2 | 85.78 | 85.59 | 91.32 | 73.65 |
| deSALT + GTF | -d 10 -x clr -s 2 -G anno.info | 85.83 | 86.86 | 92.17 | 75.89 |
| **Mouse PacBio subread, coverage = 10X, 6-9 exons events, 14180 reads/51689732 bases/243516 exons** | | | | | |
| GMAP | -f samse –cross-species -z sense_force | 75.24 | 50.13 | 35.25 | 11.48 |
| Graphmap2 | -x rnaseq | 83.09 | 82.05 | 83.48 | 46.01 |
| Minimap2 | -ax splice | 85.14 | 85.32 | 87.88 | 54.86 |
| Minimap2 + GTF | -ax splice –junc-bed anno.bed | 86.29 | 89.87 | 93.34 | 61.39 |
| deSALT | -d 10 -x clr -s 2 | 87.68 | 93.1 | 95.41 | 74.48 |
| deSALT + GTF | -d 10 -x clr -s 2 -G anno.info | 87.73 | 93.47 | 95.75 | 76.02 |
| **Mouse PacBio subread, coverage = 10X, > 9 exons events, 17084 reads/56231420 bases/296347 exons** | | | | | |
| GMAP | -f samse –cross-species -z sense_force | 83.26 | 64.81 | 27.14 | 2.74 |
| Graphmap2 | -x rnaseq | 80.92 | 83.07 | 81.58 | 33.53 |
| Minimap2 | -ax splice | 87.27 | 94.34 | 95.83 | 55.73 |
| Minimap2 + GTF | -ax splice –junc-bed anno.bed | 87.82 | 96.6 | 98.24 | 65.03 |
| deSALT | -d 10 -x clr -s 2 | 88.31 | 97.45 | 98.52 | 74.97 |
| deSALT + GTF | -d 10 -x clr -s 2 -G anno.info | 88.34 | 98.01 | 98.89 | 79.03 |
| **Mouse PacBio subread, coverage = 30X, < 6 exons events, 202805 reads/253239643 bases/510067 exons** | | | | | |
| GMAP | -f samse –cross-species -z sense_force | 69.05 | 38.03 | 44.8 | 28.92 |
| Graphmap2 | -x rnaseq | 84.75 | 79.26 | 88.79 | 64.59 |
| Minimap2 | -ax splice | 84.43 | 77.88 | 86.28 | 63.48 |
| Minimap2 + GTF | -ax splice –junc-bed anno.bed | 84.87 | 79.88 | 87.9 | 65.44 |
| deSALT | -d 10 -x clr -s 2 | 85.9 | 87.74 | 93.4 | 76.81 |
| deSALT + GTF | -d 10 -x clr -s 2 -G anno.info | 85.93 | 88.54 | 93.79 | 78.16 |
| **Mouse PacBio subread, coverage = 30X, 6-9 exons events, 42947 reads/156407534 bases/737207 exons** | | | | | |

| | | | | | |
|---|---|---|---|---|---|
| GMAP | -f samse –cross-species -z sense_force | 75.42 | 50.47 | 35.51 | 11.03 |
| Graphmap2 | -x rnaseq | 83.19 | 82.71 | 84.33 | 48.14 |
| Minimap2 | -ax splice | 85.04 | 85.32 | 88.07 | 54.52 |
| Minimap2 + GTF | -ax splice –junc-bed anno.bed | 86.22 | 89.73 | 93.36 | 60.94 |
| deSALT | -d 10 -x clr -s 2 | 87.55 | 93.31 | 95.58 | 74.96 |
| deSALT + GTF | -d 10 -x clr -s 2 -G anno.info | 87.57 | 93.55 | 95.67 | 76.06 |
| **Mouse PacBio subread, coverage = 30X, > 9 exons events, 51149 reads/169009353 bases/889311 exons** | | | | | |
| GMAP | -f samse –cross-species -z sense_force | 83.48 | 65.11 | 27.6 | 3.09 |
| Graphmap2 | -x rnaseq | 80.87 | 83.22 | 81.83 | 34.4 |
| Minimap2 | -ax splice | 87.44 | 94.49 | 95.94 | 55.62 |
| Minimap2 + GTF | -ax splice –junc-bed anno.bed | 87.82 | 96.56 | 98.21 | 64.91 |
| deSALT | -d 10 -x clr -s 2 | 88.43 | 97.58 | 98.53 | 74.89 |
| deSALT + GTF | -d 10 -x clr -s 2 -G anno.info | 88.34 | 98.06 | 98.88 | 80.12 |
| **Human PacBio subread, coverage = 4X, < 6 exons events, 35848 reads/29729832 bases/84394 exons** | | | | | |
| GMAP | -f samse –cross-species -z sense_force | 64.19 | 34.61 | 42.4 | 27.72 |
| Graphmap2 | -x rnaseq | 85.27 | 76.49 | 86.17 | 61.51 |
| Minimap2 | -ax splice | 82.41 | 68.55 | 67.9 | 49.68 |
| Minimap2 + GTF | -ax splice –junc-bed anno.bed | 83.38 | 72.62 | 74.51 | 55.38 |
| deSALT | -d 10 -x clr -s 2 | 86.22 | 81.56 | 85.51 | 67.07 |
| deSALT + GTF | -d 10 -x clr -s 2 -G anno.info | 86.41 | 83.93 | 87.4 | 70.55 |
| **Human PacBio subread, coverage = 4X, 6-9 exons events, 5522 reads/9604060 bases/39360 exons** | | | | | |
| GMAP | -f samse –cross-species -z sense_force | 73.23 | 49.35 | 34.66 | 10.41 |
| Graphmap2 | -x rnaseq | 82.42 | 81.17 | 82.03 | 40.69 |
| Minimap2 | -ax splice | 84.69 | 87.27 | 90.22 | 55.12 |
| Minimap2 + GTF | -ax splice –junc-bed anno.bed | 85.68 | 89.08 | 93.15 | 55.39 |
| deSALT | -d 10 -x clr -s 2 | 86.88 | 93.59 | 96.32 | 71.86 |
| deSALT + GTF | -d 10 -x clr -s 2 -G anno.info | 86.99 | 94.41 | 97.23 | 74.81 |
| **Human PacBio subread, coverage = 4X, > 9 exons events, 5167 reads/17475908 bases/84665 exons** | | | | | |
| GMAP | -f samse –cross-species -z sense_force | 82.76 | 64.46 | 26.96 | 2.32 |
| Graphmap2 | -x rnaseq | 81.1 | 82.2 | 79.91 | 28.69 |
| Minimap2 | -ax splice | 87.27 | 94.56 | 97.08 | 56.82 |
| Minimap2 + GTF | -ax splice –junc-bed anno.bed | 87.51 | 94.99 | 97.81 | 55.22 |

| | | | | | |
|---|---|---|---|---|---|
| deSALT | -d 10 -x clr -s 2 | 88.34 | 96.87 | 98.55 | 71.51 |
| deSALT + GTF | -d 10 -x clr -s 2 -G anno.info | 88.41 | 97.28 | 98.86 | 74.72 |
| **Human PacBio subread, coverage = 10X, < 6 exons events, 75600 reads/73059595 bases/192890 exons** | | | | | |
| GMAP | -f samse –cross-species -z sense_force | 63.84 | 34.39 | 40.36 | 26.37 |
| Graphmap2 | -x rnaseq | 85.52 | 78.54 | 88.11 | 63.21 |
| Minimap2 | -ax splice | 83.69 | 75.08 | 80.05 | 57.98 |
| Minimap2 + GTF | -ax splice –junc-bed anno.bed | 84.46 | 77.18 | 83.54 | 61.5 |
| deSALT | -d 10 -x clr -s 2 | 86.87 | 87.01 | 91.81 | 74.16 |
| deSALT + GTF | -d 10 -x clr -s 2 -G anno.info | 86.95 | 88.44 | 92.9 | 76.61 |
| **Human PacBio subread, coverage = 10X, 6-9 exons events, 13999 reads/24325764 bases/99703 exons** | | | | | |
| GMAP | -f samse –cross-species -z sense_force | 73.59 | 48.48 | 33.8 | 10.38 |
| Graphmap2 | -x rnaseq | 82.42 | 82.66 | 83.69 | 44.5 |
| Minimap2 | -ax splice | 85.48 | 87.17 | 90.41 | 55.2 |
| Minimap2 + GTF | -ax splice –junc-bed anno.bed | 85.76 | 89.19 | 93.16 | 55.72 |
| deSALT | -d 10 -x clr -s 2 | 87.89 | 94.52 | 97.3 | 75.15 |
| deSALT + GTF | -d 10 -x clr -s 2 -G anno.info | 87.97 | 95.06 | 97.78 | 77.13 |
| **Human PacBio subread, coverage = 10X, > 9 exons events, 12934 reads/43940986 bases/212358 exons** | | | | | |
| GMAP | -f samse --cross-species -z sense_force | 82.97 | 64.4 | 26.09 | 2.61 |
| Graphmap2 | -x rnaseq | 81.15 | 83.4 | 82.02 | 31.48 |
| Minimap2 | -ax splice | 87.32 | 94.52 | 96.76 | 56.77 |
| Minimap2 + GTF | -ax splice –junc-bed anno.bed | 87.43 | 94.89 | 97.83 | 54.19 |
| deSALT | -d 10 -x clr -s 2 | 88.58 | 97.34 | 98.79 | 74.12 |
| deSALT + GTF | -d 10 -x clr -s 2 -G anno.info | 88.61 | 97.58 | 98.87 | 76.16 |
| **Human PacBio subread, coverage = 30X, < 6 exons events, 208020 reads/217338313 bases/552141 exons** | | | | | |
| GMAP | -f samse --cross-species -z sense_force | 64.47 | 34.56 | 39.63 | 26.19 |
| Graphmap2 | -x rnaseq | 85.61 | 79.85 | 88.94 | 64.69 |
| Minimap2 | -ax splice | 84.51 | 78.07 | 85.3 | 61.76 |
| Minimap2 + GTF | -ax splice –junc-bed anno.bed | 84.91 | 79.46 | 87.55 | 64.42 |
| deSALT | -d 10 -x clr -s 2 | 87.34 | 89.44 | 94.37 | 77.77 |
| deSALT + GTF | -d 10 -x clr -s 2 -G anno.info | 87.39 | 90.45 | 95.06 | 79.57 |
| **Human PacBio subread, coverage = 30X, 6-9 exons events, 42098 reads/73146856 bases/300035 exons** | | | | | |
| GMAP | -f samse --cross-species -z sense_force | 73.9 | 49.04 | 33.94 | 10.93 |

| | | | | | |
|---|---|---|---|---|---|
| Graphmap2 | -x rnaseq | 82.39 | 82.77 | 84.01 | 45.25 |
| Minimap2 | -ax splice | 85.59 | 87.31 | 90.4 | 55.44 |
| Minimap2 + GTF | -ax splice –junc-bed anno.bed | 85.76 | 89.3 | 93.34 | 55.96 |
| deSALT | -d 10 -x clr -s 2 | 88.06 | 94.75 | 97.38 | 75.64 |
| deSALT + GTF | -d 10 -x clr -s 2 -G anno.info | 88.1 | 95.16 | 97.72 | 77.26 |
| **Human PacBio subread, coverage = 30X, > 9 exons events, 39257 reads/133356721 bases/644800 exons** | | | | | |
| GMAP | -f samse --cross-species -z sense_force | 82.45 | 64.31 | 26.67 | 2.77 |
| Graphmap2 | -x rnaseq | 81.27 | 83.72 | 82.33 | 32.43 |
| Minimap2 | -ax splice | 87 | 94.31 | 96.63 | 56.49 |
| Minimap2 + GTF | -ax splice –junc-bed anno.bed | 87.44 | 94.95 | 97.82 | 55.15 |
| deSALT | -d 10 -x clr -s 2 | 88.27 | 97.2 | 98.67 | 73.42 |
| deSALT + GTF | -d 10 -x clr -s 2 -G anno.info | 88.28 | 97.43 | 98.71 | 75.98 |
| **Human all protein coding genes PacBio subread, < 6 exons events, 856433 reads/1205761883 bases/2233187 exons** | | | | | |
| GMAP | -f samse --cross-species -z sense_force | 75.02 | 42.64 | 52.92 | 34.4 |
| Graphmap2 | -x rnaseq | 87.41 | 79.34 | 87.13 | 62.65 |
| Minimap2 | -ax splice | 87.25 | 80.71 | 88.79 | 66.24 |
| Minimap2 + GTF | -ax splice –junc-bed anno.bed | 87.42 | 81.96 | 89.88 | 67.52 |
| deSALT | -d 10 -x clr -s 2 | 88.55 | 89.08 | 94.76 | 77.59 |
| deSALT + GTF | -d 10 -x clr -s 2 -G anno.info | 88.56 | 89.53 | 95.12 | 78.28 |
| **Human all protein coding genes PacBio subread, 6-9 exons events, 277042 reads/472298204 bases/2002689 exons** | | | | | |
| GMAP | -f samse --cross-species -z sense_force | 75.53 | 52.43 | 36.68 | 11.22 |
| Graphmap2 | -x rnaseq | 84.15 | 81.12 | 80.37 | 38.96 |
| Minimap2 | -ax splice | 86.51 | 87.81 | 91.52 | 54.17 |
| Minimap2 + GTF | -ax splice –junc-bed anno.bed | 87.11 | 90.03 | 94.12 | 58.54 |
| deSALT | -d 10 -x clr -s 2 | 88.74 | 94.37 | 97.58 | 74.69 |
| deSALT + GTF | -d 10 -x clr -s 2 -G anno.info | 88.76 | 95.31 | 97.99 | 76.94 |
| **Human all protein coding genes PacBio subread, > 9 exons events, 256875 reads/737935259 bases/3921025 exons** | | | | | |
| GMAP | -f samse --cross-species -z sense_force | 82.34 | 63.46 | 28.15 | 3.16 |
| Graphmap2 | -x rnaseq | 81.08 | 79.87 | 75.01 | 25.03 |
| Minimap2 | -ax splice | 87.72 | 92.87 | 95.26 | 49.51 |
| Minimap2 + GTF | -ax splice –junc-bed anno.bed | 88.1 | 94.79 | 97.31 | 56.19 |
| deSALT | -d 10 -x clr -s 2 | 89.04 | 96.68 | 98.35 | 71.22 |

| | | | | | |
|---|---|---|---|---|---|
| deSALT + GTF | -d 10 -x clr -s 2 -G anno.info | 89.01 | 97.2 | 98.56 | 75.73 |
| **Simulated ONT 1D datasets** | | | | | |
| **Fruit fly ONT 1D, coverage = 4X, < 6 exons events, 28467 reads/39435293 bases/72362 exons** | | | | | |
| GMAP | -f samse –cross-species -z sense_force | 61.47 | 17.82 | 19.83 | 5.22 |
| Graphmap2 | -x rnaseq | 94.36 | 67.71 | 78.81 | 42.26 |
| Minimap2 | -ax splice | 82.28 | 53.68 | 54.23 | 31.67 |
| Minimap2 + GTF | -ax splice –junc-bed anno.bed | 82.9 | 58.51 | 59.12 | 38.05 |
| deSALT | -d 10 -x ont1d -s 2 -l 14 | 93.55 | 69.03 | 76.24 | 44.05 |
| deSALT + GTF | -d 10 -x ont1d -s 2 -l 14 -G anno.info | 94.16 | 77.71 | 84.96 | 56.03 |
| **Fruit fly ONT 1D, coverage = 4X, 6-9 exons events, 8741 reads/26204640 bases/63690 exons** | | | | | |
| GMAP | -f samse --cross-species -z sense_force | 70.89 | 23.48 | 4.4 | 0.31 |
| Graphmap2 | -x rnaseq | 94.53 | 81.16 | 80.86 | 37.85 |
| Minimap2 | -ax splice | 93.02 | 79.65 | 79.76 | 41.47 |
| Minimap2 + GTF | -ax splice –junc-bed anno.bed | 93.2 | 86.04 | 88.3 | 57.07 |
| deSALT | -d 10 -x ont1d -s 2 -l 14 | 96.63 | 89.4 | 91.99 | 55.15 |
| deSALT + GTF | -d 10 -x ont1d -s 2 -l 14 -G anno.info | 96.97 | 92.84 | 96.68 | 65.08 |
| **Fruit fly ONT 1D, coverage = 4X, > 9 exons events, 7338 reads/36202359 bases/104853 exons** | | | | | |
| GMAP | -f samse --cross-species -z sense_force | 63.83 | 20.25 | 0.26 | 0 |
| Graphmap2 | -x rnaseq | 93.87 | 83.84 | 78.78 | 26.48 |
| Minimap2 | -ax splice | 95.02 | 86.99 | 84.4 | 32.41 |
| Minimap2 + GTF | -ax splice –junc-bed anno.bed | 95.33 | 92.98 | 93.92 | 53.66 |
| deSALT | -d 10 -x ont1d -s 2 -l 14 | 96.94 | 93.1 | 95.2 | 49.33 |
| deSALT + GTF | -d 10 -x ont1d -s 2 -l 14 -G anno.info | 96.65 | 95.52 | 97.45 | 63.48 |
| **Fruit fly ONT 1D, coverage = 10X, < 6 exons events, 59321 reads/93320200 bases/158771 exons** | | | | | |
| GMAP | -f samse --cross-species -z sense_force | 64.41 | 19.4 | 20.71 | 5.38 |
| Graphmap2 | -x rnaseq | 94.81 | 73.84 | 83.82 | 48.79 |
| Minimap2 | -ax splice | 85 | 59.02 | 60.68 | 36.11 |
| Minimap2 + GTF | -ax splice –junc-bed anno.bed | 85.72 | 64.6 | 66.12 | 43.53 |
| deSALT | -d 10 -x ont1d -s 2 -l 14 | 94.62 | 75.88 | 82.23 | 51.75 |
| deSALT + GTF | -d 10 -x ont1d -s 2 -l 14 -G anno.info | 94.98 | 83.74 | 89.86 | 65.15 |

| | | | | | |
|---|---|---|---|---|---|
| **Fruit fly ONT 1D, coverage = 10X, 6-9 exons events, 21936 reads/67655362 bases/160155 exons** | | | | | |
| GMAP | -f samse --cross-species -z sense_force | 71.28 | 23.68 | 4.22 | 0.27 |
| Graphmap2 | -x rnaseq | 94.61 | 83.94 | 84.84 | 43.8 |
| Minimap2 | -ax splice | 93.8 | 81.19 | 81.54 | 42.85 |
| Minimap2 + GTF | -ax splice –junc-bed anno.bed | 93.62 | 87.29 | 89.91 | 58.55 |
| deSALT | -d 10 -x ont1d -s 2 -l 14 | 97.48 | 92.91 | 95.85 | 65.21 |
| deSALT + GTF | -d 10 -x ont1d -s 2 -l 14 -G anno.info | 97.65 | 94.9 | 98.04 | 72.22 |
| **Fruit fly ONT 1D, coverage = 10X, > 9 exons events, 18870 reads/93633101 bases/270690 exons** | | | | | |
| GMAP | -f samse --cross-species -z sense_force | 63.65 | 20.04 | 0.3 | 0.01 |
| Graphmap2 | -x rnaseq | 93.81 | 85.44 | 82.86 | 29.32 |
| Minimap2 | -ax splice | 94.9 | 87.18 | 84.89 | 32.46 |
| Minimap2 + GTF | -ax splice –junc-bed anno.bed | 95.4 | 93.35 | 94.77 | 54.5 |
| deSALT | -d 10 -x ont1d -s 2 -l 14 | 96.96 | 94.65 | 96.99 | 56.02 |
| deSALT + GTF | -d 10 -x ont1d -s 2 -l 14 -G anno.info | 96.9 | 96.48 | 98.12 | 69.49 |
| **Fruit fly ONT 1D, coverage = 30X, < 6 exons events, 175750 reads/278301970 bases/471337 exons** | | | | | |
| GMAP | -f samse --cross-species -z sense_force | 64.44 | 19.42 | 20.76 | 5.3 |
| Graphmap2 | -x rnaseq | 94.87 | 75.2 | 84.85 | 51.02 |
| Minimap2 | -ax splice | 85.31 | 59.25 | 60.82 | 36.3 |
| Minimap2 + GTF | -ax splice –junc-bed anno.bed | 85.81 | 64.82 | 66.36 | 43.68 |
| deSALT | -d 10 -x ont1d -s 2 -l 14 | 95.24 | 80.73 | 87.29 | 59.52 |
| deSALT + GTF | -d 10 -x ont1d -s 2 -l 14 -G anno.info | 95.41 | 85.68 | 91.19 | 68.67 |
| **Fruit fly ONT 1D, coverage = 30X, 6-9 exons events, 65335 reads/202340209 bases/477630 exons** | | | | | |
| GMAP | -f samse --cross-species -z sense_force | 70.93 | 23.65 | 4.21 | 0.32 |
| Graphmap2 | -x rnaseq | 94.72 | 85.39 | 86.57 | 48.21 |
| Minimap2 | -ax splice | 93.5 | 81.02 | 81.35 | 43.2 |
| Minimap2 + GTF | -ax splice –junc-bed anno.bed | 93.69 | 87.39 | 89.91 | 58.95 |
| deSALT | -d 10 -x ont1d -s 2 -l 14 | 97.21 | 94.11 | 96.87 | 70.58 |
| deSALT + GTF | -d 10 -x ont1d -s 2 -l 14 -G anno.info | 97.31 | 95.25 | 98.04 | 74.71 |
| **Fruit fly ONT 1D, coverage = 30X, > 9 exons events, 56736 reads/283160402 bases/815225 exons** | | | | | |
| GMAP | -f samse --cross-species -z sense_force | 64.05 | 20.17 | 0.33 | 0.01 |
| Graphmap2 | -x rnaseq | 93.73 | 86.01 | 83.81 | 31.43 |
| Minimap2 | -ax splice | 94.95 | 87.13 | 85.18 | 32.59 |

| | | | | | |
|---|---|---|---|---|---|
| Minimap2 + GTF | -ax splice –junc-bed anno.bed | 95.42 | 93.31 | 94.7 | 54.51 |
| deSALT | -d 10 -x ont1d -s 2 -l 14 | 97.03 | 94.98 | 97.17 | 58.9 |
| deSALT + GTF | -d 10 -x ont1d -s 2 -l 14 -G anno.info | 96.91 | 96.63 | 97.89 | 71.45 |
| **Mouse ONT 1D, coverage = 4X, < 6 exons events, 33267 reads/33137109 bases/81062 exons** | | | | | |
| GMAP | -f samse --cross-species -z sense_force | 52.82 | 8.51 | 18.01 | 3.96 |
| Graphmap2 | -x rnaseq | 83.57 | 37.82 | 53.66 | 22.25 |
| Minimap2 | -ax splice | 69.84 | 26.5 | 34.42 | 16.32 |
| Minimap2 + GTF | -ax splice –junc-bed anno.bed | 69.98 | 28.21 | 35.84 | 17.57 |
| deSALT | -d 10 -x ont1d -s 2 -l 14 | 83.21 | 43.42 | 53.48 | 25.35 |
| deSALT + GTF | -d 10 -x ont1d -s 2 -l 14 -G anno.info | 84.46 | 54.55 | 63.54 | 36.66 |
| **Mouse ONT 1D, coverage = 4X, 6-9 exons events, 6683 reads/10709673 bases/47920 exons** | | | | | |
| GMAP | -f samse --cross-species -z sense_force | 51.12 | 11.36 | 1.42 | 0.19 |
| Graphmap2 | -x rnaseq | 74.92 | 44.38 | 32.04 | 10.86 |
| Minimap2 | -ax splice | 72.41 | 45.86 | 40.55 | 18.49 |
| Minimap2 + GTF | -ax splice –junc-bed anno.bed | 73.68 | 50.95 | 47.22 | 25.01 |
| deSALT | -d 10 -x ont1d -s 2 -l 14 | 86.7 | 65.57 | 60.27 | 27.97 |
| deSALT + GTF | -d 10 -x ont1d -s 2 -l 14 -G anno.info | 88.18 | 74.28 | 73.87 | 39.29 |
| **Mouse ONT 1D, coverage = 4X, > 9 exons events, 6650 reads/21662084 bases/114907 exons** | | | | | |
| GMAP | -f samse --cross-species -z sense_force | 56.21 | 16.48 | 0.17 | 0 |
| Graphmap2 | -x rnaseq | 75.72 | 56.44 | 34.34 | 7.44 |
| Minimap2 | -ax splice | 85.14 | 73.72 | 61.41 | 17.25 |
| Minimap2 + GTF | -ax splice –junc-bed anno.bed | 87.56 | 82.09 | 75.71 | 31.22 |
| deSALT | -d 10 -x ont1d -s 2 -l 14 | 91.98 | 81.82 | 74.42 | 23.28 |
| deSALT + GTF | -d 10 -x ont1d -s 2 -l 14 -G anno.info | 92.06 | 87.12 | 1.88 | 38.8 |
| **Mouse ONT 1D, coverage = 10X, < 6 exons events, 70045 reads/79985115 bases/180952 exons** | | | | | |
| GMAP | -f samse --cross-species -z sense_force | 54.83 | 9.25 | 17.82 | 3.9 |
| Graphmap2 | -x rnaseq | 84.6 | 42.02 | 56.43 | 24.59 |
| Minimap2 | -ax splice | 72.29 | 29.92 | 38.64 | 18.88 |
| Minimap2 + GTF | -ax splice –junc-bed anno.bed | 72.56 | 31.88 | 40.43 | 20.51 |
| deSALT | -d 10 -x ont1d -s 2 -l 14 | 86.21 | 54.05 | 62.89 | 33.5 |
| deSALT + GTF | -d 10 -x ont1d -s 2 -l 14 -G anno.info | 86.84 | 63.1 | 71.61 | 45.91 |
| **Mouse ONT 1D, coverage = 10X, 6-9 exons events, 16518 reads/27559134 bases/118437 exons** | | | | | |

| | | | | | |
|---|---|---|---|---|---|
| GMAP | -f samse --cross-species -z sense_force | 52.25 | 11.96 | 1.7 | 0.19 |
| Graphmap2 | -x rnaseq | 75.96 | 47.65 | 36.6 | 14.02 |
| Minimap2 | -ax splice | 73.04 | 46.47 | 41.17 | 18.67 |
| Minimap2 + GTF | -ax splice –junc-bed anno.bed | 74.58 | 51.78 | 48.43 | 24.76 |
| deSALT | -d 10 -x ont1d -s 2 -l 14 | 88.78 | 73.11 | 72.29 | 37.87 |
| deSALT + GTF | -d 10 -x ont1d -s 2 -l 14 -G anno.info | 89.49 | 77.96 | 79.19 | 47.14 |
| **Mouse ONT 1D, coverage = 10X, > 9 exons events, 17084 reads/56231420 bases/296347 exons** | | | | | |
| GMAP | -f samse --cross-species -z sense_force | 56.24 | 16.74 | 0.18 | 0 |
| Graphmap2 | -x rnaseq | 75.34 | 58.75 | 39.81 | 9.47 |
| Minimap2 | -ax splice | 85.74 | 74.27 | 61.59 | 17.75 |
| Minimap2 + GTF | -ax splice –junc-bed anno.bed | 87.76 | 82.49 | 75.77 | 32.71 |
| deSALT | -d 10 -x ont1d -s 2 -l 14 | 93.72 | 86.96 | 83.28 | 33.79 |
| deSALT + GTF | -d 10 -x ont1d -s 2 -l 14 -G anno.info | 93.54 | 90.59 | 87.66 | 49.3 |
| **Mouse ONT 1D, coverage = 30X, < 6 exons events, 208841 reads/239854618 bases/540647 exons** | | | | | |
| GMAP | -f samse --cross-species -z sense_force | 39.27 | 6.32 | 14.06 | 2.97 |
| Graphmap2 | -x rnaseq | 84.51 | 42.92 | 57.28 | 25.54 |
| Minimap2 | -ax splice | 72.13 | 29.99 | 38.68 | 19.04 |
| Minimap2 + GTF | -ax splice –junc-bed anno.bed | 72.44 | 31.93 | 40.44 | 20.64 |
| deSALT | -d 10 -x ont1d -s 2 -l 14 | 86.71 | 59.48 | 68.31 | 40.35 |
| deSALT + GTF | -d 10 -x ont1d -s 2 -l 14 -G anno.info | 86.99 | 65.24 | 73.12 | 49.5 |
| **Mouse ONT 1D, coverage = 30X, 6-9 exons events, 49286 reads/82418147 bases/353423 exons** | | | | | |
| GMAP | -f samse --cross-species -z sense_force | 31.22 | 7.16 | 1.07 | 0.09 |
| Graphmap2 | -x rnaseq | 76.05 | 49.08 | 38.91 | 16.27 |
| Minimap2 | -ax splice | 73.56 | 46.85 | 41.52 | 19.05 |
| Minimap2 + GTF | -ax splice –junc-bed anno.bed | 75.01 | 52.24 | 48.74 | 25.69 |
| deSALT | -d 10 -x ont1d -s 2 -l 14 | 89.71 | 76.74 | 77.04 | 45.45 |
| deSALT + GTF | -d 10 -x ont1d -s 2 -l 14 -G anno.info | 89.96 | 78.92 | 79.9 | 50.42 |
| **Mouse ONT 1D, coverage = 30X, > 9 exons events, 51149 reads/169009353 bases/889311 exons** | | | | | |
| GMAP | -f samse --cross-species -z sense_force | 32.11 | 9.64 | 0.1 | 0 |
| Graphmap2 | -x rnaseq | 75.63 | 60.45 | 43.01 | 11.23 |
| Minimap2 | -ax splice | 85.96 | 74.51 | 61.87 | 18.03 |
| Minimap2 + GTF | -ax splice –junc-bed anno.bed | 87.89 | 82.73 | 76.04 | 32.45 |

| | | | | | |
|---|---|---|---|---|---|
| deSALT | -d 10 -x ont1d -s 2 -l 14 | 94.08 | 88.75 | 86.14 | 39.28 |
| deSALT + GTF | -d 10 -x ont1d -s 2 -l 14 -G anno.info | 93.93 | 91.79 | 89.17 | 54.96 |
| **Human ONT 1D, coverage = 4X, < 6 exons events, 34028 reads/28318016 bases/87076 exons** | | | | | |
| GMAP | -f samse --cross-species -z sense_force | 46.46 | 7.29 | 14.23 | 3.39 |
| Graphmap2 | -x rnaseq | 80.16 | 36.16 | 49.08 | 20.28 |
| Minimap2 | -ax splice | 63 | 23.28 | 28.98 | 13.89 |
| Minimap2 + GTF | -ax splice –junc-bed anno.bed | 63.24 | 24.71 | 30.32 | 15.08 |
| deSALT | -d 10 -x ont1d -s 2 -l 14 | 80.4 | 43.58 | 50.62 | 24.75 |
| deSALT + GTF | -d 10 -x ont1d -s 2 -l 14 -G anno.info | 81.84 | 53.06 | 59.82 | 34.9 |
| **Human ONT 1D, coverage = 4X, 6-9 exons events, 6578 reads/9907975 bases/46628 exons** | | | | | |
| GMAP | -f samse --cross-species -z sense_force | 47.59 | 10.02 | 1.32 | 0.21 |
| Graphmap2 | -x rnaseq | 70.8 | 41.19 | 29.31 | 10.78 |
| Minimap2 | -ax splice | 69.67 | 42.53 | 36.8 | 16.3 |
| Minimap2 + GTF | -ax splice –junc-bed anno.bed | 70.62 | 46.72 | 42.7 | 21.9 |
| deSALT | -d 10 -x ont1d -s 2 -l 14 | 86.66 | 66.16 | 61.25 | 28.05 |
| deSALT + GTF | -d 10 -x ont1d -s 2 -l 14 -G anno.info | 88.12 | 74.01 | 73.72 | 37.46 |
| **Human ONT 1D, coverage = 4X, > 9 exons events, 6038 reads/18291208 bases/99917 exons** | | | | | |
| GMAP | -f samse --cross-species -z sense_force | 53.87 | 15.9 | 0.17 | 0 |
| Graphmap2 | -x rnaseq | 71.71 | 52.85 | 33.29 | 7.51 |
| Minimap2 | -ax splice | 83.11 | 70.64 | 58 | 15.98 |
| Minimap2 + GTF | -ax splice –junc-bed anno.bed | 85.92 | 80.05 | 72.62 | 29.57 |
| deSALT | -d 10 -x ont1d -s 2 -l 14 | 91.71 | 81.58 | 74.81 | 24.93 |
| deSALT + GTF | -d 10 -x ont1d -s 2 -l 14 -G anno.info | 91.1 | 85.5 | 79.98 | 37.63 |
| **Human ONT 1D, coverage = 10X, < 6 exons events, 71640 reads/68392014 bases/194695 exons** | | | | | |
| GMAP | -f samse --cross-species -z sense_force | 47.8 | 7.95 | 14.32 | 3.41 |
| Graphmap2 | -x rnaseq | 81.4 | 39.62 | 51.23 | 22.32 |
| Minimap2 | -ax splice | 65.07 | 25.9 | 32.42 | 16.1 |
| Minimap2 + GTF | -ax splice –junc-bed anno.bed | 65.43 | 27.43 | 33.93 | 17.48 |
| deSALT | -d 10 -x ont1d -s 2 -l 14 | 83.7 | 53.08 | 59.85 | 32.28 |
| deSALT + GTF | -d 10 -x ont1d -s 2 -l 14 -G anno.info | 84.49 | 61.44 | 68.11 | 44.11 |
| **Human ONT 1D, coverage = 10X, 6-9 exons events, 16291 reads/25117618 bases/115369 exons** | | | | | |
| GMAP | -f samse --cross-species -z sense_force | 48.9 | 10.7 | 1.5 | 0.12 |

| | | | | | |
|---|---|---|---|---|---|
| Graphmap2 | -x rnaseq | 71.7 | 43.88 | 33.16 | 13.01 |
| Minimap2 | -ax splice | 70.85 | 43.98 | 39.06 | 16.89 |
| Minimap2 + GTF | -ax splice –junc-bed anno.bed | 71.91 | 48.27 | 44.71 | 22.88 |
| deSALT | -d 10 -x ont1d -s 2 -l 14 | 88.52 | 72.73 | 71.59 | 36.63 |
| deSALT + GTF | -d 10 -x ont1d -s 2 -l 14 -G anno.info | 89.13 | 76.61 | 77.32 | 43.42 |
| **Human ONT 1D, coverage = 10X, > 9 exons events, 15601 reads/47786596 bases/259924 exons** | | | | | |
| GMAP | -f samse --cross-species -z sense_force | 53.49 | 15.89 | 0.19 | 0 |
| Graphmap2 | -x rnaseq | 71.41 | 54.29 | 36.06 | 7.96 |
| Minimap2 | -ax splice | 83.86 | 71.47 | 58.92 | 16.7 |
| Minimap2 + GTF | -ax splice –junc-bed anno.bed | 86.34 | 80.53 | 73.65 | 30.32 |
| deSALT | -d 10 -x ont1d -s 2 -l 14 | 93.08 | 85.61 | 81.92 | 32.45 |
| deSALT + GTF | -d 10 -x ont1d -s 2 -l 14 -G anno.info | 92.5 | 88.98 | 85.39 | 45.88 |
| **Human ONT 1D, coverage = 30X, < 6 exons events, 213756 reads/204186862 bases/582406 exons** | | | | | |
| GMAP | -f samse --cross-species -z sense_force | 48.17 | 7.84 | 14.37 | 3.38 |
| Graphmap2 | -x rnaseq | 81.47 | 40.3 | 51.87 | 23.12 |
| Minimap2 | -ax splice | 65.45 | 25.95 | 32.57 | 16.12 |
| Minimap2 + GTF | -ax splice –junc-bed anno.bed | 65.64 | 27.55 | 34.14 | 17.55 |
| deSALT | -d 10 -x ont1d -s 2 -l 14 | 84.79 | 57.84 | 64.66 | 38.58 |
| deSALT + GTF | -d 10 -x ont1d -s 2 -l 14 -G anno.info | 85.13 | 63.28 | 69.44 | 47.31 |
| **Human ONT 1D, coverage = 30X, 6-9 exons events, 48744 reads/75971120 bases/345208 exons** | | | | | |
| GMAP | -f samse --cross-species -z sense_force | 49.09 | 10.68 | 1.51 | 0.15 |
| Graphmap2 | -x rnaseq | 72.08 | 45.09 | 34.49 | 14.17 |
| Minimap2 | -ax splice | 70.63 | 43.96 | 38.9 | 17.23 |
| Minimap2 + GTF | -ax splice –junc-bed anno.bed | 71.94 | 48.27 | 44.73 | 23.2 |
| deSALT | -d 10 -x ont1d -s 2 -l 14 | 88.73 | 75.23 | 75.15 | 41.17 |
| deSALT + GTF | -d 10 -x ont1d -s 2 -l 14 -G anno.info | 88.97 | 77.33 | 78.07 | 45.83 |
| **Human ONT 1D, coverage = 30X, > 9 exons events, 46635 reads/143685596 bases/778725 exons** | | | | | |
| GMAP | -f samse --cross-species -z sense_force | 53.43 | 15.81 | 0.14 | 0 |
| Graphmap2 | -x rnaseq | 71.74 | 55.61 | 38.61 | 9.36 |
| Minimap2 | -ax splice | 83.81 | 71.7 | 58.94 | 16.79 |
| Minimap2 + GTF | -ax splice –junc-bed anno.bed | 86.41 | 80.73 | 73.66 | 30.33 |
| deSALT | -d 10 -x ont1d -s 2 -l 14 | 93.24 | 86.68 | 82.79 | 36.09 |

| | | | | | |
|---|---|---|---|---|---|
| deSALT + GTF | -d 10 -x ont1d -s 2 -l 14 -G anno.info | 92.56 | 89.47 | 85.67 | 48.95 |
| **Human all protein coding genes ONT 1D, < 6 exons events, 577011 reads/863487243 bases/1822487 exons** | | | | | |
| GMAP | -f samse --cross-species -z sense_force | 57.53 | 8.87 | 13.04 | 2.61 |
| Graphmap2 | -x rnaseq | 90.33 | 39.91 | 43.62 | 19.47 |
| Minimap2 | -ax splice | 80.65 | 31.9 | 36.74 | 19.72 |
| Minimap2 + GTF | -ax splice –junc-bed anno.bed | 80.87 | 33.5 | 38.44 | 21.39 |
| deSALT | -d 10 -x ont1d -s 2 -l 14 | 87.99 | 46.93 | 50.66 | 27.78 |
| deSALT + GTF | -d 10 -x ont1d -s 2 -l 14 -G anno.info | 88.03 | 48.35 | 52.14 | 29.72 |
| **Human all protein coding genes ONT 1D, 6-9 exons events, 251586 reads/484778103 bases/1811383 exons** | | | | | |
| GMAP | -f samse --cross-species -z sense_force | 52.82 | 10.62 | 1.35 | 0.13 |
| Graphmap2 | -x rnaseq | 81.95 | 43.41 | 30.27 | 11.3 |
| Minimap2 | -ax splice | 81.01 | 47.46 | 42.1 | 20.15 |
| Minimap2 + GTF | -ax splice –junc-bed anno.bed | 81.95 | 51.71 | 47.86 | 26.59 |
| deSALT | -d 10 -x ont1d -s 2 -l 14 | 91.31 | 62.05 | 57.06 | 26.04 |
| deSALT + GTF | -d 10 -x ont1d -s 2 -l 14 -G anno.info | 91.45 | 65.05 | 61.04 | 33.03 |
| **Human all protein coding genes ONT 1D, > 9 exons events, 297731 reads/1066062435 bases/5250283 exons** | | | | | |
| GMAP | -f samse --cross-species -z sense_force | 51.74 | 12.87 | 0.11 | 0 |
| Graphmap2 | -x rnaseq | 76.1 | 49.01 | 28.92 | 6.52 |
| Minimap2 | -ax splice | 90.25 | 70.4 | 57.18 | 16.29 |
| Minimap2 + GTF | -ax splice –junc-bed anno.bed | 92.83 | 79.59 | 72.25 | 30.15 |
| deSALT | -d 10 -x ont1d -s 2 -l 14 | 95.91 | 76.66 | 65.85 | 18.88 |
| deSALT + GTF | -d 10 -x ont1d -s 2 -l 14 -G anno.info | 96.89 | 83.03 | 75.64 | 34.83 |
| **Simulated datasets by PBSim based on a real ONT dataset (PS-ONT)** | | | | | |
| **Fruit fly PS-ONT, coverage = 4X, < 6 exons events, 31184 reads/ 43804278 bases/ 75312 exons** | | | | | |
| GMAP | -f samse –cross-species -z sense_force | 93.01 | 87.22 | 92.35 | 74.7 |
| Graphmap2 | -x rnaseq | 93.26 | 94.67 | 98.38 | 88.76 |
| Minimap2 | -ax splice | 92.7 | 90.89 | 91.42 | 81.21 |
| Minimap2 + GTF | -ax splice –junc-bed anno.bed | 92.75 | 92.01 | 92.11 | 83.16 |
| deSALT | -d 10 | 93.08 | 94.04 | 95.7 | 87.45 |
| deSALT + GTF | -d 10 -G anno.info | 93.1 | 94.57 | 95.92 | 88.47 |

| | | | | | |
|---|---|---|---|---|---|
| **Fruit fly PS-ONT, coverage = 4X, 6-9 exons events, 8690 reads/ 27208439 bases/ 63241 exons** | | | | | |
| GMAP | -f samse --cross-species -z sense_force | 93.86 | 90.58 | 92.15 | 52.75 |
| Graphmap2 | -x rnaseq | 93.85 | 96.36 | 98.68 | 81.23 |
| Minimap2 | -d 10 | 93.96 | 96.28 | 99.25 | 79.28 |
| Minimap2 + GTF | -d 10 -G anno.info | 94.01 | 97.79 | 99.83 | 85.34 |
| deSALT | -d 10 -x ont1d -s 2 -l 14 | 93.96 | 97.9 | 99.61 | 86.86 |
| deSALT + GTF | -d 10 -x ont1d -s 2 -l 14 -G anno.info | 93.97 | 98.45 | 99.7 | 89.67 |
| **Fruit fly PS-ONT, coverage = 4X, > 9 exons events, 6431 reads/ 30870603 bases/ 89017 exons** | | | | | |
| GMAP | -f samse --cross-species -z sense_force | 93.83 | 90.37 | 90.83 | 30.71 |
| Graphmap2 | -x rnaseq | 93.04 | 96.04 | 97.2 | 71.42 |
| Minimap2 | -ax splice | 94.03 | 97.01 | 99.38 | 72.46 |
| Minimap2 + GTF | -ax splice –junc-bed anno.bed | 94.07 | 98.48 | 99.86 | 81.9 |
| deSALT | -d 10 | 94.09 | 98.4 | 99.64 | 82.82 |
| deSALT + GTF | -d 10 -G anno.info | 94.1 | 98.9 | 99.84 | 86.88 |
| **Fruit fly PS-ONT, coverage = 10X, < 6 exons events, 66904 reads/ 107361007 bases/ 172634 exons** | | | | | |
| GMAP | -f samse --cross-species -z sense_force | 93.22 | 88.56 | 93.7 | 75.91 |
| Graphmap2 | -x rnaseq | 93.38 | 95.41 | 98.73 | 89.59 |
| Minimap2 | -ax splice | 93.2 | 94.05 | 97.2 | 86.56 |
| Minimap2 + GTF | -ax splice –junc-bed anno.bed | 93.24 | 95.09 | 97.87 | 88.52 |
| deSALT | -d 10 | 93.25 | 96.09 | 98.12 | 91.11 |
| deSALT + GTF | -d 10 -G anno.info | 93.25 | 96.26 | 98.23 | 91.45 |
| **Fruit fly PS-ONT, coverage = 10X, 6-9 exons events, 21456 reads/ 67508122 bases/ 156152 exons** | | | | | |
| GMAP | -f samse --cross-species -z sense_force | 93.85 | 90.71 | 92.18 | 53.18 |
| Graphmap2 | -x rnaseq | 93.6 | 96.53 | 98.73 | 81.81 |
| Minimap2 | -ax splice | 93.97 | 96.36 | 99.18 | 79.67 |
| Minimap2 + GTF | -ax splice –junc-bed anno.bed | 94.01 | 97.86 | 99.86 | 85.66 |
| deSALT | -d 10 | 93.93 | 98.18 | 99.57 | 88.4 |
| deSALT + GTF | -d 10 -G anno.info | 93.94 | 98.52 | 99.69 | 90.12 |
| **Fruit fly PS-ONT, coverage = 10X, > 9 exons events, 16415 reads/ 79741214 bases/ 228471 exons** | | | | | |
| GMAP | -f samse --cross-species -z sense_force | 93.76 | 90.25 | 90.3 | 30.51 |
| Graphmap2 | -x rnaseq | 92.83 | 95.86 | 96.72 | 71.65 |
| Minimap2 | -ax splice | 94.06 | 97.05 | 99.29 | 72.68 |

| | | | | | |
|---|---|---|---|---|---|
| Minimap2 + GTF | -ax splice –junc-bed anno.bed | 94.1 | 98.52 | 99.82 | 82.49 |
| deSALT | -d 10 | 94.14 | 98.68 | 99.76 | 85.22 |
| deSALT + GTF | -d 10 -G anno.info | 94.14 | 98.87 | 99.81 | 87.71 |
| **Fruit fly PS-ONT, coverage = 30X, < 6 exons events, 190138 reads/ 315781018 bases/ 498304 exons** | | | | | |
| GMAP | -f samse --cross-species -z sense_force | 93.3 | 88.87 | 93.9 | 76.13 |
| Graphmap2 | -x rnaseq | 93.37 | 95.74 | 98.81 | 90.18 |
| Minimap2 | -ax splice | 93.26 | 94.33 | 97.31 | 86.9 |
| Minimap2 + GTF | -ax splice –junc-bed anno.bed | 93.31 | 95.41 | 97.99 | 88.97 |
| deSALT | -d 10 | 93.33 | 96.56 | 98.22 | 92.04 |
| deSALT + GTF | -d 10 -G anno.info | 93.33 | 96.65 | 98.32 | 92.18 |
| **Fruit fly PS-ONT, coverage = 30X, 6-9 exons events, 64714 reads/ 206432701 bases/ 471269 exons** | | | | | |
| GMAP | -f samse --cross-species -z sense_force | 93.88 | 90.69 | 91.99 | 54.12 |
| Graphmap2 | -x rnaseq | 93.61 | 96.65 | 98.72 | 82.62 |
| Minimap2 | -ax splice | 93.99 | 96.4 | 99.17 | 80.07 |
| Minimap2 + GTF | -ax splice –junc-bed anno.bed | 94.03 | 97.91 | 99.87 | 86.04 |
| deSALT | -d 10 | 93.94 | 98.31 | 99.56 | 89.26 |
| deSALT + GTF | -d 10 -G anno.info | 93.94 | 98.56 | 99.65 | 90.42 |
| **Fruit fly PS-ONT, coverage = 30X, > 9 exons events, 49658 reads/ 241693941 bases/ 691327 exons** | | | | | |
| GMAP | -f samse --cross-species -z sense_force | 93.78 | 90.32 | 90.66 | 30.68 |
| Graphmap2 | -x rnaseq | 92.82 | 95.9 | 96.74 | 72.18 |
| Minimap2 | -ax splice | 94.05 | 97.07 | 99.33 | 72.97 |
| Minimap2 + GTF | -ax splice –junc-bed anno.bed | 94.09 | 98.52 | 99.78 | 82.5 |
| deSALT | -d 10 | 94.13 | 98.7 | 99.75 | 85.58 |
| deSALT + GTF | -d 10 -G anno.info | 94.13 | 98.98 | 99.8 | 88.01 |
| **Mouse PS-ONT, coverage = 4X, < 6 exons events, 34821 reads/ 34453561 bases/ 79935 exons** | | | | | |
| GMAP | -f samse --cross-species -z sense_force | 89.66 | 78.97 | 85.1 | 66.83 |
| Graphmap2 | -x rnaseq | 90.14 | 86.01 | 92.67 | 77.64 |
| Minimap2 | -ax splice | 90.34 | 82.14 | 83.11 | 69.93 |
| Minimap2 + GTF | -ax splice –junc-bed anno.bed | 90.38 | 82.64 | 83.54 | 70.48 |
| deSALT | -d 10 | 90.42 | 87.3 | 89.59 | 78.42 |
| deSALT + GTF | -d 10 -G anno.info | 90.49 | 87.78 | 89.82 | 79.21 |
| **Mouse PS-ONT, coverage = 4X, 6-9 exons events, 6409 reads/ 11010775 bases/ 45986 exons** | | | | | |

| | | | | | |
|---|---|---|---|---|---|
| GMAP | -f samse --cross-species -z sense_force | 90.56 | 84.2 | 81.85 | 43.45 |
| Graphmap2 | -x rnaseq | 88.51 | 87.62 | 89.3 | 61.94 |
| Minimap2 | -ax splice | 92.36 | 91.97 | 95.49 | 65.84 |
| Minimap2 + GTF | -ax splice –junc-bed anno.bed | 92.46 | 93.44 | 97.29 | 69.53 |
| deSALT | -d 10 | 92.41 | 93.98 | 96.61 | 76.38 |
| deSALT + GTF | -d 10 -G anno.info | 92.53 | 94.92 | 97.24 | 79.47 |
| **Mouse PS-ONT, coverage = 4X, > 9 exons events, 6118 reads/ 20155054 bases/ 103104 exons** | | | | | |
| GMAP | -f samse --cross-species -z sense_force | 92.48 | 89.13 | 85.34 | 24.26 |
| Graphmap2 | -x rnaseq | 85.98 | 87.37 | 86.73 | 51.68 |
| Minimap2 | -ax splice | 93.28 | 96.6 | 98.28 | 66.82 |
| Minimap2 + GTF | -ax splice –junc-bed anno.bed | 93.34 | 97.58 | 99.12 | 72.44 |
| deSALT | -d 10 | 93.62 | 97.99 | 98.81 | 79.23 |
| deSALT + GTF | -d 10 -G anno.info | 93.73 | 98.48 | 99.2 | 83.03 |
| **Mouse PS-ONT, coverage = 10X, < 6 exons events, 70045 reads/79985115 bases/180952 exons** | | | | | |
| GMAP | -f samse --cross-species -z sense_force | 90.06 | 80.76 | 86.92 | 67.87 |
| Graphmap2 | -x rnaseq | 90.42 | 87.44 | 93.63 | 79.16 |
| Minimap2 | -ax splice | 91.39 | 87.8 | 92.99 | 77.9 |
| Minimap2 + GTF | -ax splice –junc-bed anno.bed | 91.43 | 88.35 | 93.46 | 78.59 |
| deSALT | -d 10 | 91.07 | 91.11 | 94.28 | 84.16 |
| deSALT + GTF | -d 10 -G anno.info | 91.1 | 91.4 | 94.44 | 84.62 |
| **Mouse PS-ONT, coverage = 10X, 6-9 exons events, 16518 reads/27559134 bases/118437 exons** | | | | | |
| GMAP | -f samse --cross-species -z sense_force | 90.59 | 84.38 | 82.73 | 43.96 |
| Graphmap2 | -x rnaseq | 88.81 | 88.15 | 89.92 | 63.12 |
| Minimap2 | -ax splice | 92.39 | 92.08 | 95.77 | 66.11 |
| Minimap2 + GTF | -ax splice –junc-bed anno.bed | 92.48 | 93.42 | 97.31 | 69.55 |
| deSALT | -d 10 | 92.57 | 94.57 | 96.82 | 78.6 |
| deSALT + GTF | -d 10 -G anno.info | 92.56 | 95.07 | 97.09 | 80.6 |
| **Mouse PS-ONT, coverage = 10X, > 9 exons events, 17084 reads/56231420 bases/296347 exons** | | | | | |
| GMAP | -f samse --cross-species -z sense_force | 92.33 | 88.83 | 85.21 | 22.98 |
| Graphmap2 | -x rnaseq | 86.05 | 87.39 | 86.69 | 52.38 |
| Minimap2 | -ax splice | 93.93 | 96.69 | 98.25 | 67.69 |
| Minimap2 + GTF | -ax splice –junc-bed anno.bed | 93.45 | 97.67 | 99.17 | 73.22 |

| | | | | | |
|---|---|---|---|---|---|
| deSALT | -d 10 | 93.78 | 98.29 | 99.05 | 82.19 |
| deSALT + GTF | -d 10 -G anno.info | 93.79 | 98.56 | 99.3 | 84.23 |
| **Mouse PS-ONT, coverage = 30X, < 6 exons events, 211498 reads/ 251585843 bases/ 530445 exons** | | | | | |
| GMAP | -f samse --cross-species -z sense_force | 90.18 | 81.29 | 87.16 | 68.08 |
| Graphmap2 | -x rnaseq | 90.46 | 87.88 | 93.73 | 79.58 |
| Minimap2 | -ax splice | 91.52 | 88.5 | 93.63 | 78.39 |
| Minimap2 + GTF | -ax splice –junc-bed anno.bed | 91.56 | 89.06 | 94.14 | 79.1 |
| deSALT | -d 10 | 91.11 | 91.7 | 94.49 | 84.97 |
| deSALT + GTF | -d 10 -G anno.info | 91.14 | 91.93 | 94.66 | 85.31 |
| **Mouse PS-ONT, coverage = 30X, 6-9 exons events, 47544 reads/ 82922348 bases/ 341162 exons** | | | | | |
| GMAP | -f samse --cross-species -z sense_force | 90.65 | 84.51 | 82.83 | 43.91 |
| Graphmap2 | -x rnaseq | 88.66 | 88.2 | 90.01 | 63.72 |
| Minimap2 | -ax splice | 92.44 | 92.22 | 95.75 | 66.7 |
| Minimap2 + GTF | -ax splice –junc-bed anno.bed | 92.53 | 93.62 | 97.32 | 70.28 |
| deSALT | -d 10 | 92.55 | 94.77 | 96.8 | 79.56 |
| deSALT + GTF | -d 10 -G anno.info | 92.56 | 95.23 | 97.11 | 81.11 |
| **Mouse PS-ONT, coverage = 30X, > 9 exons events, 46902 reads/ 156993902 bases/ 794133 exons** | | | | | |
| GMAP | -f samse --cross-species -z sense_force | 92.41 | 88.9 | 84.99 | 23.22 |
| Graphmap2 | -x rnaseq | 86.05 | 87.53 | 86.72 | 53.27 |
| Minimap2 | -ax splice | 93.37 | 96.73 | 93.82 | 67.93 |
| Minimap2 + GTF | -ax splice –junc-bed anno.bed | 93.44 | 97.71 | 99.14 | 73.52 |
| deSALT | -d 10 | 93.79 | 98.45 | 99.15 | 83.24 |
| deSALT + GTF | -d 10 -G anno.info | 93.79 | 98.63 | 99.28 | 84.92 |
| **Human PS-ONT, coverage = 4X, < 6 exons events, 35436 reads/ 29563500 bases/ 85612 exons** | | | | | |
| GMAP | -f samse --cross-species -z sense_force | 90.22 | 79.67 | 84.58 | 65.66 |
| Graphmap2 | -x rnaseq | 90.78 | 86.25 | 92.01 | 76.4 |
| Minimap2 | -ax splice | 90.36 | 81.68 | 81.35 | 67.47 |
| Minimap2 + GTF | -ax splice –junc-bed anno.bed | 90.41 | 82.24 | 81.86 | 68.17 |
| deSALT | -d 10 | 91.5 | 88.07 | 89.94 | 77.9 |
| deSALT + GTF | -d 10 -G anno.info | 91.63 | 88.71 | 90.31 | 78.93 |
| **Human PS-ONT, coverage = 4X, 6-9 exons events, 6125 reads/ 9777146 bases/ 43495 exons** | | | | | |
| GMAP | -f samse --cross-species -z sense_force | 91.02 | 84.74 | 82.48 | 42.01 |

| | | | | | |
|---|---|---|---|---|---|
| Graphmap2 | -x rnaseq | 87.95 | 87.59 | 88.82 | 61.04 |
| Minimap2 | -ax splice | 92.44 | 92.76 | 96.52 | 66.33 |
| Minimap2 + GTF | -ax splice –junc-bed anno.bed | 92.52 | 93.67 | 97.37 | 68.98 |
| deSALT | -d 10 | 92.94 | 95.22 | 98.35 | 76.36 |
| deSALT + GTF | -d 10 -G anno.info | 93.05 | 95.8 | 98.55 | 78.91 |
| **Human PS-ONT, coverage = 4X, > 9 exons events, 5625 reads/ 17309153 bases/ 91196 exons** | | | | | |
| GMAP | -f samse --cross-species -z sense_force | 91.78 | 88.2 | 83.79 | 22.06 |
| Graphmap2 | -x rnaseq | 85.76 | 86.79 | 86.29 | 49.07 |
| Minimap2 | -ax splice | 93.42 | 96.66 | 98.95 | 66.01 |
| Minimap2 + GTF | -ax splice –junc-bed anno.bed | 93.47 | 97.45 | 99.34 | 70.92 |
| deSALT | -d 10 | 93.53 | 97.67 | 98.97 | 78.56 |
| deSALT + GTF | -d 10 -G anno.info | 93.61 | 98.11 | 99.06 | 81.9 |
| **Human PS-ONT, coverage = 10X, < 6 exons events, 75517 reads/ 72370076 bases/ 196438 exons** | | | | | |
| GMAP | -f samse --cross-species -z sense_force | 90.64 | 81.44 | 96.17 | 67.22 |
| Graphmap2 | -x rnaseq | 91.12 | 88.12 | 93.44 | 78.27 |
| Minimap2 | -ax splice | 91.68 | 88.03 | 92.6 | 76.57 |
| Minimap2 + GTF | -ax splice –junc-bed anno.bed | 91.73 | 88.58 | 93.15 | 77.31 |
| deSALT | -d 10 | 92.33 | 92.5 | 95.67 | 84.99 |
| deSALT + GTF | -d 10 -G anno.info | 92.37 | 92.85 | 95.94 | 85.61 |
| **Human PS-ONT, coverage = 10X, 6-9 exons events, 15422 reads/ 24935645 bases/ 109504 exons** | | | | | |
| GMAP | -f samse --cross-species -z sense_force | 91.07 | 84.91 | 82.48 | 42.34 |
| Graphmap2 | -x rnaseq | 87.96 | 88.06 | 89.52 | 61.42 |
| Minimap2 | -ax splice | 92.5 | 92.85 | 96.86 | 65.87 |
| Minimap2 + GTF | -ax splice –junc-bed anno.bed | 92.58 | 93.77 | 97.63 | 68.64 |
| deSALT | -d 10 | 93.03 | 95.61 | 98.46 | 78.35 |
| deSALT + GTF | -d 10 -G anno.info | 93.14 | 96.08 | 98.66 | 80.09 |
| **Human PS-ONT, coverage = 10X, > 9 exons events, 14104 reads/ 43989123 bases/ 229617 exons** | | | | | |
| GMAP | -f samse --cross-species -z sense_force | 91.9 | 88.19 | 83.94 | 22.12 |
| Graphmap2 | -x rnaseq | 85.8 | 86.96 | 86.41 | 49.98 |
| Minimap2 | -ax splice | 93.41 | 96.68 | 98.8 | 67.87 |
| Minimap2 + GTF | -ax splice –junc-bed anno.bed | 93.48 | 97.5 | 99.26 | 72.49 |
| deSALT | -d 10 | 93.55 | 97.9 | 99.01 | 80.57 |

| | | | | | |
|---|---|---|---|---|---|
| deSALT + GTF | -d 10 -G anno.info | 93.59 | 98.08 | 98.99 | 82.49 |
| **Human PS-ONT, coverage = 30X, < 6 exons events, 215157 reads/ 215509093 bases/ /570699 exons** | | | | | |
| GMAP | -f samse --cross-species -z sense_force | 90.91 | 82.17 | 86.64 | 67.63 |
| Graphmap2 | -x rnaseq | 91.17 | 88.61 | 93.5 | 79.06 |
| Minimap2 | -ax splice | 91.93 | 88.85 | 93.38 | 77.4 |
| Minimap2 + GTF | -ax splice –junc-bed anno.bed | 91.97 | 89.47 | 94.01 | 78.24 |
| deSALT | -d 10 | 92.51 | 93.25 | 96.12 | 86.08 |
| deSALT + GTF | -d 10 -G anno.info | 92.55 | 93.58 | 96.38 | 86.65 |
| **Human PS-ONT, coverage = 30X, 6-9 exons events, 46308 reads/ 75023713 bases/ 328702 exons** | | | | | |
| GMAP | -f samse --cross-species -z sense_force | 91.09 | 85.09 | 82.64 | 43.32 |
| Graphmap2 | -x rnaseq | 88.08 | 88.22 | 89.45 | 62.17 |
| Minimap2 | -ax splice | 92.49 | 92.95 | 96.65 | 66.9 |
| Minimap2 + GTF | -ax splice –junc-bed anno.bed | 92.58 | 93.86 | 97.44 | 69.51 |
| deSALT | -d 10 | 93.1 | 95.79 | 98.46 | 79.12 |
| deSALT + GTF | -d 10 -G anno.info | 93.17 | 96.2 | 98.68 | 80.66 |
| **Human PS-ONT, coverage = 30X, > 9 exons events, 42668 reads/ 133560785 bases/ 695553 exons** | | | | | |
| GMAP | -f samse --cross-species -z sense_force | 91.86 | 88.2 | 83.83 | 22.06 |
| Graphmap2 | -x rnaseq | 85.68 | 86.9 | 86.25 | 49.82 |
| Minimap2 | -ax splice | 93.43 | 96.71 | 98.84 | 68.07 |
| Minimap2 + GTF | -ax splice –junc-bed anno.bed | 93.49 | 97.53 | 99.29 | 75.29 |
| deSALT | -d 10 | 93.58 | 97.88 | 98.91 | 80.49 |
| deSALT + GTF | -d 10 -G anno.info | 93.63 | 98.15 | 99.02 | 82.46 |
| **Human all protein coding genes PS-ONT, < 6 exons events, 797474 reads/1055126496 bases/2269595 exons** | | | | | |
| GMAP | -f samse --cross-species -z sense_force | 93.46 | 77.24 | 71.18 | 39.05 |
| Graphmap2 | -x rnaseq | 93.69 | 83.46 | 88.99 | 68.14 |
| Minimap2 | -ax splice | 94.08 | 84.02 | 85.27 | 67.81 |
| Minimap2 + GTF | -ax splice –junc-bed anno.bed | 94.28 | 85.09 | 86.28 | 68.98 |
| deSALT | -d 10 | 95.04 | 90.04 | 92.58 | 78.18 |
| deSALT + GTF | -d 10 -G anno.info | 95.05 | 90.12 | 92.55 | 78.45 |
| **Human all protein coding genes PS-ONT, 6-9 exons events, 269700 reads/516491645 bases/1938060 exons** | | | | | |
| GMAP | -f samse --cross-species -z sense_force | 93.32 | 77.29 | 69.44 | 30.51 |
| Graphmap2 | -x rnaseq | 91.04 | 84.84 | 86.24 | 52.34 |

| | | | | | |
|---|---|---|---|---|---|
| Minimap2 | -ax splice | 94.44 | 91.95 | 95.32 | 67.81 |
| Minimap2 + GTF | -ax splice –junc-bed anno.bed | 95.01 | 93.95 | 97.35 | 71.92 |
| deSALT | -d 10 | 95.56 | 95.79 | 98.2 | 79.54 |
| deSALT + GTF | -d 10 -G anno.info | 95.58 | 96.3 | 98.35 | 81.91 |
| **Human all protein coding genes PS-ONT, > 9 exons events, 299907 reads/1053041154 bases/5102557 exons** | | | | | |
| GMAP | -f samse --cross-species -z sense_force | 93.52 | 78.48 | 58.92 | 10.95 |
| Graphmap2 | -x rnaseq | 86.58 | 80.97 | 78.77 | 32.96 |
| Minimap2 | -ax splice | 95.23 | 95.87 | 97.72 | 66.79 |
| Minimap2 + GTF | -ax splice –junc-bed anno.bed | 95.52 | 97.23 | 98.75 | 73.38 |
| deSALT | -d 10 | 95.89 | 97.84 | 98.93 | 79.23 |
| deSALT + GTF | -d 10 -G anno.info | 95.87 | 98.18 | 98.84 | 83.34 |
| **Simulated datasets by NanoSim based on a real ONT dataset (NS-ONT)** | | | | | |
| **Fruit fly NS-ONT, coverage = 4X, < 6 exons events, 13069 reads/ 62848044 bases/ 36819 exons** | | | | | |
| GMAP | -f samse –cross-species -z sense_force | 91.65 | 59.37 | 22.98 | 10.75 |
| Graphmap2 | -x rnaseq | 92.15 | 73.78 | 43.07 | 23.35 |
| Minimap2 | -ax splice | 90.59 | 89.36 | 92.78 | 75.23 |
| Minimap2 + GTF | -ax splice –junc-bed anno.bed | 90.6 | 92.04 | 94.8 | 78.29 |
| deSALT | -d 10 | 89.95 | 90.52 | 94.88 | 77.85 |
| deSALT + GTF | -d 10 -G anno.info | 89.95 | 91.87 | 95.39 | 81.04 |
| **Fruit fly NS-ONT, coverage = 4X, 6-9 exons events, 8091 reads/ 45696620 bases/ 60810 exons** | | | | | |
| GMAP | -f samse --cross-species -z sense_force | 92.53 | 64.86 | 16.82 | 2.37 |
| Graphmap2 | -x rnaseq | 93.22 | 83.74 | 56.58 | 16.52 |
| Minimap2 | -ax splice | 93.47 | 93.48 | 97.21 | 67.19 |
| Minimap2 + GTF | -ax splice –junc-bed anno.bed | 93.52 | 97.29 | 99.16 | 84.97 |
| deSALT | -d 10 | 93.23 | 95.6 | 98.46 | 75.91 |
| deSALT + GTF | -d 10 -G anno.info | 93.25 | 97.54 | 98.81 | 87.53 |
| **Fruit fly NS-ONT, coverage = 4X, > 9 exons events, 12843 reads/ 89565943 bases/ 196131 exons** | | | | | |
| GMAP | -f samse --cross-species -z sense_force | 92.43 | 64.67 | 9.43 | 0.31 |
| Graphmap2 | -x rnaseq | 92.41 | 84.19 | 61.33 | 9.18 |
| Minimap2 | -ax splice | 94.33 | 94.36 | 97.92 | 50.09 |

| | | | | | |
|---|---|---|---|---|---|
| Minimap2 + GTF | -ax splice –junc-bed anno.bed | 94.39 | 98.07 | 99.66 | 78.36 |
| deSALT | -d 10 | 94.33 | 96.49 | 99.33 | 62.48 |
| deSALT + GTF | -d 10 -G anno.info | 94.35 | 98.38 | 99.62 | 82.15 |
| **Fruit fly NS-ONT, coverage = 10X, < 6 exons events, 33072 reads/ 159299063 bases/ 93644 exons** | | | | | |
| GMAP | -f samse --cross-species -z sense_force | 91.54 | 59.21 | 22.62 | 10.37 |
| Graphmap2 | -x rnaseq | 92.05 | 74.07 | 43.51 | 23.38 |
| Minimap2 | -ax splice | 90.39 | 89.42 | 92.85 | 73.21 |
| Minimap2 + GTF | -ax splice –junc-bed anno.bed | 90.41 | 92.02 | 94.79 | 78.18 |
| deSALT | -d 10 | 89.82 | 90.53 | 94.62 | 77.91 |
| deSALT + GTF | -d 10 -G anno.info | 89.82 | 91.85 | 95.05 | 81.1 |
| **Fruit fly NS-ONT, coverage = 10X, 6-9 exons events, 20277 reads/ 115757519 bases/ 152436 exons** | | | | | |
| GMAP | -f samse --cross-species -z sense_force | 92.37 | 65.05 | 16.59 | 2.14 |
| Graphmap2 | -x rnaseq | 92.84 | 83.48 | 55.74 | 16.16 |
| Minimap2 | -ax splice | 93.3 | 93.8 | 97.41 | 68.75 |
| Minimap2 + GTF | -ax splice –junc-bed anno.bed | 93.35 | 97.47 | 99.34 | 85.49 |
| deSALT | -d 10 | 93.13 | 96.01 | 98.67 | 77.81 |
| deSALT + GTF | -d 10 -G anno.info | 93.15 | 97.72 | 98.99 | 88.22 |
| **Fruit fly NS-ONT, coverage = 10X, > 9 exons events, 32654 reads/ 228509902 bases/ 498661 exons** | | | | | |
| GMAP | -f samse --cross-species -z sense_force | 92.4 | 64.85 | 9.44 | 0.33 |
| Graphmap2 | -x rnaseq | 92.2 | 82.85 | 58.8 | 8.27 |
| Minimap2 | -ax splice | 94.32 | 94.24 | 97.8 | 49.64 |
| Minimap2 + GTF | -ax splice –junc-bed anno.bed | 94.38 | 98.01 | 99.57 | 78.49 |
| deSALT | -d 10 | 94.34 | 96.4 | 99.34 | 62.57 |
| deSALT + GTF | -d 10 -G anno.info | 94.36 | 98.32 | 99.54 | 82.47 |
| **Fruit fly NS-ONT, coverage = 30X, < 6 exons events, 96378 reads/ 463284083 bases/ 272634 exons** | | | | | |
| GMAP | -f samse --cross-species -z sense_force | 91.45 | 59.24 | 22.61 | 10.57 |
| Graphmap2 | -x rnaseq | 92.01 | 73.62 | 42.93 | 23.4 |
| Minimap2 | -ax splice | 90.32 | 89.43 | 92.84 | 73.08 |
| Minimap2 + GTF | -ax splice –junc-bed anno.bed | 90.34 | 92.02 | 94.79 | 78.05 |
| deSALT | -d 10 | 89.87 | 90.62 | 94.71 | 77.98 |
| deSALT + GTF | -d 10 -G anno.info | 89.88 | 91.95 | 95.17 | 81.15 |
| **Fruit fly NS-ONT, coverage = 30X, 6-9 exons events, 58796 reads/ 333941145 bases/ 441383 exons** | | | | | |

| | | | | | |
|---|---|---|---|---|---|
| GMAP | -f samse --cross-species -z sense_force | 92.5 | 64.91 | 16.94 | 2.27 |
| Graphmap2 | -x rnaseq | 93.08 | 82.51 | 54.69 | 45.22 |
| Minimap2 | -ax splice | 93.44 | 93.59 | 97.13 | 68.02 |
| Minimap2 + GTF | -ax splice –junc-bed anno.bed | 93.48 | 97.35 | 99.22 | 85.29 |
| deSALT | -d 10 | 93.22 | 95.93 | 98.57 | 77.6 |
| deSALT + GTF | -d 10 -G anno.info | 93.23 | 97.67 | 98.9 | 88.28 |
| **Fruit fly NS-ONT, coverage = 30X, > 9 exons events, 94829 reads/ 663118680 bases/ 1444979 exons** | | | | | |
| GMAP | -f samse --cross-species -z sense_force | 92.5 | 64.79 | 9.43 | 0.31 |
| Graphmap2 | -x rnaseq | 92.4 | 81.21 | 54.52 | 7.16 |
| Minimap2 | -ax splice | 94.38 | 94.22 | 97.85 | 49.81 |
| Minimap2 + GTF | -ax splice –junc-bed anno.bed | 94.44 | 98.02 | 99.65 | 78.3 |
| deSALT | -d 10 | 94.39 | 96.37 | 99.37 | 61.98 |
| deSALT + GTF | -d 10 -G anno.info | 94.41 | 98.33 | 99.59 | 81.92 |
| **Mouse NS-ONT, coverage = 4X, < 6 exons events, 15795 reads/ 89819086 bases/ 24809 exons** | | | | | |
| GMAP | -f samse --cross-species -z sense_force | 92.76 | 35.87 | 15.99 | 9.22 |
| Graphmap2 | -x rnaseq | 93.38 | 45.1 | 27.71 | 18.52 |
| Minimap2 | -ax splice | 93.73 | 83.2 | 96.97 | 76.48 |
| Minimap2 + GTF | -ax splice –junc-bed anno.bed | 93.74 | 84.29 | 97.41 | 77.62 |
| deSALT | -d 10 | 93.73 | 83.85 | 96.61 | 77.09 |
| deSALT + GTF | -d 10 -G anno.info | 93.73 | 84.59 | 96.82 | 78.01 |
| **Mouse NS-ONT, coverage = 4X, 6-9 exons events, 3257 reads/ 15322105 bases/ 25157 exons** | | | | | |
| GMAP | -f samse --cross-species -z sense_force | 91.19 | 67.23 | 22.2 | 2.98 |
| Graphmap2 | -x rnaseq | 88.58 | 79.01 | 59.1 | 15.66 |
| Minimap2 | -ax splice | 92.88 | 91.5 | 95.18 | 61.74 |
| Minimap2 + GTF | -ax splice –junc-bed anno.bed | 92.94 | 95.09 | 97.94 | 76.45 |
| deSALT | -d 10 | 92.98 | 94.08 | 96.93 | 72.18 |
| deSALT + GTF | -d 10 -G anno.info | 93.01 | 96.04 | 97.85 | 82.41 |
| **Mouse NS-ONT, coverage = 4X, > 9 exons events, 17951 reads/ 111099250 bases/ 516001 exons** | | | | | |
| GMAP | -f samse --cross-species -z sense_force | 89.66 | 66.88 | 8.94 | 0.18 |
| Graphmap2 | -x rnaseq | 85.42 | 79.16 | 66.09 | 5.88 |
| Minimap2 | -ax splice | 93.1 | 93.22 | 97.18 | 28.37 |
| Minimap2 + GTF | -ax splice –junc-bed anno.bed | 93.26 | 96.97 | 99.06 | 55.94 |

| | | | | | |
|---|---|---|---|---|---|
| deSALT | -d 10 | 93.89 | 96.1 | 98.96 | 42.44 |
| deSALT + GTF | -d 10 -G anno.info | 93.94 | 98.14 | 99.37 | 66.13 |
| **Mouse NS-ONT, coverage = 10X, < 6 exons events, 38543 reads/218869207 bases/59554 exons** | | | | | |
| GMAP | -f samse --cross-species -z sense_force | 92.6 | 34.6 | 15.64 | 8.86 |
| Graphmap2 | -x rnaseq | 93.27 | 44.41 | 27.68 | 18.63 |
| Minimap2 | -ax splice | 93.66 | 83.19 | 96.96 | 76.76 |
| Minimap2 + GTF | -ax splice –junc-bed anno.bed | 93.66 | 84.19 | 97.34 | 77.75 |
| deSALT | -d 10 | 93.68 | 84.06 | 96.72 | 77.41 |
| deSALT + GTF | -d 10 -G anno.info | 93.68 | 84.77 | 96.94 | 78.24 |
| **Mouse NS-ONT, coverage = 10X, 6-9 exons events, 8020 reads/38227570 bases/61548 exons** | | | | | |
| GMAP | -f samse --cross-species -z sense_force | 91.1 | 67.06 | 20.27 | 2.94 |
| Graphmap2 | -x rnaseq | 88.58 | 78.23 | 57.62 | 15.34 |
| Minimap2 | -ax splice | 92.89 | 91.74 | 95.05 | 62.16 |
| Minimap2 + GTF | -ax splice –junc-bed anno.bed | 92.94 | 95.21 | 97.86 | 76.5 |
| deSALT | -d 10 | 93.03 | 94.63 | 97.81 | 72.79 |
| deSALT + GTF | -d 10 -G anno.info | 93.05 | 96.45 | 98.48 | 82.89 |
| **Mouse NS-ONT, coverage = 10X, > 9 exons events, 43440 reads/269003477 bases/1253355 exons** | | | | | |
| GMAP | -f samse --cross-species -z sense_force | 89.71 | 66.85 | 8.94 | 0.16 |
| Graphmap2 | -x rnaseq | 85.28 | 78.62 | 63.91 | 5.62 |
| Minimap2 | -ax splice | 93.24 | 93.26 | 97.21 | 28.21 |
| Minimap2 + GTF | -ax splice –junc-bed anno.bed | 93.4 | 97.02 | 99.05 | 55.48 |
| deSALT | -d 10 | 94.02 | 96.1 | 98.94 | 42.15 |
| deSALT + GTF | -d 10 -G anno.info | 94.07 | 98.05 | 99.36 | 66.27 |
| **Mouse NS-ONT, coverage = 30X, < 6 exons events, 115562 reads/ 653512033 bases/ 180119 exons** | | | | | |
| GMAP | -f samse --cross-species -z sense_force | 92.74 | 35.23 | 15.69 | 9.14 |
| Graphmap2 | -x rnaseq | 93.36 | 44.74 | 27.93 | 18.64 |
| Minimap2 | -ax splice | 93.74 | 83.02 | 96.83 | 76.31 |
| Minimap2 + GTF | -ax splice –junc-bed anno.bed | 93.74 | 84.08 | 97.26 | 77.45 |
| deSALT | -d 10 | 93.74 | 84.01 | 96.61 | 77.24 |
| deSALT + GTF | -d 10 -G anno.info | 93.74 | 84.79 | 96.84 | 78.19 |
| **Mouse NS-ONT, coverage = 30X, 6-9 exons events, 24100 reads/ 113556011 bases/ 185472 exons** | | | | | |
| GMAP | -f samse --cross-species -z sense_force | 91.05 | 66.95 | 20.76 | 2.83 |

| | | | | | |
|---|---|---|---|---|---|
| Graphmap2 | -x rnaseq | 88.39 | 77.07 | 56.22 | 14.77 |
| Minimap2 | -ax splice | 92.92 | 91.83 | 95.19 | 62.48 |
| Minimap2 + GTF | -ax splice –junc-bed anno.bed | 92.98 | 95.33 | 98.05 | 76.88 |
| deSALT | -d 10 | 93.08 | 94.83 | 97.98 | 73.24 |
| deSALT + GTF | -d 10 -G anno.info | 93.1 | 96.67 | 98.58 | 83.71 |
| **Mouse NS-ONT, coverage = 30X, > 9 exons events, 130341 reads/ 808079021 bases/ 3742468 exons** | | | | | |
| GMAP | -f samse --cross-species -z sense_force | 89.57 | 66.72 | 8.72 | 0.19 |
| Graphmap2 | -x rnaseq | 85.31 | 76.05 | 57.63 | 4.87 |
| Minimap2 | -ax splice | 93.16 | 93.19 | 96.98 | 27.98 |
| Minimap2 + GTF | -ax splice –junc-bed anno.bed | 93.32 | 96.97 | 99.01 | 55.17 |
| deSALT | -d 10 | 94.07 | 96.13 | 99.28 | 41.97 |
| deSALT + GTF | -d 10 -G anno.info | 94.12 | 98.05 | 99.48 | 65.31 |
| **Human NS-ONT, coverage = 4X, < 6 exons events, 17599 reads/ 101228979 bases/ 27897 exons** | | | | | |
| GMAP | -f samse --cross-species -z sense_force | 92.65 | 36.98 | 14.93 | 8.62 |
| Graphmap2 | -x rnaseq | 93.53 | 46.08 | 27.22 | 17.62 |
| Minimap2 | -ax splice | 93.93 | 83.76 | 96.91 | 76.78 |
| Minimap2 + GTF | -ax splice –junc-bed anno.bed | 93.93 | 84.87 | 97.44 | 77.92 |
| deSALT | -d 10 | 93.95 | 85.19 | 97.47 | 78.39 |
| deSALT + GTF | -d 10 -G anno.info | 93.95 | 85.92 | 97.71 | 79.26 |
| **Human NS-ONT, coverage = 4X, 6-9 exons events, 3736 reads/ 18143126 bases/ 27327 exons** | | | | | |
| GMAP | -f samse --cross-species -z sense_force | 90.96 | 66.44 | 21.76 | 2.52 |
| Graphmap2 | -x rnaseq | 89.87 | 76.42 | 56.56 | 14.32 |
| Minimap2 | -ax splice | 93.1 | 91.28 | 95.82 | 58.19 |
| Minimap2 + GTF | -ax splice –junc-bed anno.bed | 93.15 | 94.68 | 98.39 | 72.32 |
| deSALT | -d 10 | 93.25 | 94.35 | 98.15 | 70.96 |
| deSALT + GTF | -d 10 -G anno.info | 93.26 | 96.34 | 98.74 | 81.88 |
| **Human NS-ONT, coverage = 4X, > 9 exons events, 15668 reads/ 96675428 bases/ 441164 exons** | | | | | |
| GMAP | -f samse --cross-species -z sense_force | 89.41 | 66.89 | 9.51 | 0.26 |
| Graphmap2 | -x rnaseq | 82.57 | 75.63 | 63.51 | 6.22 |
| Minimap2 | -ax splice | 93.83 | 93.83 | 97.39 | 31.01 |
| Minimap2 + GTF | -ax splice –junc-bed anno.bed | 94.01 | 97.72 | 99.36 | 61.07 |
| deSALT | -d 10 | 94.1 | 96.33 | 99.14 | 44.36 |

| | | | | | |
|---|---|---|---|---|---|
| deSALT + GTF | -d 10 -G anno.info | 94.15 | 98.31 | 99.38 | 70.96 |
| **Human NS-ONT, coverage = 10X, < 6 exons events, 43206 reads/ 247192766 bases/ 70007 exons** | | | | | |
| GMAP | -f samse --cross-species -z sense_force | 92.69 | 38.06 | 15.36 | 8.66 |
| Graphmap2 | -x rnaseq | 93.56 | 47.05 | 27.73 | 17.8 |
| Minimap2 | -ax splice | 93.94 | 83.86 | 96.86 | 76.65 |
| Minimap2 + GTF | -ax splice –junc-bed anno.bed | 93.95 | 85.02 | 97.34 | 77.96 |
| deSALT | -d 10 | 93.96 | 85.38 | 97.46 | 78.39 |
| deSALT + GTF | -d 10 -G anno.info | 93.96 | 86.22 | 97.71 | 79.48 |
| **Human NS-ONT, coverage = 10X, 6-9 exons events, 9427 reads/ 46224956 bases/ 69805 exons** | | | | | |
| GMAP | -f samse --cross-species -z sense_force | 90.84 | 66.26 | 20.09 | 2.39 |
| Graphmap2 | -x rnaseq | 89.76 | 76.64 | 56.87 | 13.99 |
| Minimap2 | -ax splice | 92.97 | 91.21 | 95.43 | 59.98 |
| Minimap2 + GTF | -ax splice –junc-bed anno.bed | 93.02 | 94.77 | 98.4 | 73.61 |
| deSALT | -d 10 | 93.08 | 94.39 | 97.91 | 72.05 |
| deSALT + GTF | -d 10 -G anno.info | 93.09 | 96.34 | 98.75 | 82.2 |
| **Human NS-ONT, coverage = 10X, > 9 exons events, 37370 reads/ 231888845 bases/ 1061932 exons** | | | | | |
| GMAP | -f samse --cross-species -z sense_force | 89.24 | 66.75 | 9.33 | 0.18 |
| Graphmap2 | -x rnaseq | 81.97 | 72.51 | 56.38 | 5.75 |
| Minimap2 | -ax splice | 93.81 | 93.91 | 97.62 | 30.73 |
| Minimap2 + GTF | -ax splice –junc-bed anno.bed | 93.98 | 97.79 | 99.43 | 61.07 |
| deSALT | -d 10 | 94.07 | 96.38 | 99.34 | 44.59 |
| deSALT + GTF | -d 10 -G anno.info | 94.11 | 98.34 | 99.5 | 70.68 |
| **Human NS-ONT, coverage = 30X, < 6 exons events, 129512 reads/ 742391528 bases/ /208604 exons** | | | | | |
| GMAP | -f samse --cross-species -z sense_force | 92.74 | 37.48 | 15.17 | 8.52 |
| Graphmap2 | -x rnaseq | 93.59 | 46.13 | 27.29 | 17.52 |
| Minimap2 | -ax splice | 94.01 | 83.55 | 96.83 | 76.32 |
| Minimap2 + GTF | -ax splice –junc-bed anno.bed | 94.01 | 84.77 | 97.35 | 77.63 |
| deSALT | -d 10 | 94.03 | 85.28 | 97.56 | 78.29 |
| deSALT + GTF | -d 10 -G anno.info | 94.03 | 86.1 | 97.79 | 79.34 |
| **Human NS-ONT, coverage = 30X, 6-9 exons events, 27580 reads/ 134175002 bases/ 202377 exons** | | | | | |
| GMAP | -f samse --cross-species -z sense_force | 90.86 | 66.4 | 20.63 | 2.87 |
| Graphmap2 | -x rnaseq | 89.6 | 75.84 | 55.51 | 13.34 |

| | | | | | |
|---|---|---|---|---|---|
| Minimap2 | -ax splice | 92.94 | 91.38 | 95.7 | 60.42 |
| Minimap2 + GTF | -ax splice –junc-bed anno.bed | 92.98 | 94.79 | 98.3 | 74.21 |
| deSALT | -d 10 | 93.09 | 94.44 | 98.01 | 72.56 |
| deSALT + GTF | -d 10 -G anno.info | 93.1 | 96.37 | 98.66 | 83.14 |
| **Human NS-ONT, coverage = 30X, > 9 exons events, 112911 reads/ 698979855 bases/ 3200470 exons** | | | | | |
| GMAP | -f samse --cross-species -z sense_force | 89.44 | 66.9 | 9.38 | 0.22 |
| Graphmap2 | -x rnaseq | 91.73 | 70.73 | 51.06 | 4.69 |
| Minimap2 | -ax splice | 93.87 | 93.92 | 97.43 | 31.06 |
| Minimap2 + GTF | -ax splice –junc-bed anno.bed | 94.05 | 97.79 | 99.41 | 61.38 |
| deSALT | -d 10 | 94.13 | 96.32 | 99.29 | 43.98 |
| deSALT + GTF | -d 10 -G anno.info | 94.18 | 98.31 | 99.49 | 70.13 |
| **Human all protein coding genes NS-ONT, < 6 exons events, 797474 reads/1055126496 bases/2269595 exons** | | | | | |
| GMAP | -f samse --cross-species -z sense_force | 92.5 | 40.51 | 16.88 | 9.41 |
| Graphmap2 | -x rnaseq | 93.22 | 47.41 | 27.97 | 17.7 |
| Minimap2 | -ax splice | 93.7 | 83.67 | 96.57 | 75.91 |
| Minimap2 + GTF | -ax splice –junc-bed anno.bed | 93.7 | 85.01 | 97.18 | 77.36 |
| deSALT | -d 10 | 93.73 | 85.54 | 97.23 | 77.97 |
| deSALT + GTF | -d 10 -G anno.info | 93.73 | 86.37 | 97.47 | 79.08 |
| **Human all protein coding genes NS-ONT, 6-9 exons events, 269700 reads/516491645 bases/1938060 exons** | | | | | |
| GMAP | -f samse --cross-species -z sense_force | 91.05 | 66.41 | 20.49 | 2.76 |
| Graphmap2 | -x rnaseq | 89.25 | 74.79 | 52.69 | 11.77 |
| Minimap2 | -ax splice | 93.97 | 90.76 | 94.4 | 59.09 |
| Minimap2 + GTF | -ax splice –junc-bed anno.bed | 93.01 | 94.26 | 97.63 | 72.18 |
| deSALT | -d 10 | 93.05 | 94.04 | 97.4 | 71.13 |
| deSALT + GTF | -d 10 -G anno.info | 93.07 | 95.88 | 98.13 | 80.92 |
| **Human all protein coding genes NS-ONT, > 9 exons events, 299907 reads/1053041154 bases/5102557 exons** | | | | | |
| GMAP | -f samse --cross-species -z sense_force | 89.12 | 65.78 | 8.83 | 0.2 |
| Graphmap2 | -x rnaseq | 81.49 | 71.85 | 56.09 | 4.63 |
| Minimap2 | -ax splice | 92.76 | 91.31 | 95.61 | 29.16 |
| Minimap2 + GTF | -ax splice –junc-bed anno.bed | 92.97 | 95.54 | 97.87 | 58.95 |
| deSALT | -d 10 | 93.34 | 94.7 | 98.18 | 42.68 |
| deSALT + GTF | -d 10 -G anno.info | 93.4 | 96.8 | 98.44 | 68.73 |

a) This table depicts the results of the aligners on the reads from the transcripts with various numbers of exons. For each of the simulated datasets, the reads from the transcripts with < 6 exons, 6–9 exons, and >9 exons are respectively divided into three groups, and the assessment was separately implemented for all the groups.

b) Refer to the Supplementary Notes for the command lines of the benchmarked aligners.

c) Base%: the proportion of bases being correctly aligned to their ground truth positions, i.e., the mapped positions of the bases are within 5 bp of their ground truth positions.

d) Exon%: the proportion of the exons being correctly mapped. An exon in a certain read is considered to be correctly mapped only if its two boundaries are mapped within 5 bp of their ground truth positions in the reference genome.

e) Read80%: the proportion of Read80% reads. A read is considered to be a Read80% read only if it meets two conditions: $N_T/N_G > 80\%$ and $N_T/N_P > 80\%$, where $N_G$ is the number of ground truth exons within the read, $N_P$ is the number of exons predicted by the alignment, and $N_T$ is the number of true positive exons. Herein, a predicted exon is considered to be a true positive exon only if there is a ground truth exon in the read and the corresponding boundaries of the predicted exon and the ground truth exon are within 5 bp.

f) Read100%: the proportion of Read100% reads. A read is considered to be a Read100% read only if it meets two conditions: $N_T/N_G = 100\%$ and $N_T/N_P = 100\%$. It is worth noting that a Read100% read indicates that the read has a highly correct full-length alignment.

#### Supplementary Table 5. Benchmark results on the simulated reads from the genes with single/multiple isoforms [a]

| Aligner | parameters [b] | Base% [c] | Exons% [d] | Read80% [e] | Read100% [f] |
|---|---|---|---|---|---|
| **Simulated PacBio ROI datasets** | | | | | |
| **Fruit fly PacBio ROI, coverage = 4X, reads from the genes with single isoform, 14301 reads/15998364 bases/33963 exons** | | | | | |
| GMAP | -f samse --cross-species -z sense_force | 98.9 | 95.98 | 98.6 | 92.51 |
| Graphmap2 | -x rnaseq | 98.72 | 96.76 | 99.01 | 94.37 |
| Minimap2 | -ax splice | 98.88 | 97.48 | 98.61 | 95.12 |
| Minimap2 + GTF | -ax splice –junc-bed anno.bed | 98.88 | 97.59 | 98.6 | 95.29 |
| deSALT | -d 10 -x ccs | 99 | 98.13 | 99.33 | 96.36 |
| deSALT + GTF | -d 10 -x ccs -G anno.info | 99 | 98.13 | 99.29 | 96.43 |
| **Fruit fly PacBio ROI, coverage = 4X, reads from the genes with multiple isoforms, 30995 reads/86312808 bases/196424 exons** | | | | | |
| GMAP | -f samse --cross-species -z sense_force | 99.76 | 96.56 | 98.18 | 87.14 |
| Graphmap2 | -x rnaseq | 99.05 | 96.47 | 98.17 | 86.75 |
| Minimap2 | -ax splice | 99.67 | 98.1 | 98.81 | 91.7 |
| Minimap2 + GTF | -ax splice –junc-bed anno.bed | 99.68 | 98.96 | 99.23 | 94.15 |
| deSALT | -d 10 -x ccs | 99.78 | 98.76 | 99.65 | 94.3 |
| deSALT + GTF | -d 10 -x ccs -G anno.info | 99.78 | 98.78 | 99.67 | 94.38 |
| **Fruit fly PacBio ROI, coverage = 10X, reads from the genes with single isoform, 31515 reads/39591445 bases/80112 exons** | | | | | |
| GMAP | -f samse --cross-species -z sense_force | 98.9 | 96.21 | 98.43 | 92.84 |
| Graphmap2 | -x rnaseq | 98.77 | 97.28 | 99.11 | 95.12 |
| Minimap2 | -ax splice | 98.89 | 97.91 | 99.04 | 95.89 |
| Minimap2 + GTF | -ax splice –junc-bed anno.bed | 98.9 | 98.06 | 99.04 | 96.12 |
| deSALT | -d 10 -x ccs | 99.04 | 98.41 | 99.37 | 96.94 |
| deSALT + GTF | -d 10 -x ccs -G anno.info | 99.04 | 97.28/99.52 | 99.31 | 96.98 |
| **Fruit fly PacBio ROI, coverage = 10X, reads from the genes with multiple isoforms, 69775 reads/215258083 bases/481361 exons** | | | | | |
| GMAP | -f samse --cross-species -z sense_force | 99.77 | 96.69 | 97.97 | 86.54 |
| Graphmap2 | -x rnaseq | 99.12 | 96.54 | 98.13 | 86.43 |
| Minimap2 | -ax splice | 99.7 | 98.4 | 99.2 | 92.16 |
| Minimap2 + GTF | -ax splice –junc-bed anno.bed | 99.71 | 99.19 | 99.62 | 94.9 |
| deSALT | -d 10 -x ccs | 99.79 | 98.99 | 99.69 | 94.76 |

| | | | | | |
|---|---|---|---|---|---|
| deSALT + GTF | -d 10 -x ccs -G anno.info | 99.79 | 99 | 99.7 | 94.82 |
| **Fruit fly PacBio ROI, coverage = 30X, reads from the genes with single isoform, 88943 reads/118375102 bases/234312 exons** | | | | | |
| GMAP | -f samse --cross-species -z sense_force | 98.95 | 96.28 | 98.32 | 92.75 |
| Graphmap2 | -x rnaseq | 98.78 | 97.44 | 99.11 | 95.39 |
| Minimap2 | -ax splice | 98.95 | 98.24 | 99.4 | 96.41 |
| Minimap2 + GTF | -ax splice –junc-bed anno.bed | 98.95 | 98.38 | 99.4 | 96.67 |
| deSALT | -d 10 -x ccs | 99.05 | 98.53 | 99.32 | 97.13 |
| deSALT + GTF | -d 10 -x ccs -G anno.info | 99.04 | 98.49 | 99.25 | 97.19 |
| **Fruit fly PacBio ROI, coverage = 30X, reads from the genes with multiple isoforms, 199412 reads/645473928 bases/1432478 exons** | | | | | |
| GMAP | -f samse --cross-species -z sense_force | 99.77 | 96.68 | 97.87 | 86.02 |
| Graphmap2 | -x rnaseq | 99.12 | 96.56 | 98.07 | 86.12 |
| Minimap2 | -ax splice | 99.74 | 98.47 | 99.4 | 92.19 |
| Minimap2 + GTF | -ax splice –junc-bed anno.bed | 99.75 | 99.26 | 99.83 | 95.1 |
| deSALT | -d 10 -x ccs | 99.81 | 99.1 | 99.74 | 95 |
| deSALT + GTF | -d 10 -x ccs -G anno.info | 99.81 | 99.1 | 99.75 | 95.03 |
| **Mouse PacBio ROI, coverage = 4X, reads from the genes with single isoform, 12435 reads/12982861 bases/27141 exons** | | | | | |
| GMAP | -f samse --cross-species -z sense_force | 96.28 | 93.78 | 97.36 | 91.71 |
| Graphmap2 | -x rnaseq | 94.3 | 86.92 | 94.01 | 85.95 |
| Minimap2 | -ax splice | 97.4 | 93.63 | 96.24 | 91.5 |
| Minimap2 + GTF | -ax splice –junc-bed anno.bed | 97.56 | 94.86 | 96.86 | 92.55 |
| deSALT | -d 10 -x ccs | 96.2 | 93.2 | 96.41 | 91.57 |
| deSALT + GTF | -d 10 -x ccs -G anno.info | 96.25 | 93.5 | 96.54 | 92.01 |
| **Mouse PacBio ROI, coverage = 4X, reads from the genes with multiple isoforms, 34377 reads/53169001 bases/205670 exons** | | | | | |
| GMAP | -f samse --cross-species -z sense_force | 98.8 | 97.61 | 98.18 | 89.72 |
| Graphmap2 | -x rnaseq | 93.95 | 89.14 | 92.57 | 76.54 |
| Minimap2 | -ax splice | 98.46 | 96.17 | 96.25 | 88.85 |
| Minimap2 + GTF | -ax splice –junc-bed anno.bed | 99.04 | 97.74 | 97.48 | 91.05 |
| deSALT | -d 10 -x ccs | 99.03 | 97.82 | 98.12 | 91.94 |
| deSALT + GTF | -d 10 -x ccs -G anno.info | 99.03 | 97.82 | 98.11 | 92.02 |
| **Mouse PacBio ROI, coverage = 10X, reads from the genes with single isoform, 27414 reads/32112338 bases/63714 exons** | | | | | |
| GMAP | -f samse --cross-species -z sense_force | 96.28 | 91.85 | 96.4 | 89.13 |
| Graphmap2 | -x rnaseq | 94.42 | 87.41 | 94.04 | 86.41 |

| | | | | | |
|---|---|---|---|---|---|
| Minimap2 | -ax splice | 97.66 | 94.63 | 97.05 | 93.01 |
| Minimap2 + GTF | -ax splice –junc-bed anno.bed | 97.85 | 95.83 | 97.65 | 94.08 |
| deSALT | -d 10 -x ccs | 96.36 | 93.63 | 96.4 | 92.29 |
| deSALT + GTF | -d 10 -x ccs -G anno.info | 96.42 | 93.91 | 96.48 | 92.73 |
| **Mouse PacBio ROI, coverage = 10X, reads from the genes with multiple isoforms, 76084 reads/132081429 bases/500451 exons** | | | | | |
| GMAP | -f samse --cross-species -z sense_force | 98.79 | 95.03 | 95.86 | 82.52 |
| Graphmap2 | -x rnaseq | 93.86 | 89.33 | 92.3 | 76.24 |
| Minimap2 | -ax splice | 98.55 | 96.7 | 96.97 | 90.11 |
| Minimap2 + GTF | -ax splice –junc-bed anno.bed | 99.12 | 98.22 | 98.26 | 92.48 |
| deSALT | -d 10 -x ccs | 99.11 | 98.2 | 98.4 | 92.9 |
| deSALT + GTF | -d 10 -x ccs -G anno.info | 99.12 | 98.21 | 98.38 | 93.05 |
| **Mouse PacBio ROI, coverage = 30X, reads from the genes with single isoform, 77424 reads/95970680 bases/185808 exons** | | | | | |
| GMAP | -f samse --cross-species -z sense_force | 96.32 | 92.12 | 96.43 | 89.29 |
| Graphmap2 | -x rnaseq | 94.48 | 87.53 | 93.94 | 86.62 |
| Minimap2 | -ax splice | 97.65 | 94.87 | 97.26 | 93.59 |
| Minimap2 + GTF | -ax splice –junc-bed anno.bed | 97.81 | 96.01 | 97.95 | 94.76 |
| deSALT | -d 10 -x ccs | 96.34 | 93.87 | 96.46 | 92.63 |
| deSALT + GTF | -d 10 -x ccs -G anno.info | 96.39 | 94.19 | 96.56 | 93.22 |
| **Mouse PacBio ROI, coverage = 30X, reads from the genes with multiple isoforms, 215360 reads/395448314 bases/1483790 exons** | | | | | |
| GMAP | -f samse --cross-species -z sense_force | 98.8 | 95.14 | 95.75 | 82.49 |
| Graphmap2 | -x rnaseq | 93.86 | 89.43 | 92.12 | 76.21 |
| Minimap2 | -ax splice | 98.56 | 96.95 | 97.3 | 90.78 |
| Minimap2 + GTF | -ax splice –junc-bed anno.bed | 99.13 | 98.44 | 98.64 | 93.24 |
| deSALT | -d 10 -x ccs | 99.1 | 98.38 | 98.49 | 93.43 |
| deSALT + GTF | -d 10 -x ccs -G anno.info | 99.1 | 98.39 | 98.46 | 93.56 |
| **Human PacBio ROI, coverage = 4X, reads from the genes with single isoform, 12012 reads/10139594 bases/22474 exons** | | | | | |
| GMAP | -f samse --cross-species -z sense_force | 98.54 | 93.64 | 97.5 | 92.38 |
| Graphmap2 | -x rnaseq | 97.11 | 90.74 | 95.83 | 90.87 |
| Minimap2 | -ax splice | 98.79 | 95.19 | 97.07 | 94.03 |
| Minimap2 + GTF | -ax splice –junc-bed anno.bed | 98.94 | 96.63 | 97.89 | 95.45 |
| deSALT | -d 10 -x ccs | 98.96 | 96.32 | 98.14 | 95.55 |
| deSALT + GTF | -d 10 -x ccs -G anno.info | 99.07 | 97.05 | 98.43 | 96.33 |

| | | | | | |
|---|---|---|---|---|---|
| **Human PacBio ROI, coverage = 4X, reads from the genes with multiple isoforms, 34881 reads/28674816 bases/83324 exons** | | | | | |
| GMAP | -f samse --cross-species -z sense_force | 98.97 | 95.31 | 96.78 | 84.48 |
| Graphmap2 | -x rnaseq | 95.6 | 90.98 | 93.83 | 77.7 |
| Minimap2 | -ax splice | 98.62 | 96.62 | 96.7 | 89.69 |
| Minimap2 + GTF | -ax splice –junc-bed anno.bed | 99.14 | 97.4 | 97.67 | 90.57 |
| deSALT | -d 10 -x ccs | 99.24 | 97.98 | 98.47 | 92.71 |
| deSALT + GTF | -d 10 -x ccs -G anno.info | 99.26 | 98 | 98.45 | 92.85 |
| **Human PacBio ROI, coverage = 10X, reads from the genes with single isoform, 26452 reads/25006370 bases/52122 exons** | | | | | |
| GMAP | -f samse --cross-species -z sense_force | 98.59 | 93.84 | 97.46 | 92.32 |
| Graphmap2 | -x rnaseq | 97.04 | 91.02 | 95.73 | 91.36 |
| Minimap2 | -ax splice | 98.93 | 95.69 | 97.61 | 94.8 |
| Minimap2 + GTF | -ax splice –junc-bed anno.bed | 98.98 | 97.28 | 98.51 | 96.5 |
| deSALT | -d 10 -x ccs | 98.97 | 96.65 | 98.29 | 96.05 |
| deSALT + GTF | -d 10 -x ccs -G anno.info | 99.1 | 97.48 | 98.63 | 96.93 |
| **Human PacBio ROI, coverage = 10X, reads from the genes with multiple isoforms, 76887 reads/116762997 bases/485794 exons** | | | | | |
| GMAP | -f samse --cross-species -z sense_force | 99.03 | 95.56 | 96.59 | 84.13 |
| Graphmap2 | -x rnaseq | 95.49 | 91.16 | 93.73 | 77.6 |
| Minimap2 | -ax splice | 98.73 | 97.32 | 97.58 | 91.31 |
| Minimap2 + GTF | -ax splice –junc-bed anno.bed | 99.24 | 97.9 | 98.43 | 91.84 |
| deSALT | -d 10 -x ccs | 99.28 | 98.4 | 98.67 | 93.75 |
| deSALT + GTF | -d 10 -x ccs -G anno.info | 99.29 | 98.41 | 98.58 | 93.84 |
| **Human PacBio ROI, coverage = 30X, reads from the genes with single isoform, 74648 reads/74627529 bases/151031 exons** | | | | | |
| GMAP | -f samse --cross-species -z sense_force | 98.7 | 94.11 | 97.49 | 92.77 |
| Graphmap2 | -x rnaseq | 97.1 | 93.13 | 95.84 | 91.6 |
| Minimap2 | -ax splice | 98.97 | 96.14 | 98.03 | 95.47 |
| Minimap2 + GTF | -ax splice –junc-bed anno.bed | 99.11 | 97.74 | 98.93 | 97.06 |
| deSALT | -d 10 -x ccs | 99.11 | 97.06 | 98.46 | 96.4 |
| deSALT + GTF | -d 10 -x ccs -G anno.info | 99.15 | 97.72 | 98.72 | 97.38 |
| **Human PacBio ROI, coverage = 30X, reads from the genes with multiple isoforms, 217469 reads/349383354 bases/1440386 exons** | | | | | |
| GMAP | -f samse --cross-species -z sense_force | 99.02 | 95.6 | 96.56 | 84.17 |
| Graphmap2 | -x rnaseq | 95.55 | 91.27 | 93.62 | 77.52 |
| Minimap2 | -ax splice | 98.76 | 97.48 | 97.99 | 92.08 |

| | | | | | |
|---|---|---|---|---|---|
| Minimap2 + GTF | -ax splice –junc-bed anno.bed | 99.26 | 98.08 | 98.76 | 92.46 |
| deSALT | -d 10 -x ccs | 99.28 | 98.53 | 98.79 | 94.39 |
| deSALT + GTF | -d 10 -x ccs -G anno.info | 99.31 | 98.56 | 98.7 | 94.58 |
| **Human all protein coding genes PacBio ROI, reads from the genes with single isoform, 104865 reads/ 185990737 bases/ 362830 exons** | | | | | |
| GMAP | -f samse --cross-species -z sense_force | 98.68 | 94.17 | 96.9 | 89.35 |
| Graphmap2 | -x rnaseq | 96.34 | 88.36 | 94.35 | 85.74 |
| Minimap2 | -ax splice | 98.82 | 96.75 | 98.18 | 94.47 |
| Minimap2 + GTF | -ax splice –junc-bed anno.bed | 98.9 | 97.03 | 98.32 | 94.75 |
| deSALT | -d 10 -x ccs | 99.1 | 96.63 | 98.24 | 92.64 |
| deSALT + GTF | -d 10 -x ccs -G anno.info | 98.87 | 96.51 | 98.22 | 92.68 |
| **Human all protein coding genes PacBio ROI, reads from the genes with multiple isoform, 1271228 reads/ 2234007833 bases/ 8387942 exons** | | | | | |
| GMAP | -f samse --cross-species -z sense_force | 99.52 | 95.64 | 97.23 | 83.98 |
| Graphmap2 | -x rnaseq | 95.94 | 90.22 | 93.42 | 76.02 |
| Minimap2 | -ax splice | 99.33 | 97.14 | 98.42 | 90.35 |
| Minimap2 + GTF | -ax splice –junc-bed anno.bed | 99.65 | 9802 | 99.09 | 91.95 |
| deSALT | -d 10 -x ccs | 99.82 | 98.34 | 99.29 | 92.15 |
| deSALT + GTF | -d 10 -x ccs -G anno.info | 99.82 | 98.46 | 99.35 | 92.65 |
| **Simulated ONT 2D (1D$^2$) datasets** | | | | | |
| **Fruit fly ONT 2D (1D$^2$), coverage = 4X, reads from the genes with single isoform, 14295 reads/15799802 bases/34025 exons** | | | | | |
| GMAP | -f samse --cross-species -z sense_force | 92.81 | 72.32 | 83.39 | 55.54 |
| Graphmap2 | -x rnaseq | 94.48 | 87.67 | 95.4 | 77.11 |
| Minimap2 | -ax splice | 93.25 | 86.09 | 87.17 | 72.47 |
| Minimap2 + GTF | -ax splice –junc-bed anno.bed | 93.37 | 87.64 | 87.93 | 75.09 |
| deSALT | -d 10 -x ont2d -s 2 | 94.46 | 89.32 | 94.35 | 78.44 |
| deSALT + GTF | -d 10 -x ont2d -s 2 -G anno.info | 94.57 | 92.21 | 95.87 | 83.93 |
| **Fruit fly ONT 2D (1D$^2$), coverage = 4X, reads from the genes with multiple isoforms, 30150 reads/86067883 bases/195082 exons** | | | | | |
| GMAP | -f samse --cross-species -z sense_force | 94.52 | 74.49 | 69.66 | 38.2 |
| Graphmap2 | -x rnaseq | 95.1 | 92.55 | 95.23 | 71.06 |
| Minimap2 | -ax splice | 95.48 | 93.78 | 93.51 | 72.41 |
| Minimap2 + GTF | -ax splice –junc-bed anno.bed | 95.55 | 95.94 | 95.13 | 79.02 |

| | | | | | |
|---|---|---|---|---|---|
| deSALT | -d 10 -x ont2d -s 2 | 95.84 | 96.32 | 97.45 | 81.37 |
| deSALT + GTF | -d 10 -x ont2d -s 2 -G anno.info | 95.88 | 96.77 | 97.99 | 83.36 |
| **Fruit fly ONT 2D (1D$^2$), coverage = 10X, reads from the genes with single isoform, 31482 reads/39446943 bases/80732 exons** | | | | | |
| GMAP | -f samse --cross-species -z sense_force | 93.25 | 73.1 | 84.47 | 55.37 |
| Graphmap2 | -x rnaseq | 94.66 | 90.8 | 97.57 | 80.59 |
| Minimap2 | -ax splice | 94.14 | 89.61 | 94.84 | 78.23 |
| Minimap2 + GTF | -ax splice –junc-bed anno.bed | 94.27 | 91.31 | 95.73 | 81.15 |
| deSALT | -d 10 -x ont2d -s 2 | 94.78 | 92.69 | 97.18 | 83.68 |
| deSALT + GTF | -d 10 -x ont2d -s 2 -G anno.info | 94.86 | 94.37 | 97.75 | 87.53 |
| **Fruit fly ONT 2D (1D$^2$), coverage = 10X, reads from the genes with multiple isoforms, 67498 reads/215154115 bases/479977 exons** | | | | | |
| GMAP | -f samse --cross-species -z sense_force | 94.69 | 75.15 | 68.05 | 36.2 |
| Graphmap2 | -x rnaseq | 95.09 | 93.68 | 96.25 | 73.15 |
| Minimap2 | -ax splice | 95.69 | 94.83 | 96.35 | 74.35 |
| Minimap2 + GTF | -ax splice –junc-bed anno.bed | 95.76 | 96.87 | 98.05 | 81.61 |
| deSALT | -d 10 -x ont2d -s 2 | 95.94 | 97.48 | 98.53 | 85.05 |
| deSALT + GTF | -d 10 -x ont2d -s 2 -G anno.info | 95.96 | 97.75 | 98.84 | 86.32 |
| **Fruit fly ONT 2D (1D$^2$), coverage = 30X, reads from the genes with single isoform, 91597 reads/118351500 bases/239087 exons** | | | | | |
| GMAP | -f samse --cross-species -z sense_force | 93.24 | 73.41 | 83.99 | 55.02 |
| Graphmap2 | -x rnaseq | 94.66 | 91.99 | 97.93 | 82.61 |
| Minimap2 | -ax splice | 94.19 | 89.97 | 94.72 | 78.36 |
| Minimap2 + GTF | -ax splice –junc-bed anno.bed | 94.31 | 91.68 | 95.64 | 81.51 |
| deSALT | -d 10 -x ont2d -s 2 | 94.88 | 94.51 | 97.78 | 87.25 |
| deSALT + GTF | -d 10 -x ont2d -s 2 -G anno.info | 94.91 | 95.17 | 97.8 | 88.84 |
| **Fruit fly ONT 2D (1D$^2$), coverage = 30X, reads from the genes with multiple isoforms, 196740 reads/645463489 bases/1431878 exons** | | | | | |
| GMAP | -f samse --cross-species -z sense_force | 94.67 | 75.02 | 67.53 | 35.7 |
| Graphmap2 | -x rnaseq | 95.15 | 94.06 | 96.55 | 74.4 |
| Minimap2 | -ax splice | 95.72 | 95.02 | 96.46 | 75.05 |
| Minimap2 + GTF | -ax splice –junc-bed anno.bed | 95.78 | 97.04 | 98.15 | 82.22 |
| deSALT | -d 10 -x ont2d -s 2 | 95.96 | 97.74 | 98.79 | 86.45 |
| deSALT + GTF | -d 10 -x ont2d -s 2 -G anno.info | 95.98 | 97.97 | 99 | 87.39 |
| **Mouse ONT 2D (1D$^2$), coverage = 4X, reads from the genes with single isoform, 12412 reads/12823106 bases/27050 exons** | | | | | |
| GMAP | -f samse --cross-species -z sense_force | 88.16 | 63.35 | 79.66 | 50.76 |

| | | | | | |
|---|---|---|---|---|---|
| Graphmap2 | -x rnaseq | 89.79 | 74.01 | 88.53 | 66.56 |
| Minimap2 | -ax splice | 90.44 | 77.27 | 80.98 | 63.37 |
| Minimap2 + GTF | -ax splice –junc-bed anno.bed | 90.57 | 79.01 | 82.04 | 65.01 |
| deSALT | -d 10 -x ont2d -s 2 | 90.55 | 80.03 | 88.62 | 69.1 |
| deSALT + GTF | -d 10 -x ont2d -s 2 -G anno.info | 90.85 | 84.87 | 90.85 | 76.38 |
| **Mouse ONT 2D (1D$^2$), coverage = 4X, reads from the genes with multiple isoforms, 34179 reads/52761225 bases/205091 exons** | | | | | |
| GMAP | -f samse --cross-species -z sense_force | 91.08 | 69.2 | 63.74 | 36.27 |
| Graphmap2 | -x rnaseq | 89.31 | 81.19 | 83.55 | 54.8 |
| Minimap2 | -ax splice | 92.28 | 87.01 | 81.33 | 57.91 |
| Minimap2 + GTF | -ax splice –junc-bed anno.bed | 92.98 | 89.49 | 83.63 | 61.63 |
| deSALT | -d 10 -x ont2d -s 2 | 94.39 | 93.01 | 92 | 73.07 |
| deSALT + GTF | -d 10 -x ont2d -s 2 -G anno.info | 94.47 | 93.68 | 92.83 | 75.5 |
| **Mouse ONT 2D (1D$^2$), coverage = 10X, reads from the genes with single isoform, 27346 reads/31974136 bases/63843 exons** | | | | | |
| GMAP | -f samse --cross-species -z sense_force | 88.8 | 64.56 | 81.71 | 51.2 |
| Graphmap2 | -x rnaseq | 90.02 | 77.75 | 91.07 | 69.37 |
| Minimap2 | -ax splice | 91.12 | 80.88 | 88.63 | 69.04 |
| Minimap2 + GTF | -ax splice –junc-bed anno.bed | 91.32 | 82.27 | 89.77 | 70.85 |
| deSALT | -d 10 -x ont2d -s 2 | 91.36 | 85.62 | 92.66 | 76.46 |
| deSALT + GTF | -d 10 -x ont2d -s 2 -G anno.info | 91.51 | 88.3 | 93.56 | 81.36 |
| **Mouse ONT 2D (1D$^2$), coverage = 10X, reads from the genes with multiple isoforms, 75533 reads/131788448 bases/502041 exons** | | | | | |
| GMAP | -f samse --cross-species -z sense_force | 91.34 | 70.11 | 62.89 | 35.39 |
| Graphmap2 | -x rnaseq | 89.35 | 83.07 | 85.23 | 57.64 |
| Minimap2 | -ax splice | 92.95 | 89.25 | 88.1 | 63.05 |
| Minimap2 + GTF | -ax splice –junc-bed anno.bed | 93.63 | 91.71 | 90.85 | 67.14 |
| deSALT | -d 10 -x ont2d -s 2 | 94.76 | 94.8 | 94.84 | 78.19 |
| deSALT + GTF | -d 10 -x ont2d -s 2 -G anno.info | 94.79 | 95.14 | 95.26 | 79.62 |
| **Mouse ONT 2D (1D$^2$), coverage = 30X, reads from the genes with single isoform, 79605 reads/95941227 bases/189014 exons** | | | | | |
| GMAP | -f samse --cross-species -z sense_force | 88.89 | 65.11 | 81.37 | 51.56 |
| Graphmap2 | -x rnaseq | 90.12 | 78.71 | 91.38 | 70.47 |
| Minimap2 | -ax splice | 91.4 | 81.73 | 89.14 | 69.64 |
| Minimap2 + GTF | -ax splice –junc-bed anno.bed | 91.59 | 83.59 | 90.44 | 71.54 |
| deSALT | -d 10 -x ont2d -s 2 | 91.41 | 87.49 | 93.41 | 79.52 |

| | | | | | |
|---|---|---|---|---|---|
| deSALT + GTF | -d 10 -x ont2d -s 2 -G anno.info | 91.53 | 88.92 | 93.73 | 82.2 |
| **Mouse ONT 2D (1D$^2$), coverage = 30X, reads from the genes with multiple isoforms, 219808 reads/395387844 bases/1499062 exons** | | | | | |
| GMAP | -f samse --cross-species -z sense_force | 91.42 | 70 | 62.18 | 35.02 |
| Graphmap2 | -x rnaseq | 89.36 | 83.42 | 95.54 | 58.23 |
| Minimap2 | -ax splice | 93 | 89.43 | 88.36 | 63.32 |
| Minimap2 + GTF | -ax splice –junc-bed anno.bed | 93.69 | 92.03 | 90.94 | 67.59 |
| deSALT | -d 10 -x ont2d -s 2 | 94.73 | 94.96 | 95 | 78.89 |
| deSALT + GTF | -d 10 -x ont2d -s 2 -G anno.info | 94.75 | 95.24 | 95.33 | 80.15 |
| **Human ONT 2D (1D$^2$), coverage = 4X, reads from the genes with single isoform, 11992 reads/9984792 bases/22360 exons** | | | | | |
| GMAP | -f samse --cross-species -z sense_force | 90.59 | 63.81 | 81.8 | 54.48 |
| Graphmap2 | -x rnaseq | 91.94 | 75.75 | 89.66 | 70.3 |
| Minimap2 | -ax splice | 90.88 | 74.24 | 78.68 | 63.31 |
| Minimap2 + GTF | -ax splice –junc-bed anno.bed | 91.03 | 76.0 | 79.79 | 64.81 |
| deSALT | -d 10 -x ont2d -s 2 | 93.29 | 80.05 | 89.15 | 71.68 |
| deSALT + GTF | -d 10 -x ont2d -s 2 -G anno.info | 93.7 | 86.61 | 92.19 | 79.69 |
| **Human ONT 2D (1D$^2$), coverage = 4X, reads from the genes with multiple isoforms, 34602 reads/46620001 bases/200665 exons** | | | | | |
| GMAP | -f samse --cross-species -z sense_force | 90.72 | 69.57 | 62.7 | 35.95 |
| Graphmap2 | -x rnaseq | 88.6 | 81.17 | 82.56 | 54.06 |
| Minimap2 | -ax splice | 91.96 | 86.83 | 80.4 | 57.18 |
| Minimap2 + GTF | -ax splice –junc-bed anno.bed | 92.59 | 88.73 | 82.02 | 60.3 |
| deSALT | -d 10 -x ont2d -s 2 | 94.59 | 93.12 | 91.79 | 73.23 |
| deSALT + GTF | -d 10 -x ont2d -s 2 -G anno.info | 94.64 | 93.73 | 92.67 | 75.41 |
| **Human ONT 2D (1D$^2$), coverage = 10X, reads from the genes with single isoform, 26423 reads/24855256 bases/52295 exons** | | | | | |
| GMAP | -f samse --cross-species -z sense_force | 91.23 | 65.39 | 83.43 | 54.93 |
| Graphmap2 | -x rnaseq | 92.38 | 78.76 | 92.29 | 72.58 |
| Minimap2 | -ax splice | 92.29 | 79.25 | 88.14 | 69.95 |
| Minimap2 + GTF | -ax splice –junc-bed anno.bed | 92.44 | 81.42 | 89.38 | 71.18 |
| deSALT | -d 10 -x ont2d -s 2 | 94.1 | 86.32 | 94.27 | 78.79 |
| deSALT + GTF | -d 10 -x ont2d -s 2 -G anno.info | 94.35 | 90.59 | 95.69 | 85.04 |
| **Human ONT 2D (1D$^2$), coverage = 10X, reads from the genes with multiple isoforms, 76497 reads/116428189 bases/490722 exons** | | | | | |
| GMAP | -f samse --cross-species -z sense_force | 90.8 | 69.63 | 61.51 | 35.12 |
| Graphmap2 | -x rnaseq | 88.7 | 82.76 | 84.22 | 56.08 |

| | | | | | |
|---|---|---|---|---|---|
| Minimap2 | -ax splice | 92.64 | 88.75 | 87.76 | 62.03 |
| Minimap2 + GTF | -ax splice –junc-bed anno.bed | 93.26 | 90.82 | 89.68 | 65.52 |
| deSALT | -d 10 -x ont2d -s 2 | 94.93 | 94.67 | 95.04 | 78.1 |
| deSALT + GTF | -d 10 -x ont2d -s 2 -G anno.info | 94.97 | 95.07 | 95.56 | 79.68 |
| **Human ONT 2D (1D$^2$), coverage = 30X, reads from the genes with single isoform, 76874 reads/74590314 bases/154293 exons** | | | | | |
| GMAP | -f samse --cross-species -z sense_force | 91.18 | 76.08 | 87.58 | 67.12 |
| Graphmap2 | -x rnaseq | 92.53 | 79.88 | 92.66 | 73.33 |
| Minimap2 | -ax splice | 92.43 | 79.84 | 88.39 | 70.21 |
| Minimap2 + GTF | -ax splice –junc-bed anno.bed | 92.57 | 81.84 | 89.61 | 72.08 |
| deSALT | -d 10 -x ont2d -s 2 | 94.31 | 88.97 | 94.94 | 82.3 |
| deSALT + GTF | -d 10 -x ont2d -s 2 -G anno.info | 94.52 | 91.56 | 95.87 | 86.07 |
| **Human ONT 2D (1D$^2$), coverage = 30X, reads from the genes with multiple isoforms, 222523 reads/349307842 bases/1461607 exons** | | | | | |
| GMAP | -f samse --cross-species -z sense_force | 90.91 | 79.36 | 73.4 | 44.68 |
| Graphmap2 | -x rnaseq | 88.67 | 82.78 | 83.94 | 56.43 |
| Minimap2 | -ax splice | 92.74 | 89.18 | 87.91 | 62.57 |
| Minimap2 + GTF | -ax splice –junc-bed anno.bed | 93.36 | 91.23 | 89.83 | 66.24 |
| deSALT | -d 10 -x ont2d -s 2 | 94.96 | 94.9 | 95.11 | 78.71 |
| deSALT + GTF | -d 10 -x ont2d -s 2 -G anno.info | 94.99 | 95.28 | 95.55 | 80.06 |
| **Human all protein coding genes ONT 2D (1D$^2$), reads from the genes with single isoform, 88893 reads/ 185919364 bases/ 351981 exons** | | | | | |
| GMAP | -f samse --cross-species -z sense_force | 94.56 | 69.13 | 76.81 | 48.12 |
| Graphmap2 | -x rnaseq | 94.13 | 80.3 | 89.14 | 67.16 |
| Minimap2 | -ax splice | 97.1 | 89.77 | 94.05 | 74.97 |
| Minimap2 + GTF | -ax splice –junc-bed anno.bed | 97.14 | 90.89 | 94.74 | 76.52 |
| deSALT | -d 10 -x ont2d -s 2 | 97.16 | 92.52 | 95.27 | 81.6 |
| deSALT + GTF | -d 10 -x ont2d -s 2 -G anno.info | 97.06 | 92.47 | 95.27 | 82.03 |
| **Human all protein coding genes ONT 2D (1D$^2$), reads from the genes with multiple isoforms, 1012793 reads/ 2228784360 bases/ 8070099exons** | | | | | |
| GMAP | -f samse --cross-species -z sense_force | 94.31 | 70.54 | 60.24 | 32.17 |
| Graphmap2 | -x rnaseq | 91.29 | 80.99 | 82.55 | 52.88 |
| Minimap2 | -ax splice | 96.91 | 91.51 | 90.93 | 65.78 |
| Minimap2 + GTF | -ax splice –junc-bed anno.bed | 97.3 | 93.42 | 92.75 | 69.87 |
| deSALT | -d 10 -x ont2d -s 2 | 98.22 | 95.84 | 95.99 | 79.42 |
| deSALT + GTF | -d 10 -x ont2d -s 2 -G anno.info | 98.23 | 96.34 | 96.4 | 81.42 |

| Simulated PacBio subread datasets | | | | | |
|---|---|---|---|---|---|
| Fruit fly PacBio subread, coverage = 4X, reads from the genes with single isoform, 14281 reads/15851135 bases/32392 exons | | | | | |
| GMAP | -f samse --cross-species -z sense_force | 73.19 | 53.19 | 56.82 | 36.44 |
| Graphmap2 | -x rnaseq | 87.17 | 84.09 | 93069 | 71.83 |
| Minimap2 | -ax splice | 85.65 | 80.93 | 79.25 | 62.91 |
| Minimap2 + GTF | -ax splice –junc-bed anno.bed | 85.79 | 83.05 | 80.19 | 66.23 |
| deSALT | -d 10 -x clr -s 2 | 87.17 | 87.62 | 93.45 | 75.34 |
| deSALT + GTF | -d 10 -x clr -s 2 -G anno.info | 87.29 | 89.41 | 94.34 | 78.67 |
| Fruit fly PacBio subread, coverage = 4X, reads from the genes with multiple isoforms, 29888 reads/86093995 bases/180025 exons | | | | | |
| GMAP | -f samse --cross-species -z sense_force | 83.55 | 64.11 | 50.04 | 24.67 |
| Graphmap2 | -x rnaseq | 87.96 | 91.1 | 94.25 | 66.61 |
| Minimap2 | -ax splice | 88.08 | 92.53 | 87.85 | 67.08 |
| Minimap2 + GTF | -ax splice –junc-bed anno.bed | 88.15 | 94.6 | 89.56 | 73.76 |
| deSALT | -d 10 -x clr -s 2 | 88.61 | 96.35 | 96.79 | 80.71 |
| deSALT + GTF | -d 10 -x clr -s 2 -G anno.info | 88.63 | 96.74 | 97.23 | 82.29 |
| Fruit fly PacBio subread, coverage = 10X, reads from the genes with single isoform, 31433 reads/39457740 bases/76495 exons | | | | | |
| GMAP | -f samse --cross-species -z sense_force | 73.32 | 54.06 | 55.63 | 35.48 |
| Graphmap2 | -x rnaseq | 87.34 | 87.73 | 96.6 | 75.14 |
| Minimap2 | -ax splice | 86.61 | 86.36 | 91.24 | 71.71 |
| Minimap2 + GTF | -ax splice –junc-bed anno.bed | 86.77 | 88.53 | 92.23 | 75.79 |
| deSALT | -d 10 -x clr -s 2 | 87.58 | 91.43 | 96.47 | 80.89 |
| deSALT + GTF | -d 10 -x clr -s 2 -G anno.info | 87.6 | 92.22 | 96.84 | 82.54 |
| Fruit fly PacBio subread, coverage = 10X, reads from the genes with multiple isoforms, 66477 reads/215155994 bases/442303 exons | | | | | |
| GMAP | -f samse --cross-species -z sense_force | 83.74 | 64.48 | 50.13 | 23.65 |
| Graphmap2 | -x rnaseq | 87.96 | 92.66 | 95.6 | 70.07 |
| Minimap2 | -ax splice | 88.44 | 94.09 | 95.27 | 72.54 |
| Minimap2 + GTF | -ax splice –junc-bed anno.bed | 88.51 | 96.36 | 97.08 | 80.01 |
| deSALT | -d 10 -x clr -s 2 | 88.72 | 97.26 | 98.47 | 84.47 |
| deSALT + GTF | -d 10 -x clr -s 2 -G anno.info | 88.73 | 97.48 | 98.68 | 85.5 |
| Fruit fly PacBio subread, coverage = 30X, reads from the genes with single isoform, 88646 reads/118338677 bases/223878 exons | | | | | |

| | | | | | |
|---|---|---|---|---|---|
| GMAP | -f samse --cross-species -z sense_force | 73.94 | 55.27 | 56.78 | 36 |
| Graphmap2 | -x rnaseq | 87.4 | 90.3 | 97.65 | 79.29 |
| Minimap2 | -ax splice | 87.03 | 88.6 | 95.02 | 75.72 |
| Minimap2 + GTF | -ax splice –junc-bed anno.bed | 87.17 | 90.74 | 95.97 | 79.67 |
| deSALT | -d 10 -x clr -s 2 | 87.68 | 92.96 | 97.38 | 83.84 |
| deSALT + GTF | -d 10 -x clr -s 2 -G anno.info | 87.68 | 93.26 | 97.34 | 84.53 |
| **Fruit fly PacBio subread, coverage = 30X, reads from the genes with multiple isoforms, 188895 reads/645460106 bases/1316757 exons** | | | | | |
| GMAP | -f samse --cross-species -z sense_force | 84.16 | 65.2 | 50.91 | 23.38 |
| Graphmap2 | -x rnaseq | 88.04 | 93.13 | 96.07 | 71.08 |
| Minimap2 | -ax splice | 88.53 | 94.55 | 96.46 | 73.59 |
| Minimap2 + GTF | -ax splice –junc-bed anno.bed | 88.6 | 96.82 | 98.32 | 81.38 |
| deSALT | -d 10 -x clr -s 2 | 88.75 | 97.59 | 98.8 | 85.22 |
| deSALT + GTF | -d 10 -x clr -s 2 -G anno.info | 88.75 | 97.83 | 98.95 | 86.33 |
| **Mouse PacBio subread, coverage = 4X, reads from the genes with single isoform, 12407 reads/12875901 bases/25870 exons** | | | | | |
| GMAP | -f samse --cross-species -z sense_force | 68.9 | 43.9 | 51.44 | 32.43 |
| Graphmap2 | -x rnaseq | 82.84 | 71.82 | 87.64 | 63.36 |
| Minimap2 | -ax splice | 82.84 | 72.44 | 74.45 | 56.44 |
| Minimap2 + GTF | -ax splice –junc-bed anno.bed | 82.99 | 74.29 | 75.38 | 58.14 |
| deSALT | -d 10 -x clr -s 2 | 83.27 | 76.92 | 85.62 | 64.28 |
| deSALT + GTF | -d 10 -x clr -s 2 -G anno.info | 83.5 | 80.67 | 87.85 | 69.36 |
| **Mouse PacBio subread, coverage = 4X, reads from the genes with multiple isoforms, 34074 reads/52880348 bases/190710 exons** | | | | | |
| GMAP | -f samse --cross-species -z sense_force | 75.49 | 53.51 | 39.6 | 21.3 |
| Graphmap2 | -x rnaseq | 83.08 | 79.66 | 82.89 | 51.02 |
| Minimap2 | -ax splice | 84.9 | 84.77 | 75.57 | 50.99 |
| Minimap2 + GTF | -ax splice –junc-bed anno.bed | 85.54 | 87.33 | 77.79 | 54.55 |
| deSALT | -d 10 -x clr -s 2 | 87.23 | 91.84 | 89.65 | 69.07 |
| deSALT + GTF | -d 10 -x clr -s 2 -G anno.info | 87.26 | 92.62 | 90.59 | 71.82 |
| **Mouse PacBio subread, coverage = 10X, reads from the genes with single isoform, 27315 reads/31993814 bases/60557 exons** | | | | | |
| GMAP | -f samse --cross-species -z sense_force | 68.87 | 44.16 | 50.78 | 31.73 |
| Graphmap2 | -x rnaseq | 83.13 | 75.01 | 90.12 | 65.57 |
| Minimap2 | -ax splice | 83.59 | 77.42 | 84.42 | 63.76 |
| Minimap2 + GTF | -ax splice –junc-bed anno.bed | 83.77 | 79.39 | 85.63 | 65.62 |

| | | | | | |
|---|---|---|---|---|---|
| deSALT | -d 10 -x clr -s 2 | 84.11 | 82.9 | 90.82 | 72.04 |
| deSALT + GTF | -d 10 -x clr -s 2 -G anno.info | 84.22 | 85.38 | 92.16 | 75.97 |
| **Mouse PacBio subread, coverage = 10X, reads from the genes with multiple isoforms, 75118 reads/131809879 bases/465578 exons** | | | | | |
| GMAP | -f samse --cross-species -z sense_force | 75.49 | 53.64 | 37.47 | 19.39 |
| Graphmap2 | -x rnaseq | 83.24 | 81.82 | 84.89 | 53.03 |
| Minimap2 | -ax splice | 85.56 | 87.09 | 84.42 | 56.79 |
| Minimap2 + GTF | -ax splice –junc-bed anno.bed | 86.23 | 89.82 | 86.99 | 60.96 |
| deSALT | -d 10 -x clr -s 2 | 87.57 | 93.81 | 93.65 | 74.65 |
| deSALT + GTF | -d 10 -x clr -s 2 -G anno.info | 87.58 | 94.18 | 94.07 | 76.12 |
| **Mouse PacBio subread, coverage = 30X, reads from the genes with single isoform, 77032 reads/95918127 bases/176769 exons** | | | | | |
| GMAP | -f samse --cross-species -z sense_force | 69.45 | 44.82 | 50.49 | 32.01 |
| Graphmap2 | -x rnaseq | 83.29 | 77.24 | 91.47 | 67.9 |
| Minimap2 | -ax splice | 84.15 | 79.81 | 88.74 | 67 |
| Minimap2 + GTF | -ax splice –junc-bed anno.bed | 84.31 | 81.71 | 89.93 | 68.96 |
| deSALT | -d 10 -x clr -s 2 | 84.3 | 85.61 | 93.17 | 76 |
| deSALT + GTF | -d 10 -x clr -s 2 -G anno.info | 84.37 | 86.74 | 93.45 | 77.91 |
| **Mouse PacBio subread, coverage = 30X, reads from the genes with multiple isoforms, 212189 reads/395360408 bases/1383094 exons** | | | | | |
| GMAP | -f samse --cross-species -z sense_force | 75.98 | 54.41 | 37.35 | 18.9 |
| Graphmap2 | -x rnaseq | 83.25 | 82.41 | 85.49 | 53.9 |
| Minimap2 | -ax splice | 85.82 | 88.17 | 87.71 | 58.78 |
| Minimap2 + GTF | -ax splice –junc-bed anno.bed | 86.45 | 90.77 | 90.38 | 63.13 |
| deSALT | -d 10 -x clr -s 2 | 87.63 | 94.51 | 94.97 | 76.34 |
| deSALT + GTF | -d 10 -x clr -s 2 -G anno.info | 87.64 | 94.82 | 95.27 | 77.6 |
| **Human PacBio subread, coverage = 4X, reads from the genes with single isoform, 11985 reads/10043761 bases/21582 exons** | | | | | |
| GMAP | -f samse --cross-species -z sense_force | 66.55 | 39.39 | 48.94 | 33.08 |
| Graphmap2 | -x rnaseq | 85.24 | 73.77 | 89.19 | 68.12 |
| Minimap2 | -ax splice | 83.2 | 68.72 | 72.33 | 55.94 |
| Minimap2 + GTF | -ax splice –junc-bed anno.bed | 83.28 | 71.37 | 76.1 | 59.38 |
| deSALT | -d 10 -x clr -s 2 | 85.81 | 76.5 | 85.95 | 66.25 |
| deSALT + GTF | -d 10 -x clr -s 2 -G anno.info | 86.2 | 82.11 | 88.99 | 72.66 |
| **Human PacBio subread, coverage = 4X, reads from the genes having multiple isoforms, 34552 reads/46766039 bases/186837 exons** | | | | | |
| GMAP | -f samse --cross-species -z sense_force | 72.48 | 50.69 | 36.59 | 19.29 |

| | | | | | |
|---|---|---|---|---|---|
| Graphmap2 | -x rnaseq | 83.48 | 80.2 | 83.64 | 51.4 |
| Minimap2 | -ax splice | 84.52 | 84.26 | 74.29 | 49.44 |
| Minimap2 + GTF | -ax splice –junc-bed anno.bed | 85.07 | 85.62 | 80.34 | 54.06 |
| deSALT | -d 10 -x clr -s 2 | 87.23 | 91.61 | 89.03 | 68.78 |
| deSALT + GTF | -d 10 -x clr -s 2 -G anno.info | 87.32 | 92.39 | 90.13 | 71.12 |
| **Human PacBio subread, coverage = 10X, reads from the genes with single isoform, 26391 reads/24880108 bases/50077 exons** | | | | | |
| GMAP | -f samse --cross-species -z sense_force | 66.51 | 39.33 | 48.04 | 31.75 |
| Graphmap2 | -x rnaseq | 85.52 | 76.2 | 91.81 | 69.45 |
| Minimap2 | -ax splice | 84.5 | 74.4 | 82.88 | 63.16 |
| Minimap2 + GTF | -ax splice –junc-bed anno.bed | 84.65 | 76.46 | 85.33 | 66.25 |
| deSALT | -d 10 -x clr -s 2 | 86.85 | 83.49 | 92.34 | 74.29 |
| deSALT + GTF | -d 10 -x clr -s 2 -G anno.info | 87.04 | 86.82 | 93.88 | 78.59 |
| **Human PacBio subread, coverage = 10X, reads from the genes with multiple isoforms, 76142 reads/116446237 bases/454874 exons** | | | | | |
| GMAP | -f samse --cross-species -z sense_force | 72.53 | 50.94 | 34.07 | 17.53 |
| Graphmap2 | -x rnaseq | 83.58 | 81.82 | 85.24 | 52.97 |
| Minimap2 | -ax splice | 85.26 | 86.88 | 83.81 | 55.47 |
| Minimap2 + GTF | -ax splice –junc-bed anno.bed | 85.65 | 87.5 | 86.91 | 57.8 |
| deSALT | -d 10 -x clr -s 2 | 87.73 | 93.87 | 93.82 | 74.29 |
| deSALT + GTF | -d 10 -x clr -s 2 -G anno.info | 87.77 | 94.33 | 94.48 | 75.94 |
| **Human PacBio subread, coverage = 30X, reads from the genes with single isoform, 74399 reads/74561310 bases/145347 exons** | | | | | |
| GMAP | -f samse --cross-species -z sense_force | 66.68 | 39.48 | 47.43 | 31.89 |
| Graphmap2 | -x rnaseq | 85.58 | 78.09 | 93.03 | 71.06 |
| Minimap2 | -ax splice | 85.04 | 77.14 | 87.65 | 67.14 |
| Minimap2 + GTF | -ax splice –junc-bed anno.bed | 84.97 | 78.46 | 89.06 | 68.71 |
| deSALT | -d 10 -x clr -s 2 | 87.23 | 86.91 | 95.07 | 78.91 |
| deSALT + GTF | -d 10 -x clr -s 2 -G anno.info | 87.33 | 88.89 | 95.85 | 81.72 |
| **Human PacBio subread, coverage = 30X, reads from the genes with multiple isoforms, 214976 reads/349280580 bases/1351629 exons** | | | | | |
| GMAP | -f samse --cross-species -z sense_force | 72.84 | 51.44 | 33.45 | 16.96 |
| Graphmap2 | -x rnaseq | 83.64 | 82.83 | 85.63 | 53.68 |
| Minimap2 | -ax splice | 85.57 | 87.97 | 87.55 | 57.7 |
| Minimap2 + GTF | -ax splice –junc-bed anno.bed | 85.83 | 88.48 | 89.8 | 59.96 |
| deSALT | -d 10 -x clr -s 2 | 87.87 | 94.6 | 95.5 | 76.16 |

| | | | | | |
|---|---|---|---|---|---|
| deSALT + GTF | -d 10 -x clr -s 2 -G anno.info | 87.89 | 94.99 | 95.97 | 77.72 |
| **Human all protein coding genes PacBio subread, reads from the genes with single isoform, 108426 reads/ 185904069 bases/ 347355 exons** | | | | | |
| GMAP | -f samse --cross-species -z sense_force | 78.81 | 54.82 | 62.25 | 39.09 |
| Graphmap2 | -x rnaseq | 85.86 | 78.18 | 89.17 | 65.08 |
| Minimap2 | -ax splice | 87.65 | 86.98 | 93.83 | 71.78 |
| Minimap2 + GTF | -ax splice –junc-bed anno.bed | 87.75 | 88.05 | 94.44 | 72.95 |
| deSALT | -d 10 -x clr -s 2 | 87.69 | 90.17 | 94.39 | 78.2 |
| deSALT + GTF | -d 10 -x clr -s 2 -G anno.info | 87.65 | 90.3 | 94.41 | 78.65 |
| **Human all protein coding genes PacBio subread, reads from the genes with multiple isoforms, 1281924 reads/ 2230091277 bases/ 7809546 exons** | | | | | |
| GMAP | -f samse --cross-species -z sense_force | 77.24 | 55.06 | 43.66 | 22.73 |
| Graphmap2 | -x rnaseq | 84.76 | 80.12 | 83.07 | 49.79 |
| Minimap2 | -ax splice | 87.21 | 88.36 | 90.25 | 59.81 |
| Minimap2 + GTF | -ax splice –junc-bed anno.bed | 87.55 | 90.2 | 91.9 | 62.85 |
| deSALT | -d 10 -x clr -s 2 | 88.82 | 94.3 | 96.12 | 75.64 |
| deSALT + GTF | -d 10 -x clr -s 2 -G anno.info | 88.83 | 94.83 | 96.49 | 77.45 |
| **Simulated ONT 1D datasets** | | | | | |
| **Fruit fly ONT 1D, coverage = 4X, reads from the genes with single isoform, 14293 reads/15779696 bases/35183 exons** | | | | | |
| GMAP | -f samse --cross-species -z sense_force | 55 | 16.79 | 21.24 | 5.37 |
| Graphmap2 | -x rnaseq | 93.45 | 64.99 | 80.65 | 39.89 |
| Minimap2 | -ax splice | 77.7 | 53.85 | 52.75 | 29.2 |
| Minimap2 + GTF | -ax splice –junc-bed anno.bed | 78.2 | 59.01 | 56.5 | 35.37 |
| deSALT | -d 10 -x ont1d -s 2 -l 14 -s 2 | 91.41 | 64.14 | 74.17 | 37.98 |
| deSALT + GTF | -d 10 -x ont1d -s 2 -l 14 -s 2 -G anno.info | 92.5 | 74.83 | 83.75 | 52.65 |
| **Fruit fly ONT 1D, coverage = 4X, reads from the genes with multiple isoforms, 30253 reads/86062596 bases/205722 exons** | | | | | |
| GMAP | -f samse --cross-species -z sense_force | 66.52 | 20.99 | 9.96 | 2.47 |
| Graphmap2 | -x rnaseq | 94.37 | 80.56 | 78.53 | 38.28 |
| Minimap2 | -ax splice | 91.75 | 78.67 | 69.62 | 35.85 |
| Minimap2 + GTF | -ax splice –junc-bed anno.bed | 92.14 | 84.62 | 77.23 | 48.6 |
| deSALT | -d 10 -x ont1d -s 2 -l 14 -s 2 | 96.3 | 88.44 | 86.37 | 51.41 |
| deSALT + GTF | -d 10 -x ont1d -s 2 -l 14 -s 2 -G anno.info | 96.57 | 91.8 | 92.05 | 60.72 |

| | | | | | |
|---|---|---|---|---|---|
| **Fruit fly ONT 1D, coverage = 10X, reads from the genes with single isoforms, 31603 reads/39450304 bases/83839 exons** | | | | | |
| GMAP | -f samse --cross-species -z sense_force | 56.83 | 17.69 | 22.26 | 5.5 |
| Graphmap2 | -x rnaseq | 94.02 | 73.18 | 86.9 | 46.97 |
| Minimap2 | -ax splice | 80.44 | 58.74 | 59.48 | 33.02 |
| Minimap2 + GTF | -ax splice –junc-bed anno.bed | 80.99 | 64.55 | 63.95 | 40.66 |
| deSALT | -d 10 -x ont1d -s 2 -l 14 -s 2 | 93.49 | 72.62 | 81 | 45.71 |
| deSALT + GTF | -d 10 -x ont1d -s 2 -l 14 -s 2 -G anno.info | 94.14 | 83.44 | 90.37 | 64.02 |
| **Fruit fly ONT 1D, coverage = 10X, reads from the genes with multiple isoforms, 68524 reads/215158359 bases/505777 exons** | | | | | |
| GMAP | -f samse --cross-species -z sense_force | 67.63 | 21.38 | 9.1 | 2.21 |
| Graphmap2 | -x rnaseq | 94.46 | 83.36 | 82.46 | 42.67 |
| Minimap2 | -ax splice | 92.91 | 81.16 | 74.58 | 38.69 |
| Minimap2 + GTF | -ax splice –junc-bed anno.bed | 93.3 | 87.19 | 82.65 | 52.7 |
| deSALT | -d 10 -x ont1d -s 2 -l 14 -s 2 | 96.75 | 91.86 | 91.23 | 60.02 |
| deSALT + GTF | -d 10 -x ont1d -s 2 -l 14 -s 2 -G anno.info | 96.9 | 94.02 | 94.63 | 67.75 |
| **Fruit fly ONT 1D, coverage = 30X, reads from the genes with single isoform, 94542 reads/118339414 bases/251194 exons** | | | | | |
| GMAP | -f samse --cross-species -z sense_force | 56.91 | 17.82 | 22.36 | 5.35 |
| Graphmap2 | -x rnaseq | 94.05 | 75.68 | 88.67 | 50.43 |
| Minimap2 | -ax splice | 80.52 | 58.79 | 59.47 | 33.02 |
| Minimap2 + GTF | -ax splice –junc-bed anno.bed | 81.06 | 64.7 | 64.01 | 40.56 |
| deSALT | -d 10 -x ont1d -s 2 -l 14 -s 2 | 94.16 | 79.38 | 87.37 | 55.79 |
| deSALT + GTF | -d 10 -x ont1d -s 2 -l 14 -s 2 -G anno.info | 94.44 | 85.64 | 91.65 | 68.08 |
| **Fruit fly ONT 1D, coverage = 30X, reads from the genes with multiple isoforms, 203279 reads/645463167 bases/1512998 exons** | | | | | |
| GMAP | -f samse --cross-species -z sense_force | 67.68 | 21.43 | 8.99 | 2.2 |
| Graphmap2 | -x rnaseq | 94.47 | 84.16 | 83.34 | 44.92 |
| Minimap2 | -ax splice | 92.98 | 81.22 | 74.84 | 39.01 |
| Minimap2 + GTF | -ax splice –junc-bed anno.bed | 93.37 | 87.32 | 82.94 | 53.07 |
| deSALT | -d 10 -x ont1d -s 2 -l 14 -s 2 | 96.84 | 92.86 | 93.09 | 64.64 |
| deSALT + GTF | -d 10 -x ont1d -s 2 -l 14 -s 2 -G anno.info | 96.94 | 94.49 | 95.24 | 70.09 |
| **Mouse ONT 1D, coverage = 4X, reads from the genes with single isoform, 12411 reads/12792340 bases/27910 exons** | | | | | |
| GMAP | -f samse --cross-species -z sense_force | 53.16 | 10.43 | 25.52 | 4.99 |
| Graphmap2 | -x rnaseq | 83.64 | 38.28 | 66.01 | 26.23 |
| Minimap2 | -ax splice | 71.45 | 36.85 | 41.33 | 17.86 |

| | | | | | |
|---|---|---|---|---|---|
| Minimap2 + GTF | -ax splice –junc-bed anno.bed | 71.73 | 39.34 | 42.82 | 19.38 |
| deSALT | -d 10 -x ont1d -s 2 -l 14 -s 2 | 80.42 | 40.42 | 56.36 | 22.19 |
| deSALT + GTF | -d 10 -x ont1d -s 2 -l 14 -s 2 -G anno.info | 82.18 | 53.6 | 64.66 | 35.3 |
| **Mouse ONT 1D, coverage = 4X, reads from the genes with multiple isoforms, 34189 reads/52716526 bases/215979 exons** | | | | | |
| GMAP | -f samse --cross-species -z sense_force | 53.78 | 13.14 | 8.58 | 2.09 |
| Graphmap2 | -x rnaseq | 78.58 | 49.09 | 41.2 | 15.7 |
| Minimap2 | -ax splice | 76.26 | 54.58 | 38.36 | 16.37 |
| Minimap2 + GTF | -ax splice –junc-bed anno.bed | 77.5 | 60.36 | 43.25 | 21.12 |
| deSALT | -d 10 -x ont1d -s 2 -l 14 -s 2 | 88.2 | 69.15 | 57.84 | 26.61 |
| deSALT + GTF | -d 10 -x ont1d -s 2 -l 14 -s 2 -G anno.info | 89.47 | 77.03 | 69.69 | 37.28 |
| **Mouse ONT 1D, coverage = 10X, reads from the genes with single isoform, 27474 reads/31977856 bases/66240 exons** | | | | | |
| GMAP | -f samse --cross-species -z sense_force | 54.18 | 10.91 | 25.96 | 5.25 |
| Graphmap2 | -x rnaseq | 84.46 | 43.51 | 70.06 | 28.65 |
| Minimap2 | -ax splice | 73.68 | 40.19 | 46.08 | 20.29 |
| Minimap2 + GTF | -ax splice –junc-bed anno.bed | 74.01 | 43.16 | 47.89 | 22.09 |
| deSALT | -d 10 -x ont1d -s 2 -l 14 -s 2 | 84.28 | 50.1 | 63.62 | 26.97 |
| deSALT + GTF | -d 10 -x ont1d -s 2 -l 14 -s 2 -G anno.info | 85.52 | 65.68 | 74.86 | 45.52 |
| **Mouse ONT 1D, coverage = 10X, reads from the genes with multiple isoforms, 76173 reads/131797813 bases/529496 exons** | | | | | |
| GMAP | -f samse --cross-species -z sense_force | 55.05 | 13.84 | 7.43 | 1.73 |
| Graphmap2 | -x rnaseq | 78.88 | 52.46 | 43.49 | 17.44 |
| Minimap2 | -ax splice | 77.85 | 57.16 | 41.66 | 18.08 |
| Minimap2 + GTF | -ax splice –junc-bed anno.bed | 79.12 | 63.27 | 47.43 | 23.61 |
| deSALT | -d 10 -x ont1d -s 2 -l 14 -s 2 | 90.42 | 77.23 | 69.24 | 36.87 |
| deSALT + GTF | -d 10 -x ont1d -s 2 -l 14 -s 2 -G anno.info | 91.06 | 81.91 | 76.48 | 45.85 |
| **Mouse ONT 1D, coverage = 30X, reads from the genes with single isoform, 82162 reads/95919067 bases/198286 exons** | | | | | |
| GMAP | -f samse --cross-species -z sense_force | 52.78 | 10.46 | 24.45 | 4.84 |
| Graphmap2 | -x rnaseq | 84.26 | 45.41 | 71.04 | 29.65 |
| Minimap2 | -ax splice | 73.62 | 40.46 | 46 | 20.46 |
| Minimap2 + GTF | -ax splice –junc-bed anno.bed | 57.22 | 43.34 | 47.78 | 22.26 |
| deSALT | -d 10 -x ont1d -s 2 -l 14 -s 2 | 85.43 | 59.81 | 70.44 | 36.03 |
| deSALT + GTF | -d 10 -x ont1d -s 2 -l 14 -s 2 -G anno.info | 85.99 | 69.95 | 76.83 | 51.43 |
| **Mouse ONT 1D, coverage = 30X, reads from the genes with multiple isoforms, 227114 reads/395363051 bases/1585095 exons** | | | | | |

| | | | | | |
|---|---|---|---|---|---|
| GMAP | -f samse --cross-species -z sense_force | 31.25 | 7.85 | 4.34 | 1 |
| Graphmap2 | -x rnaseq | 79.01 | 53.82 | 45.1 | 18.81 |
| Minimap2 | -ax splice | 77.98 | 57.42 | 41.87 | 18.3 |
| Minimap2 + GTF | -ax splice –junc-bed anno.bed | 79.23 | 63.54 | 47.61 | 23.81 |
| deSALT | -d 10 -x ont1d -s 2 -l 14 -s 2 | 90.8 | 79.71 | 73.45 | 42.78 |
| deSALT + GTF | -d 10 -x ont1d -s 2 -l 14 -s 2 -G anno.info | 91.18 | 82.5 | 77.32 | 48.21 |
| **Human ONT 1D, coverage = 4X, reads from the genes with single isoform, 11997 reads/9944834 bases/23000 exons** | | | | | |
| GMAP | -f samse --cross-species -z sense_force | 49.37 | 8.23 | 22.53 | 4.86 |
| Graphmap2 | -x rnaseq | 84.59 | 35.78 | 65.19 | 26.43 |
| Minimap2 | -ax splice | 67.38 | 27 | 35.75 | 15.57 |
| Minimap2 + GTF | -ax splice –junc-bed anno.bed | 67.6 | 29.16 | 36.85 | 16.51 |
| deSALT | -d 10 -x ont1d -s 2 -l 14 -s 2 | 79.56 | 33.02 | 53.16 | 20.81 |
| deSALT + GTF | -d 10 -x ont1d -s 2 -l 14 -s 2 -G anno.info | 81.77 | 46.52 | 61.08 | 33.38 |
| **Human ONT 1D, coverage = 4X, reads from the genes with multiple isoforms, 34647 reads/46572365 bases/210621 exons** | | | | | |
| GMAP | -f samse --cross-species -z sense_force | 48.99 | 11.88 | 6.46 | 1.68 |
| Graphmap2 | -x rnaseq | 73.9 | 45.23 | 36.99 | 14.12 |
| Minimap2 | -ax splice | 71.38 | 49.6 | 33.18 | 14.13 |
| Minimap2 + GTF | -ax splice –junc-bed anno.bed | 72.79 | 55.33 | 37.33 | 18.4 |
| deSALT | -d 10 -x ont1d -s 2 -l 14 -s 2 | 86.35 | 67.76 | 55.98 | 26.78 |
| deSALT + GTF | -d 10 -x ont1d -s 2 -l 14 -s 2 -G anno.info | 87.68 | 74.87 | 66.54 | 35.92 |
| **Human ONT 1D, coverage = 10X, reads from the genes with single isoform, 26530 reads/24859119 bases/53998 exons** | | | | | |
| GMAP | -f samse --cross-species -z sense_force | 49.99 | 8.82 | 23.85 | 5.18 |
| Graphmap2 | -x rnaseq | 85.55 | 39.89 | 69.08 | 28.76 |
| Minimap2 | -ax splice | 69.61 | 30.46 | 40.37 | 18.11 |
| Minimap2 + GTF | -ax splice –junc-bed anno.bed | 69.85 | 32.86 | 41.75 | 19.19 |
| deSALT | -d 10 -x ont1d -s 2 -l 14 -s 2 | 84.04 | 41.56 | 60.32 | 25.42 |
| deSALT + GTF | -d 10 -x ont1d -s 2 -l 14 -s 2 -G anno.info | 85.63 | 59.81 | 71.78 | 44.71 |
| **Human ONT 1D, coverage = 10X, reads from the genes with multiple isoforms, 77002 reads/116437109 bases/515990 exons** | | | | | |
| GMAP | -f samse --cross-species -z sense_force | 49.9 | 12.47 | 5.46 | 1.41 |
| Graphmap2 | -x rnaseq | 74.31 | 47.94 | 38.81 | 15.22 |
| Minimap2 | -ax splice | 73.06 | 52.42 | 36.46 | 15.7 |
| Minimap2 + GTF | -ax splice –junc-bed anno.bed | 74.49 | 58.28 | 41.58 | 20.64 |

| | | | | | |
|---|---|---|---|---|---|
| deSALT | -d 10 -x ont1d -s 2 -l 14 -s 2 | 88.52 | 75.06 | 66.65 | 35.6 |
| deSALT + GTF | -d 10 -x ont1d -s 2 -l 14 -s 2 -G anno.info | 89.22 | 79.38 | 73.13 | 42.99 |
| **Human ONT 1D, coverage = 30X, reads from the genes with single isoform, 79328 reads/74561914 bases/161631 exons** | | | | | |
| GMAP | -f samse --cross-species -z sense_force | 51 | 9.01 | 24.12 | 5.08 |
| Graphmap2 | -x rnaseq | 85.69 | 41.51 | 70.08 | 29.8 |
| Minimap2 | -ax splice | 69.62 | 29.94 | 40.1 | 17.9 |
| Minimap2 + GTF | -ax splice –junc-bed anno.bed | 69.86 | 32.34 | 41.43 | 19.05 |
| deSALT | -d 10 -x ont1d -s 2 -l 14 -s 2 | 85.5 | 51.65 | 66.98 | 33.7 |
| deSALT + GTF | -d 10 -x ont1d -s 2 -l 14 -s 2 -G anno.info | 86.26 | 64.53 | 73.98 | 49.99 |
| **Human ONT 1D, coverage = 30X, reads from the genes with multiple isoforms, 229807 reads/349281664 bases/1544708 exons** | | | | | |
| GMAP | -f samse --cross-species -z sense_force | 49.93 | 12.37 | 5.39 | 1.42 |
| Graphmap2 | -x rnaseq | 74.52 | 48.97 | 39.21 | 16.12 |
| Minimap2 | -ax splice | 73.24 | 52.62 | 36.66 | 15.88 |
| Minimap2 + GTF | -ax splice –junc-bed anno.bed | 74.66 | 58.5 | 41.89 | 20.82 |
| deSALT | -d 10 -x ont1d -s 2 -l 14 -s 2 | 88.98 | 76.91 | 69.76 | 40.31 |
| deSALT + GTF | -d 10 -x ont1d -s 2 -l 14 -s 2 -G anno.info | 89.44 | 79.81 | 73.61 | 44.93 |
| **Human all protein coding genes ONT 1D, reads from the genes with single isoform, 91821 reads/ 185871767 bases/ 371073 exons** | | | | | |
| GMAP | -f samse --cross-species -z sense_force | 61.38 | 11.96 | 26.56 | 4.26 |
| Graphmap2 | -x rnaseq | 91.02 | 46.49 | 63.09 | 26.13 |
| Minimap2 | -ax splice | 88.23 | 54.11 | 58.26 | 26.59 |
| Minimap2 + GTF | -ax splice –junc-bed anno.bed | 88.74 | 58.6 | 61.06 | 29.57 |
| deSALT | -d 10 -x ont1d -s 2 -l 14 | 92.28 | 62.15 | 66.94 | 30.87 |
| deSALT + GTF | -d 10 -x ont1d -s 2 -l 14 -G anno.info | 92.36 | 65.12 | 68.64 | 34.09 |
| **Human all protein coding genes ONT 1D, reads from the genes with multiple isoforms, 1034507 reads/ 2228456014 bases/ 8513080 exons** | | | | | |
| GMAP | -f samse --cross-species -z sense_force | 53.42 | 11.58 | 5.28 | 1.11 |
| Graphmap2 | -x rnaseq | 81.64 | 45.98 | 34.41 | 13.17 |
| Minimap2 | -ax splice | 84.69 | 57.99 | 42.01 | 18.23 |
| Minimap2 + GTF | -ax splice –junc-bed anno.bed | 86.17 | 64.7 | 48.45 | 24.45 |
| deSALT | -d 10 -x ont1d -s 2 -l 14 | 92.14 | 67.82 | 55.14 | 24.52 |
| deSALT + GTF | -d 10 -x ont1d -s 2 -l 14 -G anno.info | 92.65 | 72.56 | 59.61 | 31.61 |
| **Simulated datasets by PBSim based on a real ONT dataset (PS-ONT)** | | | | | |

| | | | | | |
|---|---|---|---|---|---|
| **Fruit fly PS-ONT, coverage = 4X, reads from the genes with single isoform, 14395 reads/ 15809329 bases/33628 exons** | | | | | |
| GMAP | -f samse --cross-species -z sense_force | 96.08 | 86.78 | 92.23 | 74.82 |
| Graphmap2 | -x rnaseq | 92.55 | 94.75 | 98.46 | 89.9 |
| Minimap2 | -ax splice | 91.82 | 90.76 | 89.61 | 81.27 |
| Minimap2 + GTF | -ax splice –junc-bed anno.bed | 91.88 | 91.62 | 89.84 | 82.87 |
| deSALT | -d 10 | 92.28 | 93.53 | 94.41 | 86.95 |
| deSALT + GTF | -d 10 -G anno.info | 92.33 | 94.26 | 94.53 | 88.46 |
| **Fruit fly PS-ONT, coverage = 4X, reads from the genes with multiple isoforms, 31910 reads/86073991 bases/193942 exons** | | | | | |
| GMAP | -f samse --cross-species -z sense_force | 93.75 | 89.84 | 91.72 | 59.8 |
| Graphmap2 | -x rnaseq | 93.41 | 95.84 | 98.19 | 82.7 |
| Minimap2 | -ax splice | 97.27 | 95.48 | 95.97 | 78.9 |
| Minimap2 + GTF | -ax splice –junc-bed anno.bed | 93.78 | 96.93 | 96.79 | 83.64 |
| deSALT | -d 10 | 93.87 | 97.39 | 98.14 | 86.58 |
| deSALT + GTF | -d 10 -G anno.info | 93.87 | 97.87 | 98.37 | 88.48 |
| **Fruit fly PS-ONT, coverage = 10X, reads from the genes with single isoforms, 31927 reads/ 39450414 bases/ 80177 exons** | | | | | |
| GMAP | -f samse --cross-species -z sense_force | 92.24 | 87.96 | 94.4 | 75.77 |
| Graphmap2 | -x rnaseq | 92.73 | 95.36 | 98.9 | 90.48 |
| Minimap2 | -ax splice | 92.66 | 94.47 | 97.32 | 88.1 |
| Minimap2 + GTF | -ax splice –junc-bed anno.bed | 92.72 | 95.33 | 97.56 | 89.81 |
| deSALT | -d 10 | 92.66 | 96.05 | 97.61 | 91.63 |
| deSALT + GTF | -d 10 -G anno.info | 92.67 | 96.16 | 97.63 | 91.91 |
| **Fruit fly PS-ONT, coverage = 10X, reads from the genes with multiple isoforms, 72848 reads/ 215159929 bases/ 477080 exons** | | | | | |
| GMAP | -f samse --cross-species -z sense_force | 93.8 | 90.17 | 92.18 | 59.05 |
| Graphmap2 | -x rnaseq | 93.36 | 96.0 | 98.2 | 82.86 |
| Minimap2 | -ax splice | 93.86 | 96.17 | 98.21 | 80.73 |
| Minimap2 + GTF | -ax splice –junc-bed anno.bed | 93.9 | 97.6 | 99.03 | 85.75 |
| deSALT | -d 10 | 93.9 | 98.02 | 99.14 | 88.76 |
| deSALT + GTF | -d 10 -G anno.info | 93.9 | 98.32 | 99.28 | 90.02 |
| **Fruit fly PS-ONT, coverage = 30X, reads from the genes with single isoform, 92478 reads/ 118404601 bases/ 236981 exons** | | | | | |
| GMAP | -f samse --cross-species -z sense_force | 92.36 | 88.3 | 94.57 | 76.27 |
| Graphmap2 | -x rnaseq | 92.7 | 95.86 | 98.96 | 91.5 |
| Minimap2 | -ax splice | 92.66 | 94.71 | 97.43 | 88.39 |

| | | | | | |
|---|---|---|---|---|---|
| Minimap2 + GTF | -ax splice –junc-bed anno.bed | 92.72 | 95.58 | 97.69 | 90.15 |
| deSALT | -d 10 | 92.7 | 96.56 | 97.68 | 92.73 |
| deSALT + GTF | -d 10 -G anno.info | 92.7 | 96.58 | 97.71 | 92.76 |
| **Fruit fly PS-ONT, coverage = 30X, reads from the genes with multiple isoforms, 212032 reads/ 645503059 bases/ 1423919 exons** | | | | | |
| GMAP | -f samse --cross-species -z sense_force | 93.84 | 90.28 | 92.27 | 58.71 |
| Graphmap2 | -x rnaseq | 93.36 | 96.1 | 98.23 | 83.08 |
| Minimap2 | -ax splice | 93.9 | 96.28 | 98.3 | 80.9 |
| Minimap2 + GTF | -ax splice –junc-bed anno.bed | 93.94 | 97.72 | 99.12 | 86.04 |
| deSALT | -d 10 | 93.94 | 98.18 | 99.23 | 89.38 |
| deSALT + GTF | -d 10 -G anno.info | 9394 | 98.43 | 99.34 | 90.41 |
| **Mouse PS-ONT, coverage = 4X, reads from the genes with single isoform, 12508 reads/ 12833801 bases/26791 exons** | | | | | |
| GMAP | -f samse --cross-species -z sense_force | 88.25 | 79.08 | 87.43 | 69.05 |
| Graphmap2 | -x rnaseq | 88.43 | 84.35 | 92.98 | 80.94 |
| Minimap2 | -ax splice | 89.76 | 84.63 | 84.79 | 74.91 |
| Minimap2 + GTF | -ax splice –junc-bed anno.bed | 89.84 | 85.4 | 85.19 | 75.67 |
| deSALT | -d 10 | 88.4 | 85.36 | 88.42 | 78.22 |
| deSALT + GTF | -d 10 -G anno.info | 88.53 | 86.64 | 88.78 | 80.09 |
| **Mouse PS-ONT, coverage = 4X, reads from the genes with multiple isoforms, 34840 reads/ 52785589 bases/202234 exons** | | | | | |
| GMAP | -f samse --cross-species -z sense_force | 91.27 | 85.33 | 83.71 | 54.26 |
| Graphmap2 | -x rnaseq | 88.63 | 87.29 | 90.89 | 69.04 |
| Minimap2 | -ax splice | 92.03 | 91.42 | 87.45 | 66.85 |
| Minimap2 + GTF | -ax splice –junc-bed anno.bed | 92.07 | 92.35 | 88.21 | 68.79 |
| deSALT | -d 10 | 92.55 | 94.53 | 92.92 | 78.25 |
| deSALT + GTF | -d 10 -G anno.info | 92.63 | 95.01 | 93.21 | 79.62 |
| **Mouse PS-ONT, coverage = 10X, reads from the genes with single isoform, 27787 reads/ 31977891 bases/63438 exons** | | | | | |
| GMAP | -f samse --cross-species -z sense_force | 88.53 | 80.3 | 89.37 | 69.88 |
| Graphmap2 | -x rnaseq | 88.69 | 85.3 | 94.02 | 81.69 |
| Minimap2 | -ax splice | 90.64 | 89.07 | 93.62 | 82.26 |
| Minimap2 + GTF | -ax splice –junc-bed anno.bed | 90.7 | 89.88 | 94.09 | 83.12 |
| deSALT | -d 10 | 89.14 | 89.52 | 92.6 | 84.87 |
| deSALT + GTF | -d 10 -G anno.info | 89.18 | 90.02 | 92.71 | 85.78 |
| **Mouse PS-ONT, coverage = 10X, reads from the genes with multiple isoforms, 77930 reads/ 131794166 bases/ 495760 exons** | | | | | |

| | | | | | |
|---|---|---|---|---|---|
| GMAP | -f samse --cross-species -z sense_force | 91.43 | 85.95 | 84.85 | 53.34 |
| Graphmap2 | -x rnaseq | 88.79 | 87.85 | 91.36 | 69.56 |
| Minimap2 | -ax splice | 92.57 | 93.32 | 94.38 | 71.9 |
| Minimap2 + GTF | -ax splice –junc-bed anno.bed | 98.13 | 94.25 | 95.16 | 74.06 |
| deSALT | -d 10 | 92.92 | 95.91 | 96.35 | 82.38 |
| deSALT + GTF | -d 10 -G anno.info | 92.93 | 96.21 | 96.56 | 83.31 |
| **Mouse PS-ONT, coverage = 30X, reads from the genes with single isoform, 80361 reads/ 95992475 bases/ 187360 exons** | | | | | |
| GMAP | -f samse --cross-species -z sense_force | 88.63 | 80.28 | 89.5 | 69.72 |
| Graphmap2 | -x rnaseq | 88.68 | 85.52 | 93.95 | 82.35 |
| Minimap2 | -ax splice | 90.76 | 89.51 | 94.35 | 82.81 |
| Minimap2 + GTF | -ax splice –junc-bed anno.bed | 90.83 | 90.34 | 94.85 | 83.68 |
| deSALT | -d 10 | 89.13 | 90.25 | 92.75 | 85.87 |
| deSALT + GTF | -d 10 -G anno.info | 98.92 | 90.59 | 92.93 | 86.41 |
| **Mouse PS-ONT, coverage = 30X, reads from the genes with multiple isoforms, 225628 reads/ 395509618 bases/ 1478380 exons** | | | | | |
| GMAP | -f samse --cross-species -z sense_force | 91.54 | 86.26 | 84.96 | 53.07 |
| Graphmap2 | -x rnaseq | 88.77 | 88.06 | 91.41 | 69.88 |
| Minimap2 | -ax splice | 92.63 | 93.65 | 94.8 | 72.17 |
| Minimap2 + GTF | -ax splice –junc-bed anno.bed | 92.69 | 94.6 | 95.6 | 74.45 |
| deSALT | -d 10 | 92.95 | 96.22 | 96.56 | 83.15 |
| deSALT + GTF | -d 10 -G anno.info | 92.96 | 96.46 | 96.75 | 83.95 |
| **Human PS-ONT, coverage = 4X, reads from the genes with single isoform, 12052 reads/ 9998219 bases/22226 exons** | | | | | |
| GMAP | -f samse --cross-species -z sense_force | 90.34 | 79.45 | 88.35 | 71.59 |
| Graphmap2 | -x rnaseq | 90.69 | 86.43 | 93.98 | 83.89 |
| Minimap2 | -ax splice | 90.71 | 83.31 | 83.62 | 75.1 |
| Minimap2 + GTF | -ax splice –junc-bed anno.bed | 90.83 | 84.13 | 83.98 | 75.79 |
| deSALT | -d 10 | 91.28 | 87.24 | 90.49 | 81.35 |
| deSALT + GTF | -d 10 -G anno.info | 91.51 | 88.87 | 91.03 | 83.31 |
| **Human PS-ONT, coverage = 4X, reads from the genes with multiple isoforms, 35134 reads/ 46651580 bases/198077 exons** | | | | | |
| GMAP | -f samse --cross-species -z sense_force | 90.94 | 84.74 | 82.8 | 52.52 |
| Graphmap2 | -x rnaseq | 88.35 | 86.77 | 89.85 | 66.78 |
| Minimap2 | -ax splice | 91.86 | 90.83 | 86.03 | 64.42 |
| Minimap2 + GTF | -ax splice –junc-bed anno.bed | 91.9 | 91.54 | 86.64 | 66.14 |

| | | | | | |
|---|---|---|---|---|---|
| deSALT | -d 10 | 92.6 | 94.16 | 92.67 | 76.56 |
| deSALT + GTF | -d 10 -G anno.info | 92.69 | 94.58 | 92.9 | 77.9 |
| **Human PS-ONT, coverage = 10X, reads from the genes with single isoform, 26692 reads/24859142 bases/51944 exons** | | | | | |
| GMAP | -f samse --cross-species -z sense_force | 90.67 | 81.22 | 90.27 | 73.58 |
| Graphmap2 | -x rnaseq | 91.1 | 87.76 | 95.3 | 85.58 |
| Minimap2 | -ax splice | 91.81 | 89.05 | 93.48 | 83.64 |
| Minimap2 + GTF | -ax splice –junc-bed anno.bed | 91.94 | 89.99 | 93.89 | 84.31 |
| deSALT | -d 10 | 92.16 | 92.57 | 95.7 | 89.25 |
| deSALT + GTF | -d 10 -G anno.info | 92.27 | 93.53 | 96.01 | 90.54 |
| **Human PS-ONT, coverage = 10X, reads from the genes with multiple isoforms, 78351 reads/116435702 bases/483615 exons** | | | | | |
| GMAP | -f samse --cross-species -z sense_force | 91.2 | 85.45 | 93.65 | 52.04 |
| Graphmap2 | -x rnaseq | 88.44 | 87.59 | 90.77 | 67.47 |
| Minimap2 | -ax splice | 92.48 | 93.12 | 94.26 | 70.49 |
| Minimap2 + GTF | -ax splice –junc-bed anno.bed | 92.52 | 93.84 | 94.88 | 72.35 |
| deSALT | -d 10 | 92.98 | 95.76 | 96.81 | 81.43 |
| deSALT + GTF | -d 10 -G anno.info | 93.02 | 95.99 | 97.01 | 82.28 |
| **Human PS-ONT, coverage = 30X, reads from the genes with single isoform, 77134 reads/7642422 bases/152764 exons** | | | | | |
| GMAP | -f samse --cross-species -z sense_force | 90.93 | 81.75 | 90.78 | 74.02 |
| Graphmap2 | -x rnaseq | 91.11 | 88.32 | 95.49 | 86.28 |
| Minimap2 | -ax splice | 92.07 | 89.88 | 94.67 | 84.57 |
| Minimap2 + GTF | -ax splice –junc-bed anno.bed | 92.2 | 90.84 | 95.12 | 85.35 |
| deSALT | -d 10 | 92.33 | 93.63 | 96.32 | 90.87 |
| deSALT + GTF | -d 10 -G anno.info | 92.44 | 94.31 | 96.55 | 91.71 |
| **Human PS-ONT, coverage = 30X, reads from the genes with multiple isoforms, 226999 reads/ 349451169 bases/ 1442190 exons** | | | | | |
| GMAP | -f samse --cross-species -z sense_force | 91.31 | 85.79 | 93.89 | 52.03 |
| Graphmap2 | -x rnaseq | 88.42 | 97.72 | 90.64 | 67.67 |
| Minimap2 | -ax splice | 92.59 | 93.46 | 94.63 | 71.07 |
| Minimap2 + GTF | -ax splice –junc-bed anno.bed | 92.64 | 94.21 | 95.32 | 72.98 |
| deSALT | -d 10 | 93.08 | 96.02 | 97.05 | 81.98 |
| deSALT + GTF | -d 10 -G anno.info | 93.12 | 96.3 | 97.29 | 82.82 |
| **Human all protein coding genes PS-ONT, reads from the genes with single isoform, 91843 reads/ 185908018 bases/ 347970 exons** | | | | | |
| GMAP | -f samse --cross-species -z sense_force | 93.62 | 76.17 | 83.33 | 55.4 |

| | | | | | |
|---|---|---|---|---|---|
| Graphmap2 | -x rnaseq | 92.17 | 82.41 | 91.26 | 71.83 |
| Minimap2 | -ax splice | 94.76 | 91.81 | 95.81 | 79.62 |
| Minimap2 + GTF | -ax splice –junc-bed anno.bed | 94.86 | 92.61 | 96.28 | 80.7 |
| deSALT | -d 10 | 94.67 | 93.16 | 95.36 | 83.67 |
| deSALT + GTF | -d 10 -G anno.info | 94.59 | 93.1 | 95.2 | 84.01 |
| **Human all protein coding genes PS-ONT, reads from the genes with multiple isoforms, 1275238 reads/ 2438751277 bases/ 8962242 exons** | | | | | |
| GMAP | -f samse --cross-species -z sense_force | 93.44 | 77.24 | 71.18 | 39.05 |
| Graphmap2 | -x rnaseq | 90.17 | 82.38 | 85.84 | 56.26 |
| Minimap2 | -ax splice | 94.6 | 92.18 | 89.56 | 66.72 |
| Minimap2 + GTF | -ax splice –junc-bed anno.bed | 94.92 | 93.63 | 90.84 | 69.79 |
| deSALT | -d 10 | 95.55 | 95.6 | 95.06 | 78.32 |
| deSALT + GTF | -d 10 -G anno.info | 95.55 | 95.39 | 95.06 | 79.93 |
| **Simulated datasets by NanoSIm based on a real ONT dataset (NS-ONT)** | | | | | |
| **Fruit fly NS-ONT, coverage = 4X, reads from the genes with single isoform, 891 reads/ 3663249 bases/3649 exons** | | | | | |
| GMAP | -f samse --cross-species -z sense_force | 90.42 | 58.92 | 25.81 | 11.34 |
| Graphmap2 | -x rnaseq | 91.29 | 75.69 | 51.07 | 26.49 |
| Minimap2 | -ax splice | 91.01 | 90.08 | 94.84 | 70.59 |
| Minimap2 + GTF | -ax splice –junc-bed anno.bed | 91.06 | 93.26 | 95.4 | 79.35 |
| deSALT | -d 10 | 91.47 | 90.49 | 96.18 | 71.48 |
| deSALT + GTF | -d 10 -G anno.info | 91.51 | 93.01 | 96.63 | 79.01 |
| **Fruit fly NS-ONT, coverage = 4X, reads from the genes with multiple isoforms, 33111 reads/194447357 bases/290110 exons** | | | | | |
| GMAP | -f samse --cross-species -z sense_force | 92.24 | 64.11 | 16.16 | 4.64 |
| Graphmap2 | -x rnaseq | 92.53 | 82.88 | 53.24 | 16.1 |
| Minimap2 | -ax splice | 92.98 | 93.59 | 95.8 | 62.85 |
| Minimap2 + GTF | -ax splice –junc-bed anno.bed | 93.02 | 97.2 | 97.74 | 79.93 |
| deSALT | -d 10 | 92.71 | 95.62 | 97.45 | 71.59 |
| deSALT + GTF | -d 10 -G anno.info | 92.72 | 97.45 | 97.83 | 83.11 |
| **Fruit fly NS-ONT, coverage = 10X, reads from the genes with single isoforms, 2120 reads/ 8651909 bases/ 8860 exons** | | | | | |
| GMAP | -f samse --cross-species -z sense_force | 90.76 | 59.41 | 27.31 | 13.44 |
| Graphmap2 | -x rnaseq | 90.88 | 75.44 | 53.16 | 28.21 |
| Minimap2 | -ax splice | 91.19 | 89.91 | 94.1 | 71.27 |

| | | | | | |
|---|---|---|---|---|---|
| Minimap2 + GTF | -ax splice –junc-bed anno.bed | 91.23 | 92.69 | 94.81 | 78.92 |
| deSALT | -d 10 | 91.45 | 90.79 | 94.95 | 72.97 |
| deSALT + GTF | -d 10 -G anno.info | 91.48 | 92.91 | 95.52 | 79.34 |
| **Fruit fly NS-ONT, coverage = 10X, reads from the genes with multiple isoforms, 83882 reads/ 494914574 bases/ 735880 exons** | | | | | |
| GMAP | -f samse --cross-species -z sense_force | 92.14 | 64.24 | 15.91 | 4.39 |
| Graphmap2 | -x rnaseq | 92.32 | 81.95 | 52.18 | 15.63 |
| Minimap2 | -ax splice | 92.87 | 93.59 | 95.85 | 62.96 |
| Minimap2 + GTF | -ax splice –junc-bed anno.bed | 92.92 | 97.19 | 97.15 | 80.05 |
| deSALT | -d 10 | 92.65 | 95.64 | 97.43 | 72.04 |
| deSALT + GTF | -d 10 -G anno.info | 92.67 | 97.44 | 97.74 | 83.4 |
| **Fruit fly NS-ONT, coverage = 30X, reads from the genes with single isoform, 6045 reads/ 24421609 bases/ 24178 exons** | | | | | |
| GMAP | -f samse --cross-species -z sense_force | 90.53 | 59.14 | 27.2 | 13.53 |
| Graphmap2 | -x rnaseq | 90.69 | 74.37 | 49.93 | 27.66 |
| Minimap2 | -ax splice | 90.82 | 89.35 | 93.85 | 70.19 |
| Minimap2 + GTF | -ax splice –junc-bed anno.bed | 90.86 | 92.32 | 94.64 | 77.57 |
| deSALT | -d 10 | 91.31 | 90.47 | 95.22 | 72.57 |
| deSALT + GTF | -d 10 -G anno.info | 91.15 | 92.54 | 95.65 | 78.41 |
| **Fruit fly NS-ONT, coverage = 30X, reads from the genes with multiple isoforms, 243957 reads/ 1435922298 bases/ 2134817 exons** | | | | | |
| GMAP | -f samse --cross-species -z sense_force | 92.2 | 64.17 | 16.01 | 4.51 |
| Graphmap2 | -x rnaseq | 92.46 | 80.59 | 50.09 | 15.04 |
| Minimap2 | -ax splice | 92.91 | 93.53 | 95.8 | 62.89 |
| Minimap2 + GTF | -ax splice –junc-bed anno.bed | 92.96 | 97.18 | 97.75 | 79.9 |
| deSALT | -d 10 | 92.72 | 95.61 | 97.44 | 71.8 |
| deSALT + GTF | -d 10 -G anno.info | 92.73 | 97.44 | 97.77 | 83.23 |
| **Mouse NS-ONT, coverage = 4X, reads from the genes with single isoform, 7146 reads/ 51741914 bases/22843 exons** | | | | | |
| GMAP | -f samse --cross-species -z sense_force | 93.67 | 50.51 | 8.83 | 4.84 |
| Graphmap2 | -x rnaseq | 93.56 | 57.79 | 20.67 | 11.45 |
| Minimap2 | -ax splice | 94.72 | 90.26 | 98.87 | 75.16 |
| Minimap2 + GTF | -ax splice –junc-bed anno.bed | 94.73 | 92.31 | 98.84 | 78.24 |
| deSALT | -d 10 | 94.71 | 90.76 | 97.36 | 75.13 |
| deSALT + GTF | -d 10 -G anno.info | 94.72 | 92.17 | 97.38 | 77.88 |
| **Mouse NS-ONT, coverage = 4X, reads from the genes with multiple isoforms, 29856 reads/ 164498526 bases/543123 exons** | | | | | |

| | | | | | |
|---|---|---|---|---|---|
| GMAP | -f samse --cross-species -z sense_force | 90.23 | 66.17 | 14.14 | 4.15 |
| Graphmap2 | -x rnaseq | 87.5 | 78.5 | 55.9 | 12.3 |
| Minimap2 | -ax splice | 92.92 | 92.81 | 96.45 | 46.27 |
| Minimap2 + GTF | -ax splice –junc-bed anno.bed | 93.03 | 96.5 | 98.1 | 64.31 |
| deSALT | -d 10 | 93.46 | 95.67 | 97.88 | 56.19 |
| deSALT + GTF | -d 10 -G anno.info | 93.49 | 97.55 | 98.33 | 71.38 |
| **Mouse NS-ONT, coverage = 10X, reads from the genes with single isoform, 17512 reads/ 126715118 bases/54985 exons** | | | | | |
| GMAP | -f samse --cross-species -z sense_force | 93.5 | 50.67 | 9.07 | 5.03 |
| Graphmap2 | -x rnaseq | 93.52 | 58.22 | 21.55 | 12.17 |
| Minimap2 | -ax splice | 94.71 | 90.58 | 98.64 | 75.5 |
| Minimap2 + GTF | -ax splice –junc-bed anno.bed | 94.72 | 92.61 | 98.76 | 78.35 |
| deSALT | -d 10 | 94.69 | 91.1 | 97.2 | 75.17 |
| deSALT + GTF | -d 10 -G anno.info | 94.69 | 92.54 | 97.28 | 77.83 |
| **Mouse NS-ONT, coverage = 10X, reads from the genes with multiple isoforms, 72490 reads/ 399385135 bases/ 1319471 exons** | | | | | |
| GMAP | -f samse --cross-species -z sense_force | 90.22 | 66.08 | 13.63 | 3.92 |
| Graphmap2 | -x rnaseq | 87.36 | 77.91 | 54.19 | 12.03 |
| Minimap2 | -ax splice | 92.97 | 92.85 | 96.46 | 46.31 |
| Minimap2 + GTF | -ax splice –junc-bed anno.bed | 93.08 | 96.54 | 98.08 | 64.12 |
| deSALT | -d 10 | 95.35 | 95.7 | 98.05 | 56.31 |
| deSALT + GTF | -d 10 -G anno.info | 93.56 | 97.6 | 98.48 | 71.68 |
| **Mouse NS-ONT, coverage = 30X, reads from the genes with single isoform, 51904 reads/ 374294776 bases/ 164621 exons** | | | | | |
| GMAP | -f samse --cross-species -z sense_force | 99.39 | 50.72 | 9.13 | 5.23 |
| Graphmap2 | -x rnaseq | 93.55 | 58.3 | 21.67 | 12.27 |
| Minimap2 | -ax splice | 94.77 | 90.37 | 98.57 | 74.91 |
| Minimap2 + GTF | -ax splice –junc-bed anno.bed | 94.78 | 92.32 | 98.7 | 77.78 |
| deSALT | -d 10 | 94.76 | 91.05 | 97.31 | 75.12 |
| deSALT + GTF | -d 10 -G anno.info | 94.77 | 92.44 | 97.4 | 77.68 |
| **Mouse NS-ONT, coverage = 30X, reads from the genes with multiple isoforms, 218098 reads/ 1200852288 bases/ 3943437 exons** | | | | | |
| GMAP | -f samse --cross-species -z sense_force | 90.17 | 65.96 | 13.65 | 4.02 |
| Graphmap2 | -x rnaseq | 87.41 | 75.44 | 50.29 | 11.49 |
| Minimap2 | -ax splice | 92.95 | 92.78 | 96.33 | 46.23 |
| Minimap2 + GTF | -ax splice –junc-bed anno.bed | 93.06 | 96.6 | 98.04 | 64.1 |

| | | | | | |
|---|---|---|---|---|---|
| deSALT | -d 10 | 93.58 | 95.72 | 98.19 | 56.22 |
| deSALT + GTF | -d 10 -G anno.info | 93.61 | 97.62 | 98.48 | 71.22 |
| **Human NS-ONT, coverage = 4X, reads from the genes with single isoform, 3829 reads/ 22232609 bases/7506 exons** | | | | | |
| GMAP | -f samse --cross-species -z sense_force | 91.94 | 38.88 | 13.37 | 7.7 |
| Graphmap2 | -x rnaseq | 93.19 | 48.59 | 25.75 | 17.34 |
| Minimap2 | -ax splice | 94.04 | 87.14 | 97.75 | 76.99 |
| Minimap2 + GTF | -ax splice –junc-bed anno.bed | 94.05 | 88.58 | 97.96 | 78.64 |
| deSALT | -d 10 | 94.09 | 87.81 | 98.02 | 77.83 |
| deSALT + GTF | -d 10 -G anno.info | 94.09 | 88.88 | 98.12 | 79.24 |
| **Human NS-ONT, coverage = 4X, reads from the genes with multiple isoforms, 33173 reads/ 193814923 bases/488881 exons** | | | | | |
| GMAP | -f samse --cross-species -z sense_force | 90.96 | 65.59 | 13.32 | 4.09 |
| Graphmap2 | -x rnaseq | 87.76 | 74.4 | 47.83 | 11.9 |
| Minimap2 | -ax splice | 93.79 | 93.22 | 96.92 | 53.04 |
| Minimap2 + GTF | -ax splice –junc-bed anno.bed | 93.89 | 96.95 | 98.4 | 69.25 |
| deSALT | -d 10 | 93.94 | 95.72 | 98.28 | 61.55 |
| deSALT + GTF | -d 10 -G anno.info | 93.97 | 97.64 | 98.57 | 75.64 |
| **Human NS-ONT, coverage = 10X, reads from the genes with single isoform, 9447 reads/54578014 bases/21120 exons** | | | | | |
| GMAP | -f samse --cross-species -z sense_force | 92.1 | 42.43 | 14.26 | 8.47 |
| Graphmap2 | -x rnaseq | 93.25 | 54.08 | 26.38 | 16.69 |
| Minimap2 | -ax splice | 94.06 | 87.29 | 97.79 | 75.3 |
| Minimap2 + GTF | -ax splice –junc-bed anno.bed | 94.07 | 89.11 | 98.05 | 77.48 |
| deSALT | -d 10 | 94.11 | 87.96 | 97.75 | 76.29 |
| deSALT + GTF | -d 10 -G anno.info | 94.09 | 89.1 | 97.89 | 78.09 |
| **Human NS-ONT, coverage = 10X, reads from the genes with multiple isoforms, 80555 reads/470728552 bases/1179903 exons** | | | | | |
| GMAP | -f samse --cross-species -z sense_force | 90.89 | 65.46 | 13.25 | 4.01 |
| Graphmap2 | -x rnaseq | 87.51 | 71.57 | 44.59 | 11.89 |
| Minimap2 | -ax splice | 93.77 | 93.28 | 96.94 | 53.56 |
| Minimap2 + GTF | -ax splice –junc-bed anno.bed | 93.86 | 97.01 | 98.35 | 69.67 |
| deSALT | -d 10 | 93.91 | 95.76 | 98.35 | 62.21 |
| deSALT + GTF | -d 10 -G anno.info | 93.94 | 97.67 | 98.64 | 75.88 |
| **Human NS-ONT, coverage = 30X, reads from the genes with single isoform, 27786 reads/161989024 bases/59904 exons** | | | | | |
| GMAP | -f samse --cross-species -z sense_force | 91.95 | 41.52 | 13.2 | 7.6 |

| | | | | | |
|---|---|---|---|---|---|
| Graphmap2 | -x rnaseq | 93.18 | 52.52 | 25.98 | 16.52 |
| Minimap2 | -ax splice | 93.95 | 86.83 | 97.51 | 75.3 |
| Minimap2 + GTF | -ax splice –junc-bed anno.bed | 93.94 | 88.67 | 97.83 | 77.35 |
| deSALT | -d 10 | 94.1 | 87.69 | 98.02 | 76.71 |
| deSALT + GTF | -d 10 -G anno.info | 94.01 | 88.93 | 98.11 | 78.46 |
| **Human NS-ONT, coverage = 30X, reads from the genes with multiple isoforms, 242216 reads/ 1413557360 bases/ 3551546 exons** | | | | | |
| GMAP | -f samse --cross-species -z sense_force | 91.02 | 65.59 | 13.32 | 4.11 |
| Graphmap2 | -x rnaseq | 87.39 | 69.89 | 41.74 | 11.18 |
| Minimap2 | -ax splice | 93.84 | 93.28 | 96.9 | 53.53 |
| Minimap2 + GTF | -ax splice –junc-bed anno.bed | 93.94 | 97.01 | 98.36 | 69.7 |
| deSALT | -d 10 | 94.1 | 95.71 | 98.37 | 61.38 |
| deSALT + GTF | -d 10 -G anno.info | 94.02 | 97.64 | 98.65 | 75.58 |
| **Human all protein coding genes NS-ONT, reads from the genes with single isoform, 36196 reads/ 194501927 bases/ 193646 exons** | | | | | |
| GMAP | -f samse --cross-species -z sense_force | 75.01 | 71.48 | 14.27 | 7.43 |
| Graphmap2 | -x rnaseq | 90.76 | 59.95 | 32.7 | 16.52 |
| Minimap2 | -ax splice | 91.5 | 75.39 | 94.74 | 69.17 |
| Minimap2 + GTF | -ax splice –junc-bed anno.bed | 91.51 | 78.33 | 95.72 | 74.77 |
| deSALT | -d 10 | 92.24 | 84.12 | 95.93 | 72.35 |
| deSALT + GTF | -d 10 -G anno.info | 92.25 | 85.77 | 96.22 | 76.33 |
| **Human all protein coding genes NS-ONT, reads from the genes with multiple isoforms, 957556 reads/ 5604052729 bases/ 14333466 exons** | | | | | |
| GMAP | -f samse --cross-species -z sense_force | 91.18 | 64.54 | 13.38 | 4.24 |
| Graphmap2 | -x rnaseq | 86.89 | 70.9 | 43.67 | 10.93 |
| Minimap2 | -ax splice | 93.22 | 91.1 | 95.97 | 52.22 |
| Minimap2 + GTF | -ax splice –junc-bed anno.bed | 93.34 | 95.16 | 97.61 | 68.18 |
| deSALT | -d 10 | 93.52 | 94.34 | 97.75 | 60.69 |
| deSALT + GTF | -d 10 -G anno.info | 93.55 | 96.37 | 98.04 | 74.45 |

a) This table depicts the results of the aligners on the reads from the genes with single and multiple isoforms. For each of the simulated datasets, the reads from the genes with single and multiple isoforms are respectively divided into two groups, and the assessment was separately implemented both of them.

b) Refer to the Supplementary Notes for the command lines of the benchmarked aligners.

c) Base%: the proportion of bases being correctly aligned to their ground truth positions, i.e., the mapped positions of the bases are within 5 bp of their ground truth positions.

d) Exon%: the proportion of the exons being correctly mapped. An exon in a certain read is considered to be correctly mapped only if its two boundaries are mapped within 5 bp of their ground truth positions in the reference genome.

e) Read80%: the proportion of Read80% reads. A read is considered to be a Read80% read only if it meets two conditions: $N_T/N_G > 80\%$ and $N_T/N_P > 80\%$, where $N_G$ is the number of ground truth exons within the read, $N_P$ is the number of exons predicted by the alignment, and $N_T$ is the number of true positive exons. Herein, a predicted exon is considered to be a true positive exon only if there is a ground truth exon in the read and the corresponding boundaries of the predicted exon and the ground truth exon are within 5 bp.

f) Read100%: the proportion of Read100% reads. A read is considered to be a Read100% read only if it meets two conditions: $N_T/N_G = 100\%$ and $N_T/N_P = 100\%$. It is worth noting that a Read100% read indicates that the read has a highly correct full-length alignment.

#### Supplementary Table 6. Benchmark results on the reads from the highly- and lowly expressed isoforms of the human all coding genes datasets[a]

| Aligner | Parameters [b] | Base% [c] | Exons% [d] | Read80% [e] | Read100% [f] |
|---|---|---|---|---|---|
| | | **Simulated PacBio ROI datasets** | | | |
| | | **Human PacBio ROI, highly expressed, 881869 reads/ 1759973613 bases/ 5641017 exons** | | | |
| GMAP | -f samse --cross-species -z sense_force | 99.44 | 95.54 | 97.25 | 84.37 |
| Graphmap2 | -x rnaseq | 96.14 | 90.05 | 93.39 | 76.8 |
| Minimap2 | -ax splice | 99.34 | 97.34 | 98.56 | 91.14 |
| Minimap2 + GTF | -ax splice –junc-bed anno.bed | 99.6 | 98.07 | 99.05 | 92.41 |
| deSALT | -d 10 -x ccs | 99.72 | 98.29 | 99.17 | 92.31 |
| deSALT + GTF | -d 10 -x ccs -G anno.info | 99.76 | 98.37 | 99.26 | 92.66 |
| | | **Human PacBio ROI, lowly expressed, 494224 reads/ 660024957 bases/ 3109755 exons** | | | |
| GMAP | -f samse --cross-species -z sense_force | 99.49 | 95.67 | 97.14 | 84.43 |
| Graphmap2 | -x rnaseq | 95.54 | 90.32 | 93.67 | 76.69 |
| Minimap2 | -ax splice | 99.17 | 96.73 | 98.13 | 89.81 |
| Minimap2 + GTF | -ax splice –junc-bed anno.bed | 99.58 | 97.82 | 99.01 | 91.72 |
| deSALT | -d 10 -x ccs | 99.76 | 98.27 | 99.21 | 92.19 |
| deSALT + GTF | -d 10 -x ccs -G anno.info | 99.87 | 98.25 | 99.28 | 91.97 |
| | | **Simulated ONT 2D (1D$^2$) datasets** | | | |
| | | **Human ONT 2D (1D$^2$), highly expressed, 659615 reads/ 1759570903 bases/ 5403089 exons** | | | |
| GMAP | -f samse --cross-species -z sense_force | 94.82 | 70.97 | 61.72 | 33.07 |
| Graphmap2 | -x rnaseq | 91.7 | 80.7 | 82.73 | 54.22 |
| Minimap2 | -ax splice | 97.45 | 93.08 | 93.35 | 70.55 |
| Minimap2 + GTF | -ax splice –junc-bed anno.bed | 97.76 | 94.78 | 94.77 | 74.56 |
| deSALT | -d 10 -x ont2d -s 2 | 98.31 | 96.82 | 97.07 | 84.06 |
| deSALT + GTF | -d 10 -x ont2d -s 2 -G anno.info | 98.3 | 97.11 | 97.24 | 85.57 |
| | | **Human ONT 2D (1D$^2$), lowly expressed, 442071 reads/ 655132821 bases/ 3018991 exons** | | | |
| GMAP | -f samse --cross-species -z sense_force | 95.51 | 69.59 | 61.36 | 34.03 |
| Graphmap2 | -x rnaseq | 90.97 | 81.43 | 83.61 | 53.76 |

| | | | | | |
|---|---|---|---|---|---|
| Minimap2 | -ax splice | 95.46 | 88.51 | 87.95 | 60.5 |
| Minimap2 + GTF | -ax splice –junc-bed anno.bed | 96.03 | 90.69 | 90.12 | 64.23 |
| deSALT | -d 10 -x ont2d -s 2 | 97.68 | 93.71 | 94.24 | 72.3 |
| deSALT + GTF | -d 10 -x ont2d -s 2 -G anno.info | 97.37 | 94.51 | 94.92 | 75.37 |
| **Simulated PacBio subread datasets** | | | | | |
| **Human PacBio subread, highly expressed, 886829 reads/ 1759443220 bases/ 5253772 exons** | | | | | |
| GMAP | -f samse --cross-species -z sense_force | 80.1 | 58.29 | 50.46 | 27.25 |
| Graphmap2 | -x rnaseq | 85.18 | 80.12 | 78.22 | 52.79 |
| Minimap2 | -ax splice | 87.82 | 89.73 | 93.06 | 64.35 |
| Minimap2 + GTF | -ax splice –junc-bed anno.bed | 88.08 | 91.51 | 94.34 | 67.09 |
| deSALT | -d 10 -x clr -s 2 | 88.88 | 95.06 | 97.01 | 79.09 |
| deSALT + GTF | -d 10 -x clr -s 2 -G anno.info | 88.86 | 95.39 | 97.16 | 80.46 |
| **Human PacBio subread, lowly expressed, 503521 reads/ 656552126 bases/ 2903129 exons** | | | | | |
| GMAP | -f samse --cross-species -z sense_force | 70.01 | 49.19 | 35.68 | 18.31 |
| Graphmap2 | -x rnaseq | 83.94 | 79.87 | 82.35 | 47.8 |
| Minimap2 | -ax splice | 85.71 | 85.46 | 86.06 | 54.39 |
| Minimap2 + GTF | -ax splice –junc-bed anno.bed | 86.19 | 87.57 | 88.15 | 57.55 |
| deSALT | -d 10 -x clr -s 2 | 88.37 | 92.42 | 94.2 | 70.12 |
| deSALT + GTF | -d 10 -x clr -s 2 -G anno.info | 88.42 | 93.27 | 94.87 | 72.41 |
| **Simulated ONT 1D datasets** | | | | | |
| **Human ONT 1D, highly expressed, 681451 reads/ 1759269337 bases/ 5711070 exons** | | | | | |
| GMAP | -f samse --cross-species -z sense_force | 57.33 | 12.53 | 9.18 | 1.66 |
| Graphmap2 | -x rnaseq | 84.84 | 47.61 | 40.12 | 15.88 |
| Minimap2 | -ax splice | 89.42 | 63.54 | 51.44 | 22.84 |
| Minimap2 + GTF | -ax splice –junc-bed anno.bed | 90.76 | 70.67 | 58.18 | 29.89 |
| deSALT | -d 10 -x ont1d -s 2 -l 14 | 95.04 | 70.95 | 61.87 | 27.84 |
| deSALT + GTF | -d 10 -x ont1d -s 2 -l 14 -G anno.info | 95.48 | 76.16 | 66.36 | 35.82 |
| **Human ONT 1D, lowly expressed, 444877 reads/ 655058444 bases/ 3173083 exons** | | | | | |
| GMAP | -f samse --cross-species -z sense_force | 54.18 | 9.91 | 3.69 | 0.92 |

| | | | | | |
|---|---|---|---|---|---|
| Graphmap2 | -x rnaseq | 75.7 | 43.09 | 31.59 | 11.68 |
| Minimap2 | -ax splice | 72.98 | 45.77 | 30.93 | 12.88 |
| Minimap2 + GTF | -ax splice –junc-bed anno.bed | 74.56 | 53.23 | 36.16 | 17.17 |
| deSALT | -d 10 -x ont1d -s 2 -l 14 | 84.4 | 61.51 | 47.27 | 20.74 |
| deSALT + GTF | -d 10 -x ont1d -s 2 -l 14 -G anno.info | 84.98 | 65.21 | 50.71 | 25.67 |
| **Simulated datasets by PBSim based on a real ONT dataset (PS-ONT)** | | | | | |
| **Human ONT, highly expressed, 695911 reads/ 1759469285 bases/ 5328674 exons** | | | | | |
| GMAP | -f samse --cross-species -z sense_force | 94.02 | 78.1 | 72.81 | 39.7 |
| Graphmap2 | -x rnaseq | 90.59 | 82.84 | 86.86 | 58.99 |
| Minimap2 | -ax splice | 95.19 | 94.4 | 95.72 | 74.66 |
| Minimap2 + GTF | -ax splice –junc-bed anno.bed | 95.44 | 95.69 | 96.74 | 77.77 |
| deSALT | -d 10 -x ont1d -s 2 -l 14 | 95.71 | 95.78 | 96.4 | 81.38 |
| deSALT + GTF | -d 10 -x ont1d -s 2 -l 14 -G anno.info | 95.69 | 95.97 | 96.21 | 82.65 |
| **Human ONT, lowly expressed, 671170 reads/ 865190010 bases/ 3981538 exons** | | | | | |
| GMAP | -f samse --cross-species -z sense_force | 92.32 | 75.99 | 71.16 | 40.6 |
| Graphmap2 | -x rnaseq | 89.76 | 81.77 | 85.25 | 55.56 |
| Minimap2 | -ax splice | 93.44 | 89.16 | 84.04 | 60.26 |
| Minimap2 + GTF | -ax splice –junc-bed anno.bed | 93.86 | 90.77 | 85.46 | 63.02 |
| deSALT | -d 10 -x ont1d -s 2 -l 14 | 95.03 | 94.82 | 93.68 | 75.81 |
| deSALT + GTF | -d 10 -x ont1d -s 2 -l 14 -G anno.info | 95.06 | 95.3 | 93.86 | 77.6 |
| **Simulated datasets by NanoSim based on a real ONT dataset (NS-ONT)** | | | | | |
| **Human NS-ONT, highly expressed, 593850 reads/ 3480172728 bases/ 7901187exons** | | | | | |
| GMAP | -f samse --cross-species -z sense_force | 90.79 | 64.05 | 13.65 | 4.85 |
| Graphmap2 | -x rnaseq | 87.7 | 70.87 | 42.21 | 11.64 |
| Minimap2 | -ax splice | 93.42 | 91.03 | 96.52 | 55.93 |
| Minimap2 + GTF | -ax splice –junc-bed anno.bed | 93.54 | 95.06 | 98.14 | 70.54 |
| deSALT | -d 10 -s 2 -l 14 | 93.71 | 94.52 | 98.22 | 64.01 |
| deSALT + GTF | -d 10 -s 2 -l 14 -G anno.info | 93.74 | 96.43 | 98.5 | 76.25 |
| **Human ONT, lowly expressed, 393417 reads/ 2302592847 bases/ 6579043 exons** | | | | | |

| | | | | | |
|---|---|---|---|---|---|
| GMAP | -f samse --cross-species -z sense_force | 90.4 | 65.07 | 13.15 | 3.81 |
| Graphmap2 | -x rnaseq | 86.59 | 71.11 | 45.58 | 10.55 |
| Minimap2 | -ax splice | 93.42 | 91.38 | 96.6 | 49.03 |
| Minimap2 + GTF | -ax splice –junc-bed anno.bed | 93.53 | 95.47 | 98.25 | 66.34 |
| deSALT | -d 10 -s 2 -l 14 | 93.76 | 94.5 | 98.49 | 57.7 |
| deSALT + GTF | -d 10 -s 2 -l 14 -G anno.info | 93.8 | 96.66 | 98.81 | 73.13 |

a) This table depicts the results of the aligners on the reads from “highly expressed isoforms” and “lowly expressed isoforms” of the human all coding genes datasets. The assessment was separately implemented for both of the two groups of reads.

b) Refer to the Supplementary Notes for the command lines of the benchmarked aligners.

c) Base%: the proportion of bases being correctly aligned to their ground truth positions, i.e., the mapped positions of the bases are within 5 bp of their ground truth positions.

d) Exon%: the proportion of the exons being correctly mapped. An exon in a certain read is considered to be correctly mapped only if its two boundaries are mapped within 5 bp of their ground truth positions in the reference genome.

e) Read80%: the proportion of Read80% reads. A read is considered to be a Read80% read only if it meets two conditions: $N_T/N_G > 80\%$ and $N_T/N_P > 80\%$, where $N_G$ is the number of ground truth exons within the read, $N_P$ is the number of exons predicted by the alignment, and $N_T$ is the number of true positive exons. Herein, a predicted exon is considered to be a true positive exon only if there is a ground truth exon in the read and the corresponding boundaries of the predicted exon and the ground truth exon are within 5 bp.

f) Read100%: the proportion of Read100% reads. A read is considered to be a Read100% read only if it meets two conditions: $N_T/N_G = 100\%$ and $N_T/N_P = 100\%$. It is worth noting that a Read100% read indicates that the read has a highly correct full-length alignment.

#### Supplementary Table 7. Benchmark results on the reads from overlapped- and non-overlapped genes of the human all coding genes datasets[a]

| Aligner | Type | Base% [b] | Exons% [c] | Read80% [d] | Read100% [e] |
|---|---|---|---|---|---|
| **Simulated PacBio ROI datasets** | | | | | |
| deSALT | Overlap genes | 99.87 | 98.47 | 99.31 | 92.24 |
| deSALT | Non-overlap genes | 99.74 | 98.23 | 99.19 | 92.18 |
| **Simulated ONT 2D ($1D^2$) datasets** | | | | | |
| deSALT | Overlap genes | 98.36 | 96.08 | 95.94 | 79.3 |
| deSALT | Non-overlap genes | 98.09 | 95.63 | 95.93 | 79.65 |
| **Simulated PacBio subread datasets** | | | | | |
| deSALT | Overlap genes | 88.94 | 94.53 | 96.14 | 75.25 |
| deSALT | Non-overlap genes | 88.7 | 94.04 | 95.96 | 75.95 |
| **Simulated ONT 1D datasets** | | | | | |
| deSALT + GTF | Overlap genes | 92.64 | 69.05 | 56.53 | 24.84 |
| deSALT | Non-overlap genes | 92.06 | 67.27 | 56.02 | 25.07 |
| **Simulated datasets by PBSim based on a real ONT dataset (PS-ONT)** | | | | | |
| deSALT | Overlap genes | 95.71 | 96.07 | 95.78 | 79.19 |
| deSALT | Non-overlap genes | 95.44 | 95.4 | 94.96 | 78.59 |
| **Simulated datasets by NanoSim based on a real ONT dataset (NS-ONT)** | | | | | |
| deSALT | Overlap genes | 94.11 | 95.91 | 98.79 | 58.61 |
| deSALT | Non-overlap genes | 93.66 | 94.21 | 98.24 | 62.02 |

a) This table depicts the results of the aligners on the reads from the genomically overlapped and non-overlap genes of the human all coding genes datasets. The assessment was separately implemented for both of the two groups of reads.

b) Base%: the proportion of bases being correctly aligned to their ground truth positions, i.e., the mapped positions of the bases are within 5 bp of their ground truth positions.

c) Exon%: the proportion of the exons being correctly mapped. An exon in a certain read is considered to be correctly mapped only if its two boundaries are mapped within 5 bp of their ground truth positions in the reference genome.

d) Read80%: the proportion of Read80% reads. A read is considered to be a Read80% read only if it meets two conditions: $N_T/N_G > 80\%$ and $N_T/N_P > 80\%$, where $N_G$ is the number of ground truth exons within the read, $N_P$ is the number of exons predicted by the alignment, and $N_T$ is the number of true positive exons. Herein, a predicted exon is considered to be a true positive exon only if there is a ground truth exon in the read and the corresponding boundaries of the predicted exon and the ground truth exon are within 5 bp.

e) Read100%: the proportion of Read100% reads. A read is considered to be a Read100% read only if it meets two conditions: $N_T/N_G = 100\%$ and $N_T/N_P = 100\%$. It is worth noting that a Read100% read indicates that the read has a highly correct full-length alignment.

#### Supplementary Table 8. Speed and memory use on simulated human datasets with various numbers of threads [a]

| Aligner | Resource usage | 1 thread | 4 threads | 8 threads | 16 threads | 1 thread | 4 threads | 8 threads | 16 threads | 1 thread | 4 threads | 8 threads | 16 threads |
|---|---|---|---|---|---|---|---|---|---|---|---|---|---|
| | | Human PacBio ROI 4x dataset (46783 reads, 57203247 bases) | | | | Human PacBio ROI 10x dataset (103339 reads, 141769367 bases) | | | | Human PacBio ROI 30x dataset (292117 reads, 424010833 bases) | | | |
| GMAP | Wall time(M) | 50.2 | 16.2 | 7.1 | 6.7 | 132.4 | 37.1 | 17.2 | 15.7 | 406.2 | 101.4 | 58.8 | 43.8 |
| | CPU time(M) | 49.9 | 61.1 | 49.7 | 54.2 | 130.2 | 144.4 | 138.1 | 204.1 | 406.7 | 400.3 | 408.4 | 432.4 |
| | Memory(GB) | 5.17 | 5.32 | 6.13 | 6.79 | 5.43 | 5.79 | 6.67 | 7.99 | 5.46 | 5.66 | 6.69 | 7.76 |
| Graphmap2 | Wall time(M) | 832 | 209 | 98 | 105 | 1454 | 480 | 236 | 160 | 5036 | 750 | 424 | 402 |
| | CPU time(M) | 826 | 844 | 728 | 1354 | 1444 | 1347 | 1356 | 2370 | 5039 | 2980 | 3340 | 6168 |
| | Memory(GB) | 67.17 | 82.21 | 76.49 | 83.49 | 71.58 | 75.28 | 79.31 | 86.20 | 97.56 | 101.31 | 92.63 | 98.88 |
| Minimap2 | Wall time(M) | 16.3 | 4.6 | 2.6 | 2.1 | 39.8 | 10.9 | 5.8 | 4.6 | 111.6 | 31.4 | 16.5 | 12.3 |
| | CPU time(M) | 16.3 | 16.9 | 17.2 | 25.3 | 39.9 | 41.8 | 44.3 | 62.9 | 111.4 | 123.8 | 124.5 | 185.3 |
| | Memory(GB) | 13.82 | 14.58 | 15.83 | 18.13 | 13.85 | 16.64 | 16.56 | 21.52 | 14.61 | 16.16 | 18.59 | 20.74 |
| Minimap2+GTF | Wall time(M) | 17.3 | 5.1 | 2.7 | 2.2 | 40.8 | 11.1 | 6.1 | 4.4 | 118.3 | 29.6 | 16.1 | 11.6 |
| | CPU time(M) | 17.1 | 18.2 | 17.8 | 24.4 | 40.5 | 42.4 | 43.6 | 60.1 | 118.1 | 116.6 | 123.8 | 173.1 |
| | Memory(GB) | 13.32 | 14.02 | 15.27 | 17.07 | 13.98 | 15.27 | 16.04 | 18.85 | 14.78 | 16.49 | 18.04 | 20.41 |
| deSALT | Wall time(M) | 15.7 | 4.2 | 3.1 | 2.1 | 35.4 | 9.3 | 5.1 | 3.9 | 98.1 | 25.7 | 13.8 | 9.9 |
| | CPU time(M) | 15.7 | 14.3 | 15.2 | 20.1 | 35.4 | 34.1 | 34.3 | 47.6 | 98.2 | 99.1 | 100.7 | 138.4 |
| | Memory(GB) | 35.03 | 35.05 | 35.06 | 35.09 | 35.13 | 35.15 | 35.17 | 35.2 | 35.47 | 35.48 | 35.51 | 35.54 |
| deSALT+GTF | Wall time(M) | 13.2 | 4.2 | 2.9 | 2.4 | 29.7 | 8.8 | 5.1 | 3.9 | 77.1 | 22.8 | 12.2 | 8.4 |
| | CPU time(M) | 13.3 | 12.1 | 12.1 | 15.6 | 29.1 | 28.5 | 28.8 | 37.4 | 76.2 | 84.8 | 83.6 | 104 |
| | Memory(GB) | 35.03 | 35.05 | 35.06 | 35.09 | 35.13 | 35.15 | 35.17 | 35.20 | 35.47 | 35.49 | 35.51 | 35.54 |
| | | Human ONT 2D ($1D^2$) 4x dataset (46594 reads, 56604793 bases) | | | | Human ONT 2D ($1D^2$) 10x dataset (102920 reads, 141283445 bases) | | | | Human ONT 2D ($1D^2$) 30x dataset (299397 reads, 423898156 bases) | | | |
| GMAP | Wall time(M) | 304 | 72 | 36 | 21 | 753 | 184 | 96 | 59 | 2171 | 572 | 292 | 169 |
| | CPU time(M) | 301 | 298 | 294 | 294 | 748 | 745 | 757 | 824 | 2172 | 2178 | 2177 | 2210 |
| | Memory(GB) | 6.87 | 7.25 | 7.10 | 8.96 | 7.12 | 7.89 | 8.12 | 8.24 | 7.05 | 7.49 | 8.12 | 8.07 |
| Graphmap2 | Wall time(M) | 679 | 149 | 122 | 75 | 1218 | 264 | 196 | 122 | 4947 | 795 | 615 | 381 |
| | CPU time(M) | 674 | 581 | 908 | 1035 | 1212 | 1044 | 1541 | 1867 | 4958 | 3150 | 4834 | 5845 |
| | Memory(GB) | 67.16 | 72.03 | 76.29 | 81.91 | 71.51 | 73.91 | 76.68 | 81.68 | 84.35 | 88.52 | 97.43 | 95.82 |

| | | | | | | | | | | | | | |
|---|---|---|---|---|---|---|---|---|---|---|---|---|---|
| Minimap2 | Wall time(M) | 16.1 | 4.6 | 2.6 | 1.9 | 39.8 | 11.6 | 6.1 | 3.9 | 113.5 | 34.3 | 17.5 | 10.4 |
| | CPU time(M) | 16.2 | 17.9 | 18.5 | 21.5 | 39.9 | 44.1 | 44.3 | 52.2 | 113.4 | 133.8 | 133.5 | 152.9 |
| | Memory(GB) | 14.25 | 15.18 | 16.85 | 18.16 | 14.37 | 16.64 | 18.46 | 21.52 | 15.23 | 17.71 | 20.16 | 24.62 |
| Minimap2+GTF | Wall time(M) | 19.6 | 4.5 | 3.1 | 2.5 | 50.7 | 10.1 | 6.7 | 5.6 | 129.8 | 29.3 | 18.6 | 13.9 |
| | CPU time(M) | 19.1 | 15.9 | 20.3 | 28.6 | 50.2 | 38.6 | 47.4 | 74.3 | 129.9 | 114.7 | 143.6 | 210.6 |
| | Memory(GB) | 14.21 | 15.48 | 16.88 | 17.97 | 14.38 | 16.71 | 18.44 | 21.64 | 15.13 | 17.84 | 20.06 | 24.42 |
| deSALT | Wall time(M) | 17.9 | 5.8 | 3.2 | 2.3 | 42.5 | 11.8 | 6.6 | 4.1 | 110.6 | 33.1 | 16.8 | 9.8 |
| | CPU time(M) | 17.9 | 20.4 | 19.3 | 21.8 | 42.5 | 44.2 | 45.8 | 48.2 | 110.4 | 127.1 | 124.4 | 135.1 |
| | Memory(GB) | 35.04 | 35.06 | 35.08 | 35.11 | 35.13 | 35.16 | 35.21 | 35.28 | 35.18 | 35.22 | 35.27 | 35.27 |
| deSALT+GTF | Wall time(M) | 16.4 | 5.1 | 3.6 | 2.8 | 35.9 | 10.5 | 6.4 | 4.5 | 101.3 | 28.1 | 15.6 | 9.5 |
| | CPU time(M) | 15.7 | 14.8 | 17.1 | 19.7 | 35.8 | 36.2 | 38.3 | 45.2 | 101.3 | 103.4 | 102.3 | 106.4 |
| | Memory(GB) | 35.04 | 35.05 | 35.07 | 35.11 | 35.13 | 35.15 | 35.17 | 35.21 | 35.18 | 35.21 | 35.27 | 35.93 |
| | | **Human PacBio subread 4x dataset (46537 reads, 56809800 bases)** | | | | **Human PacBio subread 10x dataset (102533 reads, 141326345 bases)** | | | | **Human PacBio subread 30x dataset (289375 reads, 423841890 bases)** | | | |
| GMAP | Wall time(M) | 506 | 123 | 66 | 52 | 1411 | 348 | 184 | 159 | 4048 | 1061 | 517 | 398 |
| | CPU time(M) | 501 | 498 | 521 | 716 | 1402 | 1396 | 1453 | 2210 | 4085 | 4077 | 4086 | 4101 |
| | Memory(GB) | 6.45 | 7.12 | 7.56 | 8.31 | 7.13 | 7.84 | 7.93 | 8.78 | 7.14 | 7.77 | 8.45 | 9.16 |
| Graphmap2 | Wall time(M) | 685 | 148 | 89 | 80 | 1280 | 619 | 153 | 142 | 3344 | 763 | 383 | 358 |
| | CPU time(M) | 681 | 566 | 652 | 1103 | 1273 | 1337 | 1177 | 2118 | 3312 | 3020 | 3019 | 5576 |
| | Memory(GB) | 67.27 | 71.98 | 76.34 | 82.81 | 72.01 | 74.72 | 79.01 | 84.35 | 87.53 | 89.76 | 93.03 | 98.18 |
| Minimap2 | Wall time(M) | 15.3 | 4.2 | 2.5 | 2.2 | 39.3 | 9.6 | 5.4 | 4.6 | 110.7 | 28.1 | 15.1 | 12.9 |
| | CPU time(M) | 15.3 | 14.9 | 15.8 | 26.8 | 39.4 | 35.5 | 38.5 | 46.3 | 110.7 | 109.8 | 115.1 | 185.2 |
| | Memory(GB) | 13.72 | 15.19 | 17.04 | 19.58 | 13.95 | 16.09 | 18.44 | 21.59 | 16.25 | 18.35 | 21.77 | 26.55 |
| Minimap2+GTF | Wall time(M) | 14.7 | 4.1 | 2.4 | 2.1 | 36.1 | 9.4 | 5.3 | 4.2 | 105.7 | 26.9 | 14.9 | 11.6 |
| | CPU time(M) | 14.5 | 14.6 | 15.3 | 23.8 | 36.7 | 35.6 | 37.7 | 57.8 | 106.1 | 105.1 | 112.9 | 172.3 |
| | Memory(GB) | 13.89 | 15.48 | 17.17 | 20.15 | 14.47 | 17.09 | 19.26 | 22.85 | 15.49 | 18.42 | 21.67 | 25.12 |
| deSALT | Wall time(M) | 18.1 | 5.6 | 2.8 | 2.6 | 43.3 | 10.7 | 5.8 | 4.6 | 117.1 | 29.6 | 15.8 | 12.4 |
| | CPU time(M) | 18.2 | 19.5 | 16.9 | 27.9 | 42.1 | 39.9 | 39.5 | 57.1 | 117.1 | 113.3 | 116.5 | 175.6 |
| | Memory(GB) | 35.03 | 35.05 | 35.07 | 35.11 | 35.13 | 35.15 | 35.17 | 35.21 | 35.47 | 35.48 | 35.51 | 35.54 |
| deSALT+GTF | Wall time(M) | 14.8 | 5.4 | 3.5 | 2.7 | 34.4 | 10.4 | 6.1 | 4.5 | 98.3 | 28.5 | 15.4 | 10.6 |
| | CPU time(M) | 14.1 | 16.4 | 15.9 | 18.5 | 34.1 | 36.2 | 35.9 | 45.7 | 98.1 | 106.7 | 106.9 | 136.1 |
| | Memory(GB) | 35.03 | 35.05 | 35.07 | 35.10 | 35.13 | 35.15 | 35.17 | 35.21 | 35.47 | 35.48 | 35.51 | 35.54 |

| | | Human ONT 1D 4x dataset (46644 reads, 56517199 bases) | | | | Human ONT 1D 10x dataset (103532 reads, 141296288 bases) | | | | Human ONT 1D 30x dataset (309135 reads, 423843578 bases) | | | |
|---|---|---|---|---|---|---|---|---|---|---|---|---|---|
| GMAP | Wall time(M) | 1115 | 289 | 187 | 144 | 3824 | 978 | 766 | 652 | 11474 | 2335 | 1524 | 978 |
| | CPU time(M) | 1114 | 1128 | 1354 | 1989 | 3815 | 3901 | 5977 | 7175 | 114723 | 11480 | 11567 | 15634 |
| | Memory(GB) | 8.21 | 8.56 | 9.14 | 9.54 | 8.47 | 9.12 | 9.56 | 10.35 | 9.23 | 9.45 | 10.21 | 10.56 |
| Graphmap2 | Wall time(M) | 572 | 122 | 78 | 60 | 1133 | 243 | 146 | 117 | 3136 | 759 | 398 | 352 |
| | CPU time(M) | 569 | 476 | 608 | 892 | 1127 | 962 | 1146 | 1808 | 3097 | 2979 | 3193 | 5460 |
| | Memory(GB) | 66.81 | 70.63 | 73.16 | 78.16 | 78.49 | 73.42 | 81.92 | 79.31 | 82.47 | 86.15 | 88.38 | 98.61 |
| Minimap2 | Wall time(M) | 14.3 | 4.3 | 2.4 | 2.1 | 37.3 | 9.8 | 5.3 | 4.5 | 105.5 | 27.9 | 14.8 | 12.3 |
| | CPU time(M) | 14.3 | 15.2 | 15.4 | 24.1 | 37.4 | 37.5 | 38.4 | 62.1 | 105.5 | 109.2 | 113.4 | 182.7 |
| | Memory(GB) | 14.14 | 16.39 | 18.43 | 22.11 | 14.45 | 16.98 | 19.43 | 24.27 | 15.69 | 19.41 | 22.78 | 28.96 |
| Minimap2+GTF | Wall time(M) | 14.8 | 4.1 | 2.4 | 2.0 | 37 | 9.6 | 5.4 | 4.3 | 107.8 | 27.6 | 15.3 | 11.8 |
| | CPU time(M) | 14.3 | 14.1 | 15.1 | 23.18 | 36.9 | 36.5 | 38.6 | 59.3 | 107.5 | 108.1 | 114.3 | 175.6 |
| | Memory(GB) | 14.14 | 16.03 | 18.30 | 22.72 | 14.45 | 17.09 | 20.05 | 24.33 | 15.64 | 19.55 | 21.98 | 29.07 |
| deSALT | Wall time(M) | 19.6 | 5.8 | 3.6 | 3.2 | 50.4 | 12.4 | 6.9 | 5.7 | 135.9 | 35.2 | 19.7 | 15.7 |
| | CPU time(M) | 19.1 | 20.6 | 22.2 | 35.6 | 49.9 | 46.8 | 49.5 | 74.8 | 135.8 | 136.5 | 147.6 | 235.2 |
| | Memory(GB) | 35.03 | 35.05 | 35.06 | 35.09 | 35.14 | 35.15 | 35.17 | 35.20 | 35.48 | 35.49 | 35.53 | 35.55 |
| deSALT+GTF | Wall time(M) | 16.1 | 5.2 | 3.5 | 2.9 | 40.5 | 10.9 | 6.3 | 5.1 | 108.7 | 30.1 | 16.8 | 12.2 |
| | CPU time(M) | 15.5 | 16.1 | 16.4 | 22.1 | 38.7 | 38.2 | 38.8 | 53.9 | 108.8 | 113.4 | 118.8 | 160.6 |
| | Memory(GB) | 35.03 | 35.05 | 35.06 | 35.09 | 35.14 | 35.15 | 35.17 | 35.21 | 35.48 | 35.49 | 35.52 | 35.55 |
| | | **PS-ONT 4x dataset (47186 reads, 216047531 bases)** | | | | **PS-ONT 10x dataset (90001 reads, 525306565 bases)** | | | | **PS-ONT 30x dataset (270001 reads, 1575546383 bases)** | | | |
| GMAP | Wall time(M) | 291 | 66 | 39 | 29 | 1055 | 151 | 76 | 59 | 7836 | 606 | 250 | 214 |
| | CPU time(M) | 287 | 256 | 239 | 316 | 631 | 593 | 603 | 925 | 1820 | 2395 | 1976 | 2945 |
| | Memory(GB) | 11.25 | 11.79 | 12.36 | 13.01 | 12.32 | 12.79 | 13.45 | 14.42 | 12.03 | 13.2 | 12.57 | 13.71 |
| Graphmap2 | Wall time(M) | 730 | 157 | 85 | 79 | 1261 | 330 | 152 | 132 | 3915 | 756 | 423 | 397 |
| | CPU time(M) | 724 | 611 | 645 | 1074 | 1252 | 1305 | 1189 | 1970 | 3474 | 3001 | 3334 | 5823 |
| | Memory(GB) | 67.14 | 71.32 | 76.75 | 83.78 | 71.48 | 74.92 | 77.95 | 84.65 | 86.07 | 88.71 | 92.12 | 98.74 |
| Minimap2 | Wall time(M) | 14.9 | 5.4 | 2.5 | 2.2 | 36.6 | 11.9 | 5.4 | 4.9 | 103.2 | 30.8 | 15.1 | 12.5 |
| | CPU time(M) | 14.7 | 22.2 | 16.1 | 25.6 | 36.4 | 45.7 | 38.5 | 67.5 | 103.1 | 120.6 | 114.8 | 184.6 |
| | Memory(GB) | 13.84 | 15.55 | 17.03 | 19.27 | 14.20 | 15.97 | 17.45 | 19.34 | 15.57 | 17.81 | 20.62 | 24.44 |
| Minimap2+G | Wall time(M) | 16.1 | 5.6 | 2.9 | 2.3 | 38.2 | 11.7 | 5.7 | 4.6 | 113.3 | 36.4 | 17.3 | 14.2 |

| | | | | | | | | | | | | | |
|---|---|---|---|---|---|---|---|---|---|---|---|---|---|
| TF | CPU time(M) | 15.7 | 20.2 | 19.3 | 26.2 | 37.5 | 44.9 | 41.4 | 63.2 | 112.9 | 142.3 | 132.7 | 207.7 |
| | Memory(GB) | 13.85 | 15.63 | 17.01 | 19.48 | 14.22 | 15.83 | 17.61 | 19.28 | 15.59 | 17.82 | 20.48 | 24.44 |
| deSALT | Wall time(M) | 18.3 | 5.4 | 3.2 | 2.8 | 39.1 | 13.2 | 6.2 | 5.1 | 113.9 | 34.9 | 17.1 | 13.6 |
| | CPU time(M) | 18.1 | 18.8 | 19.9 | 29.5 | 38.9 | 49.5 | 43.1 | 61 | 113.2 | 134.6 | 125.2 | 188.4 |
| | Memory(GB) | 35.5 | 35.05 | 35.08 | 35.10 | 35.13 | 35.14 | 35.16 | 35.20 | 35.47 | 35.48 | 35.51 | 35.54 |
| deSALT+GTF | Wall time(M) | 25.1 | 5.4 | 2.8 | 2.7 | 39.5 | 12.5 | 6.8 | 5.3 | 117.3 | 33.8 | 16.7 | 14.9 |
| | CPU time(M) | 25.1 | 18.8 | 16.3 | 27.5 | 39.1 | 46.7 | 46.4 | 65.1 | 117.1 | 130.5 | 122.1 | 195.7 |
| | Memory(GB) | 35.03 | 35.05 | 35.07 | 35.10 | 35.13 | 35.14 | 35.16 | 35.20 | 35.47 | 35.48 | 35.50 | 35.54 |
| | | **NS-ONT 4x dataset (37001 reads, 56517199 bases)** | | | | **NS-ONT 10x dataset (103532 reads, 141296288 bases)** | | | | **NS-ONT 30x dataset (309135 reads, 423843578 bases)** | | | |
| GMAP | Wall time(M) | 2214 | 525 | 267 | 198 | 5961 | 1425 | 798 | 898 | 17285 | 4235 | 2064 | 1436 |
| | CPU time(M) | 2192 | 2037 | 2011 | 2882 | 5868 | 5421 | 5827 | 7037 | 17235 | 15129 | 16295 | 22330 |
| | Memory(GB) | 11.45 | 11.89 | 12.12 | 12.49 | 11.34 | 12.21 | 12.89 | 13.01 | 12.33 | 12.45 | 13.05 | 13.36 |
| Graphmap2 | Wall time(M) | 845 | 309 | 117 | 108 | 1947 | 602 | 319 | 280 | 5988 | 1751 | 983 | 1127 |
| | CPU time(M) | 842 | 1223 | 913 | 1621 | 1939 | 2378 | 2475 | 3985 | 5965 | 7800 | 7744 | 12161 |
| | Memory(GB) | 68.31 | 68.83 | 69.50 | 71.23 | 74.53 | 75.85 | 76.78 | 78.08 | 94.52 | 97.40 | 98.85 | 100.24 |
| Minimap2 | Wall time(M) | 40.1 | 10.6 | 5.2 | 4.3 | 97.3 | 21.2 | 15.3 | 9.61 | 232 | 64.6 | 42.9 | 27 |
| | CPU time(M) | 41.1 | 40.5 | 36.9 | 57.8 | 97.1 | 82.5 | 85.1 | 139.1 | 232 | 255.7 | 257.7 | 418 |
| | Memory(GB) | 13.98 | 15.24 | 16.83 | 18.59 | 14.28 | 15.89 | 17.38 | 20.19 | 14.63 | 16.79 | 18.49 | 21.03 |
| Minimap2+GTF | Wall time(M) | 52.7 | 16.2 | 7.6 | 4.6 | 108.5 | 33.7 | 19.2 | 10.2 | 247.8 | 68.1 | 47.4 | 28.8 |
| | CPU time(M) | 52.4 | 62.2 | 55.6 | 61.5 | 108.1 | 132.1 | 147.2 | 148.3 | 247.3 | 269.1 | 316.8 | 442.5 |
| | Memory(GB) | 14.01 | 15.25 | 16.59 | 18.65 | 14.29 | 15.91 | 17.49 | 20.1 | 14.64 | 16.86 | 18.41 | 21.35 |
| deSALT | Wall time(M) | 93.6 | 15.9 | 8.5 | 6.8 | 127 | 37.5 | 17.7 | 14.7 | 385.9 | 105.5 | 41.9 | 40 |
| | CPU time(M) | 92.7 | 60.3 | 60.4 | 84.4 | 125 | 145.3 | 132.4 | 198.7 | 384.7 | 413.3 | 404.7 | 560 |
| | Memory(GB) | 35.21 | 35.24 | 35.28 | 35.35 | 35.53 | 35.56 | 35.59 | 35.69 | 36.66 | 36.69 | 36.72 | 36.78 |
| deSALT+GTF | Wall time(M) | 56.4 | 15.6 | 7.1 | 6.4 | 129.3 | 37.1 | 16.3 | 14.1 | 367 | 107.7 | 42.5 | 40.4 |
| | CPU time(M) | 56.3 | 58.8 | 49.7 | 79.1 | 128.2 | 143.9 | 120.9 | 190.7 | 365 | 419.3 | 365.8 | 579.2 |
| | Memory(GB) | 35.21 | 35.24 | 35.27 | 35.35 | 35.53 | 35.56 | 35.6 | 35.67 | 36.66 | 36.69 | 36.72 | 36.79 |

a) The benchmark of time cost and memory employed the 18 simulated datasets which were composed by the randomly selected human genes with 6 sequencing models and 3 kinds of sequencing coverages (4x, 10x, 30x). The time costs of the aligners are shown in minutes their memory footprints are shown in GB.

#### Supplementary Table 9. Benchmark results on the real sequencing datasets

| Aligner | Parameters | #BaseA [c] | #BaseGA [d] | #ExonP [e] | #ExonGO [f] | #ExonGA [g] | #ReadGA [h] | Time/Memory [i] |
|---|---|---|---|---|---|---|---|---|
| **real HUMAN ONT cDNA dataset, 15152101 reads, 14134831170 bases** | | | | | | | | |
| GMAP | -f samse --cross-species -z sense_force [a] | 9948549314 | 8402753847 | 39905636 | 37625378 | 17132563 | 3107592 | 30935(618804)/ 16.53GB |
| Graphmap2 | -x rnaseq | 11783661722 | 9967808509 | 48152781 | 43761553 | 20907389 | 4884624 | 2903(58003)/ 91.35GB |
| Minimap2 | -ax splice -uf -l14 [b] | 12068583337 | 10297615525 | 50372792 | 49621879 | 25201774 | 4746517 | 124(2480)/ 33.23GB |
| Minimap2 + GTF | -ax splice –junc-bed anno.bed -uf -k14 [b] | 12001215964 | 10534272431 | 53146973 | 51456769 | 31408964 | 7109314 | 217(3832) / 30.16GB |
| deSALT | -d 10 -l 15 | 12035091373 | 10555134761 | 53626292 | 52661753 | 31887344 | 6892999 | 181(3663)/ 37.65GB |
| deSALT + GTF | -d 10 -l 15 -G anno.info | 12024891192 | 10588047517 | 54760851 | 52723260 | 33126290 | 7864913 | 166(3321) / 37.68GB |
| **real HUMAN ONT direct RNA dataset, 10302647 reads, 10614186428 bases** | | | | | | | | |
| GMAP | -f samse --cross-species -z sense_force [a] | 9147741621 | 8495880466 | 40737024 | 39159719 | 22603728 | 3592296 | 8413(195944)/ 12.69GB |
| Graphmap2 | -x rnaseq | 467815455 | 406925133 | 1045344 | 1010730 | 313600 | 125856 | 2525(52503)/ 78.34GB |
| Minimap2 | -ax splice -uf | 9545417671 | 8999091103 | 42699705 | 42314689 | 28668512 | 5764834 | 106(1944)/ 32.57GB |
| Minimap2 + GTF | -ax splice –junc-bed anno.bed -uf | 9546106081 | 8999975230 | 42716675 | 42331348 | 28684461 | 5768622 | 85.9(1973)/ 33.18GB |
| deSALT | -d 10 -l 15 -T | 9590186841 | 9048155654 | 43670386 | 43091603 | 29030919 | 5853926 | 125(2265)/ 37.52GB |
| deSALT + GTF | -d 10 -l 15 -T -G anno.info | 9575008284 | 9037514484 | 43431084 | 42870738 | 29340861 | 6075128 | 130(2494)/ 37.52GB |
| **real mouse PacBio dataset 2269795 reads, 3213849871 bases** | | | | | | | | |
| GMAP | -f samse --cross-species -z sense_force | 2256745964 | 1926549648 | 8889112 | 8341684 | 5464835 | 791874 | 10961(251547)/ 62.54GB |
| Graphmap2 | -x rnaseq | -- | -- | -- | -- | -- | -- | -- |
| Minimap2 | -ax splice | 2749969434 | 2451570407 | 10792312 | 10755064 | 8103391 | 1396677 | 79(1292)/ 31.81GB |
| Minimap2 | -ax splice –junc-bed anno.bed | 2751800088 | 2459068476 | 11089541 | 10897552 | 8332463 | 1526894 | 84 (1355) / 33.52GB |
| deSALT | -d 10 -l 15 | 2769323628 | 2475226314 | 11274040 | 11149070 | 8389832 | 1463815 | 68(972)/ 38.77GB |

| | | | | | | | | |
|---|---|---|---|---|---|---|---|---|
| deSALT + GTF | -d 10 -l 15 -G anno.info | 2764786786 | 2474976131 | 11393014 | 11134880 | 8458913 | 1552100 | 65(998) / 38.68GB |

a) The parameters of GMAP were configured by referring to the recommendtion of the PacBio team (https://github.com/Magdoll/cDNA_Cupcake/wiki/Best-practice-for-aligning-Iso-Seq-to-reference-genome:-minimap2,-deSALT,-GMAP,-STAR,-BLAT).

b) The parameters of Minimap2 were configured by referring to https://github.com/nanopore-wgs-consortium/NA12878/blob/master/RNA.md.

c) #BaseA: the number of bases being aligned.

d) #BaseGA: the number of bases aligned to the positions within annotated exons.

e) #ExonP: the number of exons predicted by the alignments (also termed “predicted exons”). Herein, the predicted exons in various reads are independently considered.

f) #ExonGO: the number of predicted exons being overlapped by annotated exons (also termed “overlapped exons”). Herein, a predicted exon is considered to be overlapped by annotated exons only if there is at least one annotated exon and at least 10 bp overlapping between the predicted exon and the annotated exon.

g) #ExonGA: the number of predicted exons being exactly matched by annotated exons (also termed “exactly matched exons”). Herein, a predicted exon is considered to be exactly matched by annotated exons only if there is an annotated exon and the positions of the two boundaries of the predicted exon are within 5 bp of the corresponding boundaries of the annotated exon.

h) #ReadGA: the number of ReadGA reads. A read is considered to be a ReadGA read only if each of the intron boundaries implied by its alignment is within 5 bp of an annotated exon. Herein, a ReadGA read indicates that the read could have a correct full-length alignment.

i) The alignment time (in minutes) and memory footprint (in GB) of the aligners. All the aligners were run in 24 threads for all datasets, and the alignment time is shown in a A(B) format, where the numbers A and B are the wall clock time and the CPU time of the aligner, respectively.

#### Supplementary Table 10. Numbers of exactly matched exons of various lengths on the real sequencing datasets [a]

| Aligner | ExonGA(20) | ExonGA(30) | ExonGA(40) | ExonGA(50) | ExonGA(60) |
|---|---|---|---|---|---|
| **real HUMAN ONT dataset, 15152101 reads, 14134831170 bases** | | | | | |
| GMAP | 9332 | 208819 | 447638 | 893442 | 1604262 |
| Graphmap2 | 11820 | 234295 | 487410 | 1004192 | 1800014 |
| Minimap2 | 1775 | 138108 | 357615 | 891225 | 1859905 |
| Minimap2 + GTF | 3222 | 196583 | 505579 | 1213615 | 2490904 |
| deSALT | 7534 | 242548 | 602534 | 1367186 | 2725536 |
| deSALT + GTF | 18950 | 373088 | 784858 | 1593077 | 2998681 |
| **real HUMAN ONT direct RNA dataset, 10302647 reads, 10614186428 bases** | | | | | |
| GMAP | 19557 | 212046 | 471247 | 1008180 | 1855456 |
| Graphmap2 | 13 | 450 | 1208 | 4756 | 20574 |
| Minimap2 | 5607 | 139301 | 374648 | 992656 | 2018190 |
| Minimap2 + GTF | 5607 | 139304 | 374677 | 992944 | 2019107 |
| deSALT | 12466 | 178329 | 439625 | 1089322 | 2138303 |
| deSALT + GTF | 25524 | 232323 | 543580 | 1220809 | 2289702 |
| **real mouse PacBio dataset 2269795 reads, 3213849871 bases** | | | | | |
| GMAP | 4111 | 45191 | 137456 | 227049 | 365372 |
| Graphmap2 | -- | -- | -- | -- | -- |
| Minimap2 | 625 | 30058 | 129886 | 256000 | 463676 |
| Minimap2 + GTF | 9016 | 74328 | 212128 | 342775 | 556015 |
| deSALT | 5068 | 57585 | 178308 | 312060 | 531633 |
| deSALT + GTF | 8805 | 74944 | 213873 | 347823 | 567746 |

a) This table describes the various ExonGA(x) (x<60) statistics of the aligners, which reflect their ability to handle relatively small exons.

#### Supplementary Table 11. Numbers of *in silico* transcript sequences made from various sets of genes [a]

| Species | # of transcripts from genes having single isoforms | # of transcripts from genes having multiple isoforms | # of transcripts from genes having short (<31 bp) exons | # of all the generated transcripts |
|---|---|---|---|---|
| Fruit fly | 3009 | 3065 | 2880 | 8954 |
| Mouse | 3072 | 4080 | 2783 | 9935 |
| Human | 2998 | 3603 | 3346 | 9947 |

a) This table describes the number of *in silico* transcript sequences being composed from various sets of randomly selected genes and used to generate simulated datasets.

#### Supplementary Table 12. Benchmark results of deSALT with different *l*-mer size[a]

| Aligner | Parameters [b] | Base% [c] | Exons% [d] | Read80% [e] | Read100% [f] |
|---|---|---|---|---|---|
| **Simulated PacBio ROI datasets** | | | | | |
| **Fruit fly PacBio ROI, coverage = 4X, 45296 reads/102311172 bases/230387 exons** | | | | | |
| deSALT | -l 14 -x ccs | 99.63 | 98.78 | 99.47 | 95.05 |
| deSALT | -l 15 -x ccs | 99.61 | 98.73 | 99.43 | 94.89 |
| deSALT | -l 17 -x ccs | 99.61 | 98.58 | 99.31 | 94.37 |
| deSALT | -l 19 -x ccs | 99.57 | 98.41 | 99.2 | 93.72 |
| **Mouse PacBio ROI, coverage = 4X, 46812 reads/66151862 bases/232811 exons** | | | | | |
| deSALT | -l 14 -x ccs | 98.41 | 97.36 | 97.29 | 91.79 |
| deSALT | -l 15 -x ccs | 98.41 | 97.33 | 97.36 | 91.65 |
| deSALT | -l 17 -x ccs | 98.43 | 97.16 | 97.29 | 91.11 |
| deSALT | -l 19 -x ccs | 98.44 | 97.03 | 97.26 | 90.69 |
| **Human PacBio ROI, coverage = 4X, 46783 reads/57203247 bases/222656 exons** | | | | | |
| deSALT | -l 14 -x ccs | 99.21 | 97.7 | 98.28 | 92.85 |
| deSALT | -l 15 -x ccs | 99.21 | 97.7 | 98.38 | 92.84 |
| deSALT | -l 17 -x ccs | 99.2 | 97.55 | 98.3 | 92.33 |
| deSALT | -l 19 -x ccs | 99.17 | 97.39 | 98.15 | 91.79 |
| **Simulated ONT 2D ($1D^2$) datasets** | | | | | |
| **Fruit fly ONT 2D ($1D^2$), coverage = 4X, 44445 reads/101867685 bases/229107 exons** | | | | | |
| deSALT | -l 14 -x ont2d | 95.48 | 95.36 | 95.79 | 80.41 |
| deSALT | -l 15 -x ont2d | 95.4 | 94.96 | 94.98 | 79.12 |
| deSALT | -l 17 -x ont2d | 95.14 | 93.84 | 92.87 | 75.56 |
| deSALT | -l 19 -x ont2d | 94.57 | 92.31 | 90.07 | 71.51 |
| **Mouse ONT 2D ($1D^2$), coverage = 4X, 46591 reads/65584331 bases/232141 exons** | | | | | |
| deSALT | -l 14 -x ont2d | 93.28 | 90.79 | 88.82 | 69.75 |
| deSALT | -l 15 -x ont2d | 93.18 | 90.3 | 88.15 | 68.3 |
| deSALT | -l 17 -x ont2d | 92.35 | 87.91 | 84.32 | 63.05 |
| deSALT | -l 19 -x ont2d | 90.93 | 84.47 | 79.03 | 56.64 |

| | | | | | |
|---|---|---|---|---|---|
| **Human ONT 2D ($1D^2$), coverage = 4X, 46594 reads/56604793 bases/223025 exons** | | | | | |
| deSALT | -l 14 -x ont2d | 93.8 | 90.73 | 88.88 | 69.96 |
| deSALT | -l 15 -x ont2d | 93.64 | 90.2 | 87.99 | 68.55 |
| deSALT | -l 17 -x ont2d | 92.51 | 87.46 | 83.53 | 62.72 |
| deSALT | -l 19 -x ont2d | 90.79 | 83.89 | 78.03 | 56.29 |
| **Simulated PacBio subread datasets** | | | | | |
| **Fruit fly PacBio subread, coverage = 4X, 44169 reads/101945130 bases/212417 exons** | | | | | |
| deSALT | -l 14 -x clr | 88.1 | 93.78 | 92.01 | 75.41 |
| deSALT | -l 15 -x clr | 88.01 | 93.13 | 90.66 | 73.29 |
| deSALT | -l 17 -x clr | 87.67 | 91.35 | 87.12 | 67.81 |
| deSALT | -l 19 -x clr | 87.14 | 89.4 | 83.45 | 62.48 |
| **Mouse PacBio subread, coverage = 4X, 46481 reads/65756249 bases/216580 exons** | | | | | |
| deSALT | -l 14 -x clr | 85.51 | 88.63 | 84.31 | 63.67 |
| deSALT | -l 15 -x clr | 85.62 | 87.83 | 82.97 | 61.6 |
| deSALT | -l 17 -x clr | 84.59 | 84.39 | 77.5 | 54.01 |
| deSALT | -l 19 -x clr | 82.84 | 79.3 | 70.1 | 45.51 |
| **Human PacBio subread, coverage = 4X, 46537 reads/56809800 bases/208419 exons** | | | | | |
| deSALT | -l 14 -x clr | 86.2 | 88.1 | 86.52 | 64.67 |
| deSALT | -l 15 -x clr | 86.06 | 87.37 | 85.55 | 63.05 |
| deSALT | -l 17 -x clr | 84.76 | 83.7 | 80.23 | 56.02 |
| deSALT | -l 19 -x clr | 82.26 | 77.91 | 82.61 | 47.71 |
| **Simulated ONT 1D datasets** | | | | | |
| **Fruit fly ONT 1D, coverage = 4X, 44546 reads/101842292 bases/240905 exons** | | | | | |
| deSALT | -l 14 -x ont1d -s 2 | 94.67 | 84.33 | 81.33 | 46.47 |
| deSALT | -l 15 -x ont1d -s 2 | 94.17 | 82.54 | 78.59 | 43.25 |
| deSALT | -l 17 -x ont1d -s 2 | 90.16 | 75.03 | 66.39 | 33.83 |
| deSALT | -l 19 -x ont1d -s 2 | 78.54 | 60.23 | 47.19 | 22.26 |
| **Mouse ONT 1D, coverage = 4X, 46600 reads/65508866 bases/243889 exons** | | | | | |
| deSALT | -l 14 -x ont1d -s 2 | 83.9 | 63.28 | 52.48 | 22.89 |

| | | | | | |
|---|---|---|---|---|---|
| deSALT | -l 15 -x ont1d -s 2 | 82.94 | 60.82 | 49.53 | 20.7 |
| deSALT | -l 17 -x ont1d -s 2 | 75.77 | 48.92 | 36.63 | 14.24 |
| deSALT | -l 19 -x ont1d -s 2 | 59.8 | 31.48 | 21.7 | 7.86 |
| **Human ONT 1D, coverage = 4X, 46644 reads/56517199 bases/233621 exons** | | | | | |
| deSALT | -l 14 -x ont1d -s 2 | 81.43 | 61.15 | 49.54 | 21.99 |
| deSALT | -l 15 -x ont1d -s 2 | 80.48 | 58.07 | 46.25 | 19.6 |
| deSALT | -l 17 -x ont1d -s 2 | 72.92 | 46.17 | 33.18 | 12.87 |
| deSALT | -l 19 -x ont1d -s 2 | 56.62 | 28.23 | 18.75 | 6.82 |
| **Simulated datasets by PBSim based on a real ONT dataset (PS-ONT)** | | | | | |
| **Fruit fly PS-ONT, coverage = 4X, 46305 reads/ 101883320 bases/ 227570 exons** | | | | | |
| deSALT | -l 14 | 93..64 | 96.11 | 96.99 | 83.84 |
| deSALT | -l 15 | 93.62 | 95.85 | 96.54 | 82.93 |
| deSALT | -l 17 | 93.52 | 95.12 | 95.1 | 80.44 |
| deSALT | -l 19 | 93.37 | 94.26 | 93.66 | 77.53 |
| **Mouse PS-ONT, coverage = 4X, 47348 reads/65619390 bases/229025 exons** | | | | | |
| deSALT | -l 14 | 91.84 | 92.63 | 91.45 | 74.82 |
| deSALT | -l 15 | 91.74 | 92.2 | 90.9 | 73.7 |
| deSALT | -l 17 | 91.45 | 90.95 | 88.86 | 69.96 |
| deSALT | -l 19 | 91.01 | 89.17 | 86.09 | 65.29 |
| **Human PS-ONT, coverage = 4X, 47186 reads/56649799 bases/220303 exons** | | | | | |
| deSALT | -l 14 | 92.44 | 92.45 | 91.6 | 74.25 |
| deSALT | -l 15 | 92.37 | 92.1 | 91.11 | 73.16 |
| deSALT | -l 17 | 92.01 | 90.65 | 88.29 | 68.94 |
| deSALT | -l 19 | 91.37 | 88.64 | 85.26 | 64.27 |
| **Simulated datasets by NanoSim based on a real ONT dataset (NS-ONT)** | | | | | |
| **Fruit fly NS-ONT, coverage = 4X, 34001 reads/ 198110605 bases/ 293758 exons** | | | | | |
| deSALT | -l 14 | 92.71 | 95.59 | 97.38 | 71.73 |
| deSALT | -l 15 | 92.69 | 95.56 | 97.42 | 71.59 |
| deSALT | -l 17 | 92.68 | 95.42 | 97.26 | 70.89 |

| | | | | | |
|---|---|---|---|---|---|
| deSALT | -l 19 | 92.62 | 95.28 | 97.07 | 70.24 |
| **Mouse NS-ONT, coverage = 4X, 37001 reads/ 216240439 bases/ 565965 exons** | | | | | |
| deSALT | -l 14 | 93.78 | 95.49 | 97.81 | 60.01 |
| deSALT | -l 15 | 93.76 | 95.48 | 97.78 | 59.85 |
| deSALT | -l 17 | 93.74 | 95.38 | 97.6 | 59.37 |
| deSALT | -l 19 | 93.7 | 95.26 | 97.24 | 58.89 |
| **Human NS-ONT, coverage = 4X, 37001 reads/216047531 bases/496386 exons** | | | | | |
| deSALT | -l 14 | 93.96 | 95.63 | 98.29 | 63.31 |
| deSALT | -l 15 | 93.96 | 95.6 | 98.25 | 63.23 |
| deSALT | -l 17 | 93.95 | 95.54 | 98.18 | 62.97 |
| deSALT | -l 19 | 93.93 | 95.35 | 98.02 | 62.42 |

a) This table depicts the results of deSALT on the 4x coverage simulated datasets from human, mouse and fruit fly with various configurations of *l* parameter.

b) Refer to the Supplementary Notes for the command lines of the benchmarked aligners.

c) Base%: the proportion of bases being correctly aligned to their ground truth positions, i.e., the mapped positions of the bases are within 5 bp of their ground truth positions.

d) Exon%: the proportion of the exons being correctly mapped. An exon in a certain read is considered to be correctly mapped only if its two boundaries are mapped within 5 bp of their ground truth positions in the reference genome.

e) Read80%: the proportion of Read80% reads. A read is considered to be a Read80% read only if it meets two conditions: $N_T/N_G > 80\%$ and $N_T/N_P > 80\%$, where $N_G$ is the number of ground truth exons within the read, $N_P$ is the number of exons predicted by the alignment, and $N_T$ is the number of true positive exons. Herein, a predicted exon is considered to be a true positive exon only if there is a ground truth exon in the read and the corresponding boundaries of the predicted exon and the ground truth exon are within 5 bp.

f) Read100%: the proportion of Read100% reads. A read is considered to be a Read100% read only if it meets two conditions: $N_T/N_G = 100\%$ and $N_T/N_P = 100\%$. It is worth noting that a Read100% read indicates that the read has a highly correct full-length alignment.

### Supplementary Notes

#### 1. Genome indexing

The reference genome is organized and indexed using RdBG-index approach, which was initially developed in deBGA [4] for short DNA-seq read alignment. This data structure indexes the unitigs of the de Bruijn graph of the reference (also called as RdBG) instead of the reference itself. Give a reference, the RdBG-index is built in three steps: 1) a de Bruijn graph of the reference is built with a user-defined $k$ parameter (default value: 22); 2) all the unitigs of RdBG are extracted; 3) RdBG-index is built to index all the k-mers of all the unitigs, moreover, it also records the sequences of the unitigs and the genomic positions of all the copies of the unitigs in the reference. In details, RdBG-index has four major data structures as follows (a schematic illustration is in Supplementary Figure 12).

1) A linear table ($UT_{seq}$) records the sequences and the IDs (called as $U_{ID}$) of all the unitigs of RdBG. Each unitig has a unique $U_{ID}$.

2) A linear table ($UT_{kmer}$) records the lexicographically sorted list of all the k-mers of RdBG. For each of the k-mers, $UT_{kmer}$ records its token and its unique position in RdBG, i.e., the $U_{ID}$ and the offset of the unitig the k-mer being at.

3) A hash table ($UT_{k'}$) records the ranges of the k-mers having identical initial k' charaters in $UT_{kmer}$. $UT_{k'}$ is straightforwardly implemented by a surjection of k'-mers to RAM space, i.e., $UT_{k'}$ records all the $4^{k'}$ k'-mers in a lexicographical order in the RAM, and each item of $UT_{k'}$ records a specific range of the k-mers of RdBG whose k' long prefixes (also called as k'-prefixes) are same to the corresponding k'-mer. It is worth noting that for some items their ranges could be null, due to that there is no such k-mers in RdBG. This is an auxiliary data structure which helps to find short token matches between a read and the reference. As a sorted list, binary search is needed to locate a specific k-mer in $UT_{kmer}$, which is quite time consuming. $UT_{k'}$ helps to narrow down the range of k-mer retrieval and accelerate the query. The configuration of parameter *k'* is critical and deSALT sets *k'* = 14 (same as that of deBGA) which is a tradeoff between speed and memory use. Thus, the size of $UT_{k'}$ is 2GB, i.e., there are $4^{14}$ items and each of them stores one integer to record the range. Herein, the ranges are recorded in a difference approach, so that only one integer is needed to record in an item and a range can be derived from two neighboring items.

4) A linear table ($UT_{pos}$) records the genomic positions of the untigis, i.e., for each of the unitigs, the starting positions of all its copies in the reference are recorded. $UT_{pos}$ enables to convert the matches on unitigs to the matches on reference.

The size of RdBG-index depends on several factors such as the numbers and the lengths of unitigs, the number of the k-mers, the size of the k'-mer hash table and the total numbers of the copies of the unitigs. For human (GRCh38, version 94), mouse (GRCm38, version 94) and fruit fly (BDGP6, version 94) reference genomes, the sizes of the index are 35GB, 31GB and 3.5GB, respectively (with paramerter *k*=22 and *k'*=14).

It is worth noting that, RdBG-index not only supports to find k-mer matches, but also supports to find the matches from a *l*-mer (*l*<*k*) to all the k-mers whose *l* long prefixes (termed as *l*-prefix) are same to the *l*-mer. If $l \leq \mathrm{k}'$, all the k'-mers having the same *l*-prefix can be directly found through RAM address as $UT_{k'}$ is a surjection, and all the k-mers with the same *l*-prefix can be found via the ranges of recorded in the corresponding items of $UT_{k'}$. If $\mathrm{k}' < l \leq \mathrm{k}$, there is a unique k'-mer (item) in $UT_{k'}$ whose recording range points to a set of k-mers potentially having the same *l*-prefix. With this range, the k-mers having the required *l*-prefix can be found by a binary searching on $UT_{kmer}$.

In practice, no significant difference is observed in the results of the long RNA-seq read alignment with various *k* parameter settings (*k*=20 to 28, data not shown). This is mainly due to the fact that the seed length used in deSALT is much shorter than *k* bp and the structure of RdBG does not change much in such a range of *k*, so that tuning the *k* parameter has little effect on the generation of alignment skeletons. In this situation, deSALT uses the setting *k*=22 in all the benchmarks, while users can also change the setting on *k* if necessary.

#### 2. Alignment skeleton generation (first-pass alignment)

##### 2.1 Match-block generation

For a given read, deSALT extracts *l*-mers (default value: *l*=15) at every *m* bp (default value: *m*=5) as seeds and finds exact matches between the seeds and the unitigs of RdBG via the RdBG-index. It is also worthnoting that *l* is a user-defined parameter, which relates to the sensitivity of seeding. For noisy long reads, *l* should not be set too large otherwise there could be few long exact matches. To investigate its effects, we configured *l* parameter with various settings (*l* = 14, 15, 17 and 19), and implemented deSALT on the 18 low-coverage (4X) simulated datasets from human, mouse and fruit fly (with the 6 simulation models). The results are in Supplementary Table 12. Considering the alignment yields, we selected the default value of *l* as 15 and used this configuration in all the benchmarks, except for simulated ONT 1D reads, which *l*=14 was used due to the high sequencing errors.

Since *l*<*k*, a seed possibly has matches to multiple unitigs. deSALT extends each of the matches in both the upstream and downstream directions to generate a maximal exact match between the read and the corresponding unitig (termed a U-MEM). Moreover, for a certain seed, only its longest U-MEM is saved for further processing and others (if applicable) are filtered out. Furthermore, some seeds could have identical U-MEMs; such U-MEMs would be merged to remove redundant ones. After the processing, a set of non-redundant U-MEMs is produced, and each of them is termed as a specific 5-tuple, $M_i = (M_i^{RS}, M_i^{RE}, M_i^{U}, M_i^{US}, M_i^{UE}), i = 1, \dots, |M|$, where $M_i^{RS}$ and $M_i^{RE}$ respectively indicate the start and end positions of the match on the read, $M_i^{U}$ indicates the unitig ID of the match, $M_i^{US}$ and $M_i^{UE}$ respectively indicate the start and end positions of the match on unitig $M_i^{U}$, and $|M|$ indicates the total number of U-MEMs.

deSALT further merges co-linear U-MEMs on same unitigs as super U-MEMs (SU-MEMs). This is implemented by the following rule: for two U-MEMs $M_1$and $M_2$ on the same unitig $\mathrm{ux}$, $M_1 = (M_1^{RS}, M_1^{RE}, M_{ux}^{U}, M_1^{US}, M_1^{UE})$ and $M_2 = (M_2^{RS}, M_2^{RE}, M_{ux}^{U}, M_2^{US}, M_2^{UE})$, if $|(M_2^{RS} - M_1^{RS}) - (M_2^{US} - M_1^{US})| < T_{SU}$, $M_1^{R} < M_2^{R}$ and $M_1^{O} < M_2^{O}$, the two U-MEMs can be merged, where $T_{SU}$ is a threshold (default value: $T_{SU} = 20$). This operation takes the advantage of the characteristics of RdBG-index that if two seeds have co-linear hits on the same unipath, they can be seen as have two highly similar sets of hits on the various copies of the hit unitig. Theefore, they can be directly merged to reduce the redundancy of the seeds, without the need of checking all their hits on the reference sequences. This has beed justified in our previous work [4]. An SU-MEM, $SM_1 = (M_1^{RS}, M_2^{RE}, M_{ux}^{U}, M_1^{US}, M_2^{UE})$, a similar 5-tuple is generated after the merging. This definition is compatible with U-MEMs and easy to use for the merging operations between a U-MEM and an SU-MEM or two SU-MEMs.

In practice, deSALT separately processes the untigs with U-MEMs. For each of the unitigs, deSALT uses a greedy approach to iteratively check the U-MEMs from upstream to downstream and merges the U-MEMs meeting the conditions to produce SU-MEMs. It is also worth noting that there are possibly a proportion of U-MEMs (if not all) that cannot be merged and remain as individual U-MEMs. The SU-MEMs and individual U-MEMs are then mapped to the reference genome through the genomic positions of the unitigs (which are recorded by the RdBG-index in advance). It is worth noting that a unitig could have multiple copies in the reference genome, so that a SU-MEM/U-MEM could be split into multiple copies and mapped to all the corresponding genomic positions. Each of the mapped SU-MEMs/U-MEMs is termed a match block (MB) and defined as a 4-tuple $MB_i = (MB_i^{RS}, MB_i^{RE}, MB_i^{GS}, MB_i^{GE}), i = 1, \dots, |MB|$, where $MB_i^{RS}$ and $MB_i^{RE}$ respectively indicate the start and end positions of the match on the read, $MB_i^{GS}$ and $MB_i^{GE}$

respectively indicate the start and end positions of the match on the reference genome, and $|MB|$ indicates the total number of MBs.

**2.2 SDP-based alignment skeleton generation**

deSALT builds a direct acyclic graph (DAG) consisting of MBs. Each MB is a vertex, and there is an edge $MB_i \rightarrow MB_j$ if the placement of the two MBs, $MB_i$ and $MB_j$, meets the following condition:

$$MB_i^{RE} - \varepsilon < MB_j^{RS} \text{ and } MB_i^{GE} - \varepsilon < MB_j^{GS} < MB_i^{GE} + \delta,$$

where $\delta$ is a user-defined parameter depicting the maximum allowed intron length (default value: $\delta$ = 200,000 bp), and $\varepsilon$ is the maximum allowed overlap length between two MBs (default value: $\varepsilon$ = 5 bp). $\varepsilon$ is helpful for handling some small repeats around the boundaries of exons well. Such repeats could make some neighboring MBs have small overlaps, and $\varepsilon$ helps to correctly link such MBs and prevent false negatives.

A weight $\mathrm{s}(MB_i \rightarrow MB_j) = w(MB_i \rightarrow MB_j) - \mathrm{p}(MB_i \rightarrow MB_j)$ is assigned to each of the edges.

Here, $w(MB_j)$ is a weight score for depicting the match length of $MB_j$:

$$w(MB_i \rightarrow MB_j) = min(MB_j^{RE} - MB_j^{RS}, MB_j^{RE} - MB_i^{RE}, MB_j^{GE} - MB_i^{GE})$$

The last two elements in the brackets are applicable when two MBs have overlapping bases.

$\mathrm{p}(MB_i \rightarrow MB_j)$ is the intron length-based penalty score for preventing an ill-defined intron:

$$\mathrm{p}(MB_i \rightarrow MB_j) = \begin{cases} \alpha|MB_j^{GS} - MB_i^{GE}|/w(MB_i \rightarrow MB_j), & MB_j^{GS} - MB_i^{GE} \geq 0 \\ min(\alpha|MB_j^{GS} - MB_i^{GE}|/w(MB_i \rightarrow MB_j), \beta) & MB_j^{GS} - MB_i^{GE} < 0 \end{cases}$$

where $\alpha$ is a constant empirically set as 2 to scale the penalty score, and $\beta = \alpha T_{SU}/|MB_j^{RE} - MB_j^{RS}|$ is a constant penalty for an edge.

deSALT performs an SDP on the MBs by the following recursion equation to generate the alignment skeletons:

$$S_{AS}(MB_j) = \max\{S_{AS}(MB_i) + w(MB_i \rightarrow MB_j)\}, i \in Precusor\{MB_j\},$$

where $Precusor\{MB_j\}$ is the precursor set of $MB_j$ in which each of its elements $MB_i$ has an edge $MB_i \rightarrow MB_j$. Moreover, we define a vertex $MB_j$ as an isolated vertex if $Precusor\{MB_j\} = \emptyset$, and set $S_{AS}(MB_j) = MB_j^{GE} - MB_j^{GS}$.

With the recursion equation, deSALT assigns a score to all the vertices and also records the selected precursors for each of the vertices. The end vertex with the highest score is then selected, and deSALT builds the best alignment skeleton from it with backtracking operations. Here, end vertices indicate the vertices without successors.

Moreover, deSALT could output multiple alignment skeletons with similar scores. Specifically, deSALT also builds alignment skeletons for the end vertices meeting the following condition: $S_{AS}(MB_h)/S_{AS}(MB_{best}) > T_{AS}$, where $MB_{best}$ is the end vertex with the highest score, and $T_{AS}$ is a user-defined threshold (default value: $T_{AS} = 0.9$). For an $MB_h$, if the built skeleton has no shared vertex with the best skeleton, deSALT saves it as an alternative skeleton; otherwise, it would be discarded.

deSALT stores the best and the alternative skeletons in a temporary file for further processing. Each of the skeletons is recorded as a permutation of MBs:

$$AS_i = MB_{i\_1} \rightarrow MB_{i\_2} \rightarrow \cdots \rightarrow MB_{i\_|AS_i|}, \ \mathrm{i} = 1, \ldots, |AS|,$$

where each $MB_{i\_j}, \mathrm{j} = 1, \ldots, |AS_i|$ indicates an MB in the skeleton, $|AS_i|$ indicates the total number of MBs the skeleton has, and $|AS|$ indicates the total number of generated alignment skeletons.

#### 3. Exon inference

##### 3.1 Draft exon generation

deSALT integrates the alignment skeletons in an iterative approach to produce a set of draft exons. Herein, the exon set is formulated as $EX = \{EX_i, i = 1, \dots, |EX|\}$, where $|EX|$ is the total number of exons, and each $EX_i$ depicts an exon as a 3-tuple: $EX_i = (EX_i^{GS}, EX_i^{GE}, EX_i^{C})$, where $EX_i^{GS}$ and $EX_i^{GE}$ are respectively the start and end positions of the exon on the reference genome, and $EX_i^{C}$ depicts the number of alignment skeletons covered on the exons.

In practice, deSALT first extracts all the MBs of all the alignment skeletons and then sorts the MBs by their starting positions on the reference genome, i.e., $MB_{i\_j}^{GS}$, $\mathrm{i} = 1, \dots, |AS|$, $\mathrm{j} = 1, \dots, |AS_i|$. The initial exon set is constructed by the mapped MBs as follows:

$$EX = \{EX_p, p = 1, \dots, \textstyle\sum_{t=1}^{|AS|}|AS_t|\}, \text{ each } EX_p = (MB_{x\_y}^{GS}, MB_{x\_y}^{GE}, 1),$$

where the subscript x and y defines a specific MB, and all $EX_p$ are sorted by their upstream position.

deSALT then iteratively merges the $EX_p$ set to update $EX$ from upstream to downstream. For each $EX_p$ $(p > 1)$, deSALT checks if it meets the following condition:

$$EX_p^{GS} - \mu < EX_q^{GE} < EX_p^{GE},$$

where $EX_q$ is the exon closest to $EX_p$ upstream, $EX_q^{GE}$ is the end position of $EX_q$, and $\mu$ is the user-defined minimum intron length (default value: $\mu = 20$). If it does, deSALT merges the two exons to generate an updated exon $EX_{q'} = (EX_q^{GS}, EX_p^{GE}, EX_p^{C} + EX_q^{C})$; otherwise $EX_p$ and $EX_q$ would be not merged.

An updated exon set is generated after processing all the $EX_p$s, and deSALT filters the redundant and ill-defined exons. An exon $EX_p$ would be filtered out if it meets one of the two following conditions:

1) $EX_p$ is completely within another $EX_q$, i.e., $EX_p^{GS} > EX_q^{GS}$ and $EX_p^{GE} < EX_q^{GE}$;

2) $EX_p^{C} = 1$ and $EX_p^{GE} - EX_p^{GS} < l + 5$, where $l$ is the seed length used in the alignment generation step.

##### 3.2 Exon refinement

There could be some small differences between the boundaries of the draft exons ($EX_p^{GS}$s and $EX_p^{GE}$s) and the real splicing sites, mainly due to the effect of sequencing errors. deSALT predicts real splicing sites around the boundaries using a local sequence-based scoring system to refine the draft exons.

For a draft exon $EX_p$, deSALT selects two local regions, $EA_p = [EX_p^{GS} - 10, EX_p^{GS} + 10]$ and $ED_p = [EX_p^{GE} - 10, EX_p^{GE} + 10]$, and uses two scoring matrixes—Acceptor Scoring Matrix and Donor Scoring Matrix (see below), to respectively score the positions in $EA_p$ and $ED_p$. The two matrixes were initially proposed by Kim *et al*. to identity splice sites and translation start sites in human genomic sequences [5]. For a given position, the corresponding matrix is used as a mask to simultaneously score the position itself, upstream 5 bp (-5 to -1 bp), and downstream 4 bp (+1 to +4 bp) according to their nucleotides and to sum them up as the score of the position. This sum score can be seen as the possibility of the base permutation at the real acceptor or donor splicing sites. deSALT chooses the positions with the highest sum scores in $EA_p$ and $ED_p$ as the predicted acceptor and donor sites, respectively, and updates the exon $EX_p$.

After the prediction, a more accurate exon set is produced to implement the second-pass read alignment.

**Acceptor Scoring Matrix**

| nucleotide \ position | -5 | -4 | -3 | -2 | -1 | 0 | 1 | 2 | 3 | 4 |
|---|---|---|---|---|---|---|---|---|---|---|
| A | -201 | 115 | 61 | 2548 | -364 | 704 | -142 | -166 | -33 | 12 |
| C | 431 | 176 | 838 | -364 | -364 | -222 | -170 | -92 | 201 | 485 |
| G | -121 | 199 | -191 | -364 | 2548 | 346 | -117 | 973 | 157 | -91 |
| T | 350 | -107 | -169 | -364 | -364 | -271 | 1030 | -137 | 47 | 9 |

**Donor Scoring Matrix**

| nucleotide \ position | -5 | -4 | -3 | -2 | -1 | 0 | 1 | 2 | 3 | 4 |
|---|---|---|---|---|---|---|---|---|---|---|
| A | -92 | 132 | -97 | 922 | -107 | -344 | -344 | 355 | 939 | 222 |
| C | 168 | 252 | 273 | -208 | -288 | -344 | -344 | -236 | -177 | -206 |
| G | -114 | -110 | -167 | -132 | 1104 | 2406 | -344 | 543 | -160 | 1145 |
| T | 103 | -138 | 143 | -202 | -225 | -344 | 2406 | -223 | -207 | 185 |

#### 4. Refined alignment (second-pass alignment)

##### 4.1 Construction of a local spliced reference sequence (LSRS)

deSALT splits a read $\mathrm{R}$ into a series of unaligned parts according to its alignment skeleton $AS_R$ (assuming $\mathrm{R}$ has a unique skeleton), in which each part is represented as one of the substrings of $\mathrm{R}$:

$$\mathrm{R}\left[1, MB_{R\_1}^{RS}\right], \mathrm{R}\left[MB_{R\_1}^{RS}, MB_{R\_2}^{RS}\right], \ldots, \mathrm{R}\left[MB_{R\_j}^{RS}, MB_{R\_k}^{RS}\right], \ldots, \mathrm{R}\left[MB_{R\_|AS_R|}^{RE}, |R|\right],$$

where $MB_{R\_j}$ and $MB_{R\_k}$ are two neighboring MBs, and $|R|$ is the length of $\mathrm{R}$.

For each $\mathrm{R}\left[MB_{R\_j}^{RS}, MB_{R\_k}^{RS}\right]$, deSALT recognizes the exon meeting the following two conditions as the spanning exon: 1) the exon is between $MB_{R\_j}^{GS}$ and $MB_{R\_k}^{GS}$; 2) there is at least one short match between $\mathrm{R}\left[MB_{R\_j}^{RS}, MB_{R\_k}^{RS}\right]$ and the exon. Specifically, assuming that the exons containing $MB_{R\_j}$ and $MB_{R\_k}$ are respectively $EX_{R\_j}\left(EX_{R\_j}^{GS}, EX_{R\_j}^{GE}, EX_{R\_j}^{C}\right)$ and $EX_{R\_k}\left(EX_{R\_k}^{GS}, EX_{R\_k}^{GE}, EX_{R\_k}^{C}\right)$, deSALT selects the two local genomic regions $\left(MB_{R\_j}^{GS}, EX_{R\_j}^{GE}\right)$ and $\left(EX_{R\_k}^{GS}, MB_{R\_k}^{GS}\right)$ as the first and last spanning exons for $\mathrm{R}\left[MB_{R\_j}^{RS}, MB_{R\_k}^{RS}\right]$. Furthermore, for each of the exons within $EX_{R\_j}^{GE}$ and $EX_{R\_k}^{GS}$, if there is at least one *ls*-mer match between it and $\mathrm{R}\left[MB_{R\_j}^{RS}, MB_{R\_k}^{RS}\right]$, the exon is also recognized as a spanning exon. The *ls*-mer matches are quickly retrieved by a hash table-based index which indexes all the *ls*-mers of read $\mathrm{R}$. Here, *ls* is a user-defined parameter (default value: *ls*=8). deSALT then stiches all the spanning exons by their genomic positions to compose a sequence as the LSRS for $\mathrm{R}\left[MB_{R\_j}^{RS}, MB_{R\_k}^{RS}\right]$.

For $\mathrm{R}\left[1, MB_{R\_1}^{RS}\right]$, deSALT first recognizes the local genomic region $\left(EX_{R\_1}^{GS}, MB_{R\_1}^{GS}\right)$ as a spanning exon. Furthermore, deSALT checks five inferred exons upstream $EX_{R\_1}$ and selects the one(s) with at least one *ls*-mer match to $\mathrm{R}\left[1, MB_{R\_1}^{RS}\right]$ as the spanning exon(s) as well. The LSRS is then composed with all the spanning exons.

For $\mathrm{R}\left[MB_{R\_|AS_R|}^{RE}, |R|\right]$, the LSRS is composed in a similar way to the LSRS of $\mathrm{R}\left[1, MB_{R\_1}^{RS}\right]$, the difference being that deSALT checks five inferred exons downstream $EX_{R\_|AS_R|}$.

deSALT splits a read with multiple alignment skeletons multiple times with the various skeletons and separately produces a set of LSRSs for each of the skeletons.

##### 4.2 Alignment of the read part against the LSRS

deSALT employs a fast SIMD-based global alignment implementation [6] to align read parts against their LSRSs. For each of the alignments, the CIGAR string is investigated. If there are large deletions, deSALT updates the LSRS by removing the bases corresponding to the large deletion(s) and realigns the read part. Herein, most of the large deletions are due to the exons having alternative splicing sites. For such exons, their whole sequences are inferred by deSALT in the exon inference step and composed into LSRSs. However, the exon sequences could be partially included in some gene isoforms due to their alternative splicing sites, so the reads from those transcripts only have partial sequences. In this situation, there could be large deletions when the reads are aligned against the LSRSs involving the exons having alternative splicing sites. A schematic illustration is in Supplementary Figure 13.

For a certain read, deSALT integrates the alignment of all its read parts into a full-length read alignment if the read has a unique alignment skeleton. If more than one alignment skeleton is applicable, multiple full-length read alignments will be produced for a read. Full-length read alignments are scored by summing the scores of all the alignments of its read parts. For reads with multiple alignments, deSALT outputs the alignment with the highest score as the primary alignment and the other alignments as secondary alignments.

#### 5. Availability of datasets

##### 5.1 The availability of the human ONT datasets

http://s3.amazonaws.com/nanopore-human-wgs/rna/fastq/NA12878-cDNA-1D.pass.dedup.fastq

##### 5.2 The availability of the human ONT dRNA datasets

http://s3.amazonaws.com/nanopore-human-wgs/rna/fastq/NA12878-DirectRNA.pass.dedup.fastq.gz

##### 5.3 The availability of the mouse PacBio dataset (Accession Number: SRR6238555)

ftp://ftp.ncbi.nlm.nih.gov/sra/sra-instant/reads/ByRun/sra/SRR/SRR623/SRR6238555/SRR6238555.sra

##### 5.4 The availability of the simulated datasets

https://drive.google.com/drive/folders/1jk1ddv_QGozumnO_S_f-1IJlOehL3SyW?usp=sharing

These datasets are respectively generated by PBSim with the following command lines:

1) PacBio ROI-like datasets:

```
pbsim  Transcripome_File
    --data-type CCS \
    --model_qc model_qc_clr  \
    --length-mean 6000 \
    --length-min 100
    --difference-ratio 75:5:20 \
    --accuracy-mean 0.98 \
    --accuracy-min 0.8 \
    --depth 4/10/30
```

2) PacBio subread-like datasets:

```
pbsim  Transcripome_File
```

```
    --data-type CLR \
    --model_qc model_qc_clr  \
    --length-mean 7800 \
    --length-min 100
    --difference-ratio 1:12:2 \
    --accuracy-mean 0.85 \
    --accuracy-min 0.75 \
    --depth 4/10/30
```

3) ONT 2D (1D$^2$)-like datasets:

```
pbsim  Transcripome_File
    --model_qc model_qc_clr  \
    --length-mean 7800 \
    --length-min 100
    --difference-ratio 33:36:31 \
    --accuracy-mean 0.87 \
    --accuracy-min 0.8 \
    --depth 4/10/30
```

4) ONT1D-like datasets:

```
pbsim  Transcripome_File
    --model_qc model_qc_clr  \
    --length-mean 7800 \
    --length-min 100
    --difference-ratio 48:15:37 \
    --accuracy-mean 0.75 \
    --accuracy-min 0.7 \
    --depth 4/10/30
```

5) PBsim sampling-based simulation datasets:

```
pbsim Transcripome_File –depth 4/10/30 --sample-fastq sample.fastq
```

6) NanoSim simulation datasets:

```
python read_analysis.py genome -i sample.fa -rg reference.fa -a {minimap2, LAST}
python simulator.py genome -rg reference.fa -c model_prefix -n read_Count
```

#### 6. Availability of references

##### 6.1 The availability of the reference genomes

http://hgdownload.soe.ucsc.edu/goldenPath/hg38/bigZips/, human, GRCh38
http://hgdownload.soe.ucsc.edu/goldenPath/mm10/bigZips/, mouse, GRCm38
http://hgdownload.soe.ucsc.edu/goldenPath/dm6/bigZips/, fruit fly, DM6

##### 6.2 The availability of the gene annotations

ftp://ftp.ensembl.org/pub/release-94/gtf/homo_sapiens/, human, version 94
ftp://ftp.ensembl.org/pub/release-94/gtf/mus_musculus, mouse, version 94
ftp://ftp.ensembl.org/pub/release-94/gtf/drosophila_melanogaster, fruit fly, version 94

##### 6.3 The availability of the RdBG-index

https://drive.google.com/file/d/11E2j1X5jGqKNVtyNHsfPjqnu-fVBgyoi/view?usp=sharing, human GRCh38
https://drive.google.com/file/d/1tipOySE-_tmLI4jiy3GZdTj08RhJMXcl/view?usp=sharing, mouse, GRCm38
https://drive.google.com/file/d/1ZUg-Yc7oRQQjjJdh4_lJcDGomIVNfJBs/view?usp=sharing, fruit fly, DM6

#### 7. Implementation of benchmarking

All the benchmarks were implemented on a server with Intel Xeon E4280 CPU at 2.0GHZ and 1 Terabytes RAM, running Linux Ubuntu 16.04. We referred to a tutorial proposed by the PacBio team to configure the parameters of GMAP and Minimap2 to align the PacBio reads. The tutorial is available at:

https://github.com/Magdoll/cDNA_Cupcake/wiki/Best-practice-for-aligning-Iso-Seq-to-reference-genome:-minimap2,-deSALT,-GMAP,-STAR,-BLAT

The parameters of GMAP and Minimap2 were configured to align ONT reads by referring to a previous study:

https://github.com/nanopore-wgs-consortium/NA12878/blob/master/RNA.md

The alignment results on simulated and real datasets were evaluated by two in-house scripts, respectively, which are available at:

https://github.com/hitbc/deSALT

The command lines and versions of the aligners used in the benchmarking were as follows.

##### 7.1 Command lines of deSALT (version 1.4)

Genome indexing:

```
deSALT index Genome_File Genome_index_dir
```

Read alignment (default settings)

```
deSALT aln -o out.sam Genome_Index_Dir Fastq_File ref.fa
```

Read alignment (important parameters)

```
deSALT aln -l seeding-lmer -a local_hash_kmer -s seed_step -c min_chain_score -G GTF_info -B batch_size -x read_type -f temp-file-perfix -o out.sam Genome_Index_Dir Fastq_File
```

##### 7.2 Command lines of Minimap2 (version 2.17-r941)

Genome indexing:

```
minimap2 -ax splice -d ref.mmi ref.fa
```

Read alignment:

```
minimap2 -ax splice -t 24 ref.mmi Fastq_File > aln.sam
```

##### 7.3 Command lines of GMAP (version 2017-10-12)

Genome indexing:

```
gmap_build -d <genome> ref.fa
```

Read alignment:

```
gmap -D Genome_Index_Dir -d Genome_Database -f samse -n 0 -t 24 --cross-species --max-intronlength 200000 -z sense_force Fastq_File > out.sam 2> out.sam.log
```

**7.4 Command lines of Graphmap2 (version 0.3.0)**

Genome indexing:

```
graphmap2 align -I -r ref.fa
```

Read alignment:

```
graphmap2 align -x rnaseq -r ref.fa -d Fastq_File -o aln.sam
```
